# Temporal Complexity of the BOLD-Signal In Preterm Versus Term Infants

**DOI:** 10.1101/2023.12.08.570818

**Authors:** Allison Eve Mella, Tamara Vanderwal, Steven P Miller, Alexander Mark Weber

## Abstract

Preterm birth causes alterations in cerebral development. We calculated the Hurst exponent (H) - a measure of temporal complexity - from resting state functional magnetic resonance signal in preterm and term born infants. Anatomical, fMRI, and diffusion weighted imaging data from 716 neonates born between 23-43 weeks gestational age were obtained from the Developing Human Connectome Project. H was assessed in brain tissues and 13 resting state networks. H significantly increased with age, and earlier birth age contributed to lower H values. In most brain regions, H begins below 0.5 at preterm age and crosses 0.5 at term age. Motor and sensory networks demonstrated the greatest increase in H. Correlations between indirect measures of myelination and H were moderate. Overall, H appears to reflect developmental processes in the neonatal brain, as BOLD signal in the preterm infant transforms from anticorrelated to correlated but is reduced compared to term born infants.

## Introduction

Infants born before completing the full term of 37 weeks GA are defined as being preterm (***World Health Organization, 2023***). There are numerous biological, social and environmental factors that can potentially lead to preterm birth (***Vogel et al., 2018***). Preterm infants are more susceptible to dysfunctions in their sensory, motor and cognitive systems, and experience a higher prevalence of developmental impairments (***Ream and Lehwald, 2018*; *Gao et al., 2017*; *Luu et al., 2017***). This is due to interrupted brain maturation stages, and from early exposure to the harsh external environment (***Doi et al., 2022***).

Infants born preterm have important alterations in brain maturation that are reflected in more immature white matter microstructure, smaller regional brain volumes, and abnormal connectivity. These alterations in brain maturation may contribute to poorer neurodevelopmental outcomes in this cohort. MRI studies assessing functional properties of the preterm brain have also found lower functional connectivity and less distributed networks (i.e., more focal) (***Gao et al., 2017***). However, adult-like resting state networks (RSNs) have been observed in infants as early as 24 weeks GA, including visual, auditory, somatosensory, motor, parietal, cingulate, temporal, default mode, prefrontal, frontoparietal, executive, thalamic and cerebellar regions (***Doria et al., 2010*; *Fransson et al., 2007*; *Smyser et al., 2010***).

The blood oxygen level dependent (BOLD) signal in resting-state functional MRI (rs-fMRI) can be analysed beyond spatial connectivity measurements by looking at characteristics of the time-signal, such as determining its level of signal complexity (***Dong et al., 2018***). Temporal complexity of the BOLD signal is thought to be a product of brain criticality (***O’Byrne and Jerbi, 2022***). The brain and its neural systems are theorized to transition between critical states of order and disorder to maximize information processing (***Zimmern, 2020***). The critical brain hypothesis can describe how the brain dynamically processes information through phase transitions, as there must be equilibrium between order and disorder (***O’Byrne and Jerbi, 2022***). These critical phases are characterized by long range temporal correlations (LRTC) thought to reflect a system primed to optimally process information by balancing between stability and flexibility (***Cruz et al., 2021*; *Meisel et al., 2017*; *Moran et al., 2019***). Neurological disorders, impairments, or various brain states are associated with disruptions or deviations in the critical state (***Campbell and Weber, 2022*; *Cruz et al., 2021*; *Moran et al., 2019***).

The Hurst exponent (H) measures a signal’s fractal properties to quantify the long memory processes that occur within the signal itself (***Campbell and Weber, 2022*; *Dona et al., 2017***). Fractal qualities exist in the BOLD signal by way of self-repeating structures in the time series (***Campbell and Weber, 2022***). H indicates the level of temporal complexity of a signal, with values ranging between 0 and 1. H values closer to 1 are considered complex but more ordered, with greater long-term auto-correlations. H values closer to 0 are described as chaotic with long-term switching between high and low values in adjacent pairs (***Díaz M. and Córdova, 2022*; *Eke et al., 2002, 2000***). Thus, H values > 0.5 are said to have long-term memory and have persistent and auto-correlated behaviour, while H values equal to 0.5 describe random noise and contain little to no correlation in the signal, and H < 0.5 are said to have short-term memory and have anti-correlated behaviour (***Díaz M. and Córdova, 2022*; *Dong et al., 2018*; *Eke et al., 2002*; *Maxim et al., 2005***).

In preterm born infants, brain criticality has been studied using electroencephalogram (EEG) and fMRI, with various types of analyses such as H, power-law, spectral density and/or phase locking (***Zimmern, 2020***). In brief, these studies have shown that LRTC (***Hartley et al., 2012***) and EEG bursts are predictive of neurodevelopmental outcomes (***Iyer et al., 2015***). Using EEG, term born infants at 6 and 12 months were found to have LRTCs, demonstrating that these spatiotemporal dynamics are found in the first year of life (***Jannesari et al., 2020***). An fMRI-EEG study in term born infants found scale-free power-law exponents were network-specific (i.e. different for each network) (***Fransson et al., 2013***). To the best of our knowledge, however, no study to date has investigated the temporal complexity of BOLD signals in a preterm cohort compared to term born infants.

We assessed BOLD signal complexity by computing H values from rs-fMRI data from term and preterm infants. Using open science data from the Developing Human Connectome Project, we analysed rs-fMRI scans of 716 infants born between 23 and 42.71 weeks GA. We used median birth ages of 29, 35 and 41 weeks GA to illustrate cohorts of very preterm, moderately preterm, and term healthy controls, respectively. H was calculated in various brain tissues and 13 resting state networks. We also computed fractional anisotropy (FA) and radial diffusivity (RD) from diffusion weighted imaging (DWI) data, and tested for associations between H and these indirect measures of myelination. We hypothesised that (1) H would increase with scan age with differences across resting state networks, (2) preterm infants would have lower H values compared to the term born infants at term equivalent age (TEA), and (3) there would be associations between H and DTI measures suggesting that H development mirrors that of myelination.

## Results

### Participant Demographics

After excluding rs-fMRI scans with meanFD ≥ 0.05, 97 scans were removed from an initial set of 803. Of the initial study population of 716 distinct neonates, this resulted in a subset of 641 born between 23 and 42.71 weeks GA. Most of these exclusions were from the THC cohort (from 518 to 454 subjects). The resulting preterm group was composed of 187 infants including 81 infants with preterm only scans, 41 infants with term-equivalent only scans, and 65 infants with both preterm and term-equivalent aged scans. Moreover, the preterm cohort was subdivided according to the WHO thresholds into very preterm (VPT, infants born < 32 weeks GA) and moderately preterm (MPT, infants born ≤ 37 weeks GA but > 32 weeks GA). Within the VPT group there were 70 infants with preterm scans, of whom 37 also had TEA scans. Within the MPT group there were 76 infants with preterm scans, of whom 28 also had TEA scans. For the term healthy control group (THC) there were 518 infants included. All THC infants had one scan each (no repetitions). These participant details are shown in ***Table 1***.

**Table 1.** Participant demographics. Sex (male/female) group numbers and median ages and ranges are shown. VPT = very preterm, MPT = moderately preterm, THC = term healthy control, GA = gestational age, PMA = postmenstrual age.

| Cohort | n | Sex (M/F) | Birth Age / GA in Weeks (Range) | Scan Age / PMA in Weeks (Range) |
| --- | --- | --- | --- | --- |
| VPT | 82 | 43/39 | 29.07 (23.00 to 31.86) | Preterm, n = 70<br>32.64 (26.71 to 36.71)<br>TEA, n = 49<br>40.86 (38.14 to 44.57) |
| MPT | 105 | 58/47 | 34.86 (32.14 to 37.00) | Preterm, n = 76<br>35.57 (33.14 to 37.00)<br>TEA, n = 57<br>40.57 (37.14 to 45.14) |
| THC | 454 | 248/206 | 40.14 (37.14 to 42.71) | 41.29<br>(37.43 to 44.86) |

### H in All Subjects

H values in the whole brain are visually shown in ***Figure 1*** in a sample preterm born infant, the same infant at TEA, and a term born control. A corresponding sample Welch PSD is also displayed for a fitted frequency range of 0.08 to 0.16 Hz. When comparing the H values between males and females in the whole group - and in the VPT, MPT and THC groups separately - there were no significant differences.

**Figure 1.**
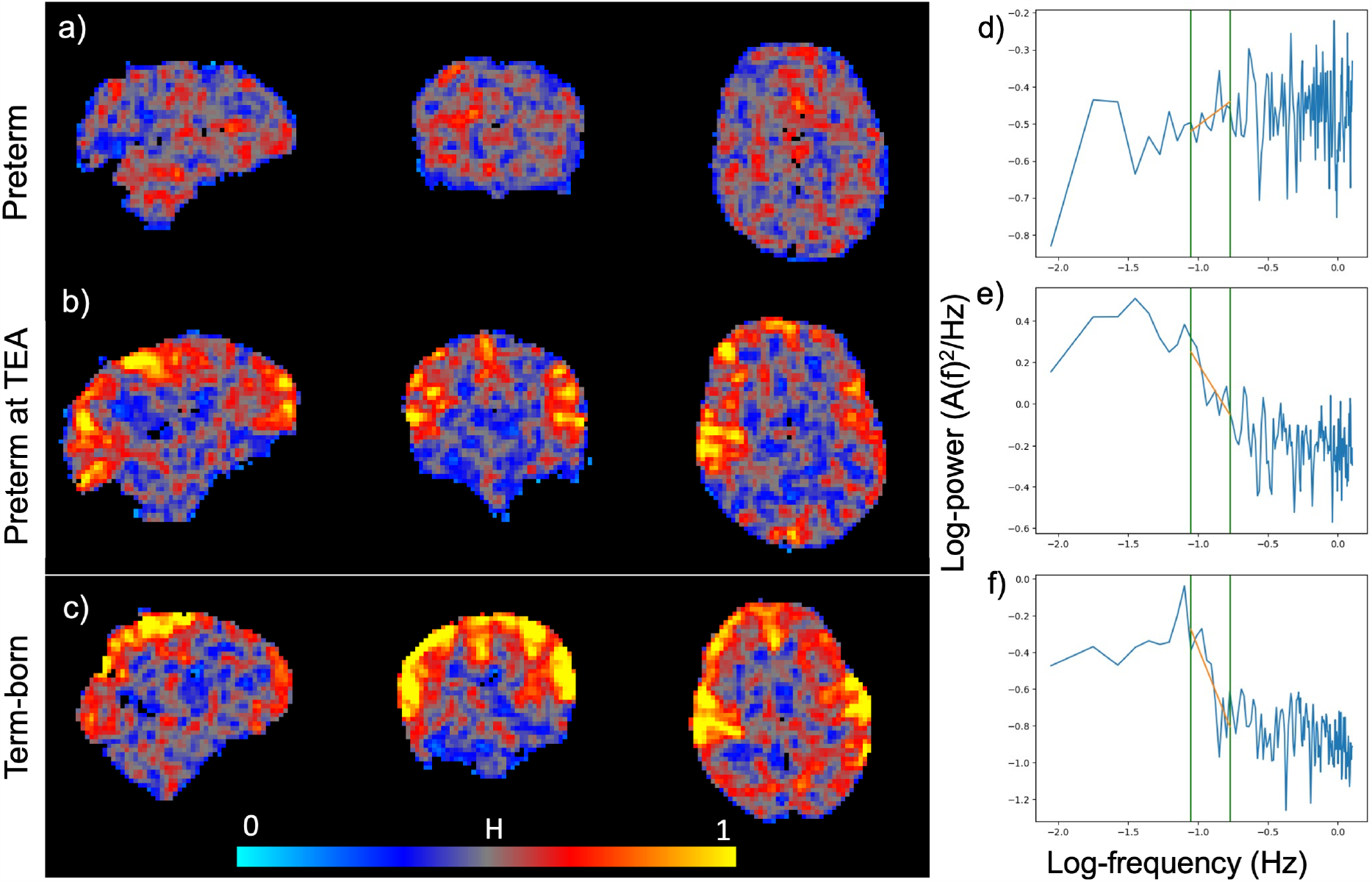
Higher H values in the whole brain with increasing gestation age in a single term-born infant and compared to a preterm infant both at 35 weeks PMA and at TEA, with their corresponding Welch PSDs. A value of 0 indicates low H and 1 indicates high H. Comparison of H values shown in a) a sample preterm infant scanned at 35 weeks PMA, b) the same infant at term of 41 weeks PMA and c) a single term control infant at 41 weeks PMA. Voxel location 32,33,25. Hurst maps were smoothed at 4mm for visualization purposes. Whole-brain averaged Welch’s power spectral densities shown (blue) with fitted frequency range of 0.08 to 0.16 (orange). d) to f) are from the same subjects/time-points as a) to c).

#### Grey Matter

H values increased with increasing scan age (PMA) when looking at the entire cohort in the GM (see ***Figure 2***.a). Similarly, H values increased with increasing birth age (GA) in the GM (see ***Figure 2***.b). Raw scan age and H were significantly and positively correlated in the GM with an r value of 0.633 (CI [0.586, 0.675], p<0.0001). Similarly, raw birth age and H in the GM were significantly positively correlated with an r value of 0.532 (CI [0.478, 0.584], p<0.0001).

**Figure 2.**
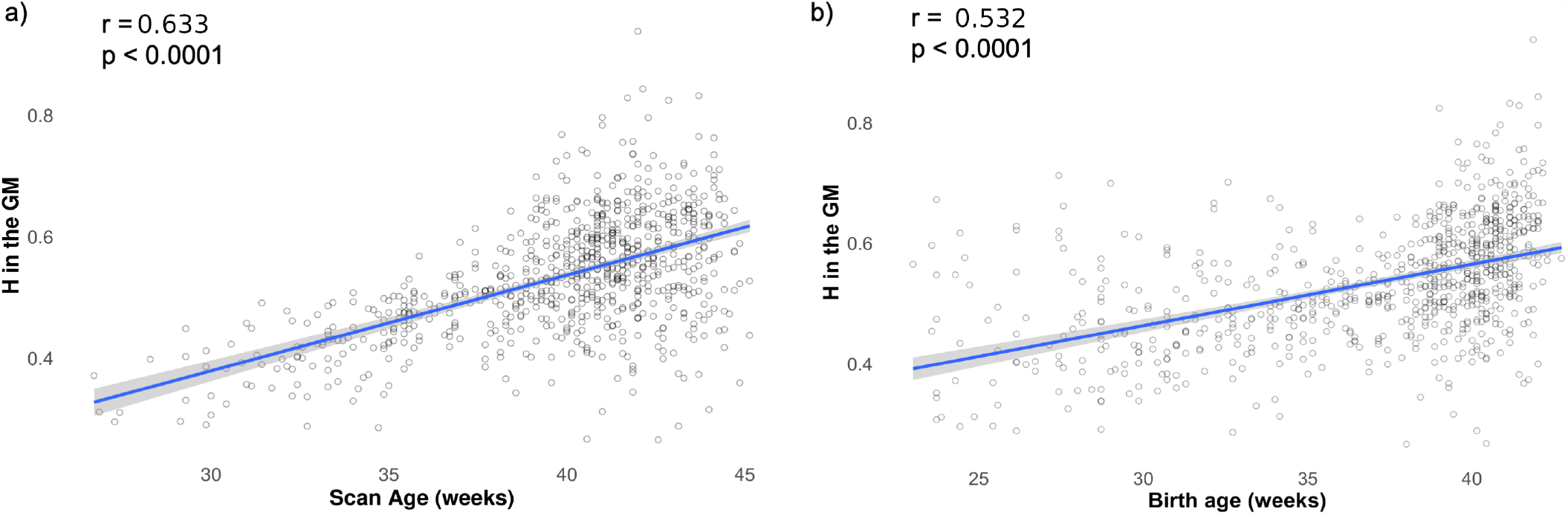
Whole group H in the GM with scan age and birth age in the full sample. Total H values in the grey matter from all subjects/time-points vs. a) scan age (PMA) in weeks; and b) birth age (GA) in weeks. Blue line represents the line of best fit for a linear regression, while the grey bands represent 95% confidence intervals.

The linear mixed effects model for H in the GM with scan age and birth age as fixed effects (***Equation 2***) was found to have a total substantial explanatory power with conditional r^2^ = 0.62, and marginal r^2^ = 0.43. Furthermore, within this model: the effects of scan age were found to be statistically significant and positive, birth age was found to be statistically significant and positive and *scanage* ∗ *birthage* was found to be statistically non-significant and negative. The statistical values are provided in ***Table 2***.

**Table 2.** Mixed linear effects model results. Results from the whole group in the grey matter, white matter and RSN. Calculated b values, SE (standard error), t-value and p-values from the linear mixed effect models. All estimates are significant excluding the grey matter scan age * birth age interaction term, and the white matter intercept.

| Parameters<br>(fixed effects) | b | SE | t-value | p-value |
| --- | --- | --- | --- | --- |
| Grey Matter, df = 700 |  |  |  |  |
| Intercept | -0.467 | 0.203 | -2.30 | 0.022 |
| Scan Age | 0.021 | 0.005 | 4.17 | < 0.001 |
| Birth Age | 0.013 | 0.006 | 2.12 | 0.035 |
| Scan age * Birth age | -2.22x10<sup>-4</sup> | 1.56x10<sup>-4</sup> | -1.42 | 0.155 |
| White Matter, df = 700 |  |  |  |  |
| Intercept | -0.190 | 0.136 | -1.39 | 0.164 |
| Scan Age | 0.014 | 0.003 | 4.01 | 0.001 |
| Birth Age | 0.013 | 0.004 | 3.04 | 0.002 |
| Scan age * Birth age | -2.43x10<sup>-4</sup> | 1.04x10<sup>-4</sup> | -2.33 | 0.020 |
| RSN (Motor Medial as Reference), df = 9124 |  |  |  |  |
| Intercept | -1.33 | 0.161 | -8.27 | < 0.001 |
| Scan Age | 0.047 | 0.004 | 11.83 | < 0.001 |
| Birth Age | 0.032 | 0.005 | 6.32 | < 0.001 |

When scan age was set at 41 weeks PMA, the slope of birth age and H in the GM was 0.00435. In other words, for every extra week of gestation, H measured at TEA increases by 0.00435 in the grey matter.

#### White Matter

The linear mixed effects model for H in the WM with scan age and birth age as fixed effects (***Equation 2***) was found to have a total substantial explanatory power with conditional r^2^ = 0.40, and marginal r^2^ = 0.31. Within this model the effects of scan age were statistically significant and positive, birth age was statistically significant and positive, *scanage* ∗ *birthage* was statistically significant and negative. The values are presented in ***Table 2***.

#### Resting State Networks

The linear mixed effects model for H with scan age, birth age, and RSN as fixed effects (***Equation 3***) was found to have a total substantial explanatory power with conditional r^2^ = 0.83, and marginal r^2^ = 0.49. Within this model, the effects of scan age and birth age were each statistically significant and positive and birth age was statistically significant and positive. These values are presented in ***Table 2***.

### H in Preterm Group Only

#### Tissues

H significantly increased (p<0.001) from preterm to TEA in the grey matter from 0.453 [0.446, 0.459] to 0.537 [0.531, 0.544], in the white matter from 0.433 [0.427, 0.439] to 0.462 [0.457, 0.468], and the combined RSN from 0.451 [0.444, 0.458] to 0.530 [0.523, 0.538]. H in the combined RSN and GM increased at similar rates and both at faster rates than the WM with slopes of 0.0128, 0.0136 and 0.00473, respectively (see ***Figure 3***.a). The GM and combined RSNs slopes cross H = 0.5 at approximately 39 weeks PMA. When comparing the VPT and MPT groups at 35 weeks PMA the VPT had significantly lower (p < 0.001) H values in the GM, WM and combined RSN than the MPT as shown in ***Figure 3***.c). In the GM the VPT had a mean H value of 0.448 [0.439,0.456], while the MPT had a mean value of 0.465 [0.4560.473]. In the WM the VPT and MPT had mean values 0.422 [0.414,0.429] and 0.448 [0.441,0.455], respectively. In the combined RSN, the VPT and MPT had mean values of 0.445 [0.437,0.454] and 0.463 [0.453,0.472], respectively. These values are provided in Supplementary Table 1.

**Figure 3.**
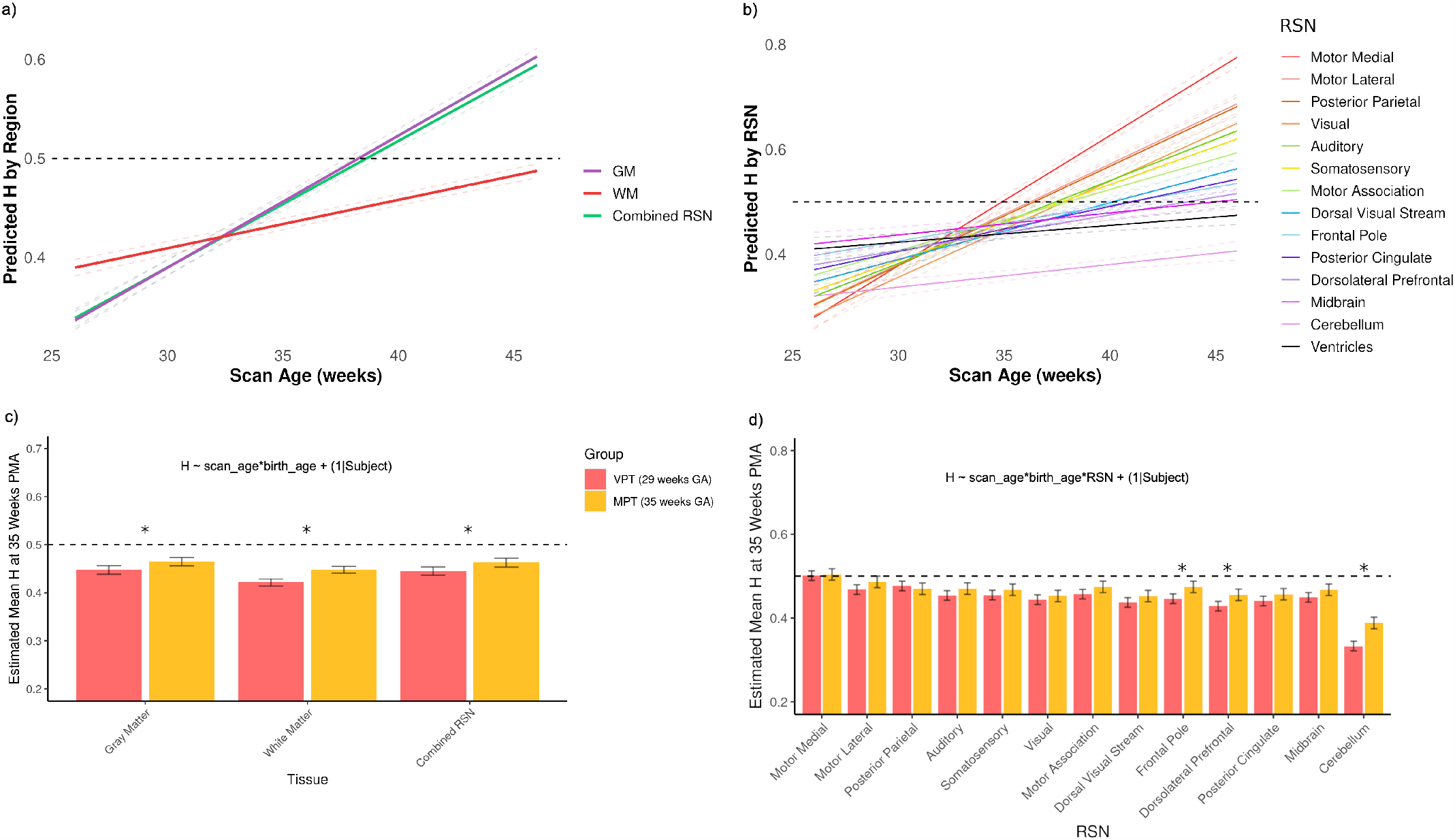
H analysis in the preterm infant group. Predicted H values in the a) tissues and b) RSNs from 25 to 45 weeks PMA. Comparison between the VPT group at 29 weeks GA and MPT at 35 weeks GA in the c) tissues and d) RSN at 35 weeks PMA. The groups are illustrated with an * to show significant comparisons (p < 0.05). The horizontal dashed line indicates H = 0.5.

#### Resting State Networks

The 13 RSNs (***Figure 4***) were analyzed to determine which network showed the greatest increase in H values from preterm to term equivalent age, using both single and longitudinal scans. All RSNs increased with scan age at different rates. The ventricles are also plotted with the RSNs to signify a ‘noise floor’, and the slope for each network surpassed the ventricle slope at different PMAs (see ***Figure 3***.b)) and ***Table 3***). The motor medial (0.0248), motor lateral (0.0191) and posterior parietal (0.0189) networks had the greatest increase in H with scan age (PMA). The dorsolateral prefrontal (0.0068), cerebellum (0.0043) and midbrain (0.0042) networks had the lowest increase in H with scan age (PMA). In ***Table 3***, the midbrain (26.0 weeks PMA), frontal pole (29.4 weeks PMA) and motor association (31.9 weeks PMA) intercept the ventricular slope the earliest, while the dorsal visual stream (34.3 weeks PMA), visual (34.4 weeks PMA) and the dorsolateral prefrontal (34.4 weeks PMA) networks intercept it the latest. In analyzing the preterm groups separately, the VPT had lower H in all RSNs as shown in Figure 4. d). This was significant in only 3 RSNs, however: the frontal pole, dorsolateral prefrontal, and cerebellum networks. The estimated marginal mean H values and p-values at 35 weeks PMA comparison for the VPT and MPT groups for all tissue and RSNs are in Supplementary Table 1.

**Table 3.** Resting state network slopes and ventricle intersection age (PMA). H slopes in each of the RSN in descending order and calculated PMA intersection with the ventricle slope. The ventricle slope is 0.0032.

| RSN | Slope | Ventricle Crossing (Weeks PMA) |
| --- | --- | --- |
| Motor Medial | 0.0248 | 32.1 |
| Motor Lateral | 0.0191 | 32.6 |
| Posterior Parietal | 0.0189 | 32.8 |
| Visual | 0.0183 | 34.4 |
| Auditory | 0.0158 | 33.2 |
| Somatosensory | 0.0145 | 33.1 |
| Motor Association | 0.0117 | 31.9 |
| Dorsal Visual Stream | 0.0108 | 34.3 |
| Posterior Cingulate | 0.0086 | 33.3 |
| Frontal Pole | 0.0069 | 29.4 |
| Dorsolateral Prefrontal | 0.0068 | 34.4 |
| Cerebellum | 0.0043 | N/A |
| Midbrain | 0.0042 | 26 |

**Figure 4.**
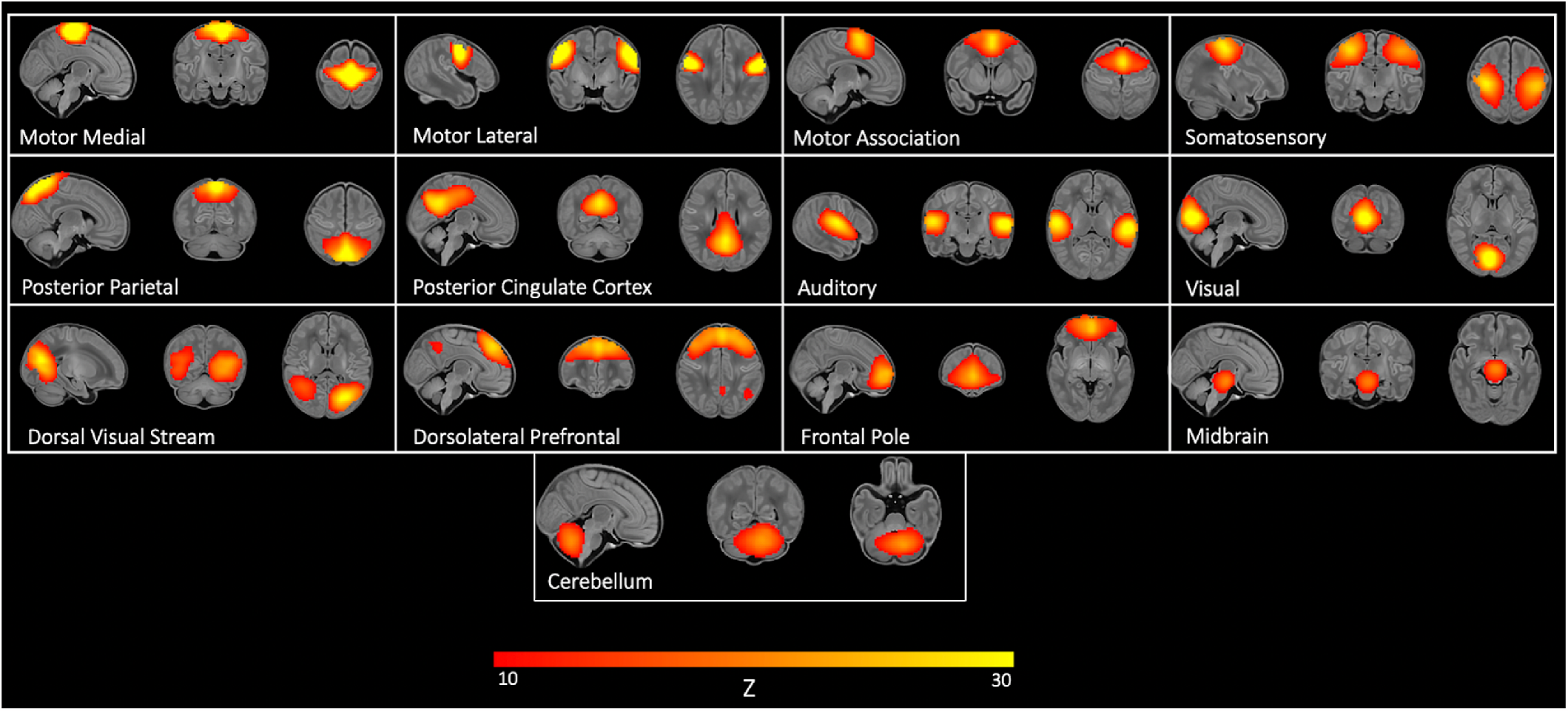
Resting state networks. The 13 identified RSNs from 30 ICA components. The networks are composed of motor, sensory, frontal, midbrain and cerebellum regions.

### H: Preterm vs Term

H values in the preterm groups of VPT and MPT at TEA were compared to the THC in the brain tissues and RSNs at 41 weeks PMA (***Figure 5***). Both preterm groups were lower than the THC and the VPT group was significantly lower than the MPT group in all regions. These comparisons were significant in the GM, WM, combined RSNs, and all individual RSN excluding the motor medial, posterior parietal, posterior cingulate and midbrain networks. The mean H values and p-values from the VPT versus MPT group comparison at 41 weeks PMA are provided in Supplementary Table 2).

**Figure 5.**
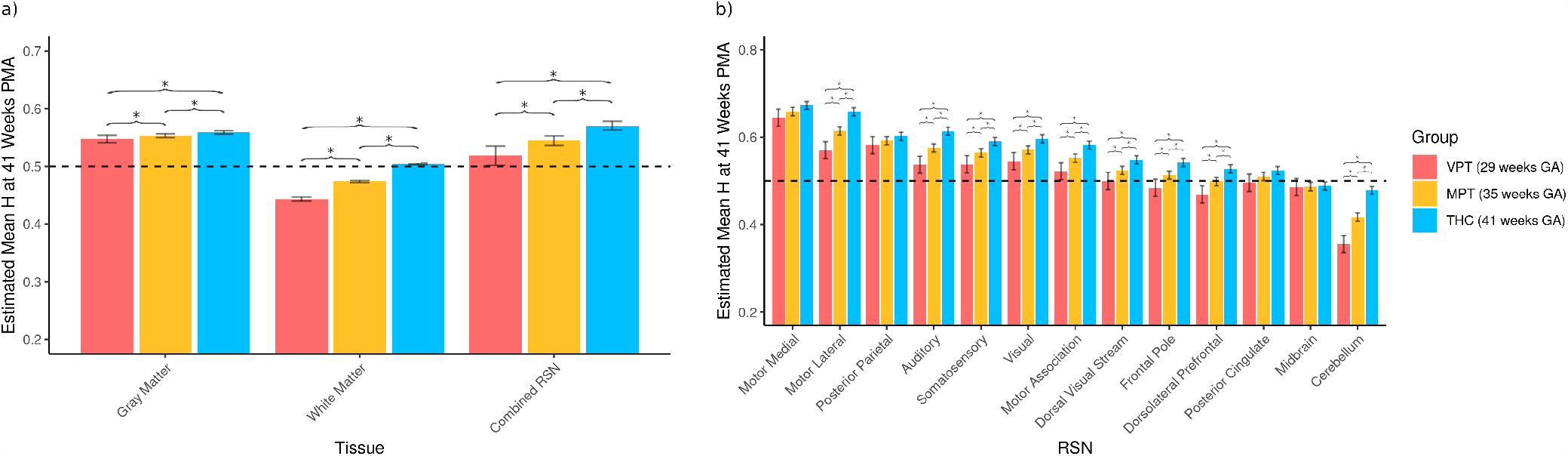
H analysis at term age comparison. H analysis in the preterm group at TEA compared to the term group in the a) tissues and b) RSNs. The mixed linear effects model in a) predicts H with scan age and birth age as fixed effects and in b) predicts H with scan age, birth age and the RSNs as fixed effects and both models account Subjects as random effects. The groups are illustrated with p-values (*) to show significant comparisons. The horizontal dashed line indicates H = 0.5.

### DTI Correlations

The calculated FA at 41 weeks PMA in the 35 weeks GA preterm group was 0.187 [0.183, 0.190] compared to at 41 weeks GA group FA significantly increased (p < 0.05) to 0.191 [0.185, 0.198]. RD at 41 weeks PMA comparison was not significantly different with mean values at 35 weeks GA of 0.00109 and at 41 weeks GA of 0.00107. FA and RD trends along with the white matter regions of interest are shown in the Supplementary Figure 2. DTI measures of FA and RD were correlated to H values in the preterm group (***Figure 6***). There was a significant positive correlation (p < 0.0001) between FA and H with an R value of 0.419 in the brain’s total white matter. RD and H had a significant negative correlation (p <0.0001) with an R value of -0.348. These associations were also present across the 11 white matter regions shown in the Supplementary Figure 3 and Supplementary Table 3. The motor lateral, somatosensory, and motor medial regions had the strongest correlations of FA and H with r values of 0.606, 0.589 and 0.552, respectively. R values for RD and H were the highest in the motor medial, motor lateral and somatosensory networks with values of -0.565, -0.499, and -0.475, respectively.

**Figure 6.**
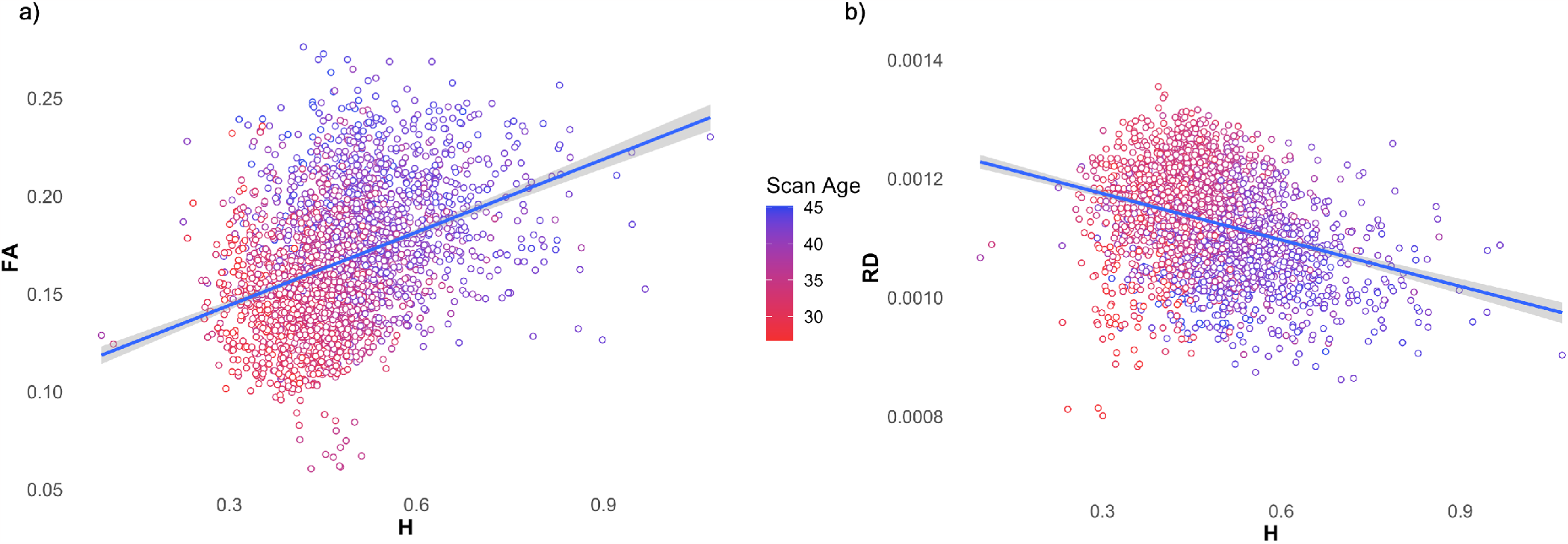
H and DTI correlations in the total white matter for all subjects and all scans. DTI measures of a) FA (fractional anisotropy) and b) RD (radial diffusivity) correlated to H values.

## Discussion

### Summary of Results

In this study we investigated whether infants born preterm demonstrated differences in BOLD-signal temporal complexity compared to term infants, using the Hurst exponent in various brain tissues and 13 resting state networks. First, we found that earlier preterm birth age was associated with lower H values at both preterm scans and later at scans conducted when the preterm infants reached term-equivalent ages. This was true in all brain tissues and networks, and was most pronounced in the very preterm group (VPT) which consistently had the lowest H values. Second, H values also demonstrated a network effect such that H in motor and sensory networks increased ahead of other networks. Third, we found moderate significant correlations between H and white matter estimates of FA (positive) and H and RD (negative). These results suggest that developmental changes in H are in some way associated with white matter development during early infancy.

### H and Myelination

We observed that H increases in motor and sensory RSN prior to other networks, a pattern that is consistent with functional connectivity studies in development across brain activity measures (***Cao et al., 2016*; *Doria et al., 2010*; *Eyre et al., 2021***). The earlier maturation of motor and sensory regions may be essential for the newborn’s survival in the external environment and/or to trigger the formation of other cortical networks via spontaneous neuronal activity and input from the sensory system (***Cao et al., 2016***). Auditory, visual and somatosensory systems become established at 33 to 35 weeks as evoked responses are maturing (***Kostović and Jovanov-Milošević, 2006***). An fMRI study with term infants by ***Fransson et al. (2013)*** found that power law exponents were greater in sensory, auditory and visual networks than other higher order networks. We posit that the higher H values and greater slopes for H in motor and sensory networks compared to the prefrontal and frontal networks at both preterm and term ages might reflect this same earlier functional maturation and pattern. The cerebellar RSN had low H values in all the groups compared to the other networks. Other studies suggest that the cerebellum demonstrates increased growth relative to the cerebral cortex, from 24 to 40 weeks; the maturation of the cerebellum is disrupted in preterm neonates due to a variety of conditions including hemorrhage, morphine exposure, and dysmaturation (***Volpe, 2009*; *Spoto et al., 2021***). This may explain why cerebellum H values are particularly low in the preterm group.

***Herzmann et al. (2019)*** assessed the functional connectivity in the cerebellum in comparing term to very preterm infants. They found decreased positive and negative functional connectivity correlation in the VPT group which they attributed in part to suppressed BOLD signal fluctuations that are associated with preterm birth (***Herzmann et al., 2019***). Here we show that cerebellar H values in the THC group in the current study were also low (below <0.5). If an analysis of H were conducted beyond the term ages, we might observe greater BOLD complexity in the cerebellum. Overall, we find that the cerebellum network in our analysis displayed the lowest H values; however, it is unclear what the contributing mechanism would be that differentiates this region from the others, and suggests future work is needed. In comparing how H develops in the RSNs to the ventricles, it is evident that all RSNs except the cerebellum eventually surpass the ventricles during preterm age (25-34 weeks PMA). H values could signify the functional organization that occurs during the 3rd trimester of pregnancy. In this current study we compared our RSN values to the ventricles to indicate the baseline level of H signalling. Since all but one RSN surpasses the ventricles, we can infer that the H measure of temporal complexity in the BOLD signal is not just fMRI signal noise.

We discovered modest but significant correlations between fractional anisotropy with H and radial diffusivity with H. These correlations were consistent in both the total white matter and individual white matter regions. Myelination is thought to commence at 20 to 28 weeks gestation, and to continue into the first year of life (***Smyser et al., 2011***). Previous studies have explored the relationship between myelination and functional connectivity. There is a theory that structural connectivity precedes - and is essential to - the formation of functional connectivity networks. For example, ***Vo Van et al***. (***2022***) showed that functional RSNs exist in more underdeveloped forms compared to more highly developed structural connectivity at the third trimester. Another study which utilised rs-fcMRI and DTI tractography in preterm infants at TEA showed that motor and visual functional connectivity was associated with WM maturation (***Weinstein et al., 2016***). However, it has also been theorized that activity dependent neuronal signals trigger myelination through sufficient blood and oxygen levels (***Yuen et al., 2014***). To date, it is clear that development of the brain’s structure and function are highly integrated, but they are not identical (***Smyser et al., 2011***). Furthermore, it is necessary to explore the relationship between BOLD complexity and myelination throughout the human lifespan.

### H Interpretations of Temporal Complexity in Infants

Our results provide a novel rs-fMRI method to investigate criticality and complexity of the BOLD signal in infants, and these findings build on prior work in EEG. For example, ***Hartley et al***. (***2012***) identified long range signalling with H in EEG in very preterm infants and concluded that the immature brain is capable of non-trivial electrical activity. Results here confirm the presence of longrange temporal correlations that are higher - and we infer more developed - in all our infants by term-age. We found short memory series in our preterm group (H < 0.5) and at term we found mostly long memory series (H > 0.5). Similar to our methods, ***Fransson et al. (2013)*** used Welch’s method for power spectral density estimation in a term born cohort with fMRI and speculated that the observed increase of power law exponent in their multiple sensory networks demonstrated neuronal processes with longer internal memory. Furthermore, an increase in power law exponents in a functional network is thought to signify more information flow and storage for longer time lengths (***Fransson et al., 2013***). Based on this, we postulate that the earlier the infant is born, the less internal signaling memory is permitted to promote information flow and storage given lower H values. With EEG and according to the critical brain hypothesis, it is apparent that there is a transition between the preterm infant asymmetric discontinuous brain activity to term age symmetrically continuous activity (***Meisel et al., 2017*; *Jannesari et al., 2020***). In another EEG study, periods of bursting neuronal activity became shorter and by 30 weeks PMA low voltage activity was apparent to demonstrate the presence of continuous activity (***Colonnese and Khazipov, 2012***). With the use of rs-fMRI and H measurements, the preterm infants here may exemplify the critical state transition from asynchronous/anti-correlated to synchronous/correlated processing. This type of BOLD signal processing information cannot be revealed through functional connectivity measurements. Temporal complexity (as measured by H) provides a unique way to investigate and characterize the BOLD signal.

### H as a biomarker for Neurodevelopment

The very and moderately born preterm groups were found to be significantly different at both preterm and term age comparisons. At term age, the THC group had higher H values than preterm infants at TEA. Furthermore, the VPT had the lowest H values overall. Structural abnormalities have been identified in infants born less than 32 weeks, with reduced overall brain volume (***Ream and Lehwald, 2018***), and we now show temporal complexity alterations in the functional signal in the preterm group. Clinically, infants born less than 32 weeks GA have higher NICU admission rates, need more respiratory support, and are at higher risk of neurodevelopmental delays than preterm infants born 34 to 37 weeks GA (***Smyrni et al., 2021*; *Chen et al., 2022***). When compared to term born children, VPT children will likely show lower scores in executive function measures (***Rogers et al., 2018***). These intellectual challenges in children born preterm can persist into adolescence and adulthood (***Ment et al., 2009***). Alternative information processing patterns and inferior synchronicity and criticality has been found in extremely preterm children (< 27 weeks GA) compared to term born children at 10 years, demonstrated with structural, diffusion and functional MRI (***Padilla et al., 2020***). This suggests that the extremely preterm group may have impaired adaptability to external and internal stimuli (***Padilla et al., 2020***). We found that the term born infants had decreased complexity with higher H values which suggest more organized and correlated BOLD signalling (***Díaz M. and Córdova, 2022*; *Dong et al., 2018*; *Wink et al., 2008***). The preterm groups had lower H values which could signify less organized, more chaotic, and anti-correlated signalling within their functional systems (***Díaz M. and Córdova, 2022*; *Dong et al., 2018*; *Wink et al., 2008***). When we assessed H in the GM, we discovered that the earlier the infant was born, H decreased by 0.00435 per week by TEA. This is in line with the clinical literature where there are more negative adverse outcomes for the infant with earlier gestational age (***Smyrni et al., 2021***). At this broad level of clinical acuity, our findings map onto the higher risk cohort. Longitudinal studies with both preterm and term infants in measuring H would better assess if these differences remain throughout the lifetime, and if preterm birth permanently impacts temporal complexity.

With regards to H trends in aging, it was found that in a cohort of 116 healthy adults aged 19 to 85 years, H was significantly correlated with age in the grey matter (***Dong et al., 2018***). This is the same general trend that we find when we correlate H with scan age and birth age for all subjects. In the current infant study, H values ranged from 0.33 to 0.65. Previous studies with H measurements in the brain have all been greater than 0.5, creating the belief that the brain only produced long memory signals of 0.5 < H < 1 (***Wink et al., 2008*; *Ciuciu et al., 2014*; *He, 2011*; *Churchill et al., 2016*; *Dong et al., 2018*; *Lei et al., 2013***). Our results in the preterm cohort contrast with these previous works. The low H values observed here could reflect a unique developmental stage in the brain that simply does not exist at any other point in life, and could include contributions from neural, vascular and other complex processes. Moreover, the use of non-traditional rs-fMRI analysis is recommended for H and other time series metrics.

### Comparison to Other H Studies

We assumed that H in the tissues would be greater than 0.5, as is seen in all adult studies to date and in a previous study of ours in preterm infants. However, we found that H was less than 0.5 at preterm age in all the regions we assessed. In our previous work, we evaluated a very preterm group of 98 (64 scanned at preterm age) infants at preterm and TEA with a sampling rate of 3s and only 100 volumes (***Drayne et al., 2022, 2023***). In that study, H was found to be between 0.67-0.70 in various RSNs. In the current study, we have replicated our previous findings that H increases with age and that certain RSNs show more rapid changes in H than others (***Drayne et al., 2022, 2023***). This previous work was the first study to assess BOLD temporal complexity with rs-fMRI in a preterm group. The present study used 641 subjects (187 preterm), and a sampling rate of 0.39s and 2,300 volumes. Due to the limited number of frequency values we could use to calculate H using Welch’s method, our previous study obtained the power-law slope using the entire frequency range. In the current study, we limited the calculation of the slope over the range of 0.08 to 0.16, which previously used in adults is believed to better represent brain signalling in the BOLD signal (***Dona et al., 2017***). Thus, we were able to build and expand on these previous findings by using a higher sampling rate and highly sampled rs-fMRI, a larger infant population and explore comparisons between preterm groups and to term born infants. For these reasons, we believe that this current study may better reflect the true H value in this population.

H measurements in the brain are method and calculation dependent. We chose to calculate the PSD using Welch’s method as it has been demonstrated to be more advantageous over alternative methods (***Rubin et al., 2013***) and was used in our previous studies (***Campbell and Weber, 2022*; *Drayne et al., 2022, 2023***). Based on the criteria of sensitivity to spikes, activation, and tissue type Welch’s outperformed other more commonly used methods such as detrended fluctuation, rescaled range and wavelet analysis (***Rubin et al., 2013***). ***Churchill et al***. (***2016***), suggested that this finding is dependent on the number of volumes in the fMRI scans and deduced that detrended fluctuation analysis is more robust for shorter time windows. However, in order to conduct fractal analysis, a higher sampling rate or approximately 500 volumes are proposed to improve accuracy of the fractal method analysis and to prevent incorrect estimates (***Campbell and Weber, 2022*; *Eke et al., 2002*; *Maxim et al., 2005***). This optimal higher sampling rate is noted to be a TR less than 1 second (***Dilharreguy et al., 2003***) and greater reliability and more detailed analysis are provided with scans of approximately 13 minutes (***Soares et al., 2016***). Our fMRI data included a long timewindow for a total of 2,300 volumes or 15 minutes scan length with a TR of 392 ms.

### Limitations

As stated in the Introduction, preterm birth is a complex, multifactorial phenomenon with both medical and sociocultural factors which were not controlled for in this study. There are a variety of metrics to measure brain complexity (***O’Byrne and Jerbi, 2022***), and thus it becomes more difficult to directly compare results between studies. The use of the term brain complexity and the significance of the results produced is heavily dependent on the methods of neuroimaging and computation of values utilized. We have tried to synthesize our findings with other work in the field, but these efforts are limited by the specificity and diversity of complexity related measures. Radiologic scores were given to each scan ranging from 1 (normal appearance) to 5 (incidental findings possible/likely significance for imaging analysis). There were 24 VPT, 13 MPT and 16 THC that had a radiologic score of 5, meaning that there is a possibility of brain injuries or lesions affecting analysis. We did not exclude these infants from analysis as the preterm group already was significantly smaller than the term born group; however, we found no significant difference when including DVARS in our linear mixed effect model. Moreover, our large group number differences between preterm to term born infants is a disadvantage. Additionally, the preterm group only had a small number of longitudinal scans available in the preterm group. All the infants from the Developing Human Connectome Project except 6 THC were scanned during natural sleep to minimize motion during scan acquisition. There are implications for the type of ‘resting’ activity that occurs during awake versus sleep states (***Eyre et al., 2021***).

### Future Directions and Conclusion

It would be advantageous in future work to have more equally sized groups when comparing the preterm born infants to term born controls. Additionally, it would be interesting to investigate whether white matter injuries or hemorrhages within the preterm groups would impact H values as the BOLD signal could be impacted due to brain injuries.

We investigated the Hurst exponent as a measure of temporal complexity of rs-fMRI signaling in comparing preterm born infants to term born infants. Our results indicate that H values below 0.5 are found in infants which has not yet been previously identified in the literature. We also showed somewhat curiously that H begins below 0.5 at preterm age and crosses 0.5 at term age in most regions. This could mean that the brain signalling develops from anti-correlated to correlated processing. In our interpretation, preterm birth interrupts development in the third trimester of gestation leading to disrupted functional signal organization at a foundational, BOLD-signal based level. The results of this study suggest that the Hurst exponent can be used to better understand and index brain development in the preterm and term newborns.

## Methods and Materials

### dHCP Dataset

Data for this study were obtained from the third release of the open source Developing Human Connectome Project, KCL-Imperial-Oxford Consortium (dHCP) (http://www.developingconnectome.org/project/) and ethics were approved by the UK National Research Ethics Authority (***Makropoulos et al., 2018***). Participants were recruited from St. Thomas Hospital and scanned at The Evelina Newborn Imaging Centre (***Edwards et al., 2022***). Preterm infants born < 32 weeks GA were defined as very preterm infants (VPT, n = 88) and those born ≤ 37 weeks GA were defined as moderately preterm infants (MPT, n = 110). Term healthy controls were defined as infants born > 37 weeks GA (THC, n = 518). It was reported that no babies in the preterm and full-term group had major brain injury.

### MRI Acquisition

Images were acquired on a 3T Philips Achieva system (Philips Medical Systems) with a dedicated neonatal 32 channel phased array head coil (***Hughes et al., 2017***). Structural, resting state fMRI (rs-fMRI) and diffusion tensor imaging (DTI) scans of the participants were analyzed for this study. Most of the participants were scanned during natural sleep after bottle feeding and swaddling, 6 were sedated with chloral hydrate. The total scan time was 63 minutes. Preterm infants were defined as infants born ≤ 37 weeks GA and term birth and term age equivalence was defined as > 37 weeks GA. The majority of infants were scanned once. A subset of preterm infants were scanned twice at preterm and TEA (n = 69). For further details see ***Edwards et al***. (***2022***).

#### Structural Parameters

Anatomical images were acquired with the following parameters: T2-weighted (spatial resolution = 0.8 mm isotropic; FOV = 145 × 145 × 108 mm; TR = 12 s; TE = 156 ms; SENSE factor 2.11 (axial) and 2.58 (sagittal)), and T1-weighted (spatial resolution = 0.8 mm isotropic; FOV = 145 × 122 × 100 mm; TR = 4,795 ms; TI = 1,740 ms; TE = 8.7 ms; SENSE factor 2.27 (axial) and 2.66 (sagittal)). The structural images used for this study were downloaded already preprocessed (***Fitzgibbon et al., 2020***).

#### rs-fMRI Parameters

Resting state fMRI (rs-fMRI) was acquired with high temporal resolution echo planar imaging using the following parameters: spatial resolution = 2.15 mm isotropic; TR = 392 ms; TE = 38 ms; multiband factor = 9; 2,300 volumes; and scan time = 15 minutes. In-plane acceleration or partial Fourier was not used. rs-fMRI images analyzed for this study were downloaded already preprocessed (***Fitzgibbon et al., 2020***), and registered into the volumetric brain template (***Schuh et al., 2018***). For further details regarding the fMRI acquisition, see ***Fitzgibbon et al***. (***2020***) and ***Edwards et al***. (***2022***).

#### DTI Parameters

Diffusion MRI was acquired with echo planar imaging using the following parameters: in-plane resolution = 1.5 × 1.5 mm; 3 mm slices with 1.5 overlap; TR = 3,800 ms; TE = 90 ms; multiband factor = 4x; SENSE factor = 1.2; and partial Fourier factor = 0.86. For each subject 20 b0s were acquired and 3 different b-value shells (b = 400 s/mm^2^: 64, b = 1,000 s/mm^2^: 88, b = 2,600 s/mm^2^: 128) in 4 phase encoding directions (AP, PA, RL, and LR). The dMRI images analysed for this study were downloaded already preprocessed ***Bastiani et al***. (***2019***). More details can be found at ***Bastiani et al***. (***2019***) and ***Edwards et al***. (***2022***).

### MRI Processing

#### fMRI Processing

Resting state fMRI scans underwent further processing through applying the dHCP provided FSL- FIX components to regress out noise and motion components (***Griffanti et al., 2014*; *Salimi-Khorshidi et al., 2014***). Briefly, FSL-FIX is an automated independent component analysis-based denoising technique (ICA-FIX). ICA-FIX auto-identifies ICA components carrying artifacts/noise from a manually pre-trained component set. For additional details, see ***Fitzgibbon et al***. (***2020***). Mean framewise displacement (meanFD) was calculated for each subject using FD values provided by dHCP (found in “sub-SUBID_ses-SESSID_motion.tsv” file). The meanFD per scan was computed first by translating rotational displacements to translational displacements by projection to the surface of a 50 mm radius sphere (***Power et al., 2012***). The average of the sum of the absolute values of the differentiated realignment estimates (by backwards differences) at every time point were then calculated to give meanFD (***Power et al., 2012***). Subjects with meanFD > 0.5 were excluded from further analysis.

#### Group-level Resting State Networks

A subgroup of 52 scans was selected by stratified random sampling for ICA analysis to manually identify group level resting state networks (RSN). rs-fMRI scans were first registered to a 40-week template (***Schuh, 2017*; *Schuh et al., 2018***), were highpass filtered for 100s/0.01Hz to remove nonneuronal noise, and spatially smoothed at 8mm full width at half maximum using a Gaussian kernel. Group ICA was then performed using FSL’s MELODIC with 30 components. After visual inspection by AMW and AEM, and comparison to other published studies with preterm infants, 13 RSNs were identified. These included six lateralized networks that were combined to create three bilateral networks. The 13 networks identified were the motor medial, motor lateral, motor association, somatosensory, auditory, posterior parietal, posterior cingulate cortex, visual, dorsal visual stream, frontal pole, dorsolateral prefrontal, midbrain and cerebellum. These RSNs were used to create masks for averaging the Hurst exponent and DTI metrics (in combination with white matter segmentation masks). These steps are shown in ***Figure 7***.

**Figure 7.**
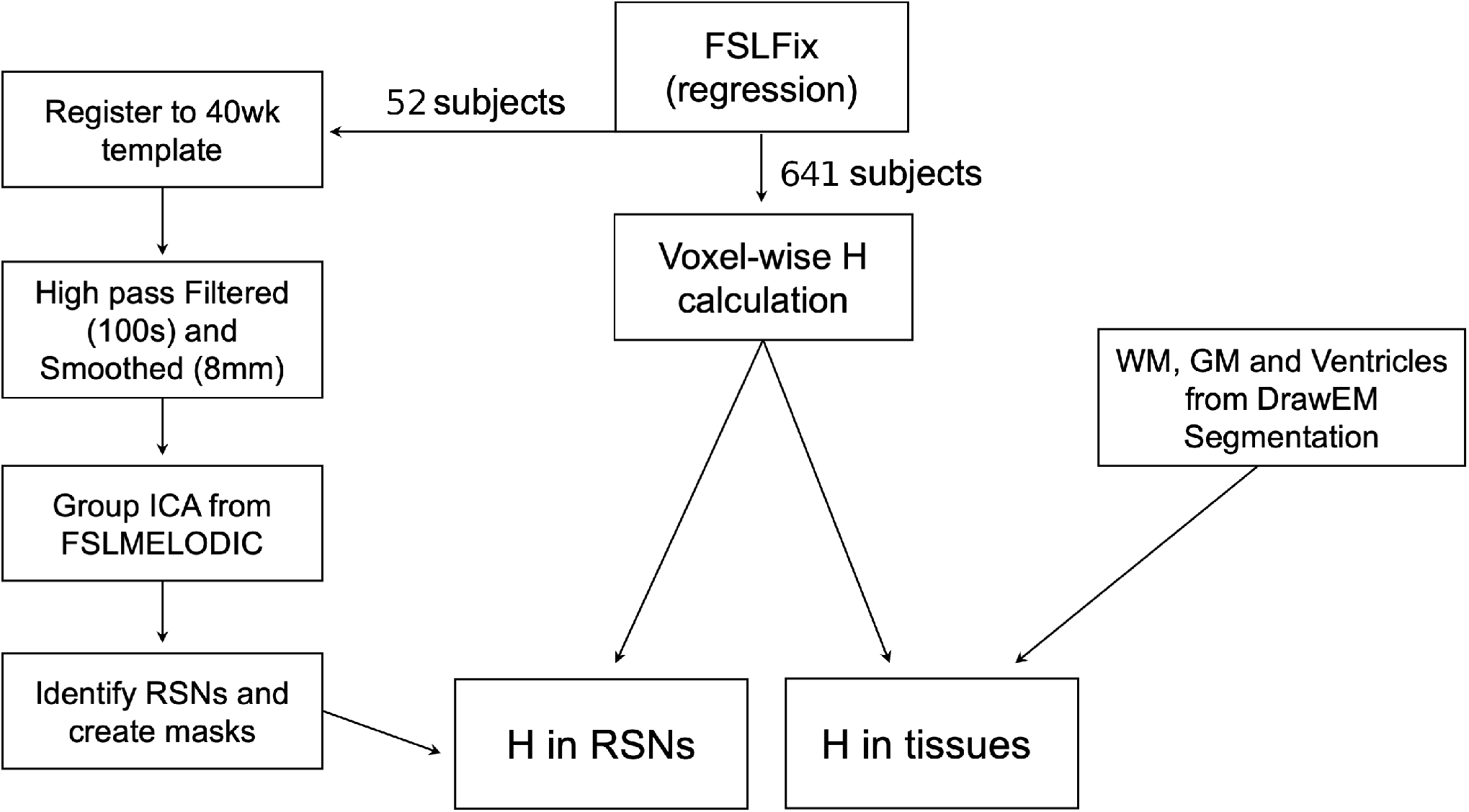
Hurst exponent calculation in the regions of interest. Flowchart for calculating H in the RSNs and grey and white matter from ICA analysis of 30 components and DrawEm.

#### Hurst Exponent Calculation

fMRI BOLD complexity was determined with H measurements that were calculated from the power spectrum density (PSD) of the BOLD signal. Our methods to calculate H were replicated from our previous work (***Drayne et al., 2022, 2023***). The PSD describes the power of the fMRI signal and can be calculated using various methods. Welch’s method to quantify the PSD has been shown to be advantageous over other methods in having higher sensitivity to activation and to the type of tissue (***Rubin et al., 2013***). The PSD was estimated by using the ‘welch’ command from Python’s ‘Scipy.Signal’ library (***Virtanen et al., 2020***). The data were divided into successive eight windows with 50% overlap, and averaged (***Welch, 1967***). The slope of the PSD was determined between the frequency range of 0.08 and 0.16 (***Dona et al., 2017***). These parameters are mirrored from ***Rubin et al. (2013)***. After determining the PSD, H was calculated from ***Equation 1*** where *β* is the negative of the slope from the log-power log-frequency plot of a power spectral density.

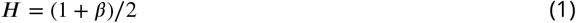

*β* here is assumed to be a signal in the class of fractional Gaussian noise (fGn) (as opposed to fractional Brownian motion (fBm)). ***Bullmore et al. (2004)*** described that human fMRI signals can sometimes begin as fluctuating fBm signals with increasing variance, but are then transformed to stationary fGn signals with constant variance when the fMRI data is motion corrected. fBm signals cannot be completely removed, however, and as first conceptualized by ***Eke et al. (2000)***, signals are hereby assumed to be fGn between 0 < H < 1 and fBm between 1 < H < 2, also known as an ‘extended’ Hurst. H was first calculated in all voxels in the brain, then averaged within individual RSNs through the previously created RSN masks. RSNs masks were combined to calculate a total RSN H value. H in the ventricles grey and white matter were calculated from segmentations obtained by the DrawEm method (***Makropoulos et al., 2014***). These steps are shown in ***Figure 7***.

#### DTI Analysis

Diffusion measures of fractional anisotropy (FA) and radial diffusivity (RD) were calculated in the brain’s white matter for the preterm group using the preterm age (≤ 37 weeks GA) scan. The RSNs masks created from ICA analysis were used as regions of interest (ROIs) to determine diffusion measurements in the white matter. The cerebellum and midbrain RSN masks were excluded after the DrawEm segmentation failed to segment out white and grey matter in these regions (***Makropoulos et al., 2014***). This left 11 white matter regions for analysis: motor medial, motor lateral, motor association, somatosensory, auditory, posterior parietal, posterior cingulate cortex, visual, dorsal visual stream, frontal pole, and dorsolateral prefrontal. These WM region masks were also combined to provide a total white brain measure.

### Statistics

Statistical analyses were completed using R and RStudio (v4.0) (R Core Team, 2022; RStudio Team). Birth age was used as a continuous variable to analyse the VPT, MPT and THC as groups were sorted by median birth age: VPT infants were analysed at 29 weeks GA, MPT infants at 35 weeks GA and THC at 41 weeks GA. Mixed linear effects models (shown in ***Equation 2*** and ***Equation 3***) were created using subjects as random effects to account for differences in within- and between-subject variance given that some preterm born infants were scanned twice (at preterm age and term, n = 85) and others were scanned only once (at preterm age, n = 69 or term, n = 44). Scan age, birth age and RSNs were set as fixed effects in the models.

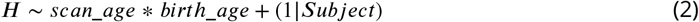

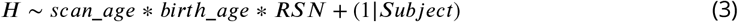

Standardised parameters were obtained by fitting the model on a standardised version of the dataset. 95% Confidence Intervals (CIs) and p-values were computed using a Wald t-distribution approximation. Marginal means and trends were calculated and Holm’s method was used for pvalue adjustment. H values in the ventricles, GM, WM, combined RSN and individual RSNs were assessed by comparing the preterm group to term age equivalent (TEA) and preterm group at TEA to THC. This was done by estimating H from median postmenstrual age scans. The preterm group (n = 198) was analysed by assessing longitudinal H values in the brain tissues and RSN. The preterm scans at 35 weeks PMA were compared to the TEA scans at 41 weeks PMA using estimated marginal means (‘emmeans’) (***Russell V. Lenth, 2022***). Preterm groups of the VPT and MPT were compared with each other at the scan age of 35 weeks PMA. RSN slopes were calculated using estimated marginal trends (‘emtrends’) (***Russell V. Lenth, 2022***) to determine which has the greatest increase with age in the entire preterm group. The VPT and MPT groups at term age were each compared to the THC group at the scan age of 41 weeks PMA in the brain tissues and RSNs. The longitudinal scans which contained a subset of preterm infants that were scanned twice to provide longitudinal data (n = 69) at preterm and TEA were also analysed to confirm whether H values trends were consistent (Supplementary Data). Sex differences were assessed between biological males and females in the brain tissues, RSNs and sexes in the infant groups. Overall means in the sexes were calculated and compared. In the preterm group, FA and RD were assessed longitudinally in both the total WM and individual regions. White matter microstructural associations to H in the preterm group were assessed by computing Pearson correlations between FA and H and between RD and H.

## Supporting information

All supplementary data

## Acknowledgements

We would like to thank the Developing Human Connectome Project, KCL-Imperial-Oxford Consortium for the use of their data funded by the European Research Council under the European Union Seventh Framework Programme (FP/2007-2013) / ERC Grant Agreement no. [319456]. We are grateful to the families who generously supported this trial.

## References

Bastiani M, Andersson JLR, Cordero-Grande L, Murgasova M, Hutter J, Price AN, Makropoulos A, Fitzgibbon SP, Hughes E, Rueckert D, Victor S, Rutherford M, Edwards AD, Smith SM, Tournier JD, Hajnal JV, Jbabdi S, Sotiropoulos SN. Automated Processing Pipeline for Neonatal Diffusion MRI in the Developing Human Connectome Project. NeuroImage. 2019 Jan; 185:750–763. doi: 10.1016/j.neuroimage.2018.05.064.

Bullmore E, Fadili J, Maxim V, Şendur L, Whitcher B, Suckling J, Brammer M, Breakspear M. Wavelets and Functional Magnetic Resonance Imaging of the Human Brain. NeuroImage. 2004 Jan; 23:S234–S249. doi: 10.1016/j.neuroimage.2004.07.012.

Campbell O, Vanderwal T, Weber AM. Fractal-Based Analysis of fMRI BOLD Signal During Naturalistic Viewing Conditions. Frontiers in Physiology. 2021; 12:809943. doi: 10.3389/fphys.2021.809943.

Campbell OL, Weber AM. Monofractal Analysis of Functional Magnetic Resonance Imaging: An Introductory Review. Human Brain Mapping. 2022 Jun; 43(8):2693–2706. doi: 10.1002/hbm.25801.

Cao M, He Y, Dai Z, Liao X, Jeon T, Ouyang M, Chalak L, Bi Y, Rollins N, Dong Q, Huang H. Early Development of Functional Network Segregation Revealed by Connectomic Analysis of the Preterm Human Brain. Cerebral Cortex. 2016 Mar; p. bhw038. doi: 10.1093/cercor/bhw038.

Chen Z, Xiong C, Liu H, Duan J, Kang C, Yao C, Chen K, Chen Y, Liu Y, Liu M, Zhou A. Impact of Early Term and Late Preterm Birth on Infants’ Neurodevelopment: Evidence from a Cohort Study in Wuhan, China. BMC Pediatrics. 2022 Dec; 22(1):251. doi: 10.1186/s12887-022-03312-3.

Churchill NW, Spring R, Grady C, Cimprich B, Askren MK, Reuter-Lorenz PA, Jung MS, Peltier S, Strother SC, Berman MG. The Suppression of Scale-Free fMRI Brain Dynamics across Three Different Sources of Effort: Aging, Task Novelty and Task Difficulty. Scientific Reports. 2016 Aug; 6(1):30895. doi: 10.1038/srep30895.

Ciuciu P, Abry P, He BJ. Interplay between Functional Connectivity and Scale-Free Dynamics in Intrinsic fMRI Networks. NeuroImage. 2014 Jul; 95:248–263. doi: 10.1016/j.neuroimage.2014.03.047.

Colonnese M, Khazipov R. Spontaneous Activity in Developing Sensory Circuits: Implications for Resting State fMRI. NeuroImage. 2012 Oct; 62(4):2212–2221. doi: 10.1016/j.neuroimage.2012.02.046.

Cruz G, Grent-’t-Jong T, Krishnadas R, Palva JM, Palva S, Uhlhaas PJ. Long Range Temporal Correlations (LRTCs) in MEG-data during Emerging Psychosis: Relationship to Symptoms, Medication-Status and Clinical Trajectory. NeuroImage: Clinical. 2021; 31:102722. doi: 10.1016/j.nicl.2021.102722.

Díaz MHA, Córdova F. On the Meaning of Hurst Entropy Applied to EEG Data Series. Procedia Computer Science. 2022; 199:1385–1392. doi: 10.1016/j.procs.2022.01.175.

Dilharreguy B, Jones RA, Moonen CTW. Influence of fMRI Data Sampling on the Temporal Characterization of the Hemodynamic Response. NeuroImage. 2003 Aug; 19(4):1820–1828. doi: 10.1016/S1053-8119(03)00289-1.

Dimitrova R, Pietsch M, Ciarrusta J, Fitzgibbon SP, Williams LZJ, Christiaens D, Cordero-Grande L, Batalle D, Makropoulos A, Schuh A, Price AN, Hutter J, Teixeira RP, Hughes E, Chew A, Falconer S, Carney O, Egloff A, Tournier JD, McAlonan G, et al. Preterm Birth Alters the Development of Cortical Microstructure and Morphology at Term-Equivalent Age. NeuroImage. 2021 Nov; 243:118488. doi: 10.1016/j.neuroimage.2021.118488.

Doi M, Usui N, Shimada S. Prenatal Environment and Neurodevelopmental Disorders. Frontiers in Endocrinology. 2022 Mar; 13:860110. doi: 10.3389/fendo.2022.860110.

Dona O, Hall GB, Noseworthy MD. Temporal Fractal Analysis of the Rs-BOLD Signal Identifies Brain Abnormalities in Autism Spectrum Disorder. PLOS ONE. 2017 Dec; 12(12):e0190081. doi: 10.1371/journal.pone.0190081.

Dong J, Jing B, Ma X, Liu H, Mo X, Li H. Hurst Exponent Analysis of Resting-State fMRI Signal Complexity across the Adult Lifespan. Frontiers in Neuroscience. 2018 Feb; 12:34. doi: 10.3389/fnins.2018.00034.

Doria V, Beckmann CF, Arichi T, Merchant N, Groppo M, Turkheimer FE, Counsell SJ, Murgasova M, Aljabar P, Nunes RG, Larkman DJ, Rees G, Edwards AD. Emergence of Resting State Networks in the Preterm Human Brain. Proceedings of the National Academy of Sciences. 2010 Nov; 107(46):20015–20020. doi: 10.1073/pnas.1007921107.

Drayne JP, Campbell O, Chau C, Miller S, Grunau R, Weber AM. Fractal Analysis of the BOLD Signal in Preterm Infants Scanned Shortly After Birth and at Term-Equivalent Age. In: International Society of Magnetic Resonance in Medicine London, UK; 2022..

Drayne JP, Mella AE, McLean MA, Ufkes S, Chau V, Guo T, Branson HM, Kelly E, Miller SP, Grunau RE, Weber AM. Long-Range Temporal Correlation Development in Resting-State fMRI Signal in Preterm Infants: Scanned Shortly After Birth and at Term-Equivalent Age. Developmental Cognitive Neuroscience. 2023 Nov; Under Review.

Edwards AD, Rueckert D, Smith SM, Abo Seada S, Alansary A, Almalbis J, Allsop J, Andersson J, Arichi T, Arulkumaran S, Bastiani M, Batalle D, Baxter L, Bozek J, Braithwaite E, Brandon J, Carney O, Chew A, Christiaens D, Chung R, et al. The Developing Human Connectome Project Neonatal Data Release. Frontiers in Neuroscience. 2022 May; 16:886772. doi: 10.3389/fnins.2022.886772.

Eke A, Hermán P, Bassingthwaighte J, Raymond G, Percival D, Cannon M, Balla I, Ikrényi C. Physiological Time Series: Distinguishing Fractal Noises from Motions. Pflügers Archiv - European Journal of Physiology. 2000 Feb; 439(4):403–415. doi: 10.1007/s004249900135.

Eke A, Herman P, Kocsis L, Kozak LR. Fractal Characterization of Complexity in Temporal Physiological Signals. Physiological Measurement. 2002 Feb; 23(1):R1–R38. doi: 10.1088/0967-3334/23/1/201.

Esteban FJ, Padilla N, Sanz-Cortés M, De Miras JR, Bargalló N, Villoslada P, Gratacós E. Fractal-Dimension Analysis Detects Cerebral Changes in Preterm Infants with and without Intrauterine Growth Restriction. NeuroImage. 2010 Dec; 53(4):1225–1232. doi: 10.1016/j.neuroimage.2010.07.019.

Eyre M, Fitzgibbon SP, Ciarrusta J, Cordero-Grande L, Price AN, Poppe T, Schuh A, Hughes E, O’Keeffe C, Bran-don J, Cromb D, Vecchiato K, Andersson J, Duff EP, Counsell SJ, Smith SM, Rueckert D, Hajnal JV, Arichi T, O’Muircheartaigh J, et al. The Developing Human Connectome Project: Typical and Disrupted Perinatal Func-tional Connectivity. Brain. 2021 Aug; 144(7):2199–2213. doi: 10.1093/brain/awab118.

Fitzgibbon SP, Harrison SJ, Jenkinson M, Baxter L, Robinson EC, Bastiani M, Bozek J, Karolis V, Cordero Grande L, Price AN, Hughes E, Makropoulos A, Passerat-Palmbach J, Schuh A, Gao J, Farahibozorg SR, O’Muircheartaigh J, Ciarrusta J, O’Keeffe C, Brandon J, et al. The Developing Human Connectome Project (dHCP) Automated Resting-State Functional Processing Framework for Newborn Infants. NeuroImage. 2020 Dec; 223:117303. doi: 10.1016/j.neuroimage.2020.117303.

Fransson P, Metsäranta M, Blennow M, Åden U, Lagercrantz H, Vanhatalo S. Early Development of Spatial Patterns of Power-Law Frequency Scaling in fMRI Resting-State and EEG Data in the Newborn Brain. Cerebral Cortex. 2013 Mar; 23(3):638–646. doi: 10.1093/cercor/bhs047.

Fransson P, Skiöld B, Horsch S, Nordell A, Blennow M, Lagercrantz H, Åden U. Resting-State Networks in the Infant Brain. Proceedings of the National Academy of Sciences. 2007 Sep; 104(39):15531–15536. doi: 10.1073/pnas.0704380104.

Gao W, Lin W, Grewen K, Gilmore JH. Functional Connectivity of the Infant Human Brain: Plastic and Modifiable. The Neuroscientist. 2017 Apr; 23(2):169–184. doi: 10.1177/1073858416635986.

Griffanti L, Salimi-Khorshidi G, Beckmann CF, Auerbach EJ, Douaud G, Sexton CE, Zsoldos E, Ebmeier KP, Filippini N, Mackay CE, Moeller S, Xu J, Yacoub E, Baselli G, Ugurbil K, Miller KL, Smith SM. ICA-based Artefact Removal and Accelerated fMRI Acquisition for Improved Resting State Network Imaging. NeuroImage. 2014 Jul; 95:232–247. doi: 10.1016/j.neuroimage.2014.03.034.

Hartley C, Berthouze L, Mathieson SR, Boylan GB, Rennie JM, Marlow N, Farmer SF. Long-Range Temporal Correlations in the EEG Bursts of Human Preterm Babies. PLoS ONE. 2012 Feb; 7(2):e31543. doi: 10.1371/jour-nal.pone.0031543.

He BJ. Scale-Free Properties of the Functional Magnetic Resonance Imaging Signal during Rest and Task. Journal of Neuroscience. 2011 Sep; 31(39):13786–13795. doi: 10.1523/JNEUROSCI.2111-11.2011.

Herzmann CS, Snyder AZ, Kenley JK, Rogers CE, Shimony JS, Smyser CD. Cerebellar Functional Connectivity in Term- and Very Preterm-Born Infants. Cerebral Cortex. 2019 Mar; 29(3):1174–1184. doi: 10.1093/cer-cor/bhy023.

Hughes EJ, Winchman T, Padormo F, Teixeira R, Wurie J, Sharma M, Fox M, Hutter J, Cordero-Grande L, Price AN, Allsop J, Bueno-Conde J, Tusor N, Arichi T, Edwards AD, Rutherford MA, Counsell SJ, Hajnal JV. A Dedicated Neonatal Brain Imaging System: A Dedicated Neonatal Brain Imaging System. Magnetic Resonance in Medicine. 2017 Aug; 78(2):794–804. doi: 10.1002/mrm.26462.

Iyer PM, Egan C, Pinto-Grau M, Burke T, Elamin M, Nasseroleslami B, Pender N, Lalor EC, Hardiman O. Functional Connectivity Changes in Resting-State EEG as Potential Biomarker for Amyotrophic Lateral Sclerosis. PLOS ONE. 2015 Jun; 10(6):e0128682. doi: 10.1371/journal.pone.0128682.

Jannesari M, Saeedi A, Zare M, Ortiz-Mantilla S, Plenz D, Benasich AA. Stability of Neuronal Avalanches and Long-Range Temporal Correlations during the First Year of Life in Human Infants. Brain Structure and Function. 2020 Apr; 225(3):1169–1183. doi: 10.1007/s00429-019-02014-4.

Kostović I, Jovanov-Milošević N. The Development of Cerebral Connections during the First 20–45 Weeks’ Gestation. Seminars in Fetal and Neonatal Medicine. 2006 Dec; 11(6):415–422. doi: 10.1016/j.siny.2006.07.001.

Lei X, Zhao Z, Chen H. Extraversion Is Encoded by Scale-Free Dynamics of Default Mode Network. NeuroImage. 2013 Jul; 74:52–57. doi: 10.1016/j.neuroimage.2013.02.020.

Luu TM, Rehman Mian MO, Nuyt AM. Long-Term Impact of Preterm Birth. Clinics in Perinatology. 2017 Jun; 44(2):305–314. doi: 10.1016/j.clp.2017.01.003.

Makropoulos A, Gousias IS, Ledig C, Aljabar P, Serag A, Hajnal JV, Edwards AD, Counsell SJ, Rueckert D. Automatic Whole Brain MRI Segmentation of the Developing Neonatal Brain. IEEE Transactions on Medical Imaging. 2014 Sep; 33(9):1818–1831. doi: 10.1109/TMI.2014.2322280.

Makropoulos A, Robinson EC, Schuh A, Wright R, Fitzgibbon S, Bozek J, Counsell SJ, Steinweg J, Vecchiato K, Passerat-Palmbach J, Lenz G, Mortari F, Tenev T, Duff EP, Bastiani M, Cordero-Grande L, Hughes E, Tusor N, Tournier JD, Hutter J, et al. The Developing Human Connectome Project: A Minimal Processing Pipeline for Neonatal Cortical Surface Reconstruction. NeuroImage. 2018 Jun; 173:88–112. doi: 10.1016/j.neuroimage.2018.01.054.

Maxim V, Sendur L, Fadili J, Suckling J, Gould R, Howard R, Bullmore E. Fractional Gaussian Noise, Functional MRI and Alzheimer’s Disease. NeuroImage. 2005 Mar; 25(1):141–158. doi: 10.1016/j.neuroimage.2004.10.044.

Meisel C, Bailey K, Achermann P, Plenz D. Decline of Long-Range Temporal Correlations in the Human Brain during Sustained Wakefulness. Scientific Reports. 2017 Sep; 7(1):11825. doi: 10.1038/s41598-017-12140-w.

Ment LR, Hirtz D, Hüppi PS. Imaging Biomarkers of Outcome in the Developing Preterm Brain. The Lancet Neurology. 2009 Nov; 8(11):1042–1055. doi: 10.1016/S1474-4422(09)70257-1.

Moran JK, Michail G, Heinz A, Keil J, Senkowski D. Long-Range Temporal Correlations in Resting State Beta Oscillations Are Reduced in Schizophrenia. Frontiers in Psychiatry. 2019 Jul; 10:517. doi: 10.3389/fp-syt.2019.00517.

O’Byrne J, Jerbi K. How Critical Is Brain Criticality? Trends in Neurosciences. 2022 Nov; 45(11):820–837. doi: 10.1016/j.tins.2022.08.007.

Padilla N, Saenger VM, Van Hartevelt TJ, Fernandes HM, Lennartsson F, Andersson JLR, Kringelbach M, Deco G, Åden U. Breakdown of Whole-brain Dynamics in Preterm-born Children. Cerebral Cortex. 2020 Mar; 30(3):1159–1170. doi: 10.1093/cercor/bhz156.

Power JD, Barnes KA, Snyder AZ, Schlaggar BL, Petersen SE. Spurious but Systematic Correlations in Functional Connectivity MRI Networks Arise from Subject Motion. NeuroImage. 2012 Feb; 59(3):2142–2154. doi: 10.1016/j.neuroimage.2011.10.018.

Ream MA, Lehwald L. Neurologic Consequences of Preterm Birth. Current Neurology and Neuroscience Reports. 2018 Aug; 18(8):48. doi: 10.1007/s11910-018-0862-2.

Rogers CE, Lean RE, Wheelock MD, Smyser CD. Aberrant Structural and Functional Connectivity and Neurodevelopmental Impairment in Preterm Children. Journal of Neurodevelopmental Disorders. 2018 Dec; 10(1):38. doi: 10.1186/s11689-018-9253-x.

Rubin D, Fekete T, Mujica-Parodi LR. Optimizing Complexity Measures for fMRI Data: Algorithm, Artifact, and Sensitivity. PLoS ONE. 2013 May; 8(5):e63448. doi: 10.1371/journal.pone.0063448.

Russell V Lenth, Emmeans: Estimated Marginal Means, Aka Least-Squares Means; 2022.

Salimi-Khorshidi G, Douaud G, Beckmann CF, Glasser MF, Griffanti L, Smith SM. Automatic Denoising of Functional MRI Data: Combining Independent Component Analysis and Hierarchical Fusion of Classifiers. NeuroImage. 2014 Apr; 90:449–468. doi: 10.1016/j.neuroimage.2013.11.046.

Schuh A. Computational Models of the Morphology of the Developing Neonatal Human Brain.. 2017 Sep; doi: 10.25560/58880.

Schuh A, Makropoulos A, Robinson EC, Cordero-Grande L, Hughes E, Hutter J, Price AN, Murgasova M, Teixeira RPAG, Tusor N, Steinweg JK, Victor S, Rutherford MA, Hajnal JV, Edwards AD, Rueckert D. Unbiased Construction of a Temporally Consistent Morphological Atlas of Neonatal Brain Development. Neuroscience; 2018.

Smyrni N, Koutsaki M, Petra M, Nikaina E, Gontika M, Strataki H, Davora F, Bouza H, Damianos G, Skouteli H, Mastroyianni S, Dalivigka Z, Dinopoulos A, Tzaki M, Papavasiliou A. Moderately and Late Preterm Infants: Short-and Long-Term Outcomes From a Registry-Based Cohort. Frontiers in Neurology. 2021 Feb; 12:628066. doi: 10.3389/fneur.2021.628066.

Smyser CD, Inder TE, Shimony JS, Hill JE, Degnan AJ, Snyder AZ, Neil JJ. Longitudinal Analysis of Neural Network Development in Preterm Infants. Cerebral Cortex. 2010 Dec; 20(12):2852–2862. doi: 10.1093/cercor/bhq035.

Smyser CD, Snyder AZ, Neil JJ. Functional Connectivity MRI in Infants: Exploration of the Functional Organization of the Developing Brain. NeuroImage. 2011 Jun; 56(3):1437–1452. doi: 10.1016/j.neuroimage.2011.02.073.

Soares JM, Magalhães R, Moreira PS, Sousa A, Ganz E, Sampaio A, Alves V, Marques P, Sousa N. A Hitchhiker’s Guide to Functional Magnetic Resonance Imaging. Frontiers in Neuroscience. 2016 Nov; 10. doi: 10.3389/fnins.2016.00515.

Spoto G, Amore G, Vetri L, Quatrosi G, Cafeo A, Gitto E, Nicotera AG, Di Rosa G. Cerebellum and Prematurity: A Complex Interplay Between Disruptive and Dysmaturational Events. Frontiers in Systems Neuroscience. 2021 Jun; 15:655164. doi: 10.3389/fnsys.2021.655164.

Virtanen P, Gommers R, Oliphant TE, Haberland M, Reddy T, Cournapeau D, Burovski E, Peterson P, Weckesser W, Bright J, van der Walt SJ, Brett M, Wilson J, Millman KJ, Mayorov N, Nelson ARJ, Jones E, Kern R, Larson E, Carey CJ, et al. SciPy 1.0: Fundamental Algorithms for Scientific Computing in Python. Nature Methods. 2020 Mar; 17(3):261–272. doi: 10.1038/s41592-019-0686-2.

Vo Van P, Alison M, Morel B, Beck J, Bednarek N, Hertz-Pannier L, Loron G. Advanced Brain Imaging in Preterm Infants: A Narrative Review of Microstructural and Connectomic Disruption. Children. 2022 Mar; 9(3):356. doi: 10.3390/children9030356.

Vogel JP, Chawanpaiboon S, Moller AB, Watananirun K, Bonet M, Lumbiganon P. The Global Epidemiology of Preterm Birth. Best Practice & Research Clinical Obstetrics & Gynaecology. 2018 Oct; 52:3–12. doi: 10.1016/j.bpobgyn.2018.04.003.

Volpe JJ. Cerebellum of the Premature Infant: Rapidly Developing, Vulnerable, Clinically Important. Journal of Child Neurology. 2009 Sep; 24(9):1085–1104. doi: 10.1177/0883073809338067.

Weinstein M, Ben-Sira L, Moran A, Berger I, Marom R, Geva R, Gross-Tsur V, Leitner Y, Ben Bashat D. The Motor and Visual Networks in Preterm Infants: An fMRI and DTI Study. Brain Research. 2016 Jul; 1642:603–611. doi: 10.1016/j.brainres.2016.04.052.

Welch P. The Use of Fast Fourier Transform for the Estimation of Power Spectra: A Method Based on Time Averaging over Short, Modified Periodograms. IEEE Transactions on Audio and Electroacoustics. 1967 Jun; 15(2):70–73. doi: 10.1109/TAU.1967.1161901.

Wink AM, Bullmore E, Barnes A, Bernard F, Suckling J. Monofractal and Multifractal Dynamics of Low Frequency Endogenous Brain Oscillations in Functional MRI. Human Brain Mapping. 2008; 29(7):791–801. doi: 10.1002/hbm.20593.

World Health Organization, Preterm Birth; 2023.

Yuen TJ, Silbereis JC, Griveau A, Chang SM, Daneman R, Fancy SPJ, Zahed H, Maltepe E, Rowitch DH. Oligodendrocyte-Encoded HIF Function Couples Postnatal Myelination and White Matter Angiogenesis. Cell. 2014 Jul; 158(2):383–396. doi: 10.1016/j.cell.2014.04.052.

Zimmern V. Why Brain Criticality Is Clinically Relevant: A Scoping Review. Frontiers in Neural Circuits. 2020 Aug; 14:54. doi: 10.3389/fncir.2020.00054.

