## Supplementary material for "Temporal Complexity of the BOLD-Signal In Preterm Versus Term Infants": All supplementary data: eLife_RAnalysis.pdf

### dHCP H Study of Temporal Complexity (Nov, 2023)

21 November, 2023

#### Intro

We are assessing a measure of the BOLD temporal complexity, H, in preterm born infants compared to term born controls.

Manuscript Title: **Temporal complexity of BOLD-signal in preterm versus term infants**

Study Questions: 1) How does H develop over time?
2) Do different RSNs develop differently with regards to H values?
3) Do term born infants have greater H compared to preterm infants at scan age?
4) Is H associated with indirect measures of myelination (FA and RD)?

Linear mixed effects models will be created to take into account of random effects and interacting terms. Significant differences will be determined through using estimated marginal means (emmeans) and trends (emtrends). Multiple comparisons will be controlled for using Holm's method.

#### Load data

```
# Data Loading
##CSV file load
ClinicalInfo <- read.csv("csvfiles/ClinicalInfo.csv")
H_Tissue <- read.csv("csvfiles/new_Hvalues.csv")
H_RSN <- read.csv("csvfiles/new_HvalinRSN_wFIX.csv")
#Data merging
fulldata <- merge(ClinicalInfo,H_Tissue)
fulldata <- merge(fulldata, H_RSN)
```

After loading our data, we have 803 rows.

We will exclude subjects with large motion (meanFD)

```
meanFD <- read.csv("csvfiles/meanFD.csv")
newdata <- merge(fulldata, meanFD)
newdata <- newdata %>% filter(meanFD < 0.5)
```

Which leaves us with 706 rows. That means we lost 97 rows.

#### GAfactors

Sorting the participants into 3 groups according to their gestational age (i.e. birth age).
VPT = very preterm
MPT = moderately preterm
THC = term healthy control

```r
fulldata <- fulldata %>% mutate(GAfactor = case_when(
    birth_age <= 32  ~ "VPT",
    birth_age > 32 & birth_age <= 37 ~ "MPT",
    birth_age >37 ~ "THC"
    ))
fulldata$GAfactor <- as.factor(fulldata$GAfactor)
fulldata$GAfactor <- factor(fulldata$GAfactor, levels = c("VPT", "MPT", "THC"))
```

##### Setting ages for when we will estimate marginal means and trends

Using median birth ages in the VPT, MPT and THC groups
VPT = 29 weeks GA
MPT = 35 weeks GA
THC = 41 weeks GA

preterm comparison age = 35 weeks PMA
term comparison age = 41 weeks PMA

#### Initial Linear Mixed Effects Model Checks

Our main model will be Hurst with fixed effects: Birth Age, Scan Age, and ROI

Subjects will be set as random effects to take into account multiple scans of the same participant.

The first question we have is should Birth Age and Scan Age be interactive terms? (i.e. Birth Age + Scan Age or Birth Age * Scan Age)

We will use an anova of the two models to decide:

```r
# Setting up our linear mixed effects model. Determining whether we will need to to take
# into account of PMA and GA.
# First: Model with no interaction:
h.model1 <- lmer(data = fulldata_long, H ~ scan_age + birth_age + (1|Subject), REML = F)
```

```r
#Nect: Model with interaction
h.model.int2 <- lmer(data = fulldata_long, H ~ scan_age*birth_age + (1|Subject), REML = F)
```

```r
# test for significance of interaction
GAPMAanova <- anova(h.model.int2, h.model1)
GAPMAanova
```

```
## Data: fulldata_long
## Models:
## h.model1: H ~ scan_age + birth_age + (1 | Subject)
## h.model.int2: H ~ scan_age * birth_age + (1 | Subject)
##              npar    AIC    BIC logLik deviance  Chisq Df Pr(>Chisq)
## h.model1        5 -20163 -20128  10086   -20173
## h.model.int2    6 -20176 -20134  10094   -20188 15.261  1  9.364e-05 ***
## ---
## Signif. codes:  0 '***' 0.001 '**' 0.01 '*' 0.05 '.' 0.1 ' ' 1
```

```r
# If p > 0.05, there is no difference between these models, so we wouldn't need to include
# PMA and GA as interactive terms
```

As $p < 0.05$, (check: TRUE), we will include Birth Age and Scan Age as interactive terms.

### Demographics

```
distinctn_old <- oldfulldata %>% distinct(Subject) %>% summarise(Observations = n())
```

Our original number of distinct subjects was 716

```
distinctn <- fulldata %>% distinct(Subject) %>% summarise (Observations = n())
```

Our number of distinct subjects after filtering for meanFD is: 641

```
##              latexvariable latexvalue
## 1      Initial_Number_Rows        803
## 2 Number_Rows_After_meanFD        706
## 3      Number_of_rows_lost         97
```

#### Range of GA

The minimum and maximum gestational age of our entire cohort is 23 and 42.71, respectively.

#### Group Sample Sizes

The initial amount of distinct subjects from each GAfactor group was:

| GAfactor | Observations |
|---|---:|
| VPT | 88 |
| MPT | 110 |
| THC | 518 |

After meanFD filtering, we are left with:

| GAfactor | Observations |
|---|---:|
| VPT | 82 |
| MPT | 105 |
| THC | 454 |

The number of preterm infants with both preterm and term scans is:

| GAfactor | Observations |
|---|---:|
| VPT | 37 |
| MPT | 28 |

Thus, the resulting preterm group was composed of 187 infants including 81 infants with preterm only scans, 41 infants with term-equivalent only scans, and 65 infants with both preterm and term-equivalent aged scans.

```
## (polygon[GRID.polygon.11], polygon[GRID.polygon.12], polygon[GRID.polygon.13], polygon[GRID.polygon.
```

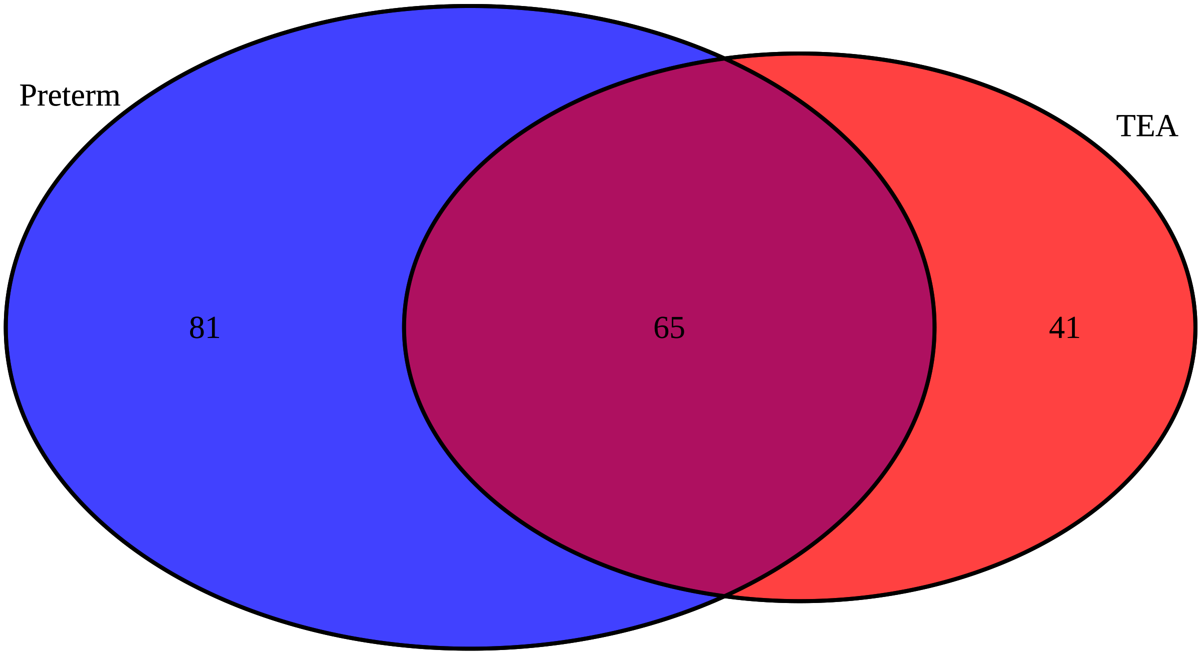


#### Basic Stats

Non-parametric stats because our data is not normally distributed.

##### Birth ages

Preterm birth median age in VPT and MPT

| GAfactor | Observations | Median.BirthAge | Range |
|---|---|---|---|
| VPT | 82 | 29.07 | 23.00 |
| VPT | 82 | 29.07 | 31.86 |
| MPT | 105 | 34.86 | 32.14 |
| MPT | 105 | 34.86 | 37.00 |
| THC | 454 | 40.14 | 37.14 |
| THC | 454 | 40.14 | 42.71 |

##### Scan ages/ PMA (Median)

Preterm scan median age in VPT and MPT

| GAfactor | Observations | Median.Scanage | Range |
|---|---|---|---|
| VPT | 70 | 32.64 | 26.71 |
| VPT | 70 | 32.64 | 36.71 |
| MPT | 76 | 35.57 | 33.14 |
| MPT | 76 | 35.57 | 37.00 |

TEA scan median age in VPT and MPT

| GAfactor | Observations | Median.Scanage | Range |
|---|---|---|---|
| VPT | 49 | 40.86 | 38.14 |
| VPT | 49 | 40.86 | 44.57 |
| MPT | 57 | 40.57 | 37.14 |
| MPT | 57 | 40.57 | 45.14 |

Preterm scan median age in combined preterm group

| Observations | Median.Scanage |
|---|---|
| 138 | 34.86 |

TEA scan median age in combined preterm group

| Observations | Median.Scanage |
|---|---|
| 49 | 40.43 |

Term scan median age in all groups

| GAfactor | Observations | Median.Scanage | Range |
|---|---|---|---|
| VPT | 16 | 41.14 | 38.14 |
| VPT | 16 | 41.14 | 44.29 |
| MPT | 33 | 40.29 | 37.14 |
| MPT | 33 | 40.29 | 45.14 |
| THC | 454 | 41.29 | 37.43 |
| THC | 454 | 41.29 | 44.86 |

THC scan median age

| Observations | Median.Scanage |
|---|---|
| 454 | 41.29 |

##### Birth Age/Gestational Age

| GAfactor | Observations | Median.Birthage | Range | n |
|---|---|---|---|---|
| VPT | 82 | 29.07 | 23.00 | 82 |
| VPT | 82 | 29.07 | 31.86 | 82 |
| MPT | 105 | 34.86 | 32.14 | 105 |
| MPT | 105 | 34.86 | 37.00 | 105 |
| THC | 454 | 40.14 | 37.14 | 454 |
| THC | 454 | 40.14 | 42.71 | 454 |

##### Birth Weight

| GAfactor | Observations | Median.Birthweight |
|---|---|---|
| VPT | 82 | 1.17 |
| MPT | 105 | 2.18 |
| THC | 454 | 3.40 |

##### Head Circumference

| GAfactor | Observations | Median.Headcircumference |
|---|---|---|
| VPT | 82 | 28 |
| MPT | 105 | 32 |
| THC | 454 | 35 |

##### Sedation

| GAfactor | Observations |
|---|---|
| THC | 6 |

##### Radiology scores

5=Incidental finding with possible / likely significance for both clinical and imaging analysis (e.g. Major lesions within white matter cortex, cerebellum and or basal ganglia; small head / brain < 1 st percentile

##### Radiology Score of 1:

Total = 320

Breakdown by GAfactor:

| GAfactor | Observations |
|---|---|
| VPT | 29 |
| MPT | 39 |
| THC | 252 |

**Radiology Score of 2:**

Total = 198

Breakdown by GAfactor:

| GAfactor | Observations |
|---|---|
| VPT | 22 |
| MPT | 33 |
| THC | 143 |

**Radiology Score of 3:**

Total = 82

Breakdown by GAfactor:

| GAfactor | Observations |
|---|---|
| VPT | 16 |
| MPT | 30 |
| THC | 36 |

**Radiology Score of 4:**

Total = 15

Breakdown by GAfactor:

| GAfactor | Observations |
|---|---|
| VPT | 7 |
| MPT | 2 |
| THC | 6 |

**Radiology Score of 5:**

Total = 53

Breakdown by GAfactor:

| GAfactor | Observations |
|---|---|
| VPT | 24 |
| MPT | 13 |
| THC | 16 |

**Sexes**

**Female**

Total = 292

Breakdown by GAfactor:

| GAfactor | Observations |
|---|---|
| VPT | 39 |
| MPT | 47 |
| THC | 206 |

**Male**

Total = 349

Breakdown by GAfactor:

| GAfactor | Observations |
|---|---|
| VPT | 43 |
| MPT | 58 |
| THC | 248 |

##### Birth age and scan age colinearity?

Preterm (combined) and term group scan age histogram

Checking to see that both groups are approximately scanned at the same age to make sure we are not extrapolating data when predicted H values at set scan age.

Emmeans at 41 weeks, controls for colinearity issue.

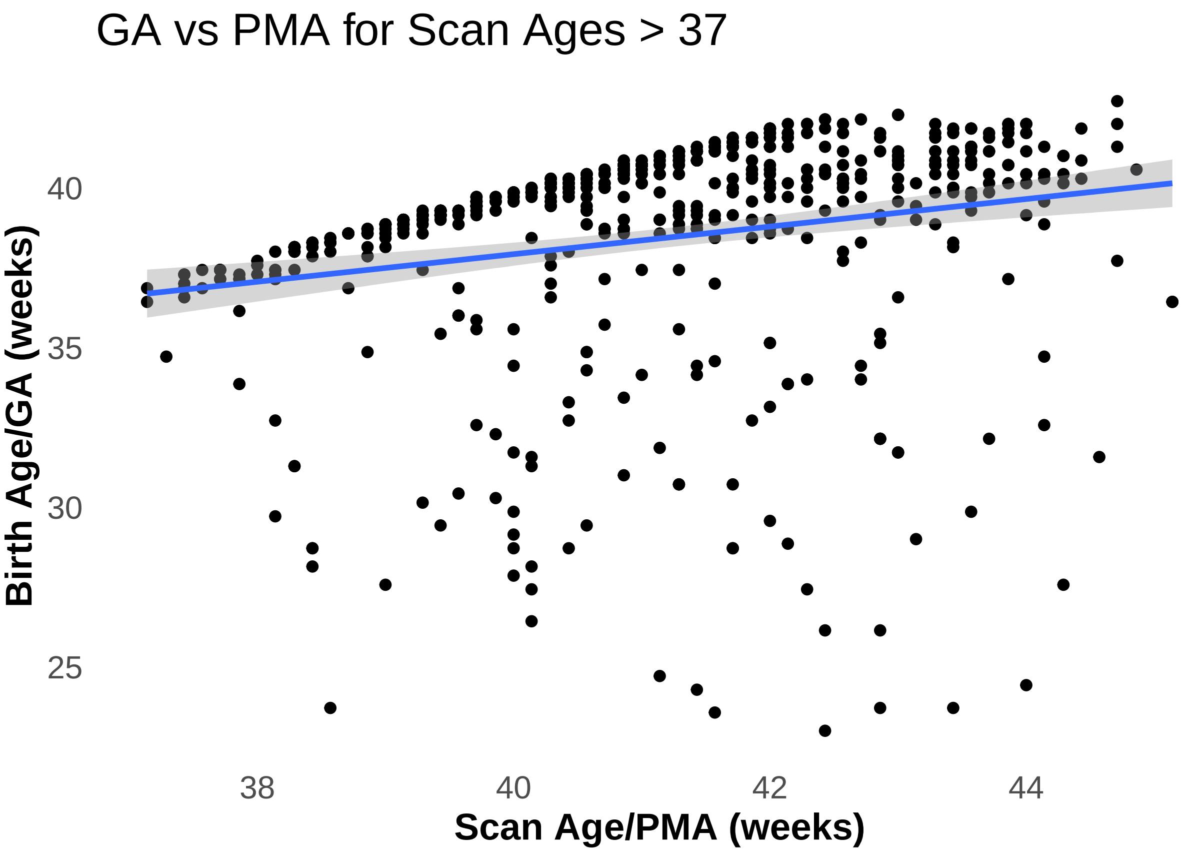


Preterm groups and term group scan age histogram

```
RemovedpretermGAfactors %>% distinct(Subject, scan_age, GAfactor) %>%
  filter(scan_age > 37) %>%
  ggplot(aes(x=scan_age, color=GAfactor)) +
  geom_histogram(fill="white",alpha=0.5, position="identity") +
```

```r
  theme_minimalism() +
  scale_color_manual(values = c("indianred1", "goldenrod1", "deepskyblue"),
                     name = "Group", labels = c("VPT", "MPT", "THC")) +
  xlab("Scan Age/PMA (weeks)") +
  ylab("Count") +
  ggtitle("PMA Histogram for Scan Ages > 37")
```

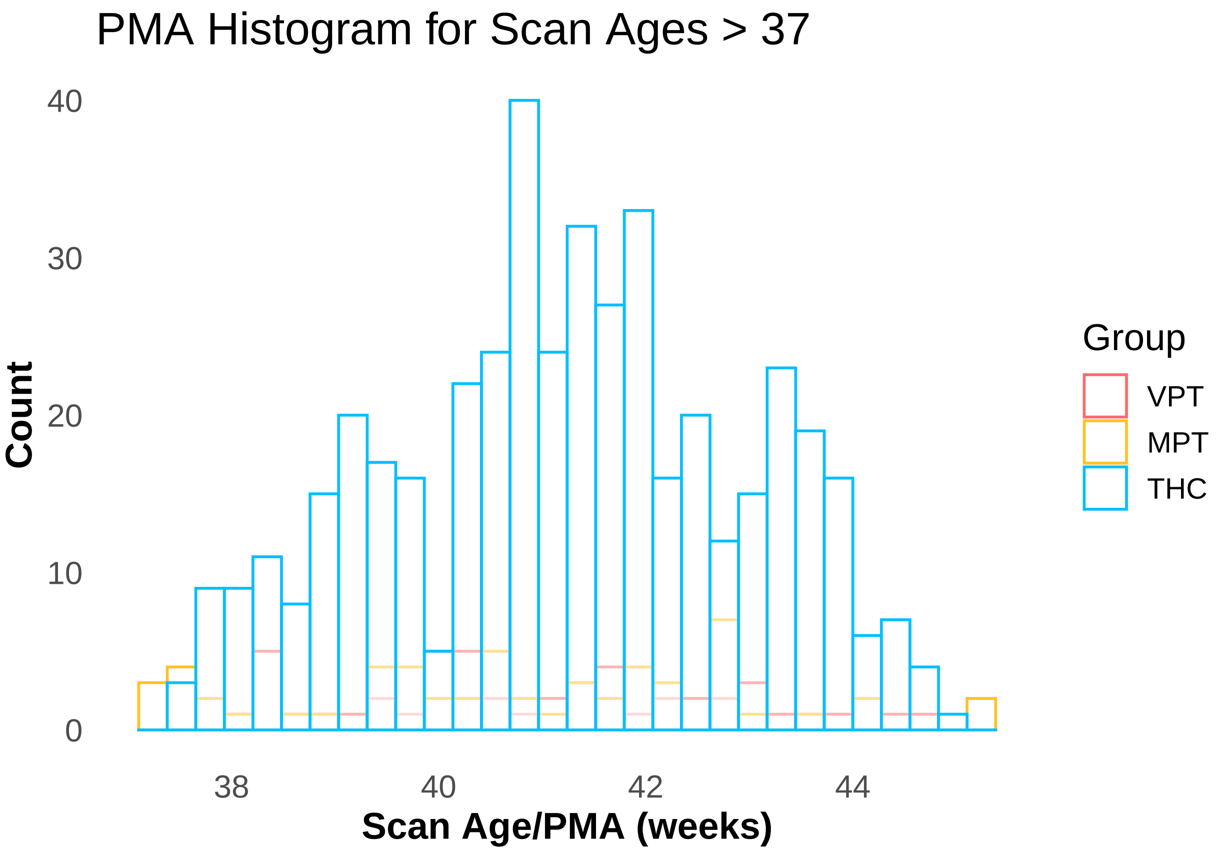


Correlation test

| estimate | p.value | conf.low | conf.high | method |
|---|---|---|---|---|
| 0.205 | 0 | 0.125 | 0.283 | Pearson's product-moment correlation |

#### Linear Mixed Effects Model: GM in Entire Cohort

Looking at GA as a continuous variable in the grey matter.

Let's set-up our model for H in the grey matter, with scan age and birth age as interative terms

```r
GMwholecohortmodel <- lmer(data = fulldata, HinGM ~ scan_age*birth_age +
                           (1|Subject), REML = F)
```

Making sure birth age and scan age should be interative terms

| | | npar | AIC | BIC | logLik | deviance | Chisq | Df | Pr(>Chisq) | |
|---|---|---|---|---|---|---|---|---|---|---|
| 2 | GMwholecohortmodel | 6 | -1688.801 | -1661.443 | 850.4006 | -1700.801 | 32.45479 | 2 | <0.001 | *** |

Summary of full model:

| | Estimate | CI (lower) | CI (upper) | Std. Error | t value |
|---|---|---|---|---|---|
| (Intercept) | -0.4671698 | -0.8712290 | -0.0542560 | 0.2031004 | -2.300192 |
| scan_age | 0.0211753 | 0.0108272 | 0.0312549 | 0.0050792 | 4.168975 |
| birth_age | 0.0134637 | 0.0006257 | 0.0261284 | 0.0063613 | 2.116481 |
| scan_age:birth_age | -0.0002223 | -0.0005326 | 0.0000933 | 0.0001561 | -1.424348 |

Report blurb:

We fitted a linear mixed model (estimated using ML and nloptwrap optimizer) to predict HinGM with scan_age and birth_age (formula: HinGM ~ scan_age * birth_age). The model included Subject as random effect (formula: ~1 | Subject). The model's total explanatory power is substantial (conditional R2 = 0.62) and the part related to the fixed effects alone (marginal R2) is of 0.43. The model's intercept, corresponding to scan_age = 0 and birth_age = 0, is at -0.47 (95% CI [-0.87, -0.07], t(700) = -2.30, p = 0.022). Within this model:

- The effect of scan age is statistically significant and positive (beta = 0.02, 95% CI [0.01, 0.03], t(700) = 4.17, p < .001; Std. beta = 0.46, 95% CI [0.38, 0.55])
- The effect of birth age is statistically significant and positive (beta = 0.01, 95% CI [9.74e-04, 0.03], t(700) = 2.12, p = 0.035; Std. beta = 0.22, 95% CI [0.14, 0.29])
- The effect of scan age × birth age is statistically non-significant and negative (beta = -2.22e-04, 95% CI [-5.29e-04, 8.41e-05], t(700) = -1.42, p = 0.155; Std. beta = -0.04, 95% CI [-0.09, 0.01])

Standardized parameters were obtained by fitting the model on a standardized version of the dataset. 95% Confidence Intervals (CIs) and p-values were computed using a Wald t-distribution approximation.

#### Raw correlation

H in GM vs birth age:

| estimate | statistic | p.value | parameter | conf.low | conf.high | method | alternative |
|---|---|---|---|---|---|---|---|
| 0.5326697 | 16.69968 | 0 | 704 | 0.4776565 | 0.583522 | Pearson's product-moment correlation | two.sided |

H in GM vs scan age:

| estimate | statistic | p.value | parameter | conf.low | conf.high | method | alternativ |
|---|---|---|---|---|---|---|---|
| 0.6326102 | 21.67294 | 0 | 704 | 0.5861854 | 0.6748943 | Pearson's product-moment correlation | two.sided |

```
(HGMvsGA <- ggplot(data = fulldata, aes(x = birth_age, y = HinGM)) +
  geom_point(shape=1, alpha = 1/3) + geom_smooth(method = "lm", se=T) +
  theme_minimalism() +
  xlab("Birth age (weeks)") +
  ylab("H in the GM") +
  theme(legend.title=element_blank()))
```

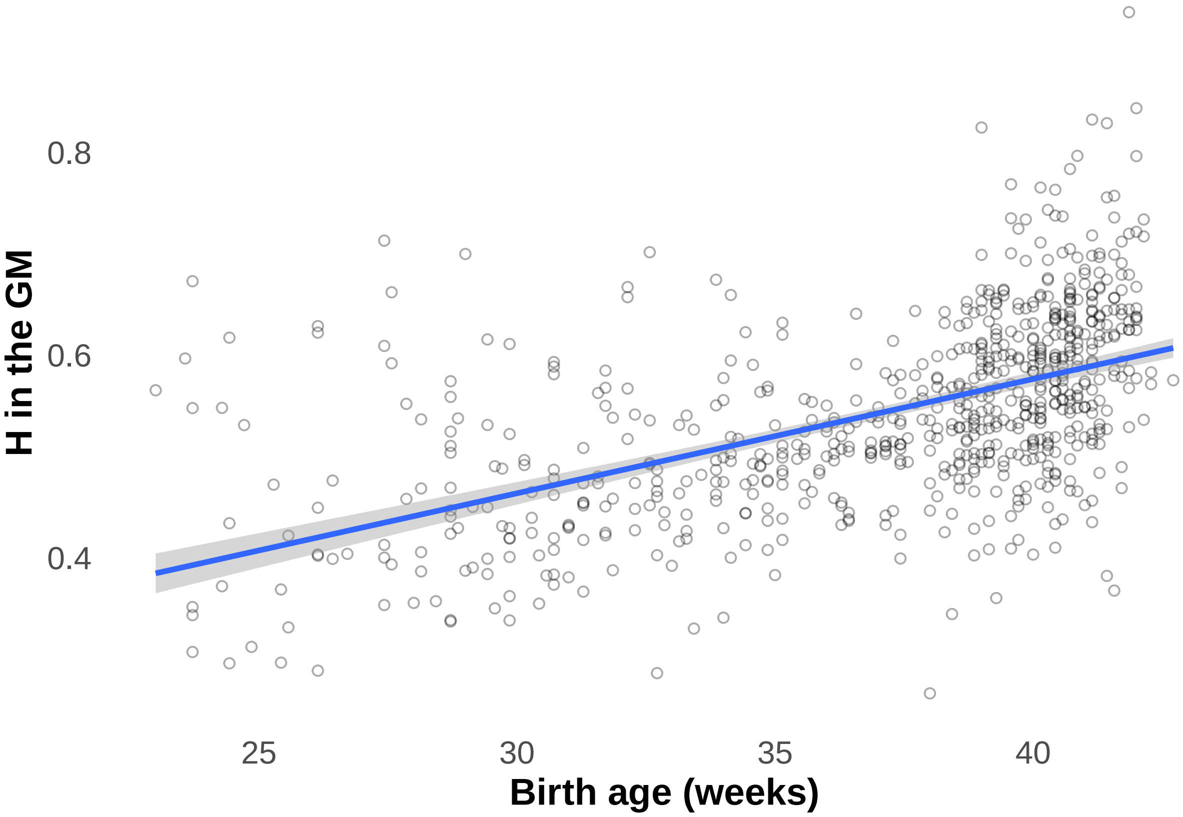


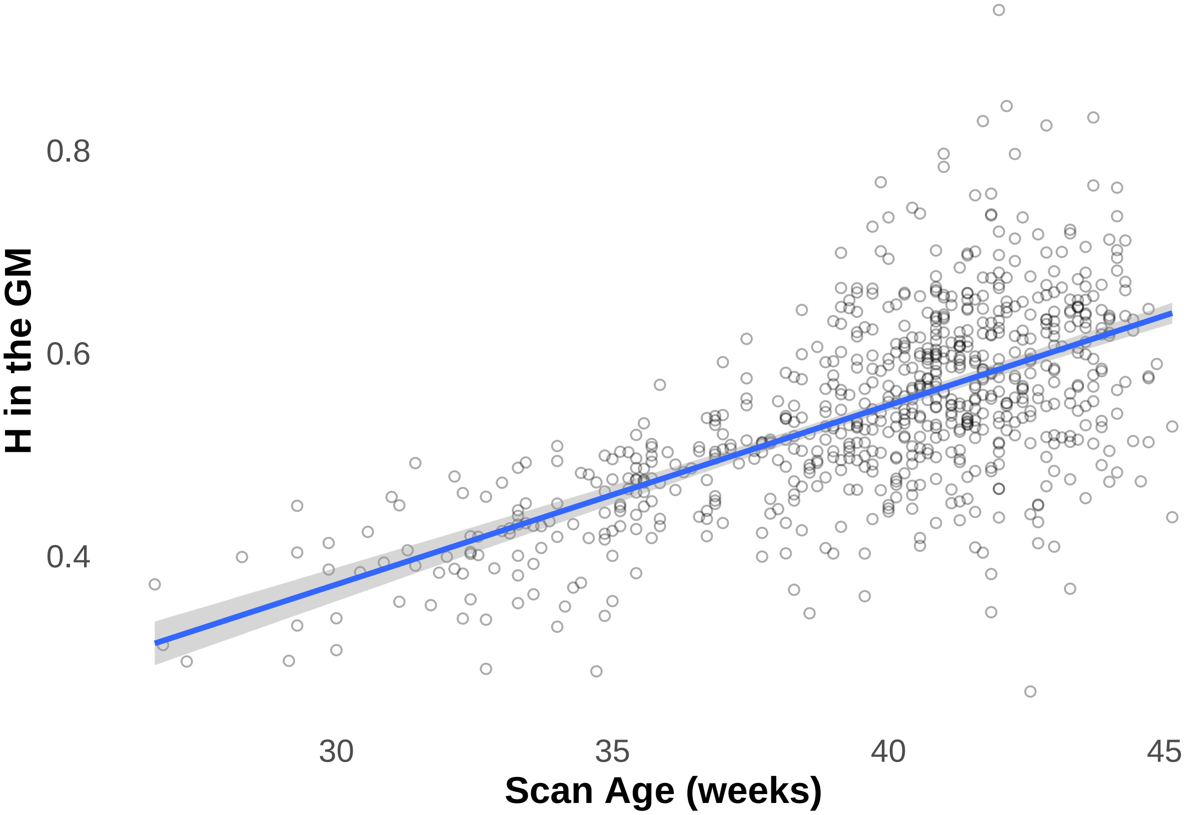


Whole group GM plot

##### How does H develop with respect to birth age?

Let's look at the estimated marginal trend, keeping scan age constant at 41weeks ("term age")

```
##  scan_age birth_age.trend       SE  df lower.CL upper.CL
##        41         0.00435 0.000811 705  0.00276  0.00594
## 
## Degrees-of-freedom method: kenward-roger 
## Confidence level used: 0.95
```

So if we scan a baby at term age (41 weeks), the rate of change of H in GM with respect to birth age is 0.0043

#### Linear Mixed Effects Model: White Matter

Model, summary and report:

```
## Linear mixed model fit by maximum likelihood  ['lmerMod']
## Formula: HinWM ~ scan_age * birth_age + (1 | Subject)
##    Data: fulldata
## 
##      AIC      BIC   logLik deviance df.resid 
##  -2292.1  -2264.7   1152.0  -2304.1      700 
## 
## Scaled residuals: 
##     Min      1Q  Median      3Q     Max 
## -5.3816 -0.4769  0.1323  0.6127  3.1630 
## 
## Random effects:
##  Groups   Name        Variance  Std.Dev.
##  Subject  (Intercept) 0.0002873 0.01695 
##  Residual             0.0019558 0.04422 
## Number of obs: 706, groups:  Subject, 641
## 
## Fixed effects:
##                      Estimate Std. Error t value
## (Intercept)        -0.1896559  0.1360378  -1.394
## scan_age            0.0137454  0.0034303   4.007
## birth_age           0.0128231  0.0042153   3.042
## scan_age:birth_age -0.0002425  0.0001041  -2.330
## 
## Correlation of Fixed Effects:
##             (Intr) scan_g brth_g
## scan_age    -0.990              
## birth_age   -0.983  0.959       
## scn_g:brth_  0.988 -0.981 -0.992
```

```
## We fitted a linear mixed model (estimated using ML and nloptwrap optimizer) to
## predict HinWM with scan_age and birth_age (formula: HinWM ~ scan_age *
## birth_age). The model included Subject as random effect (formula: ~1 |
## Subject). The model's total explanatory power is substantial (conditional R2 =
## 0.40) and the part related to the fixed effects alone (marginal R2) is of 0.31.
## The model's intercept, corresponding to scan_age = 0 and birth_age = 0, is at
## -0.19 (95% CI [-0.46, 0.08], t(700) = -1.39, p = 0.164). Within this model:
## 
##   - The effect of scan age is statistically significant and positive (beta =
## 0.01, 95% CI [7.01e-03, 0.02], t(700) = 4.01, p < .001; Std. beta = 0.29, 95%
```

```
## CI [0.19, 0.39])
##   - The effect of birth age is statistically significant and positive (beta =
## 0.01, 95% CI [4.55e-03, 0.02], t(700) = 3.04, p = 0.002; Std. beta = 0.26, 95%
## CI [0.17, 0.34])
##   - The effect of scan age × birth age is statistically significant and negative
## (beta = -2.43e-04, 95% CI [-4.47e-04, -3.82e-05], t(700) = -2.33, p = 0.020;
## Std. beta = -0.07, 95% CI [-0.12, -0.01])
##
## Standardized parameters were obtained by fitting the model on a standardized
## version of the dataset. 95% Confidence Intervals (CIs) and p-values were
## computed using a Wald t-distribution approximation.
```

#### Linear Mixed Effects Model: RSN

Model summary:

```
## Linear mixed model fit by maximum likelihood  ['lmerMod']
## Formula: H ~ scan_age * birth_age * RSN + (1 | Subject)
##    Data: fulldata_long
##
##      AIC      BIC   logLik deviance df.resid
## -27795.6 -27410.9  13951.8 -27903.6     9124
##
## Scaled residuals:
##     Min      1Q  Median      3Q     Max
## -7.1659 -0.5355 -0.0280  0.5049  7.2052
##
## Random effects:
##  Groups   Name        Variance Std.Dev.
##  Subject  (Intercept) 0.004390 0.06625
##  Residual             0.002215 0.04706
## Number of obs: 9178, groups:  Subject, 641
##
## Fixed effects:
##                                                   Estimate Std. Error t value
## (Intercept)                                     -1.329e+00  1.608e-01  -8.267
## scan_age                                         4.687e-02  3.963e-03  11.828
## birth_age                                        3.212e-02  5.081e-03   6.321
## RSNH_MotorLateral                                3.992e-01  1.934e-01   2.064
## RSNH_Visual                                      4.880e-01  1.934e-01   2.523
## RSNH_Auditory                                    7.409e-01  1.934e-01   3.830
## RSNH_MotorAssociation                            4.689e-01  1.934e-01   2.424
## RSNH_PosteriorParietal                           8.216e-01  1.934e-01   4.247
## RSNH_Somatosensory                               7.363e-01  1.934e-01   3.806
## RSNH_DorsalVisualStream                          8.792e-01  1.934e-01   4.545
## RSNH_PosteriorCingulateCortex                    7.680e-01  1.934e-01   3.970
## RSNH_FrontalPole                                 6.741e-01  1.934e-01   3.485
## RSNH_Midbrain                                    7.202e-01  1.934e-01   3.723
## RSNH_DorsolateralPrefrontal                      7.330e-01  1.934e-01   3.789
## RSNH_Cerebellum                                  2.554e-01  1.934e-01   1.320
## scan_age:birth_age                              -7.392e-04  1.234e-04  -5.989
## scan_age:RSNH_MotorLateral                      -1.501e-02  4.899e-03  -3.065
## scan_age:RSNH_Visual                            -1.531e-02  4.899e-03  -3.124
## scan_age:RSNH_Auditory                          -2.357e-02  4.899e-03  -4.812
```

```
## scan_age:RSNH_MotorAssociation                    -1.615e-02  4.899e-03  -3.297
## scan_age:RSNH_PosteriorParietal                   -2.066e-02  4.899e-03  -4.218
## scan_age:RSNH_Somatosensory                       -2.209e-02  4.899e-03  -4.508
## scan_age:RSNH_DorsalVisualStream                  -2.614e-02  4.899e-03  -5.336
## scan_age:RSNH_PosteriorCingulateCortex            -2.249e-02  4.899e-03  -4.590
## scan_age:RSNH_FrontalPole                         -2.205e-02  4.899e-03  -4.501
## scan_age:RSNH_Midbrain                            -2.060e-02  4.899e-03  -4.205
## scan_age:RSNH_DorsolateralPrefrontal              -2.406e-02  4.899e-03  -4.911
## scan_age:RSNH_Cerebellum                          -1.869e-02  4.899e-03  -3.815
## birth_age:RSNH_MotorLateral                       -7.359e-03  5.961e-03  -1.235
## birth_age:RSNH_Visual                             -1.280e-02  5.961e-03  -2.147
## birth_age:RSNH_Auditory                           -1.576e-02  5.961e-03  -2.643
## birth_age:RSNH_MotorAssociation                   -2.971e-03  5.961e-03  -0.498
## birth_age:RSNH_PosteriorParietal                  -2.386e-02  5.961e-03  -4.003
## birth_age:RSNH_Somatosensory                      -1.565e-02  5.961e-03  -2.625
## birth_age:RSNH_DorsalVisualStream                 -1.771e-02  5.961e-03  -2.971
## birth_age:RSNH_PosteriorCingulateCortex           -1.152e-02  5.961e-03  -1.932
## birth_age:RSNH_FrontalPole                        -4.652e-03  5.961e-03  -0.780
## birth_age:RSNH_Midbrain                           -4.215e-03  5.961e-03  -0.707
## birth_age:RSNH_DorsolateralPrefrontal             -7.825e-03  5.961e-03  -1.313
## birth_age:RSNH_Cerebellum                          9.613e-03  5.961e-03   1.613
## scan_age:birth_age:RSNH_MotorLateral               2.993e-04  1.477e-04   2.026
## scan_age:birth_age:RSNH_Visual                     3.507e-04  1.477e-04   2.375
## scan_age:birth_age:RSNH_Auditory                   4.843e-04  1.477e-04   3.279
## scan_age:birth_age:RSNH_MotorAssociation           1.341e-04  1.477e-04   0.908
## scan_age:birth_age:RSNH_PosteriorParietal          5.565e-04  1.477e-04   3.768
## scan_age:birth_age:RSNH_Somatosensory              4.352e-04  1.477e-04   2.947
## scan_age:birth_age:RSNH_DorsalVisualStream         4.747e-04  1.477e-04   3.214
## scan_age:birth_age:RSNH_PosteriorCingulateCortex  2.870e-04  1.477e-04   1.944
## scan_age:birth_age:RSNH_FrontalPole                1.746e-04  1.477e-04   1.182
## scan_age:birth_age:RSNH_Midbrain                   7.198e-05  1.477e-04   0.487
## scan_age:birth_age:RSNH_DorsolateralPrefrontal    2.581e-04  1.477e-04   1.747
## scan_age:birth_age:RSNH_Cerebellum                -4.593e-05  1.477e-04  -0.311
```

Report:

```
## We fitted a linear mixed model (estimated using ML and nloptwrap optimizer) to
## predict H with scan_age, birth_age and RSN (formula: H ~ scan_age * birth_age *
## RSN). The model included Subject as random effect (formula: ~1 | Subject). The
## model's total explanatory power is substantial (conditional R2 = 0.83) and the
## part related to the fixed effects alone (marginal R2) is of 0.49. The model's
## intercept, corresponding to scan_age = 0, birth_age = 0 and RSN =
## H_MotorMedial, is at -1.33 (95% CI [-1.64, -1.01], t(9124) = -8.27, p < .001).
## Within this model:
##
##   - The effect of scan age is statistically significant and positive (beta =
## 0.05, 95% CI [0.04, 0.05], t(9124) = 11.83, p < .001; Std. beta = 0.59, 95% CI
## [0.53, 0.65])
##   - The effect of birth age is statistically significant and positive (beta =
## 0.03, 95% CI [0.02, 0.04], t(9124) = 6.32, p < .001; Std. beta = 0.11, 95% CI
## [0.05, 0.18])
##   - The effect of RSN [H_MotorLateral] is statistically significant and positive
## (beta = 0.40, 95% CI [0.02, 0.78], t(9124) = 2.06, p = 0.039; Std. beta =
## -0.26, 95% CI [-0.31, -0.21])
##   - The effect of RSN [H_Visual] is statistically significant and positive (beta
```

```
## = 0.49, 95% CI [0.11, 0.87], t(9124) = 2.52, p = 0.012; Std. beta = -0.69, 95%
## CI [-0.74, -0.64])
##   - The effect of RSN [H_Auditory] is statistically significant and positive
## (beta = 0.74, 95% CI [0.36, 1.12], t(9124) = 3.83, p < .001; Std. beta = -0.59,
## 95% CI [-0.64, -0.54])
##   - The effect of RSN [H_MotorAssociation] is statistically significant and
## positive (beta = 0.47, 95% CI [0.09, 0.85], t(9124) = 2.42, p = 0.015; Std.
## beta = -0.75, 95% CI [-0.80, -0.70])
##   - The effect of RSN [H_PosteriorParietal] is statistically significant and
## positive (beta = 0.82, 95% CI [0.44, 1.20], t(9124) = 4.25, p < .001; Std. beta
## = -0.57, 95% CI [-0.62, -0.52])
##   - The effect of RSN [H_Somatosensory] is statistically significant and positive
## (beta = 0.74, 95% CI [0.36, 1.12], t(9124) = 3.81, p < .001; Std. beta = -0.71,
## 95% CI [-0.76, -0.66])
##   - The effect of RSN [H_DorsalVisualStream] is statistically significant and
## positive (beta = 0.88, 95% CI [0.50, 1.26], t(9124) = 4.54, p < .001; Std. beta
## = -1.03, 95% CI [-1.08, -0.98])
##   - The effect of RSN [H_PosteriorCingulateCortex] is statistically significant
## and positive (beta = 0.77, 95% CI [0.39, 1.15], t(9124) = 3.97, p < .001; Std.
## beta = -1.14, 95% CI [-1.19, -1.09])
##   - The effect of RSN [H_FrontalPole] is statistically significant and positive
## (beta = 0.67, 95% CI [0.29, 1.05], t(9124) = 3.48, p < .001; Std. beta = -1.03,
## 95% CI [-1.08, -0.98])
##   - The effect of RSN [H_Midbrain] is statistically significant and positive
## (beta = 0.72, 95% CI [0.34, 1.10], t(9124) = 3.72, p < .001; Std. beta = -1.31,
## 95% CI [-1.36, -1.26])
##   - The effect of RSN [H_DorsolateralPrefrontal] is statistically significant and
## positive (beta = 0.73, 95% CI [0.35, 1.11], t(9124) = 3.79, p < .001; Std. beta
## = -1.17, 95% CI [-1.22, -1.12])
##   - The effect of RSN [H_Cerebellum] is statistically non-significant and
## positive (beta = 0.26, 95% CI [-0.12, 0.63], t(9124) = 1.32, p = 0.187; Std.
## beta = -1.75, 95% CI [-1.80, -1.70])
##   - The effect of scan age × birth age is statistically significant and negative
## (beta = -7.39e-04, 95% CI [-9.81e-04, -4.97e-04], t(9124) = -5.99, p < .001;
## Std. beta = -0.10, 95% CI [-0.14, -0.07])
##   - The effect of scan age × RSN [H_MotorLateral] is statistically significant
## and negative (beta = -0.02, 95% CI [-0.02, -5.41e-03], t(9124) = -3.06, p =
## 0.002; Std. beta = -0.12, 95% CI [-0.19, -0.05])
##   - The effect of scan age × RSN [H_Visual] is statistically significant and
## negative (beta = -0.02, 95% CI [-0.02, -5.70e-03], t(9124) = -3.12, p = 0.002;
## Std. beta = -0.07, 95% CI [-0.14, -7.25e-04])
##   - The effect of scan age × RSN [H_Auditory] is statistically significant and
## negative (beta = -0.02, 95% CI [-0.03, -0.01], t(9124) = -4.81, p < .001; Std.
## beta = -0.17, 95% CI [-0.24, -0.10])
##   - The effect of scan age × RSN [H_MotorAssociation] is statistically
## significant and negative (beta = -0.02, 95% CI [-0.03, -6.55e-03], t(9124) =
## -3.30, p < .001; Std. beta = -0.34, 95% CI [-0.41, -0.27])
##   - The effect of scan age × RSN [H_PosteriorParietal] is statistically
## significant and negative (beta = -0.02, 95% CI [-0.03, -0.01], t(9124) = -4.22,
## p < .001; Std. beta = -1.32e-03, 95% CI [-0.07, 0.07])
##   - The effect of scan age × RSN [H_Somatosensory] is statistically significant
## and negative (beta = -0.02, 95% CI [-0.03, -0.01], t(9124) = -4.51, p < .001;
## Std. beta = -0.18, 95% CI [-0.25, -0.11])
##   - The effect of scan age × RSN [H_DorsalVisualStream] is statistically
```

```
## significant and negative (beta = -0.03, 95% CI [-0.04, -0.02], t(9124) = -5.34,
## p < .001; Std. beta = -0.26, 95% CI [-0.33, -0.19])
##   - The effect of scan age × RSN [H_PosteriorCingulateCortex] is statistically
## significant and negative (beta = -0.02, 95% CI [-0.03, -0.01], t(9124) = -4.59,
## p < .001; Std. beta = -0.36, 95% CI [-0.43, -0.29])
##   - The effect of scan age × RSN [H_FrontalPole] is statistically significant and
## negative (beta = -0.02, 95% CI [-0.03, -0.01], t(9124) = -4.50, p < .001; Std.
## beta = -0.47, 95% CI [-0.54, -0.40])
##   - The effect of scan age × RSN [H_Midbrain] is statistically significant and
## negative (beta = -0.02, 95% CI [-0.03, -0.01], t(9124) = -4.20, p < .001; Std.
## beta = -0.55, 95% CI [-0.61, -0.48])
##   - The effect of scan age × RSN [H_DorsolateralPrefrontal] is statistically
## significant and negative (beta = -0.02, 95% CI [-0.03, -0.01], t(9124) = -4.91,
## p < .001; Std. beta = -0.44, 95% CI [-0.51, -0.37])
##   - The effect of scan age × RSN [H_Cerebellum] is statistically significant and
## negative (beta = -0.02, 95% CI [-0.03, -9.08e-03], t(9124) = -3.81, p < .001;
## Std. beta = -0.62, 95% CI [-0.69, -0.55])
##   - The effect of birth age × RSN [H_MotorLateral] is statistically
## non-significant and negative (beta = -7.36e-03, 95% CI [-0.02, 4.33e-03],
## t(9124) = -1.23, p = 0.217; Std. beta = 0.18, 95% CI [0.12, 0.24])
##   - The effect of birth age × RSN [H_Visual] is statistically significant and
## negative (beta = -0.01, 95% CI [-0.02, -1.11e-03], t(9124) = -2.15, p = 0.032;
## Std. beta = 0.04, 95% CI [-0.01, 0.10])
##   - The effect of birth age × RSN [H_Auditory] is statistically significant and
## negative (beta = -0.02, 95% CI [-0.03, -4.07e-03], t(9124) = -2.64, p = 0.008;
## Std. beta = 0.14, 95% CI [0.08, 0.20])
##   - The effect of birth age × RSN [H_MotorAssociation] is statistically
## non-significant and negative (beta = -2.97e-03, 95% CI [-0.01, 8.71e-03],
## t(9124) = -0.50, p = 0.618; Std. beta = 0.09, 95% CI [0.04, 0.15])
##   - The effect of birth age × RSN [H_PosteriorParietal] is statistically
## significant and negative (beta = -0.02, 95% CI [-0.04, -0.01], t(9124) = -4.00,
## p < .001; Std. beta = -0.07, 95% CI [-0.13, -0.01])
##   - The effect of birth age × RSN [H_Somatosensory] is statistically significant
## and negative (beta = -0.02, 95% CI [-0.03, -3.96e-03], t(9124) = -2.62, p =
## 0.009; Std. beta = 0.07, 95% CI [8.08e-03, 0.12])
##   - The effect of birth age × RSN [H_DorsalVisualStream] is statistically
## significant and negative (beta = -0.02, 95% CI [-0.03, -6.02e-03], t(9124) =
## -2.97, p = 0.003; Std. beta = 0.05, 95% CI [-0.01, 0.10])
##   - The effect of birth age × RSN [H_PosteriorCingulateCortex] is statistically
## non-significant and negative (beta = -0.01, 95% CI [-0.02, 1.69e-04], t(9124) =
## -1.93, p = 0.053; Std. beta = -4.91e-03, 95% CI [-0.06, 0.05])
##   - The effect of birth age × RSN [H_FrontalPole] is statistically
## non-significant and negative (beta = -4.65e-03, 95% CI [-0.02, 7.03e-03],
## t(9124) = -0.78, p = 0.435; Std. beta = 0.09, 95% CI [0.03, 0.15])
##   - The effect of birth age × RSN [H_Midbrain] is statistically non-significant
## and negative (beta = -4.22e-03, 95% CI [-0.02, 7.47e-03], t(9124) = -0.71, p =
## 0.480; Std. beta = -0.05, 95% CI [-0.11, 2.70e-03])
##   - The effect of birth age × RSN [H_DorsolateralPrefrontal] is statistically
## non-significant and negative (beta = -7.82e-03, 95% CI [-0.02, 3.86e-03],
## t(9124) = -1.31, p = 0.189; Std. beta = 0.10, 95% CI [0.04, 0.15])
##   - The effect of birth age × RSN [H_Cerebellum] is statistically non-significant
## and positive (beta = 9.61e-03, 95% CI [-2.07e-03, 0.02], t(9124) = 1.61, p =
## 0.107; Std. beta = 0.31, 95% CI [0.26, 0.37])
##   - The effect of (scan age × birth age) × RSN [H_MotorLateral] is statistically
```

```
## significant and positive (beta = 2.99e-04, 95% CI [9.78e-06, 5.89e-04], t(9124)
## = 2.03, p = 0.043; Std. beta = 0.04, 95% CI [1.35e-03, 0.08])
##   - The effect of (scan age × birth age) × RSN [H_Visual] is statistically
## significant and positive (beta = 3.51e-04, 95% CI [6.12e-05, 6.40e-04], t(9124)
## = 2.37, p = 0.018; Std. beta = 0.05, 95% CI [8.45e-03, 0.09])
##   - The effect of (scan age × birth age) × RSN [H_Auditory] is statistically
## significant and positive (beta = 4.84e-04, 95% CI [1.95e-04, 7.74e-04], t(9124)
## = 3.28, p = 0.001; Std. beta = 0.07, 95% CI [0.03, 0.11])
##   - The effect of (scan age × birth age) × RSN [H_MotorAssociation] is
## statistically non-significant and positive (beta = 1.34e-04, 95% CI [-1.55e-04,
## 4.24e-04], t(9124) = 0.91, p = 0.364; Std. beta = 0.02, 95% CI [-0.02, 0.06])
##   - The effect of (scan age × birth age) × RSN [H_PosteriorParietal] is
## statistically significant and positive (beta = 5.56e-04, 95% CI [2.67e-04,
## 8.46e-04], t(9124) = 3.77, p < .001; Std. beta = 0.08, 95% CI [0.04, 0.12])
##   - The effect of (scan age × birth age) × RSN [H_Somatosensory] is statistically
## significant and positive (beta = 4.35e-04, 95% CI [1.46e-04, 7.25e-04], t(9124)
## = 2.95, p = 0.003; Std. beta = 0.06, 95% CI [0.02, 0.10])
##   - The effect of (scan age × birth age) × RSN [H_DorsalVisualStream] is
## statistically significant and positive (beta = 4.75e-04, 95% CI [1.85e-04,
## 7.64e-04], t(9124) = 3.21, p = 0.001; Std. beta = 0.07, 95% CI [0.03, 0.11])
##   - The effect of (scan age × birth age) × RSN [H_PosteriorCingulateCortex] is
## statistically non-significant and positive (beta = 2.87e-04, 95% CI [-2.46e-06,
## 5.77e-04], t(9124) = 1.94, p = 0.052; Std. beta = 0.04, 95% CI [-3.39e-04,
## 0.08])
##   - The effect of (scan age × birth age) × RSN [H_FrontalPole] is statistically
## non-significant and positive (beta = 1.75e-04, 95% CI [-1.15e-04, 4.64e-04],
## t(9124) = 1.18, p = 0.237; Std. beta = 0.02, 95% CI [-0.02, 0.06])
##   - The effect of (scan age × birth age) × RSN [H_Midbrain] is statistically
## non-significant and positive (beta = 7.20e-05, 95% CI [-2.18e-04, 3.61e-04],
## t(9124) = 0.49, p = 0.626; Std. beta = 9.93e-03, 95% CI [-0.03, 0.05])
##   - The effect of (scan age × birth age) × RSN [H_DorsolateralPrefrontal] is
## statistically non-significant and positive (beta = 2.58e-04, 95% CI [-3.14e-05,
## 5.48e-04], t(9124) = 1.75, p = 0.081; Std. beta = 0.04, 95% CI [-4.34e-03,
## 0.08])
##   - The effect of (scan age × birth age) × RSN [H_Cerebellum] is statistically
## non-significant and negative (beta = -4.59e-05, 95% CI [-3.35e-04, 2.44e-04],
## t(9124) = -0.31, p = 0.756; Std. beta = -6.34e-03, 95% CI [-0.05, 0.03])
##
## Standardized parameters were obtained by fitting the model on a standardized
## version of the dataset. 95% Confidence Intervals (CIs) and p-values were
## computed using a Wald t-distribution approximation.
```

Report in just the Motor Medial as example:

```
## We fitted a linear mixed model (estimated using ML and nloptwrap optimizer) to
## predict H_MotorMedial with scan_age and birth_age (formula: H_MotorMedial ~
## scan_age * birth_age). The model included Subject as random effect (formula: ~1
## | Subject). The model's total explanatory power is substantial (conditional R2
## = 0.59) and the part related to the fixed effects alone (marginal R2) is of
## 0.38. The model's intercept, corresponding to scan_age = 0 and birth_age = 0,
## is at -0.96 (95% CI [-1.58, -0.33], t(700) = -3.02, p = 0.003). Within this
## model:
##
##   - The effect of scan age is statistically significant and positive (beta =
## 0.04, 95% CI [0.02, 0.05], t(700) = 4.81, p < .001; Std. beta = 0.53, 95% CI
```

```
## [0.44, 0.62])
##   - The effect of birth age is statistically non-significant and positive (beta =
## 0.02, 95% CI [-1.64e-04, 0.04], t(700) = 1.95, p = 0.052; Std. beta = 0.06, 95%
## CI [-0.01, 0.14])
##   - The effect of scan age × birth age is statistically non-significant and
## negative (beta = -4.35e-04, 95% CI [-9.12e-04, 4.25e-05], t(700) = -1.79, p =
## 0.074; Std. beta = -0.05, 95% CI [-0.10, 4.64e-03])
##
## Standardized parameters were obtained by fitting the model on a standardized
## version of the dataset. 95% Confidence Intervals (CIs) and p-values were
## computed using a Wald t-distribution approximation.
```

### Just Preterm Group:

The preterm group will be assessed by comparison of the preterm scans to term age (TEA) scans and between the VPT and MPT groups.

First, let's remove the THC group from the data (not shown)

Double checking that we are only analyzing preterm infants.

This number should be 0: 0

#### Group Numbers

Number of preterm and TEA scans.

| Observations | GAfactor2 | n |
|---:|---|---:|
| 1 | PT | 146 |
| 1 | T | 102 |
| 2 | T | 2 |

2 preterm infants scanned twice at TEA (CC00098AN17 and CC00688XX21).

#### Preterm vs TEA Comparison

Will now compare the preterm group at preterm age (35 weeks) and at TEA (41 weeks) in the grey matter, white matter, combined RSN, and individual RSN.
These ages were used as they the median values of the preterm group at preterm and TEA scan age (PMA).The estimated marginal means will be calculated. Linear mixed effects models are created with scan age.

##### Grey Matter H comparison Preterm vs TEA

Estimated Marginal Means:

```
##  scan_age emmean     SE  df lower.CL upper.CL
##      34.8  0.452 0.0043 249    0.444    0.461
##      41.0  0.539 0.0051 255    0.529    0.549
##
## Degrees-of-freedom method: kenward-roger
## Confidence level used: 0.95
```

Comparison:

```
##  contrast                     estimate      SE  df t.ratio p.value
##  scan_age41 - scan_age34.785   0.0862 0.00598 174  14.426  <.0001
##
## Degrees-of-freedom method: kenward-roger
```

Slopes / Trends:

```
##  birth_age scan_age.trend       SE  df lower.CL upper.CL
##       31.9         0.0139 0.000962 174    0.012   0.0158
##
## Degrees-of-freedom method: kenward-roger
## Confidence level used: 0.95
```

##### White Matter H comparison Preterm vs TEA

Estimated marginal means:

```
##  scan_age emmean      SE  df lower.CL upper.CL
##      34.8  0.432 0.00357 250    0.425    0.439
##      41.0  0.465 0.00424 255    0.456    0.473
##
## Degrees-of-freedom method: kenward-roger
## Confidence level used: 0.95
```

Contrast:

```
##  contrast                    estimate      SE  df t.ratio p.value
##  scan_age41 - scan_age34.785   0.0332 0.00501 177   6.637  <.0001
##
## Degrees-of-freedom method: kenward-roger
```

Slopes:

```
##  birth_age scan_age.trend       SE  df lower.CL upper.CL
##       31.9        0.00535 0.000806 177  0.00376  0.00694
##
## Degrees-of-freedom method: kenward-roger
## Confidence level used: 0.95
```

##### Combined RSN H comparison Preterm vs TEA

Estimated marginal Means

```
##  scan_age emmean      SE  df asymp.LCL asymp.UCL
##      34.8  0.451 0.00359 Inf     0.444     0.458
##      41.0  0.530 0.00386 Inf     0.523     0.538
##
## Degrees-of-freedom method: asymptotic
## Confidence level used: 0.95
```

Contrast

```
##  contrast                    estimate      SE  df z.ratio p.value
##  scan_age41 - scan_age34.785   0.0794 0.00293 Inf  27.067  <.0001
##
## Degrees-of-freedom method: asymptotic
```

Slopes:

```
##  birth_age scan_age.trend       SE  df asymp.LCL asymp.UCL
##       31.9         0.0128 0.000472 Inf    0.0118    0.0137
##
## Degrees-of-freedom method: asymptotic
## Confidence level used: 0.95
```

#### Individual RSN H comparison Preterm vs TEA

Estimated marginal means

| scan_age | RSN | emmean | SE | df | asymp.LCL | asymp.UCL |
|---|---|---|---|---|---|---|
| 34.785 | H_MotorMedial | 0.4972965 | 0.0050732 | Inf | 0.4873533 | 0.5072398 |
| 41.000 | H_MotorMedial | 0.6513978 | 0.0056894 | Inf | 0.6402468 | 0.6625489 |
| 34.785 | H_MotorLateral | 0.4725454 | 0.0050732 | Inf | 0.4626021 | 0.4824887 |
| 41.000 | H_MotorLateral | 0.5912447 | 0.0056894 | Inf | 0.5800936 | 0.6023958 |
| 34.785 | H_Visual | 0.4441406 | 0.0050732 | Inf | 0.4341974 | 0.4540839 |
| 41.000 | H_Visual | 0.5579841 | 0.0056894 | Inf | 0.5468330 | 0.5691351 |
| 34.785 | H_Auditory | 0.4582713 | 0.0050732 | Inf | 0.4483281 | 0.4682146 |
| 41.000 | H_Auditory | 0.5562771 | 0.0056894 | Inf | 0.5451260 | 0.5674282 |
| 34.785 | H_MotorAssociation | 0.4628031 | 0.0050732 | Inf | 0.4528598 | 0.4727464 |
| 41.000 | H_MotorAssociation | 0.5352476 | 0.0056894 | Inf | 0.5240965 | 0.5463987 |
| 34.785 | H_PosteriorParietal | 0.4693499 | 0.0050732 | Inf | 0.4594066 | 0.4792932 |
| 41.000 | H_PosteriorParietal | 0.5868875 | 0.0056894 | Inf | 0.5757364 | 0.5980385 |
| 34.785 | H_Somatosensory | 0.4577165 | 0.0050732 | Inf | 0.4477732 | 0.4676598 |
| 41.000 | H_Somatosensory | 0.5476931 | 0.0056894 | Inf | 0.5365421 | 0.5588442 |
| 34.785 | H_DorsalVisualStream | 0.4421055 | 0.0050732 | Inf | 0.4321623 | 0.4520488 |
| 41.000 | H_DorsalVisualStream | 0.5092567 | 0.0056894 | Inf | 0.4981056 | 0.5204078 |
| 34.785 | H_PosteriorCingulateCortex | 0.4464596 | 0.0050732 | Inf | 0.4365163 | 0.4564029 |
| 41.000 | H_PosteriorCingulateCortex | 0.5000086 | 0.0056894 | Inf | 0.4888575 | 0.5111596 |
| 34.785 | H_FrontalPole | 0.4580443 | 0.0050732 | Inf | 0.4481010 | 0.4679876 |
| 41.000 | H_FrontalPole | 0.5007079 | 0.0056894 | Inf | 0.4895569 | 0.5118590 |
| 34.785 | H_Midbrain | 0.4570630 | 0.0050732 | Inf | 0.4471197 | 0.4670062 |
| 41.000 | H_Midbrain | 0.4832679 | 0.0056894 | Inf | 0.4721168 | 0.4944190 |
| 34.785 | H_DorsolateralPrefrontal | 0.4397283 | 0.0050732 | Inf | 0.4297850 | 0.4496715 |
| 41.000 | H_DorsolateralPrefrontal | 0.4819674 | 0.0056894 | Inf | 0.4708164 | 0.4931185 |
| 34.785 | H_Cerebellum | 0.3582271 | 0.0050732 | Inf | 0.3482839 | 0.3681704 |
| 41.000 | H_Cerebellum | 0.3850083 | 0.0056894 | Inf | 0.3738572 | 0.3961594 |

Contrast:

| contrast | RSN | estimate | SE | df | z.ratio | p.value |
|---|---|---|---|---|---|---|
| scan_age41 - scan_age34.785 | H_MotorMedial | 0.1541013 | 0.0057422 | Inf | 26.836827 | 0.0e+00 |
| scan_age41 - scan_age34.785 | H_MotorLateral | 0.1186993 | 0.0057422 | Inf | 20.671544 | 0.0e+00 |
| scan_age41 - scan_age34.785 | H_Visual | 0.1138434 | 0.0057422 | Inf | 19.825895 | 0.0e+00 |
| scan_age41 - scan_age34.785 | H_Auditory | 0.0980058 | 0.0057422 | Inf | 17.067760 | 0.0e+00 |
| scan_age41 - scan_age34.785 | H_MotorAssociation | 0.0724445 | 0.0057422 | Inf | 12.616249 | 0.0e+00 |
| scan_age41 - scan_age34.785 | H_PosteriorParietal | 0.1175376 | 0.0057422 | Inf | 20.469229 | 0.0e+00 |
| scan_age41 - scan_age34.785 | H_Somatosensory | 0.0899766 | 0.0057422 | Inf | 15.669479 | 0.0e+00 |
| scan_age41 - scan_age34.785 | H_DorsalVisualStream | 0.0671512 | 0.0057422 | Inf | 11.694409 | 0.0e+00 |
| scan_age41 - scan_age34.785 | H_PosteriorCingulateCortex | 0.0535490 | 0.0057422 | Inf | 9.325581 | 0.0e+00 |
| scan_age41 - scan_age34.785 | H_FrontalPole | 0.0426636 | 0.0057422 | Inf | 7.429896 | 0.0e+00 |
| scan_age41 - scan_age34.785 | H_Midbrain | 0.0262049 | 0.0057422 | Inf | 4.563604 | 6.2e-06 |
| scan_age41 - scan_age34.785 | H_DorsolateralPrefrontal | 0.0422392 | 0.0057422 | Inf | 7.355972 | 0.0e+00 |
| scan_age41 - scan_age34.785 | H_Cerebellum | 0.0267812 | 0.0057422 | Inf | 4.663952 | 6.2e-06 |

##### RSN Slopes Preterm to TEA

Determining which RSN has the greatest increase with age in the preterm group.

Trends in descending order:

| RSN | scan_age.trend | std.error | df | statistic | p.value |
|---|---|---|---|---|---|
| H_MotorMedial | 0.0247951 | 0.0009239 | Inf | 26.836827 | 0.0e+00 |
| H_MotorLateral | 0.0190988 | 0.0009239 | Inf | 20.671544 | 0.0e+00 |
| H_PosteriorParietal | 0.0189119 | 0.0009239 | Inf | 20.469229 | 0.0e+00 |
| H_Visual | 0.0183175 | 0.0009239 | Inf | 19.825895 | 0.0e+00 |
| H_Auditory | 0.0157692 | 0.0009239 | Inf | 17.067760 | 0.0e+00 |
| H_Somatosensory | 0.0144773 | 0.0009239 | Inf | 15.669479 | 0.0e+00 |
| H_MotorAssociation | 0.0116564 | 0.0009239 | Inf | 12.616249 | 0.0e+00 |
| H_DorsalVisualStream | 0.0108047 | 0.0009239 | Inf | 11.694409 | 0.0e+00 |
| H_PosteriorCingulateCortex | 0.0086161 | 0.0009239 | Inf | 9.325581 | 0.0e+00 |
| H_FrontalPole | 0.0068646 | 0.0009239 | Inf | 7.429896 | 0.0e+00 |
| H_DorsolateralPrefrontal | 0.0067963 | 0.0009239 | Inf | 7.355972 | 0.0e+00 |
| H_Cerebellum | 0.0043091 | 0.0009239 | Inf | 4.663952 | 3.1e-06 |
| H_Midbrain | 0.0042164 | 0.0009239 | Inf | 4.563604 | 5.0e-06 |

##### Investigating when RSNs Surpass Ventricle

Let's treat H in ventricles as a 'noise ground floor'

When do various RSN's surpass the ventricle?

Scan age (weeks) when RSN crosses Ventricles:

| brain_regions | intersect |
|---|---|
| H_Midbrain | 26.00007 |
| H_FrontalPole | 29.43576 |
| H_MotorAssociation | 31.90526 |
| H_MotorMedial | 32.06010 |
| H_MotorLateral | 32.64073 |
| H_PosteriorParietal | 32.81864 |
| H_Somatosensory | 33.07707 |
| H_Auditory | 33.20849 |
| H_PosteriorCingulateCortex | 33.30734 |
| H_DorsalVisualStream | 34.30565 |
| H_Visual | 34.40928 |
| H_DorsolateralPrefrontal | 34.43257 |
| H_Cerebellum | 45.99993 |

#### VPT vs MPT Preterm Group Comparison

Assessing the very preterm and moderately born preterm groups separately to determine if H values between groups are significantly different at preterm age and term age. The overall slopes will be compared in the regions. Only testing at preterm age since we will be doing the term age comparison later with the THC.

##### Grey matter

Estimated marginal means:

| scan_age | birth_age | emmean | SE | df | lower.CL | upper.CL |
|---|---|---|---|---|---|---|
| 35 | 29 | 0.4478977 | 0.0046357 | 178.9145 | 0.4387501 | 0.4570453 |
| 35 | 35 | 0.4635938 | 0.0061066 | 256.0643 | 0.4515682 | 0.4756195 |

Contrast:

| contrast | estimate | SE | df | t.ratio | p.value |
|---|---|---|---|---|---|
| scan_age35 birth_age35 - scan_age35 birth_age29 | 0.0156962 | 0.0067856 | 232.5648 | 2.313163 | 0.0215871 |

Slopes:

| birth_age | scan_age.trend | p.value |
|---:|---:|---:|
| 29 | 0.0155961 | 0 |
| 35 | 0.0119698 | 0 |

Contrast of slopes

| contrast | estimate | SE | df | t.ratio | p.value |
|:---|---:|---:|---:|---:|---:|
| birth_age35 - birth_age29 | -0.0036263 | 0.0013347 | 178.2051 | -2.716843 | 0.0072407 |

##### White Matter

Estimated marginal means:

| scan_age | birth_age | emmean | SE | df | lower.CL | upper.CL |
|---:|---:|---:|---:|---:|---:|---:|
| 35 | 29 | 0.4207576 | 0.0038298 | 177.9728 | 0.4131999 | 0.4283154 |
| 35 | 35 | 0.4458795 | 0.0050813 | 256.0573 | 0.4358730 | 0.4558859 |

Contrast:

| contrast | estimate | SE | df | t.ratio | p.value |
|:---|---:|---:|---:|---:|---:|
| scan_age35 birth_age35 - scan_age35 birth_age29 | 0.0251218 | 0.0056268 | 232.5068 | 4.46467 | 1.25e-05 |

Slopes / trends:

| birth_age | scan_age.trend | p.value |
|---:|---:|---:|
| 29 | 0.0071093 | 0.0000000 |
| 35 | 0.0034056 | 0.0051771 |

Contrast of trends:

| contrast | estimate | SE | df | [illegible] |
|:---|---:|---:|---:|---:|
| scan_age36.917380952381 birth_age35 - scan_age36.917380952381 birth_age29 | -0.0037037 | 0.0011184 | 181.0012 | -3[illegible] |

##### Combined RSN

Estimated marginal means:

| scan_age | birth_age | emmean | SE | df | asymp.LCL | asymp.UCL |
|---:|---:|---:|---:|---:|---:|---:|
| 35 | 29 | 0.4453484 | 0.0044423 | Inf | 0.4366418 | 0.4540551 |
| 35 | 35 | 0.4626926 | 0.0047970 | Inf | 0.4532907 | 0.4720945 |

Contrast:

| contrast | estimate | SE | df | z.ratio | p.value |
|:---|---:|---:|---:|---:|---:|
| scan_age35 birth_age35 - scan_age35 birth_age29 | 0.0173441 | 0.0058805 | Inf | 2.949418 | 0.0031837 |

Slopes

| birth_age | scan_age.trend | p.value |
|---:|---:|---:|
| 29 | 0.0143188 | 0 |
| 35 | 0.0110557 | 0 |

Contrast of slopes:

| contrast | estimate | SE | df | z.rat[illegible] |
|:---|---:|---:|---:|---:|
| scan_age36.917380952381 birth_age35 - scan_age36.917380952381 birth_age29 | -0.0032631 | 0.0006613 | Inf | -4.93437[illegible] |

##### Individual RSNs

Emmeans:

| scan_age | birth_age | RSN | emmean | SE | df | asymp.LCL | asymp.UCL |
|---:|---:|---|---:|---:|---:|---:|---:|
| 35 | 29 | H_MotorMedial | 0.5012853 | 0.0058739 | Inf | 0.4897726 | 0.5127979 |
| 35 | 35 | H_MotorMedial | 0.5041095 | 0.0069984 | Inf | 0.4903929 | 0.5178260 |
| 35 | 29 | H_MotorLateral | 0.4679305 | 0.0058739 | Inf | 0.4564179 | 0.4794432 |
| 35 | 35 | H_MotorLateral | 0.4862813 | 0.0069984 | Inf | 0.4725648 | 0.4999978 |
| 35 | 29 | H_Visual | 0.4439999 | 0.0058739 | Inf | 0.4324873 | 0.4555126 |
| 35 | 35 | H_Visual | 0.4525828 | 0.0069984 | Inf | 0.4388663 | 0.4662993 |
| 35 | 29 | H_Auditory | 0.4538910 | 0.0058739 | Inf | 0.4423783 | 0.4654037 |
| 35 | 35 | H_Auditory | 0.4702419 | 0.0069984 | Inf | 0.4565254 | 0.4839585 |
| 35 | 29 | H_MotorAssociation | 0.4570128 | 0.0058739 | Inf | 0.4455001 | 0.4685255 |
| 35 | 35 | H_MotorAssociation | 0.4744699 | 0.0069984 | Inf | 0.4607534 | 0.4881865 |
| 35 | 29 | H_PosteriorParietal | 0.4766016 | 0.0058739 | Inf | 0.4650890 | 0.4881143 |
| 35 | 35 | H_PosteriorParietal | 0.4698984 | 0.0069984 | Inf | 0.4561819 | 0.4836150 |
| 35 | 29 | H_Somatosensory | 0.4547554 | 0.0058739 | Inf | 0.4432428 | 0.4662681 |
| 35 | 35 | H_Somatosensory | 0.4675356 | 0.0069984 | Inf | 0.4538190 | 0.4812521 |
| 35 | 29 | H_DorsalVisualStream | 0.4372332 | 0.0058739 | Inf | 0.4257206 | 0.4487459 |
| 35 | 35 | H_DorsalVisualStream | 0.4523734 | 0.0069984 | Inf | 0.4386569 | 0.4660899 |
| 35 | 29 | H_PosteriorCingulateCortex | 0.4406951 | 0.0058739 | Inf | 0.4291824 | 0.4522077 |
| 35 | 35 | H_PosteriorCingulateCortex | 0.4567226 | 0.0069984 | Inf | 0.4430060 | 0.4704391 |
| 35 | 29 | H_FrontalPole | 0.4460689 | 0.0058739 | Inf | 0.4345563 | 0.4575816 |
| 35 | 35 | H_FrontalPole | 0.4743727 | 0.0069984 | Inf | 0.4606562 | 0.4880893 |
| 35 | 29 | H_Midbrain | 0.4493370 | 0.0058739 | Inf | 0.4378244 | 0.4608497 |
| 35 | 35 | H_Midbrain | 0.4675012 | 0.0069984 | Inf | 0.4537847 | 0.4812178 |
| 35 | 29 | H_DorsolateralPrefrontal | 0.4283563 | 0.0058739 | Inf | 0.4168437 | 0.4398690 |
| 35 | 35 | H_DorsolateralPrefrontal | 0.4553595 | 0.0069984 | Inf | 0.4416430 | 0.4690761 |
| 35 | 29 | H_Cerebellum | 0.3328869 | 0.0058739 | Inf | 0.3213742 | 0.3443996 |
| 35 | 35 | H_Cerebellum | 0.3881566 | 0.0069984 | Inf | 0.3744401 | 0.4018732 |

Contrast

| contrast | RSN | estimate | p.value |
|---|---|---:|---:|
| scan_age35 birth_age35 - scan_age35 birth_age29 | H_MotorMedial | 0.0028242 | 0.8747375 |
| scan_age35 birth_age35 - scan_age35 birth_age29 | H_MotorLateral | 0.0183508 | 0.2413679 |
| scan_age35 birth_age35 - scan_age35 birth_age29 | H_Visual | 0.0085829 | 0.8747375 |
| scan_age35 birth_age35 - scan_age35 birth_age29 | H_Auditory | 0.0163509 | 0.3116070 |
| scan_age35 birth_age35 - scan_age35 birth_age29 | H_MotorAssociation | 0.0174571 | 0.2555396 |
| scan_age35 birth_age35 - scan_age35 birth_age29 | H_PosteriorParietal | -0.0067032 | 0.8747375 |
| scan_age35 birth_age35 - scan_age35 birth_age29 | H_Somatosensory | 0.0127802 | 0.4652634 |
| scan_age35 birth_age35 - scan_age35 birth_age29 | H_DorsalVisualStream | 0.0151401 | 0.3141289 |
| scan_age35 birth_age35 - scan_age35 birth_age29 | H_PosteriorCingulateCortex | 0.0160275 | 0.3116070 |
| scan_age35 birth_age35 - scan_age35 birth_age29 | H_FrontalPole | 0.0283038 | **0.0060627** |
| scan_age35 birth_age35 - scan_age35 birth_age29 | H_Midbrain | 0.0181642 | 0.2413679 |
| scan_age35 birth_age35 - scan_age35 birth_age29 | H_DorsolateralPrefrontal | 0.0270032 | **0.0099676** |
| scan_age35 birth_age35 - scan_age35 birth_age29 | H_Cerebellum | 0.0552697 | **0.0000000** |

Slopes:

| birth_age | RSN | scan_age.trend | std.error | df | statistic | p.value |
|---|---|---|---|---|---|---|
| 29 | H_MotorMedial | 0.0255811 | 0.0008240 | Inf | 31.044486 | 0.0000000 |
| 35 | H_MotorMedial | 0.0239271 | 0.0013821 | Inf | 17.311585 | 0.0000000 |
| 29 | H_MotorLateral | 0.0195830 | 0.0008240 | Inf | 23.765335 | 0.0000000 |
| 35 | H_MotorLateral | 0.0185643 | 0.0013821 | Inf | 13.431479 | 0.0000000 |
| 29 | H_Visual | 0.0208466 | 0.0008240 | Inf | 25.298749 | 0.0000000 |
| 35 | H_Visual | 0.0155250 | 0.0013821 | Inf | 11.232576 | 0.0000000 |
| 29 | H_Auditory | 0.0162608 | 0.0008240 | Inf | 19.733599 | 0.0000000 |
| 35 | H_Auditory | 0.0152265 | 0.0013821 | Inf | 11.016559 | 0.0000000 |
| 29 | H_MotorAssociation | 0.0133523 | 0.0008240 | Inf | 16.203961 | 0.0000000 |
| 35 | H_MotorAssociation | 0.0097838 | 0.0013821 | Inf | 7.078718 | 0.0000000 |
| 29 | H_PosteriorParietal | 0.0210348 | 0.0008240 | Inf | 25.527248 | 0.0000000 |
| 35 | H_PosteriorParietal | 0.0165678 | 0.0013821 | Inf | 11.987052 | 0.0000000 |
| 29 | H_Somatosensory | 0.0159920 | 0.0008240 | Inf | 19.407453 | 0.0000000 |
| 35 | H_Somatosensory | 0.0128048 | 0.0013821 | Inf | 9.264473 | 0.0000000 |
| 29 | H_DorsalVisualStream | 0.0131404 | 0.0008240 | Inf | 15.946853 | 0.0000000 |
| 35 | H_DorsalVisualStream | 0.0082256 | 0.0013821 | Inf | 5.951336 | 0.0000000 |
| 29 | H_PosteriorCingulateCortex | 0.0110032 | 0.0008240 | Inf | 13.353153 | 0.0000000 |
| 35 | H_PosteriorCingulateCortex | 0.0059803 | 0.0013821 | Inf | 4.326820 | 0.0000151 |
| 29 | H_FrontalPole | 0.0083948 | 0.0008240 | Inf | 10.187669 | 0.0000000 |
| 35 | H_FrontalPole | 0.0051751 | 0.0013821 | Inf | 3.744218 | 0.0001810 |
| 29 | H_Midbrain | 0.0063490 | 0.0008240 | Inf | 7.704944 | 0.0000000 |
| 35 | H_Midbrain | 0.0018617 | 0.0013821 | Inf | 1.346930 | 0.1780027 |
| 29 | H_DorsolateralPrefrontal | 0.0087735 | 0.0008240 | Inf | 10.647294 | 0.0000000 |
| 35 | H_DorsolateralPrefrontal | 0.0046131 | 0.0013821 | Inf | 3.337671 | 0.0008448 |
| 29 | H_Cerebellum | 0.0050752 | 0.0008240 | Inf | 6.159071 | 0.0000000 |
| 35 | H_Cerebellum | 0.0034633 | 0.0013821 | Inf | 2.505715 | 0.0122204 |

#### Plots:

##### GM, WM and Combined RSN Plot

Plot of H in the tissues in the preterm group. The horizontal line H=0.5

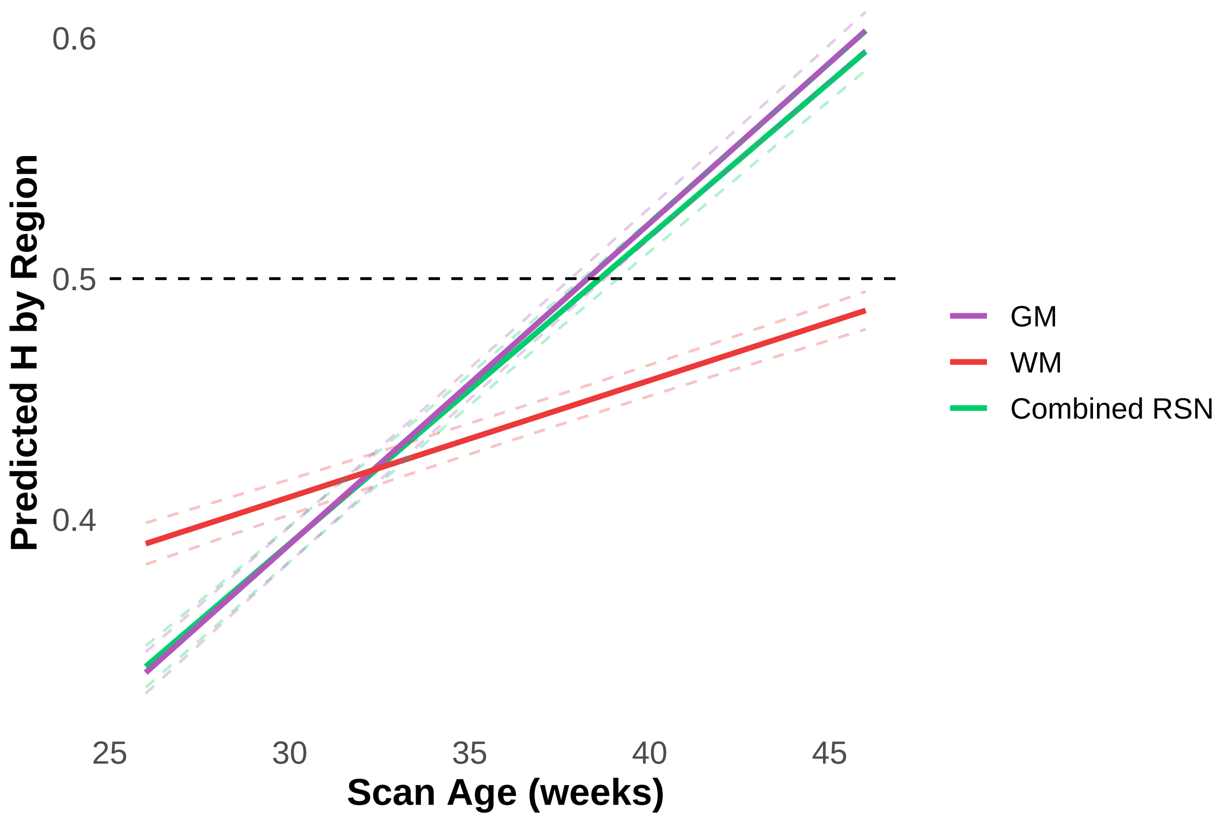


**Individual RSN Preterm Group Plot**

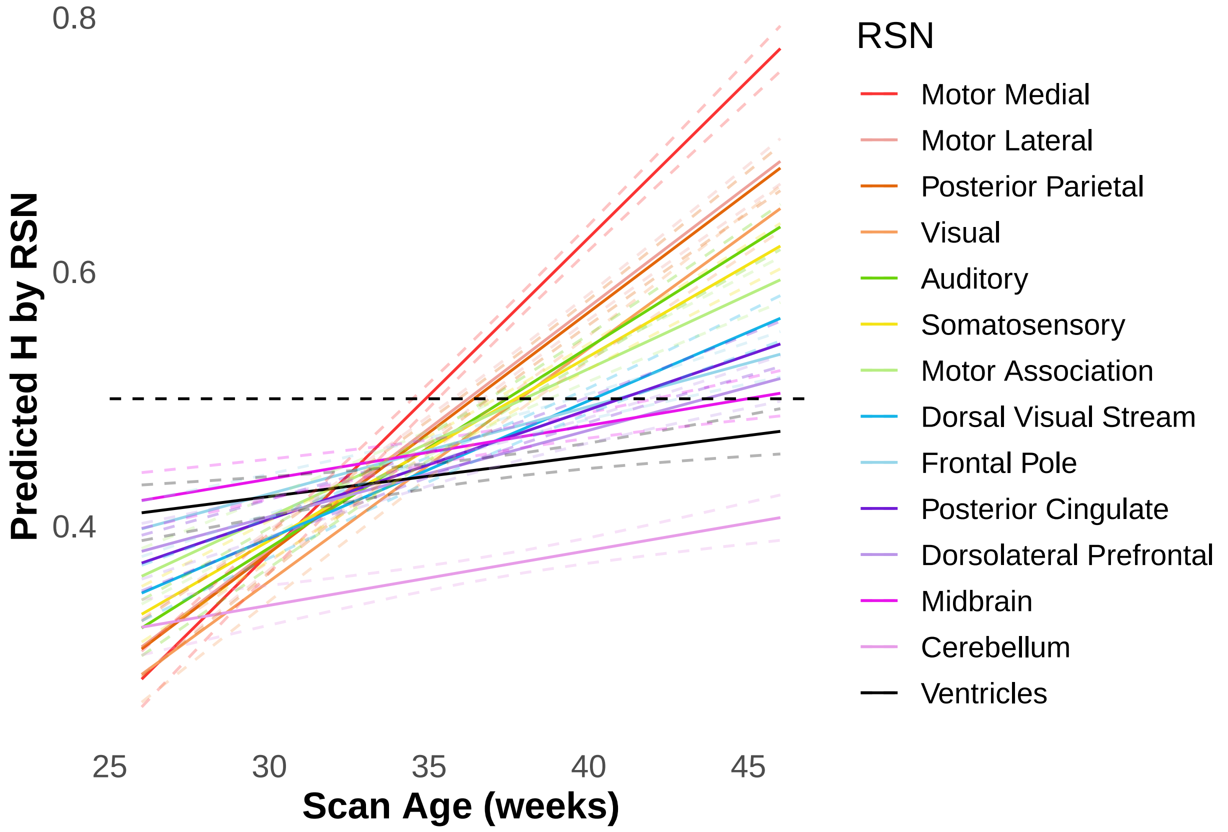

**VPT and MPT RSN Barchart at 35 weeks**

Comparing the preterm groups VPT and MPT at preterm age. Later will compare these groups at term to THC.

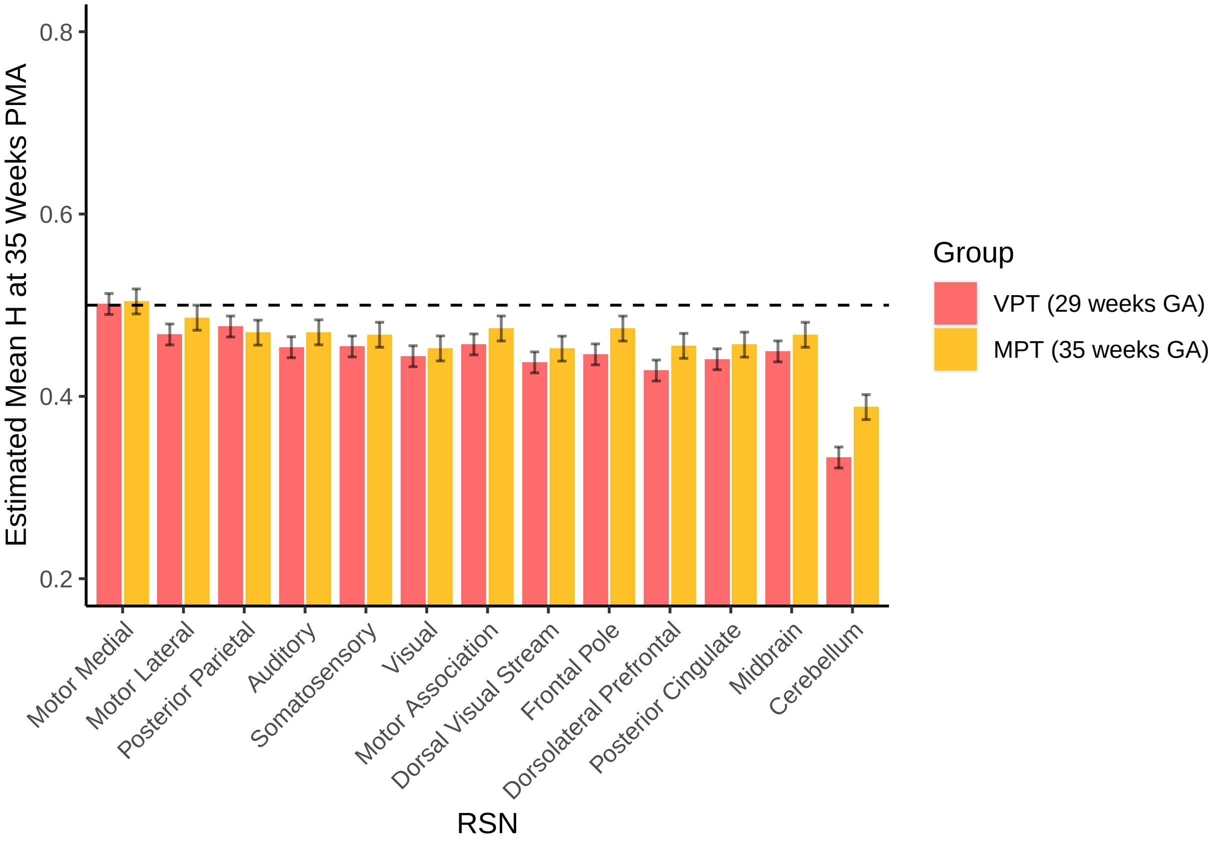


**VPT and MPT Tissue Barchart at 35 weeks**

Visual assessment of H values in the tissues compared in the groups. Need to add which are significant differences.

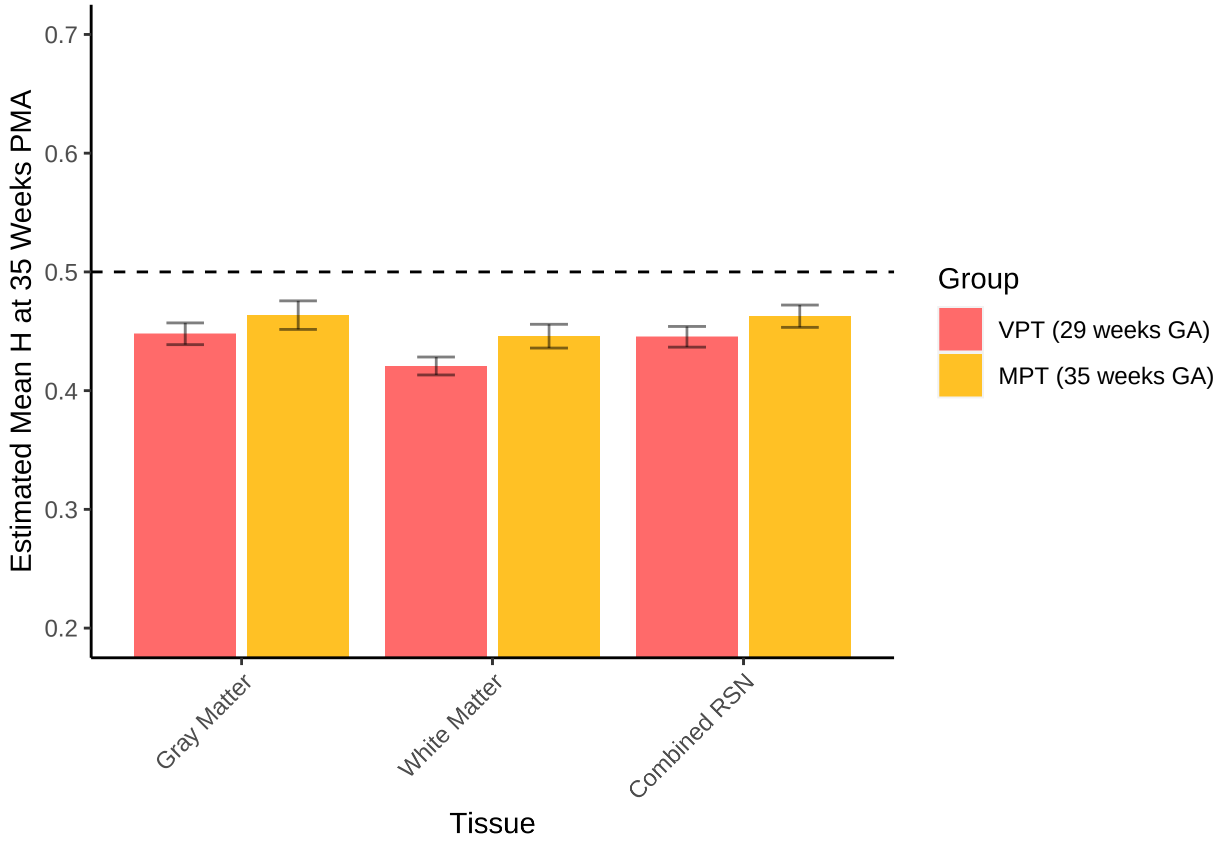


---

#### All Group Comparisons:

Do term born infants have greater H compared to preterm infants at scan age?

VPT at TEA vs MPT at TEA vs THC

Assessing the very preterm and moderately born preterm groups at term age to the term born controls.

Removing all the preterm scans.

Median age when looking at all scans > 37 weeks: 40.14285714

##### GM H Comparison VPT vs MPT vs THC

| scan_age | birth_age | emmean | SE | df | asymp.LCL | asymp.UCL |
|---:|---:|---:|---:|---|---:|---:|
| 41 | 29 | 0.5472094 | 0.0032403 | Inf | 0.5408585 | 0.5535603 |
| 41 | 35 | 0.5529663 | 0.0016189 | Inf | 0.5497933 | 0.5561393 |
| 41 | 41 | 0.5587231 | 0.0014488 | Inf | 0.5558835 | 0.5615628 |

Contrast:

| contrast | estimate | SE | df | z.ratio | p.value |
|---|---:|---:|---|---:|---:|
| scan_age41 birth_age35 - scan_age41 birth_age29 | 0.0057569 | 0.0019180 | Inf | 3.00158 | 0.0080575 |
| scan_age41 birth_age41 - scan_age41 birth_age29 | 0.0115138 | 0.0038359 | Inf | 3.00158 | 0.0080575 |
| scan_age41 birth_age41 - scan_age41 birth_age35 | 0.0057569 | 0.0019180 | Inf | 3.00158 | 0.0080575 |

#### WM H Comparison VPT vs MPT vs THC

| scan_age | birth_age | emmean | SE | df | asymp.LCL | asymp.UCL |
|---:|---:|---:|---:|---|---:|---:|
| 41 | 29 | 0.4432085 | 0.0018136 | Inf | 0.4396538 | 0.4467631 |
| 41 | 35 | 0.4738559 | 0.0009066 | Inf | 0.4720790 | 0.4756329 |
| 41 | 41 | 0.5045034 | 0.0008112 | Inf | 0.5029134 | 0.5060934 |

| contrast | estimate | SE | df | z.ratio | p.value |
|---|---:|---:|---|---:|---:|
| scan_age41 birth_age35 - scan_age41 birth_age29 | 0.0306475 | 0.0010732 | Inf | 28.55764 | 0 |
| scan_age41 birth_age41 - scan_age41 birth_age29 | 0.0612949 | 0.0021464 | Inf | 28.55764 | 0 |
| scan_age41 birth_age41 - scan_age41 birth_age35 | 0.0306475 | 0.0010732 | Inf | 28.55764 | 0 |

#### Combined RSN H Comparison VPT vs MPT vs THC

| scan_age | birth_age | emmean | SE | df | asymp.LCL | asymp.UCL |
|---:|---:|---:|---:|---|---:|---:|
| 41 | 29 | 0.5187201 | 0.0084967 | Inf | 0.5020668 | 0.5353734 |
| 41 | 35 | 0.5446592 | 0.0041346 | Inf | 0.5365557 | 0.5527628 |
| 41 | 41 | 0.5705984 | 0.0039776 | Inf | 0.5628024 | 0.5783944 |

| contrast | estimate | SE | df | z.ratio | p.value |
|---|---:|---:|---|---:|---:|
| scan_age41 birth_age35 - scan_age41 birth_age29 | 0.0259392 | 0.0051878 | Inf | 5.00002 | 1.7e-06 |
| scan_age41 birth_age41 - scan_age41 birth_age29 | 0.0518784 | 0.0103756 | Inf | 5.00002 | 1.7e-06 |
| scan_age41 birth_age41 - scan_age41 birth_age35 | 0.0259392 | 0.0051878 | Inf | 5.00002 | 1.7e-06 |

#### Individual RSN H Comparison VPT vs MPT vs THC

| scan_age | birth_age | RSN | emmean | SE | df | asymp.LCL | asymp.UCL |
|---:|---:|---|---:|---:|---|---:|---:|
| 41 | 29 | H_MotorMedial | 0.6441664 | 0.0100222 | Inf | 0.6245232 | 0.6638096 |
| 41 | 35 | H_MotorMedial | 0.6583701 | 0.0048768 | Inf | 0.6488117 | 0.6679286 |
| 41 | 41 | H_MotorMedial | 0.6725739 | 0.0046920 | Inf | 0.6633777 | 0.6817701 |
| 41 | 29 | H_MotorLateral | 0.5703375 | 0.0100222 | Inf | 0.5506943 | 0.5899806 |
| 41 | 35 | H_MotorLateral | 0.6141829 | 0.0048768 | Inf | 0.6046245 | 0.6237414 |
| 41 | 41 | H_MotorLateral | 0.6580284 | 0.0046920 | Inf | 0.6488322 | 0.6672246 |
| 41 | 29 | H_Visual | 0.5450012 | 0.0100222 | Inf | 0.5253580 | 0.5646443 |
| 41 | 35 | H_Visual | 0.5705744 | 0.0048768 | Inf | 0.5610160 | 0.5801329 |
| 41 | 41 | H_Visual | 0.5961477 | 0.0046920 | Inf | 0.5869514 | 0.6053439 |
| 41 | 29 | H_Auditory | 0.5368239 | 0.0100222 | Inf | 0.5171808 | 0.5564671 |
| 41 | 35 | H_Auditory | 0.5751990 | 0.0048768 | Inf | 0.5656406 | 0.5847575 |
| 41 | 41 | H_Auditory | 0.6135741 | 0.0046920 | Inf | 0.6043779 | 0.6227704 |
| 41 | 29 | H_MotorAssociation | 0.5219864 | 0.0100222 | Inf | 0.5023433 | 0.5416296 |
| 41 | 35 | H_MotorAssociation | 0.5519272 | 0.0048768 | Inf | 0.5423688 | 0.5614857 |
| 41 | 41 | H_MotorAssociation | 0.5818680 | 0.0046920 | Inf | 0.5726718 | 0.5910643 |
| 41 | 29 | H_PosteriorParietal | 0.5815963 | 0.0100222 | Inf | 0.5619531 | 0.6012395 |
| 41 | 35 | H_PosteriorParietal | 0.5918621 | 0.0048768 | Inf | 0.5823037 | 0.6014205 |
| 41 | 41 | H_PosteriorParietal | 0.6021279 | 0.0046920 | Inf | 0.5929317 | 0.6113241 |
| 41 | 29 | H_Somatosensory | 0.5378278 | 0.0100222 | Inf | 0.5181846 | 0.5574710 |
| 41 | 35 | H_Somatosensory | 0.5640799 | 0.0048768 | Inf | 0.5545214 | 0.5736383 |
| 41 | 41 | H_Somatosensory | 0.5903319 | 0.0046920 | Inf | 0.5811357 | 0.5995281 |
| 41 | 29 | H_DorsalVisualStream | 0.4996798 | 0.0100222 | Inf | 0.4800366 | 0.5193229 |
| 41 | 35 | H_DorsalVisualStream | 0.5238136 | 0.0048768 | Inf | 0.5142552 | 0.5333721 |
| 41 | 41 | H_DorsalVisualStream | 0.5479475 | 0.0046920 | Inf | 0.5387512 | 0.5571437 |
| 41 | 29 | H_PosteriorCingulateCortex | 0.4957606 | 0.0100222 | Inf | 0.4761174 | 0.5154038 |
| 41 | 35 | H_PosteriorCingulateCortex | 0.5098505 | 0.0048768 | Inf | 0.5002920 | 0.5194089 |
| 41 | 41 | H_PosteriorCingulateCortex | 0.5239403 | 0.0046920 | Inf | 0.5147441 | 0.5331366 |
| 41 | 29 | H_FrontalPole | 0.4841255 | 0.0100222 | Inf | 0.4644823 | 0.5037687 |
| 41 | 35 | H_FrontalPole | 0.5132041 | 0.0048768 | Inf | 0.5036456 | 0.5227625 |
| 41 | 41 | H_FrontalPole | 0.5422827 | 0.0046920 | Inf | 0.5330865 | 0.5514789 |
| 41 | 29 | H_Midbrain | 0.4855891 | 0.0100222 | Inf | 0.4659459 | 0.5052323 |
| 41 | 35 | H_Midbrain | 0.4871265 | 0.0048768 | Inf | 0.4775681 | 0.4966849 |
| 41 | 41 | H_Midbrain | 0.4886639 | 0.0046920 | Inf | 0.4794677 | 0.4978601 |
| 41 | 29 | H_DorsolateralPrefrontal | 0.4692114 | 0.0100222 | Inf | 0.4495682 | 0.4888546 |
| 41 | 35 | H_DorsolateralPrefrontal | 0.4985351 | 0.0048768 | Inf | 0.4889767 | 0.5080936 |
| 41 | 41 | H_DorsolateralPrefrontal | 0.5278588 | 0.0046920 | Inf | 0.5186626 | 0.5370551 |
| 41 | 29 | H_Cerebellum | 0.3558582 | 0.0100222 | Inf | 0.3362151 | 0.3755014 |
| 41 | 35 | H_Cerebellum | 0.4172000 | 0.0048768 | Inf | 0.4076416 | 0.4267585 |
| 41 | 41 | H_Cerebellum | 0.4785419 | 0.0046920 | Inf | 0.4693456 | 0.4877381 |

Contrast

| contrast | RSN | estimate | p.value |
|---|---|---|---|
| scan_age41 birth_age35 - scan_age41 birth_age29 | H_MotorMedial | 0.0142037 | 0.2433627 |
| scan_age41 birth_age41 - scan_age41 birth_age29 | H_MotorMedial | 0.0284075 | 0.2433627 |
| scan_age41 birth_age41 - scan_age41 birth_age35 | H_MotorMedial | 0.0142037 | 0.2433627 |
| scan_age41 birth_age35 - scan_age41 birth_age29 | H_MotorLateral | 0.0438455 | **0.0000000** |
| scan_age41 birth_age41 - scan_age41 birth_age29 | H_MotorLateral | 0.0876909 | **0.0000000** |
| scan_age41 birth_age41 - scan_age41 birth_age35 | H_MotorLateral | 0.0438455 | **0.0000000** |
| scan_age41 birth_age35 - scan_age41 birth_age29 | H_Visual | 0.0255732 | **0.0005268** |
| scan_age41 birth_age41 - scan_age41 birth_age29 | H_Visual | 0.0511465 | **0.0005268** |
| scan_age41 birth_age41 - scan_age41 birth_age35 | H_Visual | 0.0255732 | **0.0005268** |
| scan_age41 birth_age35 - scan_age41 birth_age29 | H_Auditory | 0.0383751 | **0.0000000** |
| scan_age41 birth_age41 - scan_age41 birth_age29 | H_Auditory | 0.0767502 | **0.0000000** |
| scan_age41 birth_age41 - scan_age41 birth_age35 | H_Auditory | 0.0383751 | **0.0000000** |
| scan_age41 birth_age35 - scan_age41 birth_age29 | H_MotorAssociation | 0.0299408 | **0.0000298** |
| scan_age41 birth_age41 - scan_age41 birth_age29 | H_MotorAssociation | 0.0598816 | **0.0000298** |
| scan_age41 birth_age41 - scan_age41 birth_age35 | H_MotorAssociation | 0.0299408 | **0.0000298** |
| scan_age41 birth_age35 - scan_age41 birth_age29 | H_PosteriorParietal | 0.0102658 | 0.5605569 |
| scan_age41 birth_age41 - scan_age41 birth_age29 | H_PosteriorParietal | 0.0205316 | 0.5605569 |
| scan_age41 birth_age41 - scan_age41 birth_age35 | H_PosteriorParietal | 0.0102658 | 0.5605569 |
| scan_age41 birth_age35 - scan_age41 birth_age29 | H_Somatosensory | 0.0262521 | **0.0003752** |
| scan_age41 birth_age41 - scan_age41 birth_age29 | H_Somatosensory | 0.0525041 | **0.0003752** |
| scan_age41 birth_age41 - scan_age41 birth_age35 | H_Somatosensory | 0.0262521 | **0.0003752** |
| scan_age41 birth_age35 - scan_age41 birth_age29 | H_DorsalVisualStream | 0.0241338 | **0.0012027** |
| scan_age41 birth_age41 - scan_age41 birth_age29 | H_DorsalVisualStream | 0.0482677 | **0.0012027** |
| scan_age41 birth_age41 - scan_age41 birth_age35 | H_DorsalVisualStream | 0.0241338 | **0.0012027** |
| scan_age41 birth_age35 - scan_age41 birth_age29 | H_PosteriorCingulateCortex | 0.0140899 | 0.2433627 |
| scan_age41 birth_age41 - scan_age41 birth_age29 | H_PosteriorCingulateCortex | 0.0281797 | 0.2433627 |
| scan_age41 birth_age41 - scan_age41 birth_age35 | H_PosteriorCingulateCortex | 0.0140899 | 0.2433627 |
| scan_age41 birth_age35 - scan_age41 birth_age29 | H_FrontalPole | 0.0290786 | **0.0000484** |
| scan_age41 birth_age41 - scan_age41 birth_age29 | H_FrontalPole | 0.0581572 | **0.0000484** |
| scan_age41 birth_age41 - scan_age41 birth_age35 | H_FrontalPole | 0.0290786 | **0.0000484** |
| scan_age41 birth_age35 - scan_age41 birth_age29 | H_Midbrain | 0.0015374 | 1.0000000 |
| scan_age41 birth_age41 - scan_age41 birth_age29 | H_Midbrain | 0.0030748 | 1.0000000 |
| scan_age41 birth_age41 - scan_age41 birth_age35 | H_Midbrain | 0.0015374 | 1.0000000 |
| scan_age41 birth_age35 - scan_age41 birth_age29 | H_DorsolateralPrefrontal | 0.0293237 | **0.0000446** |
| scan_age41 birth_age41 - scan_age41 birth_age29 | H_DorsolateralPrefrontal | 0.0586474 | **0.0000446** |
| scan_age41 birth_age41 - scan_age41 birth_age35 | H_DorsolateralPrefrontal | 0.0293237 | **0.0000446** |
| scan_age41 birth_age35 - scan_age41 birth_age29 | H_Cerebellum | 0.0613418 | **0.0000000** |
| scan_age41 birth_age41 - scan_age41 birth_age29 | H_Cerebellum | 0.1226836 | **0.0000000** |
| scan_age41 birth_age41 - scan_age41 birth_age35 | H_Cerebellum | 0.0613418 | **0.0000000** |

Slopes:

| birth_age | RSN | scan_age.trend | SE | df | asymp.LCL | asymp.UCL |
|---:|---|---:|---:|---:|---:|---:|
| 29 | H_MotorMedial | 0.0342424 | 0.0054266 | Inf | 0.0236064 | 0.0448783 |
| 35 | H_MotorMedial | 0.0222780 | 0.0024957 | Inf | 0.0173866 | 0.0271694 |
| 41 | H_MotorMedial | 0.0103136 | 0.0026996 | Inf | 0.0050225 | 0.0156047 |
| 29 | H_MotorLateral | 0.0278141 | 0.0054266 | Inf | 0.0171781 | 0.0384500 |
| 35 | H_MotorLateral | 0.0174287 | 0.0024957 | Inf | 0.0125373 | 0.0223201 |
| 41 | H_MotorLateral | 0.0070433 | 0.0026996 | Inf | 0.0017521 | 0.0123344 |
| 29 | H_Visual | 0.0332561 | 0.0054266 | Inf | 0.0226201 | 0.0438920 |
| 35 | H_Visual | 0.0226673 | 0.0024957 | Inf | 0.0177759 | 0.0275587 |
| 41 | H_Visual | 0.0120786 | 0.0026996 | Inf | 0.0067874 | 0.0173697 |
| 29 | H_Auditory | 0.0232076 | 0.0054266 | Inf | 0.0125717 | 0.0338436 |
| 35 | H_Auditory | 0.0158705 | 0.0024957 | Inf | 0.0109791 | 0.0207618 |
| 41 | H_Auditory | 0.0085333 | 0.0026996 | Inf | 0.0032421 | 0.0138244 |
| 29 | H_MotorAssociation | 0.0193348 | 0.0054266 | Inf | 0.0086988 | 0.0299707 |
| 35 | H_MotorAssociation | 0.0102851 | 0.0024957 | Inf | 0.0053937 | 0.0151764 |
| 41 | H_MotorAssociation | 0.0012353 | 0.0026996 | Inf | -0.0040558 | 0.0065265 |
| 29 | H_PosteriorParietal | 0.0365959 | 0.0054266 | Inf | 0.0259600 | 0.0472318 |
| 35 | H_PosteriorParietal | 0.0262342 | 0.0024957 | Inf | 0.0213428 | 0.0311256 |
| 41 | H_PosteriorParietal | 0.0158725 | 0.0026996 | Inf | 0.0105813 | 0.0211636 |
| 29 | H_Somatosensory | 0.0216634 | 0.0054266 | Inf | 0.0110275 | 0.0322993 |
| 35 | H_Somatosensory | 0.0156283 | 0.0024957 | Inf | 0.0107369 | 0.0205197 |
| 41 | H_Somatosensory | 0.0095933 | 0.0026996 | Inf | 0.0043021 | 0.0148844 |
| 29 | H_DorsalVisualStream | 0.0200209 | 0.0054266 | Inf | 0.0093850 | 0.0306569 |
| 35 | H_DorsalVisualStream | 0.0142772 | 0.0024957 | Inf | 0.0093858 | 0.0191686 |
| 41 | H_DorsalVisualStream | 0.0085334 | 0.0026996 | Inf | 0.0032423 | 0.0138245 |
| 29 | H_PosteriorCingulateCortex | 0.0148589 | 0.0054266 | Inf | 0.0042230 | 0.0254948 |
| 35 | H_PosteriorCingulateCortex | 0.0101500 | 0.0024957 | Inf | 0.0052586 | 0.0150414 |
| 41 | H_PosteriorCingulateCortex | 0.0054411 | 0.0026996 | Inf | 0.0001500 | 0.0107323 |
| 29 | H_FrontalPole | 0.0141592 | 0.0054266 | Inf | 0.0035233 | 0.0247951 |
| 35 | H_FrontalPole | 0.0067738 | 0.0024957 | Inf | 0.0018824 | 0.0116652 |
| 41 | H_FrontalPole | -0.0006117 | 0.0026996 | Inf | -0.0059028 | 0.0046795 |
| 29 | H_Midbrain | -0.0017213 | 0.0054266 | Inf | -0.0123572 | 0.0089146 |
| 35 | H_Midbrain | -0.0004856 | 0.0024957 | Inf | -0.0053770 | 0.0044058 |
| 41 | H_Midbrain | 0.0007500 | 0.0026996 | Inf | -0.0045411 | 0.0060412 |
| 29 | H_DorsolateralPrefrontal | 0.0111846 | 0.0054266 | Inf | 0.0005487 | 0.0218206 |
| 35 | H_DorsolateralPrefrontal | 0.0064800 | 0.0024957 | Inf | 0.0015886 | 0.0113714 |
| 41 | H_DorsolateralPrefrontal | 0.0017755 | 0.0026996 | Inf | -0.0035157 | 0.0070666 |
| 29 | H_Cerebellum | 0.0131259 | 0.0054266 | Inf | 0.0024899 | 0.0237618 |
| 35 | H_Cerebellum | 0.0015371 | 0.0024957 | Inf | -0.0033543 | 0.0064285 |
| 41 | H_Cerebellum | -0.0100517 | 0.0026996 | Inf | -0.0153428 | -0.0047605 |

#### Preterm Groups vs THC Plots

##### Group RSN Bar Graph

Visual assessment of H values in the RSN compared in the groups.

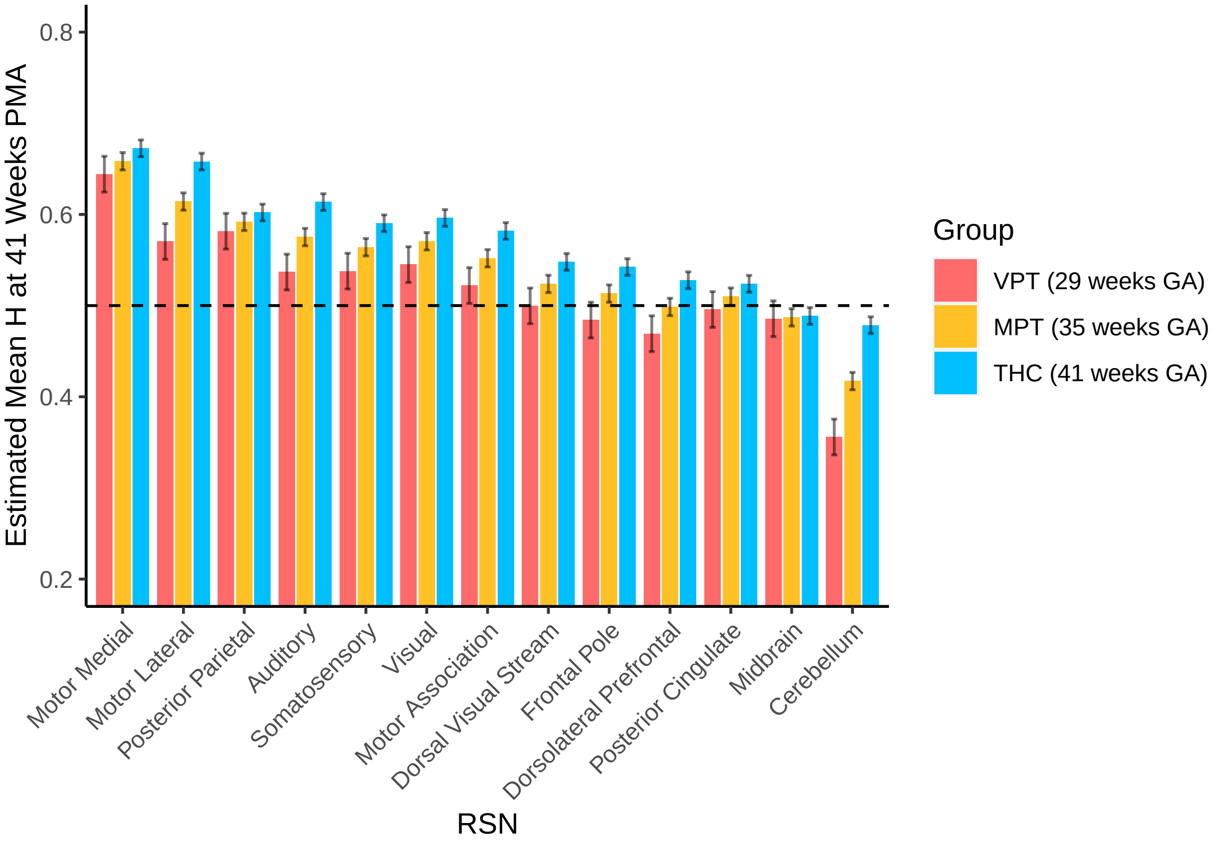


**Tissues Bar Graph**

Visual assessment of H values in the tissues compared in the groups.

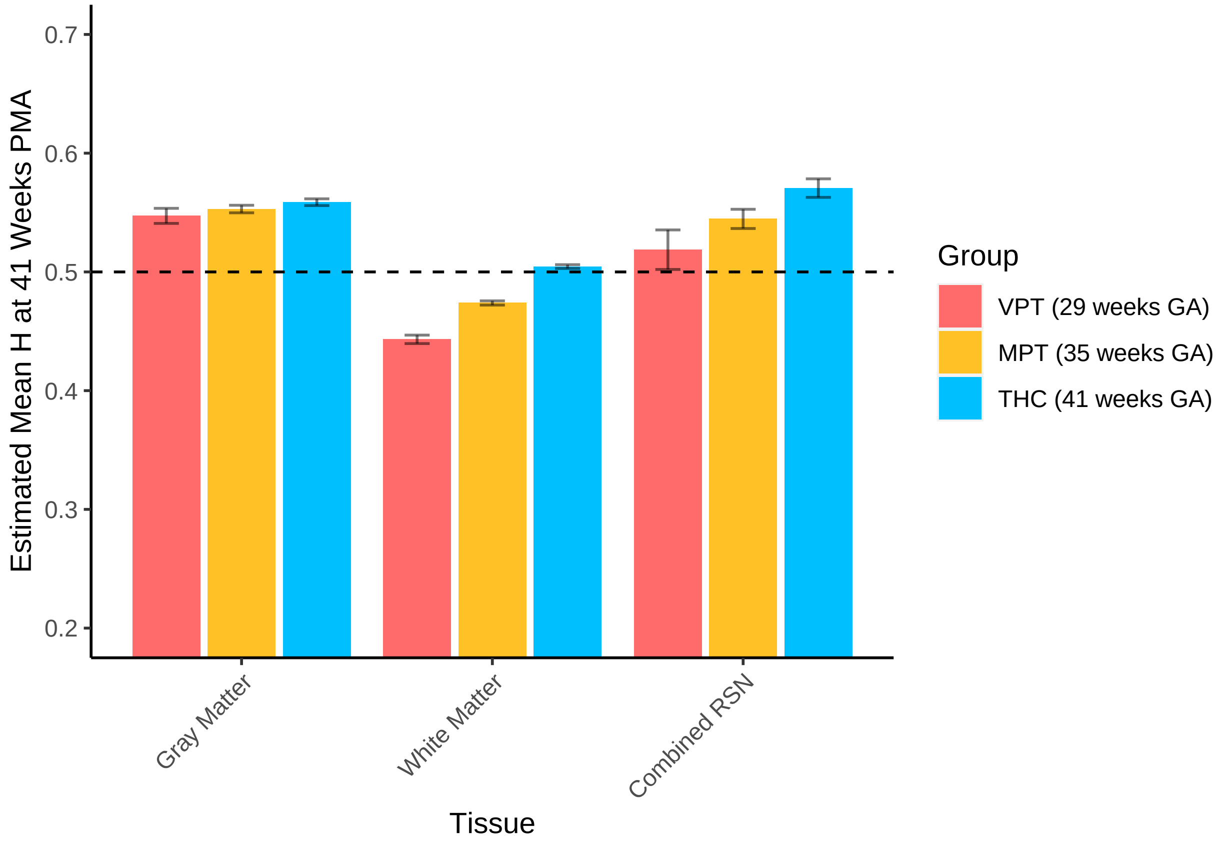


---

#### Is H associated with indirect measures of myelination (FA and RD)?

WM measures of FA and RD were calculated in the total white matter and in the RSNs.

Load data (hidden)

##### FA Analysis

###### FA comparison Total WM Preterm vs TEA

| scan_age | birth_age | emmean | SE | df | lower.CL | upper.CL |
|---:|---:|---:|---:|---:|---:|---:|
| 41 | 35 | 0.1866434 | 0.0016640 | 287.4906 | 0.1833683 | 0.1899186 |
| 41 | 41 | 0.1914910 | 0.0033521 | 266.5714 | 0.1848911 | 0.1980909 |

Contrast:

| contrast | estimate | p.value |
|---|---:|---:|
| scan_age41 birth_age41 - scan_age41 birth_age35 | 0.0048476 | **0.0184535** |

**FA vs scan age (PMA) plot**

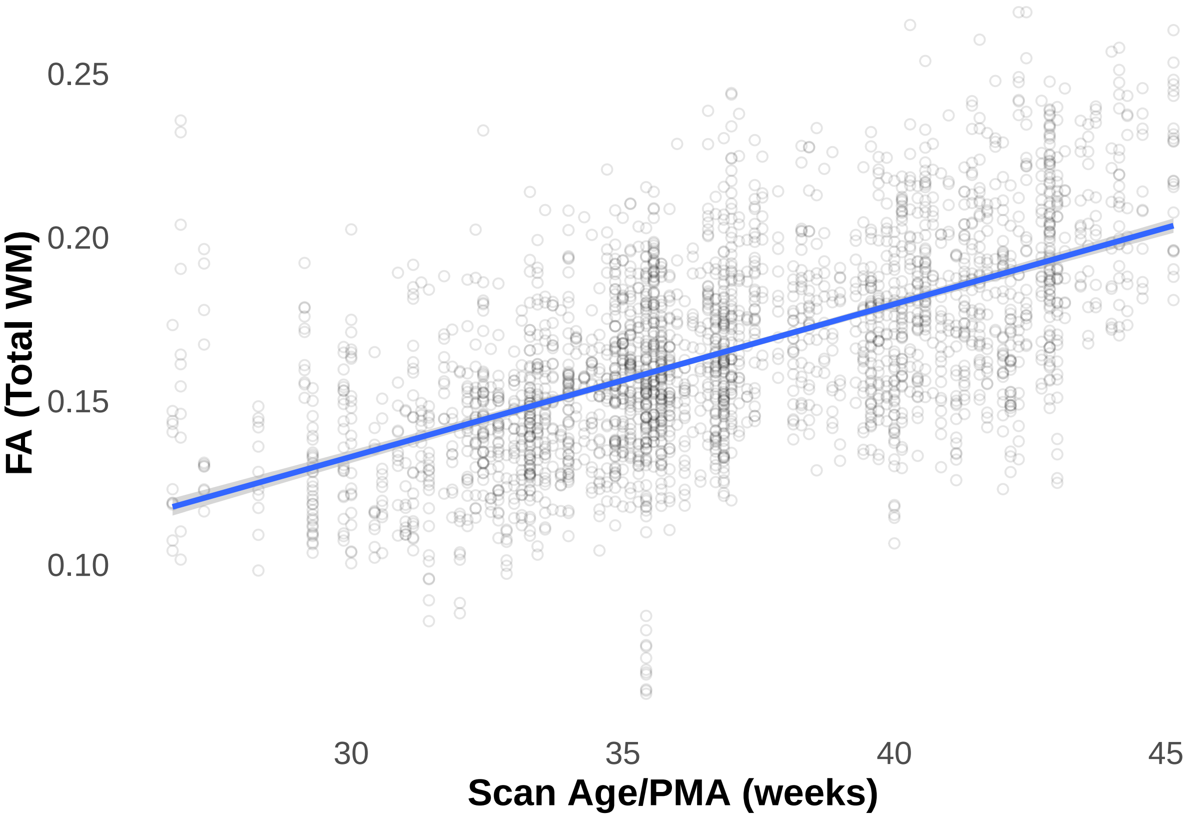


**FA comparison individual RSN Preterm vs TEA**

| scan_age | RSN | emmean | SE | df | lower.CL | upper.CL |
|---:|---|---:|---:|---:|---:|---:|
| 35 | Auditory | 0.1652154 | 0.0013978 | 391.2769 | 0.1624673 | 0.1679635 |
| 41 | Auditory | 0.1826610 | 0.0015781 | 605.6427 | 0.1795617 | 0.1857603 |
| 35 | Dorsal_Visual_Stream | 0.1603900 | 0.0013978 | 391.2769 | 0.1576419 | 0.1631381 |
| 41 | Dorsal_Visual_Stream | 0.1838964 | 0.0015781 | 605.6427 | 0.1807972 | 0.1869957 |
| 35 | Dorsolateral_Prefrontal | 0.1423323 | 0.0013978 | 391.2769 | 0.1395842 | 0.1450804 |
| 41 | Dorsolateral_Prefrontal | 0.1636586 | 0.0015781 | 605.6427 | 0.1605593 | 0.1667579 |
| 35 | Frontal_Pole | 0.1385895 | 0.0013978 | 391.2769 | 0.1358414 | 0.1413376 |
| 41 | Frontal_Pole | 0.1602939 | 0.0015781 | 605.6427 | 0.1571946 | 0.1633932 |
| 35 | LateralMotor | 0.1665454 | 0.0013978 | 391.2769 | 0.1637973 | 0.1692935 |
| 41 | LateralMotor | 0.2046735 | 0.0015781 | 605.6427 | 0.2015742 | 0.2077728 |
| 35 | MedialMotor | 0.1650782 | 0.0013957 | 389.1401 | 0.1623342 | 0.1678222 |
| 41 | MedialMotor | 0.2017077 | 0.0015781 | 605.6018 | 0.1986085 | 0.2048069 |
| 35 | MotorAssociation | 0.1536773 | 0.0013957 | 389.1401 | 0.1509333 | 0.1564213 |
| 41 | MotorAssociation | 0.1822827 | 0.0015781 | 605.6018 | 0.1791835 | 0.1853819 |
| 35 | Posterior_Cingular_Cortex | 0.1896896 | 0.0013978 | 391.2769 | 0.1869415 | 0.1924377 |
| 41 | Posterior_Cingular_Cortex | 0.2152956 | 0.0015781 | 605.6427 | 0.2121963 | 0.2183949 |
| 35 | Posterior_Parietal | 0.1337868 | 0.0013957 | 389.1401 | 0.1310428 | 0.1365308 |
| 41 | Posterior_Parietal | 0.1548599 | 0.0015781 | 605.6018 | 0.1517607 | 0.1579591 |
| 35 | Somatosensory | 0.1595581 | 0.0013978 | 391.2769 | 0.1568100 | 0.1623062 |
| 41 | Somatosensory | 0.2017429 | 0.0015781 | 605.6427 | 0.1986436 | 0.2048422 |
| 35 | Visual | 0.1484377 | 0.0013978 | 391.2769 | 0.1456896 | 0.1511858 |
| 41 | Visual | 0.1644413 | 0.0015781 | 605.6427 | 0.1613420 | 0.1675406 |

Contrast:

| contrast | estimate | p.value |
|---|---|---|
| scan_age41 Auditory - scan_age35 Auditory | 0.0174456 | **0.0000000** |
| scan_age35 Dorsal_Visual_Stream - scan_age35 Auditory | -0.0048254 | **0.0013125** |
| scan_age35 Dorsal_Visual_Stream - scan_age41 Auditory | -0.0222710 | **0.0000000** |
| scan_age41 Dorsal_Visual_Stream - scan_age35 Auditory | 0.0186810 | **0.0000000** |
| scan_age41 Dorsal_Visual_Stream - scan_age41 Auditory | 0.0012354 | 1.0000000 |
| scan_age41 Dorsal_Visual_Stream - scan_age35 Dorsal_Visual_Stream | 0.0235064 | **0.0000000** |
| scan_age35 Dorsolateral_Prefrontal - scan_age35 Auditory | -0.0228831 | **0.0000000** |
| scan_age35 Dorsolateral_Prefrontal - scan_age41 Auditory | -0.0403287 | **0.0000000** |
| scan_age35 Dorsolateral_Prefrontal - scan_age35 Dorsal_Visual_Stream | -0.0180577 | **0.0000000** |
| scan_age35 Dorsolateral_Prefrontal - scan_age41 Dorsal_Visual_Stream | -0.0415641 | **0.0000000** |
| scan_age41 Dorsolateral_Prefrontal - scan_age35 Auditory | -0.0015568 | 1.0000000 |
| scan_age41 Dorsolateral_Prefrontal - scan_age41 Auditory | -0.0190024 | **0.0000000** |
| scan_age41 Dorsolateral_Prefrontal - scan_age35 Dorsal_Visual_Stream | 0.0032686 | 0.4302443 |
| scan_age41 Dorsolateral_Prefrontal - scan_age41 Dorsal_Visual_Stream | -0.0202378 | **0.0000000** |
| scan_age41 Dorsolateral_Prefrontal - scan_age35 Dorsolateral_Prefrontal | 0.0213263 | **0.0000000** |
| scan_age35 Frontal_Pole - scan_age35 Auditory | -0.0266259 | **0.0000000** |
| scan_age35 Frontal_Pole - scan_age41 Auditory | -0.0440715 | **0.0000000** |
| scan_age35 Frontal_Pole - scan_age35 Dorsal_Visual_Stream | -0.0218005 | **0.0000000** |
| scan_age35 Frontal_Pole - scan_age41 Dorsal_Visual_Stream | -0.0453069 | **0.0000000** |
| scan_age35 Frontal_Pole - scan_age35 Dorsolateral_Prefrontal | -0.0037428 | **0.0357698** |
| scan_age35 Frontal_Pole - scan_age41 Dorsolateral_Prefrontal | -0.0250691 | **0.0000000** |
| scan_age41 Frontal_Pole - scan_age35 Auditory | -0.0049216 | **0.0132909** |
| scan_age41 Frontal_Pole - scan_age41 Auditory | -0.0223672 | **0.0000000** |
| scan_age41 Frontal_Pole - scan_age35 Dorsal_Visual_Stream | -0.0000962 | 1.0000000 |
| scan_age41 Frontal_Pole - scan_age41 Dorsal_Visual_Stream | -0.0236026 | **0.0000000** |
| scan_age41 Frontal_Pole - scan_age35 Dorsolateral_Prefrontal | 0.0179616 | **0.0000000** |
| scan_age41 Frontal_Pole - scan_age41 Dorsolateral_Prefrontal | -0.0033648 | 0.5690552 |
| scan_age41 Frontal_Pole - scan_age35 Frontal_Pole | 0.0217044 | **0.0000000** |
| scan_age35 LateralMotor - scan_age35 Auditory | 0.0013300 | 1.0000000 |
| scan_age35 LateralMotor - scan_age41 Auditory | -0.0161156 | **0.0000000** |
| scan_age35 LateralMotor - scan_age35 Dorsal_Visual_Stream | 0.0061554 | **0.0000069** |
| scan_age35 LateralMotor - scan_age41 Dorsal_Visual_Stream | -0.0173510 | **0.0000000** |
| scan_age35 LateralMotor - scan_age35 Dorsolateral_Prefrontal | 0.0242131 | **0.0000000** |
| scan_age35 LateralMotor - scan_age41 Dorsolateral_Prefrontal | 0.0028868 | 0.7835758 |
| scan_age35 LateralMotor - scan_age35 Frontal_Pole | 0.0279559 | **0.0000000** |
| scan_age35 LateralMotor - scan_age41 Frontal_Pole | 0.0062516 | **0.0003047** |
| scan_age41 LateralMotor - scan_age35 Auditory | 0.0394581 | **0.0000000** |
| scan_age41 LateralMotor - scan_age41 Auditory | 0.0220125 | **0.0000000** |
| scan_age41 LateralMotor - scan_age35 Dorsal_Visual_Stream | 0.0442835 | **0.0000000** |
| scan_age41 LateralMotor - scan_age41 Dorsal_Visual_Stream | 0.0207770 | **0.0000000** |
| scan_age41 LateralMotor - scan_age35 Dorsolateral_Prefrontal | 0.0623412 | **0.0000000** |
| scan_age41 LateralMotor - scan_age41 Dorsolateral_Prefrontal | 0.0410149 | **0.0000000** |
| scan_age41 LateralMotor - scan_age35 Frontal_Pole | 0.0660840 | **0.0000000** |
| scan_age41 LateralMotor - scan_age41 Frontal_Pole | 0.0443796 | **0.0000000** |
| scan_age41 LateralMotor - scan_age35 LateralMotor | 0.0381280 | **0.0000000** |
| scan_age35 MedialMotor - scan_age35 Auditory | -0.0001372 | 1.0000000 |
| scan_age35 MedialMotor - scan_age41 Auditory | -0.0175828 | **0.0000000** |
| scan_age35 MedialMotor - scan_age35 Dorsal_Visual_Stream | 0.0046882 | **0.0019996** |
| scan_age35 MedialMotor - scan_age41 Dorsal_Visual_Stream | -0.0188182 | **0.0000000** |
| scan_age35 MedialMotor - scan_age35 Dorsolateral_Prefrontal | 0.0227459 | **0.0000000** |
| scan_age35 MedialMotor - scan_age41 Dorsolateral_Prefrontal | 0.0014196 | 1.0000000 |
| scan_age35 MedialMotor - scan_age35 Frontal_Pole | 0.0264887 | **0.0000000** |
| scan_age35 MedialMotor - scan_age41 Frontal_Pole | 0.0047843 | **0.0176392** |
| scan_age35 MedialMotor - scan_age35 LateralMotor | -0.0014672 | 1.0000000 |
| scan_age35 MedialMotor - scan_age41 LateralMotor | -0.0395953 | **0.0000000** |
| scan_age41 MedialMotor - scan_age35 Auditory | 0.0364923 | **0.0000000** |
| scan_age41 MedialMotor - scan_age41 Auditory | 0.0190467 | **0.0000000** |

**FA Slopes in RSN Preterm vs TEA**

| RSN | scan_age.trend | std.error | df | statistic | p.value |
|---|---|---|---|---|---|
| Somatosensory | 0.0070308 | 0.0001954 | 2490.515 | 35.98728 | 0 |
| LateralMotor | 0.0063547 | 0.0001954 | 2490.515 | 32.52654 | 0 |
| MedialMotor | 0.0061049 | 0.0001951 | 2491.526 | 31.28379 | 0 |
| MotorAssociation | 0.0047676 | 0.0001951 | 2491.526 | 24.43065 | 0 |
| Posterior_Cingular_Cortex | 0.0042677 | 0.0001954 | 2490.515 | 21.84418 | 0 |
| Dorsal_Visual_Stream | 0.0039177 | 0.0001954 | 2490.515 | 20.05302 | 0 |
| Frontal_Pole | 0.0036174 | 0.0001954 | 2490.515 | 18.51570 | 0 |
| Dorsolateral_Prefrontal | 0.0035544 | 0.0001954 | 2490.515 | 18.19321 | 0 |
| Posterior_Parietal | 0.0035122 | 0.0001951 | 2491.526 | 17.99762 | 0 |
| Auditory | 0.0029076 | 0.0001954 | 2490.515 | 14.88260 | 0 |
| Visual | 0.0026673 | 0.0001954 | 2490.515 | 13.65249 | 0 |

**FA plot in the RSN**

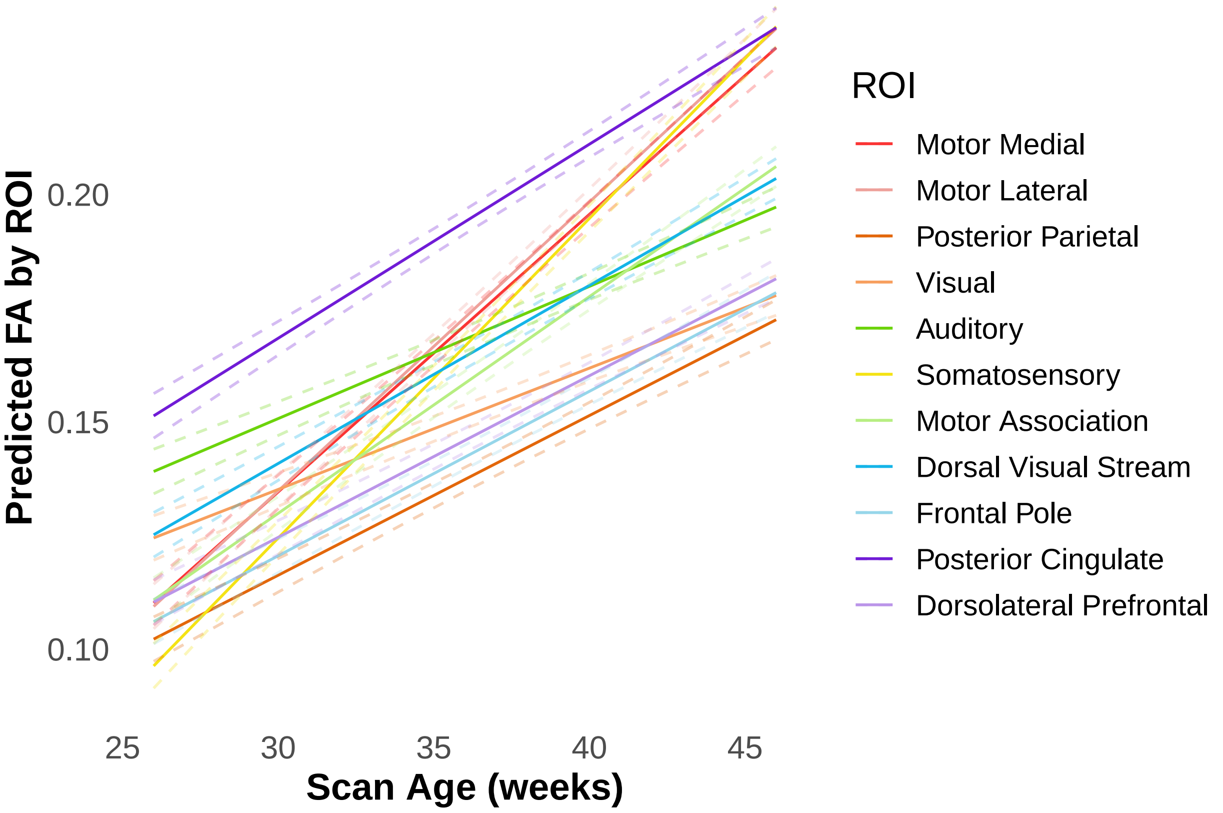


**FA and H Correlation test**

**Total WM**:

| estimate | statistic | p.value | parameter | conf.low | conf.high | method | alternativ |
|---|---|---|---|---|---|---|---|
| 0.4189898 | 23.56545 | 0 | 2608 | 0.3868409 | 0.4501214 | Pearson's product-moment correlation | two.sided |

**White Matter Regions**:

| RSN | estimate | statistic | p.value | parameter | conf.low | conf.high | method |
|---|---|---|---|---|---|---|---|
| LateralMotor | 0.6055746 | 11.665513 | 0 | 235 | 0.5181274 | 0.6804923 | Pearson's product- |
| Somatosensory | 0.5891108 | 11.176130 | 0 | 235 | 0.4991521 | 0.6665061 | Pearson's product- |
| MedialMotor | 0.5515703 | 10.158368 | 0 | 236 | 0.4564212 | 0.6342469 | Pearson's product- |
| Posterior_Parietal | 0.5234531 | 9.437706 | 0 | 236 | 0.4245508 | 0.6100106 | Pearson's product- |
| Auditory | 0.5065448 | 9.006103 | 0 | 235 | 0.4052746 | 0.5955339 | Pearson's product- |
| MotorAssociation | 0.4699760 | 8.179539 | 0 | 236 | 0.3646040 | 0.5634636 | Pearson's product- |
| Dorsal_Visual_Stream | 0.4608583 | 7.960601 | 0 | 235 | 0.3542311 | 0.5556564 | Pearson's product- |
| Dorsolateral_Prefrontal | 0.4249342 | 7.196138 | 0 | 235 | 0.3145359 | 0.5239907 | Pearson's product- |
| Visual | 0.4114515 | 6.920356 | 0 | 235 | 0.2997368 | 0.5120351 | Pearson's product- |
| Frontal_Pole | 0.3699419 | 6.104163 | 0 | 235 | 0.2545097 | 0.4749808 | Pearson's product- |
| Posterior_Cingular_Cortex | 0.3524966 | 5.774306 | 0 | 235 | 0.2356515 | 0.4592959 | Pearson's product- |

**FA H correlation preterm vs TEA**:

Preterm:

| estimate | statistic | p.value | parameter | conf.low | conf.high | method | alternative |
|---|---|---|---|---|---|---|---|
| 0.1594192 | 6.225008 | 0 | 1486 | 0.109489 | 0.2085469 | Pearson's product-moment correlation | two.sided |

Term:

| estimate | statistic | p.value | parameter | conf.low | conf.high | method | alternativ |
|---|---|---|---|---|---|---|---|
| 0.2031904 | 6.77226 | 0 | 1065 | 0.1449436 | 0.2600338 | Pearson's product-moment correlation | two.sided |

**FA and H Correlation Plot**

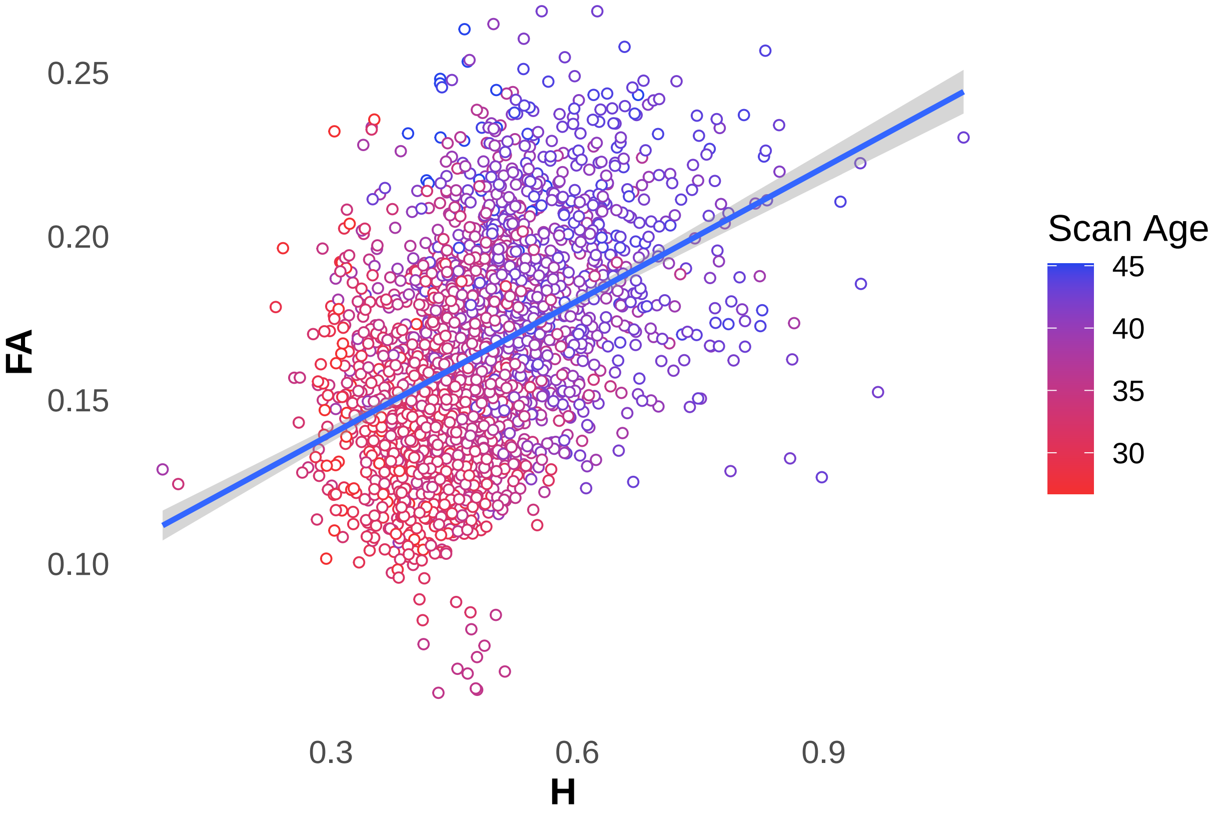

**FA and H Correlation RSN Plot**

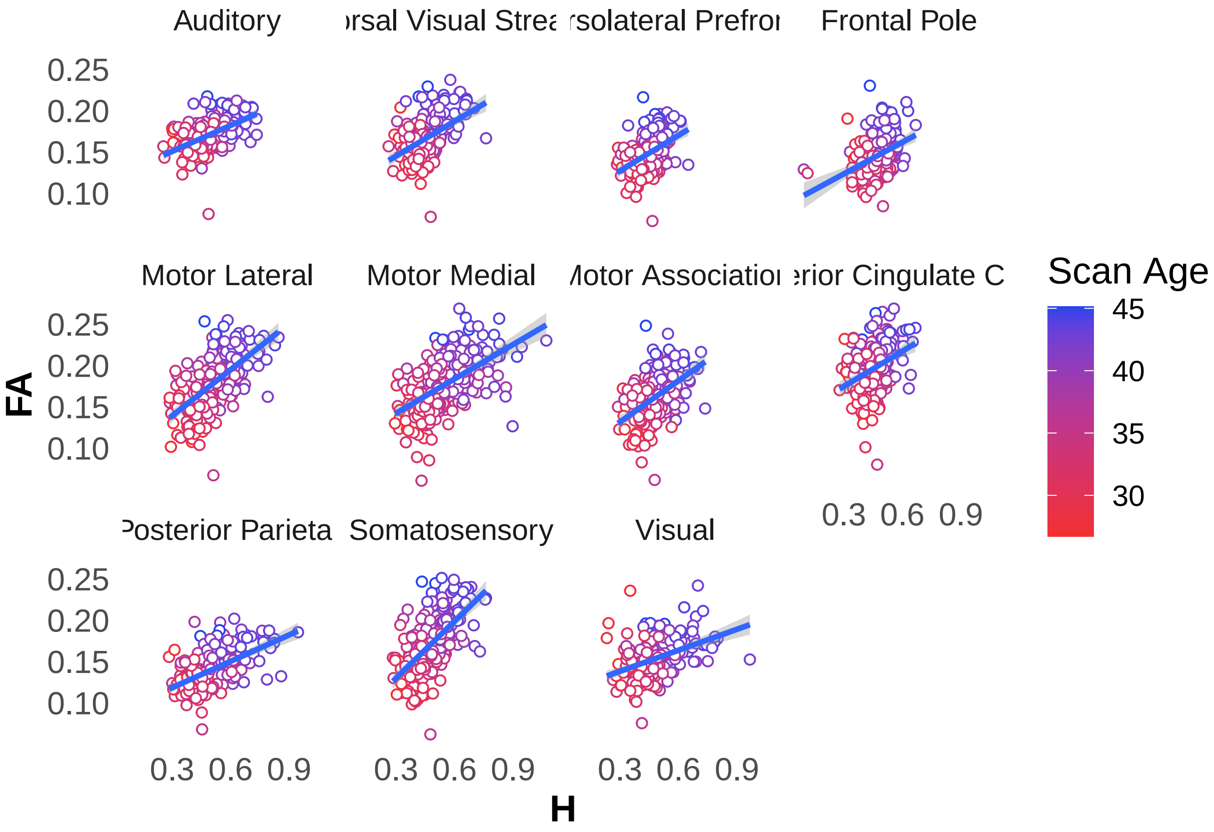


#### RD Analysis

**RD comparison Total WM Preterm vs TEA:**

| scan_age | birth_age | emmean | SE | df | lower.CL | upper.CL |
|---|---|---|---|---|---|---|
| 41 | 35 | 0.0010866 | 5.20e-06 | 252.721 | 0.0010763 | 0.0010970 |
| 41 | 41 | 0.0010735 | 1.07e-05 | 237.834 | 0.0010525 | 0.0010945 |

Contrast:

| contrast | estimate | p.value |
|---|---|---|
| scan_age41 birth_age41 - scan_age41 birth_age35 | -1.31e-05 | **0.0458228** |

**RD vs scan age (PMA) plot**

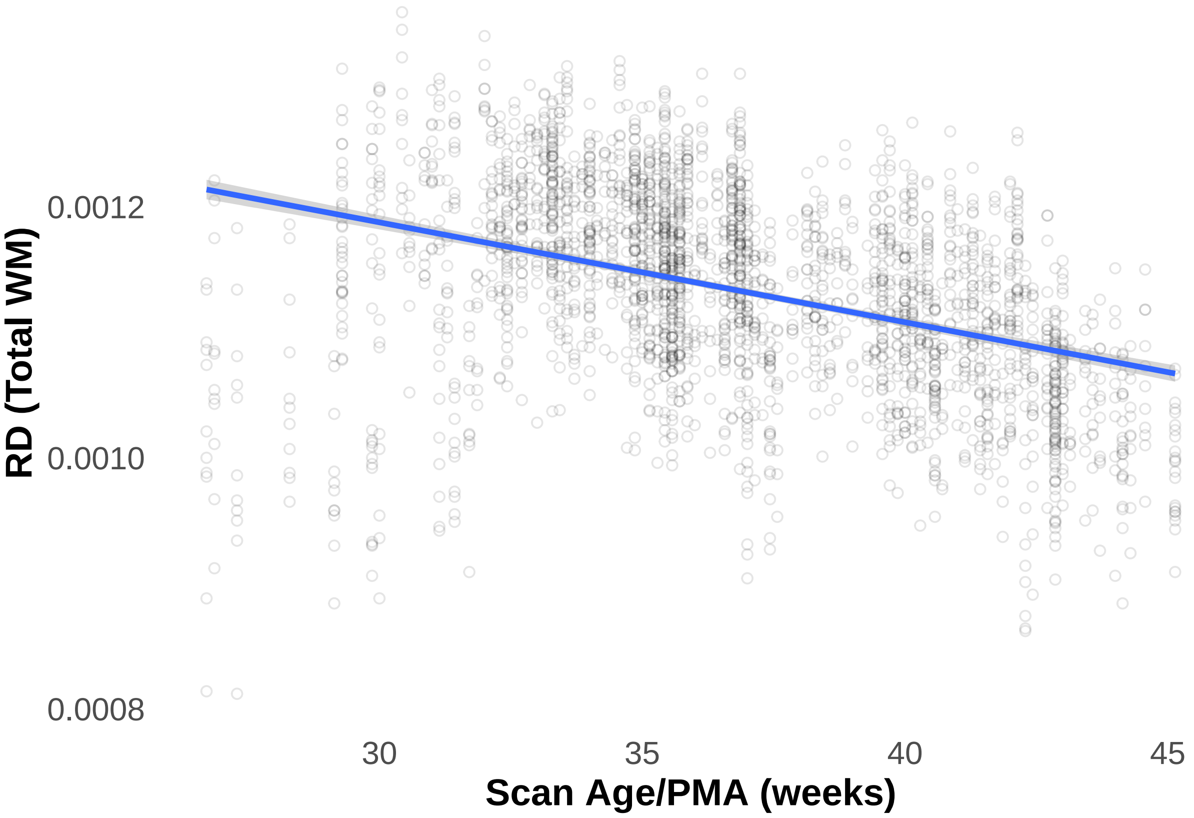


**RD comparison individual RSN Preterm vs TEA**

| scan_age | RSN | emmean | SE | df | lower.CL | upper.CL |
|---|---|---|---|---|---|---|
| 35 | Auditory | 0.0011218 | 5.0e-06 | 398.9043 | 0.0011120 | 0.0011316 |
| 41 | Auditory | 0.0011006 | 5.6e-06 | 621.1871 | 0.0010895 | 0.0011116 |
| 35 | Dorsal__Visual__Stream | 0.0011697 | 5.0e-06 | 398.9043 | 0.0011599 | 0.0011795 |
| 41 | Dorsal__Visual__Stream | 0.0011462 | 5.6e-06 | 621.1871 | 0.0011352 | 0.0011573 |
| 35 | Dorsolateral__Prefrontal | 0.0011778 | 5.0e-06 | 398.9043 | 0.0011680 | 0.0011875 |
| 41 | Dorsolateral__Prefrontal | 0.0011590 | 5.6e-06 | 621.1871 | 0.0011479 | 0.0011700 |
| 35 | Frontal__Pole | 0.0011286 | 5.0e-06 | 398.9043 | 0.0011189 | 0.0011384 |
| 41 | Frontal__Pole | 0.0011179 | 5.6e-06 | 621.1871 | 0.0011069 | 0.0011290 |
| 35 | LateralMotor | 0.0011341 | 5.0e-06 | 398.9043 | 0.0011243 | 0.0011438 |
| 41 | LateralMotor | 0.0010611 | 5.6e-06 | 621.1871 | 0.0010501 | 0.0010722 |
| 35 | MedialMotor | 0.0011099 | 5.0e-06 | 396.6905 | 0.0011001 | 0.0011196 |
| 41 | MedialMotor | 0.0010167 | 5.6e-06 | 621.1459 | 0.0010057 | 0.0010277 |
| 35 | MotorAssociation | 0.0011599 | 5.0e-06 | 396.6905 | 0.0011501 | 0.0011696 |
| 41 | MotorAssociation | 0.0011097 | 5.6e-06 | 621.1459 | 0.0010987 | 0.0011207 |
| 35 | Posterior__Cingular__Cortex | 0.0011154 | 5.0e-06 | 398.9043 | 0.0011057 | 0.0011252 |
| 41 | Posterior__Cingular__Cortex | 0.0010708 | 5.6e-06 | 621.1871 | 0.0010597 | 0.0010818 |
| 35 | Posterior__Parietal | 0.0011856 | 5.0e-06 | 396.6905 | 0.0011758 | 0.0011953 |
| 41 | Posterior__Parietal | 0.0011529 | 5.6e-06 | 621.1459 | 0.0011419 | 0.0011640 |
| 35 | Somatosensory | 0.0011773 | 5.0e-06 | 398.9043 | 0.0011675 | 0.0011870 |
| 41 | Somatosensory | 0.0010953 | 5.6e-06 | 621.1871 | 0.0010842 | 0.0011063 |
| 35 | Visual | 0.0011407 | 5.0e-06 | 398.9043 | 0.0011309 | 0.0011504 |
| 41 | Visual | 0.0011009 | 5.6e-06 | 621.1871 | 0.0010899 | 0.0011120 |

| contrast | RSN | estimate | p.value |
|---|---|---|---|
| scan_age41 - scan_age35 | Auditory | -2.12e-05 | **0.0000015** |
| scan_age41 - scan_age35 | Dorsal_Visual_Stream | -2.35e-05 | **0.0000001** |
| scan_age41 - scan_age35 | Dorsolateral_Prefrontal | -1.88e-05 | **0.0000165** |
| scan_age41 - scan_age35 | Frontal_Pole | -1.07e-05 | **0.0111350** |
| scan_age41 - scan_age35 | LateralMotor | -7.29e-05 | **0.0000000** |
| scan_age41 - scan_age35 | MedialMotor | -9.32e-05 | **0.0000000** |
| scan_age41 - scan_age35 | MotorAssociation | -5.02e-05 | **0.0000000** |
| scan_age41 - scan_age35 | Posterior_Cingular_Cortex | -4.46e-05 | **0.0000000** |
| scan_age41 - scan_age35 | Posterior_Parietal | -3.27e-05 | **0.0000000** |
| scan_age41 - scan_age35 | Somatosensory | -8.20e-05 | **0.0000000** |
| scan_age41 - scan_age35 | Visual | -3.97e-05 | **0.0000000** |

##### RD Slopes in RSN Preterm vs TEA

| RSN | scan_age.trend | std.error | df | statistic | p.value |
|---|---|---|---|---|---|
| Frontal_Pole | -1.80e-06 | 7e-07 | 2491.566 | -2.540332 | 0.0111350 |
| Dorsolateral_Prefrontal | -3.10e-06 | 7e-07 | 2491.566 | -4.468371 | 0.0000082 |
| Auditory | -3.50e-06 | 7e-07 | 2491.566 | -5.043531 | 0.0000005 |
| Dorsal_Visual_Stream | -3.90e-06 | 7e-07 | 2491.566 | -5.583138 | 0.0000000 |
| Posterior_Parietal | -5.40e-06 | 7e-07 | 2492.596 | -7.769251 | 0.0000000 |
| Visual | -6.60e-06 | 7e-07 | 2491.566 | -9.438729 | 0.0000000 |
| Posterior_Cingular_Cortex | -7.40e-06 | 7e-07 | 2491.566 | -10.609672 | 0.0000000 |
| MotorAssociation | -8.40e-06 | 7e-07 | 2492.596 | -11.935292 | 0.0000000 |
| LateralMotor | -1.22e-05 | 7e-07 | 2491.566 | -17.331371 | 0.0000000 |
| Somatosensory | -1.37e-05 | 7e-07 | 2491.566 | -19.491854 | 0.0000000 |
| MedialMotor | -1.55e-05 | 7e-07 | 2492.596 | -22.164433 | 0.0000000 |

**RD plot in the RSN**

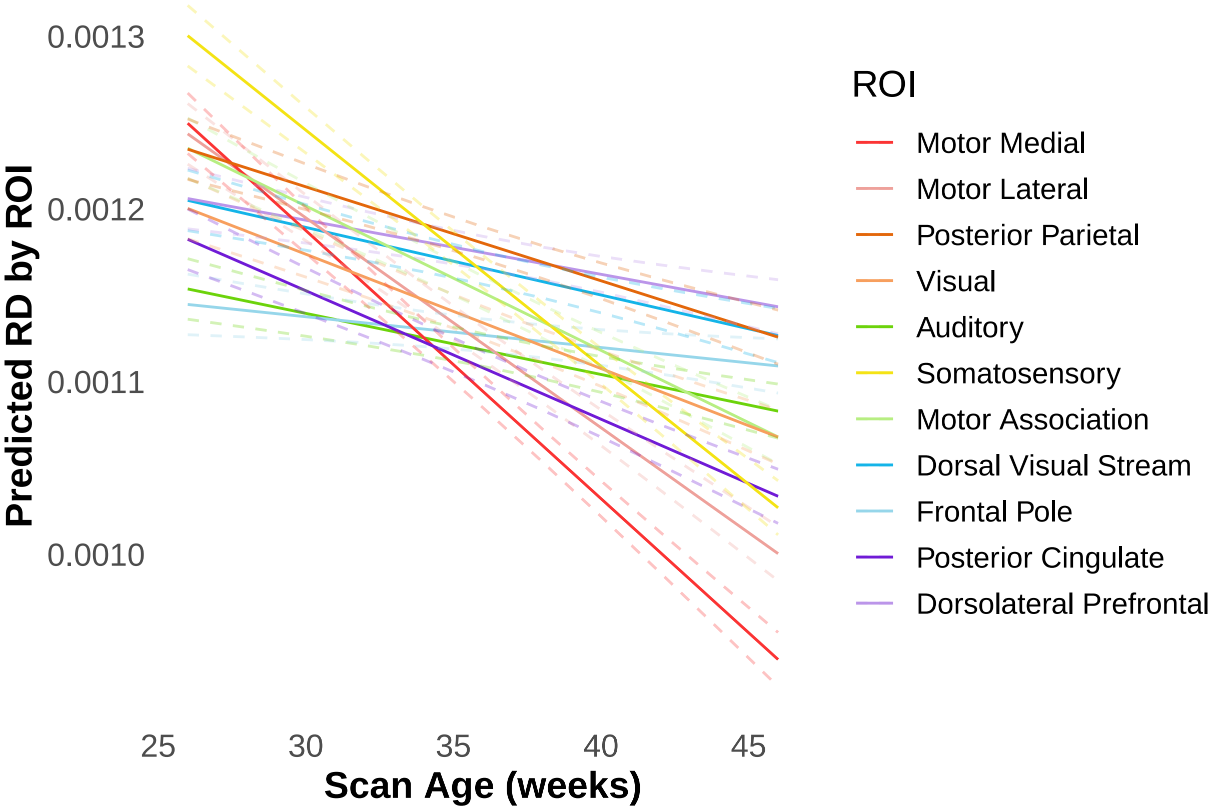


#### RD and H Correlation tests

**Total WM**

| estimate | statistic | p.value | parameter | conf.low | conf.high | method |
|---|---|---|---|---|---|---|
| -0.348385 | -18.98063 | 0 | 2608 | -0.3816511 | -0.3142174 | Pearson's product-moment correlat |

**White Matter Regions**

| RSN | estimate | statistic | p.value | parameter | conf.low |
|---|---|---|---|---|---|
| Auditory | -0.2107038 | -3.304208 | **0.0011012** | 235 | -0.3292926 |
| Dorsal_Visual_Stream | -0.1512207 | -2.345138 | **0.0198530** | 235 | -0.2733828 |
| Dorsolateral_Prefrontal | -0.1311752 | -2.028405 | **0.0436470** | 235 | -0.2543538 |
| Frontal_Pole | -0.0084357 | -0.129322 | 0.8972135 | 235 | -0.1357201 |
| LateralMotor | -0.4745088 | -8.263648 | **0.0000000** | 235 | -0.5676171 |
| MedialMotor | -0.5654581 | -10.532220 | **0.0000000** | 236 | -0.6461582 |
| MotorAssociation | -0.3377944 | -5.513373 | **0.0000001** | 236 | -0.4458068 |
| Posterior_Cingular_Cortex | -0.2999297 | -4.819729 | **0.0000026** | 235 | -0.4116275 |
| Posterior_Parietal | -0.3219963 | -5.224873 | **0.0000004** | 236 | -0.4314905 |
| Somatosensory | -0.4990559 | -8.828350 | **0.0000000** | 235 | -0.5890271 |
| Visual | -0.2593886 | -4.117274 | **0.0000531** | 235 | -0.3744421 |

| RSN | estimate | statistic | p.value | parameter | conf.low | conf.high | method |
|---|---|---|---|---|---|---|---|
| Frontal_Pole | -0.0084357 | -0.129322 | 0.8972135 | 235 | -0.1357201 | 0.1191226 | Pearson's p |
| Dorsolateral_Prefrontal | -0.1311752 | -2.028405 | **0.0436470** | 235 | -0.2543538 | -0.0038086 | Pearson's p |
| Dorsal_Visual_Stream | -0.1512207 | -2.345138 | **0.0198530** | 235 | -0.2733828 | -0.0242578 | Pearson's p |
| Auditory | -0.2107038 | -3.304208 | **0.0011012** | 235 | -0.3292926 | -0.0855711 | Pearson's p |
| Visual | -0.2593886 | -4.117274 | **0.0000531** | 235 | -0.3744421 | -0.1364691 | Pearson's p |
| Posterior_Cingular_Cortex | -0.2999297 | -4.819729 | **0.0000026** | 235 | -0.4116275 | -0.1793543 | Pearson's p |
| Posterior_Parietal | -0.3219963 | -5.224873 | **0.0000004** | 236 | -0.4314905 | -0.2031527 | Pearson's p |
| MotorAssociation | -0.3377944 | -5.513373 | **0.0000001** | 236 | -0.4458068 | -0.2200863 | Pearson's p |
| LateralMotor | -0.4745088 | -8.263648 | **0.0000000** | 235 | -0.5676171 | -0.3694159 | Pearson's p |
| Somatosensory | -0.4990559 | -8.828350 | **0.0000000** | 235 | -0.5890271 | -0.3968641 | Pearson's p |
| MedialMotor | -0.5654581 | -10.532220 | **0.0000000** | 236 | -0.6461582 | -0.4722536 | Pearson's p |

##### RD and H Correlation Plot

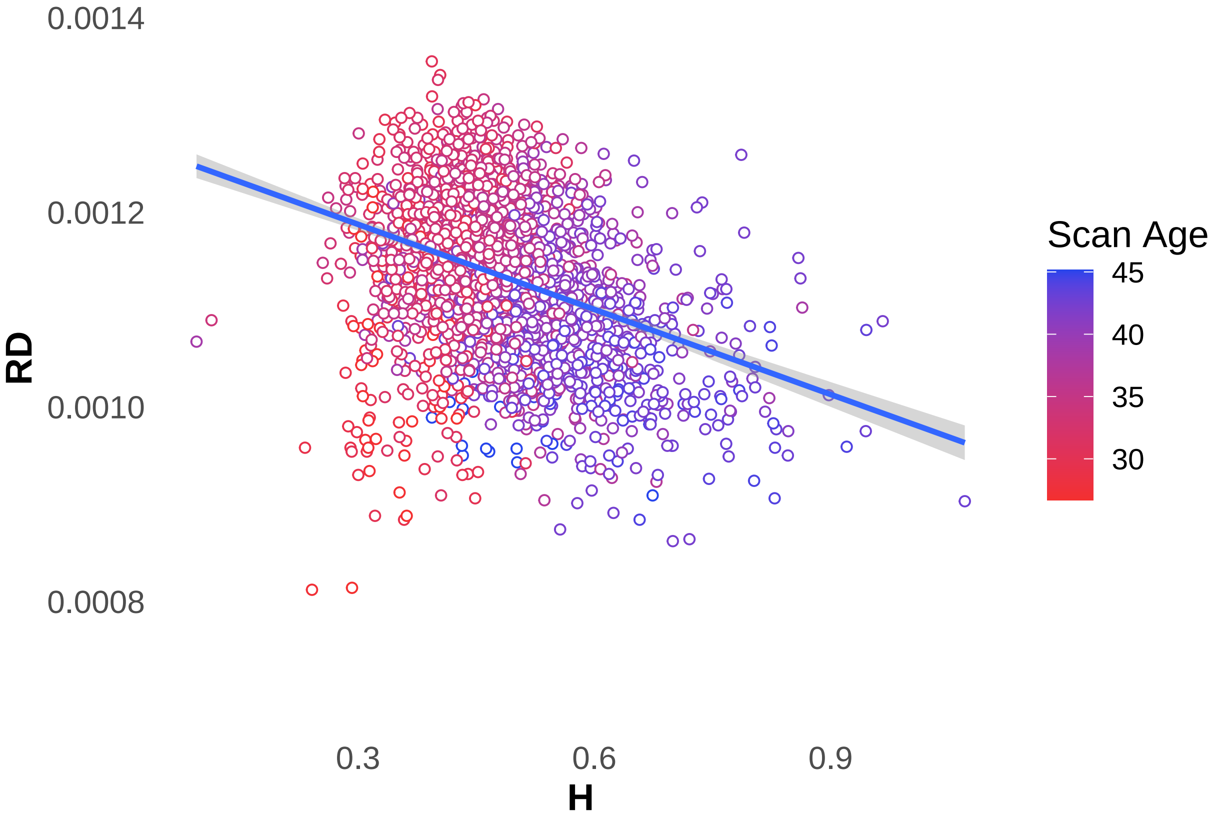

**RD and H Correlation RSN Plot**

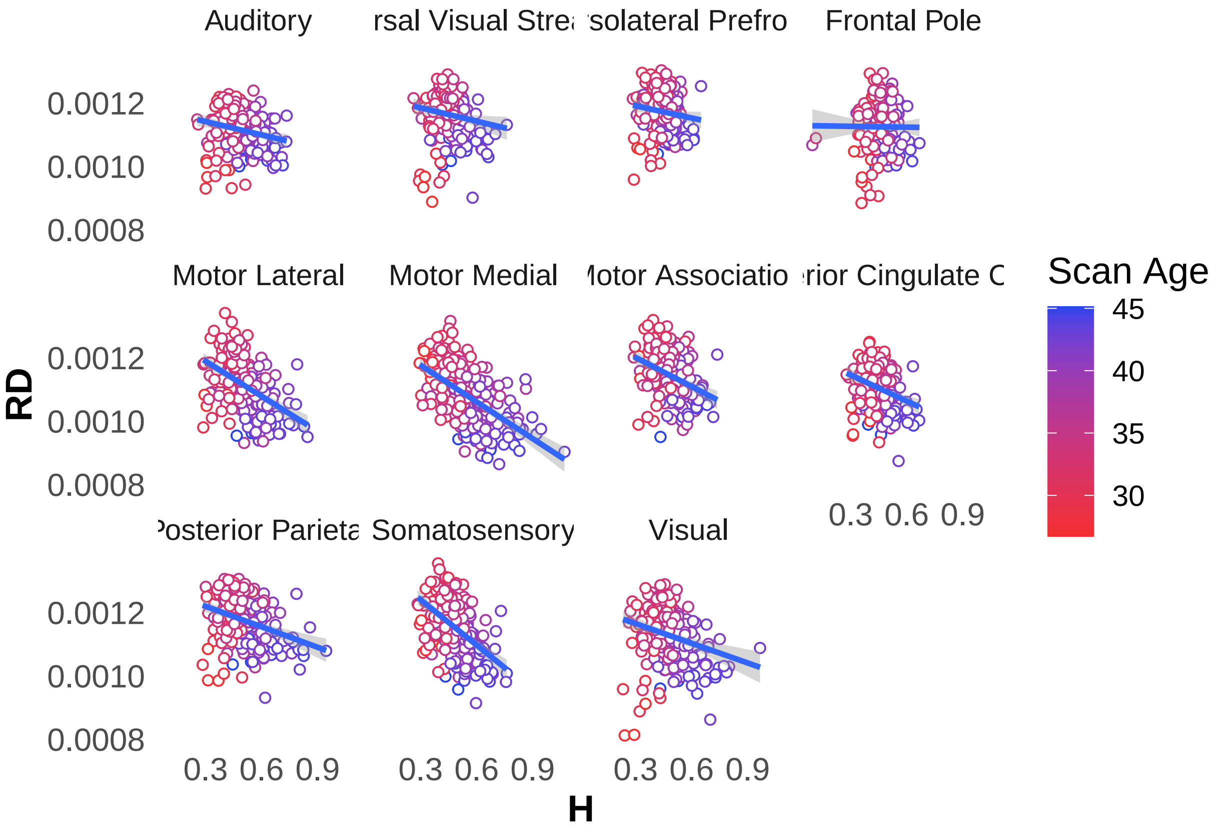


### Supplementary Analysis

#### Sex Differences Analysis

Comparing both sexes in total group Better to look at overall means, sexes may differ at a specific time point.

##### Grey Matter sex differences emmeans

| sex | emmean | SE | df | asymp.LCL | asymp.UCL |
|---|---|---|---|---|---|
| male | 0.5516298 | 0.0049329 | Inf | 0.5419615 | 0.5612980 |
| female | 0.5452315 | 0.0053929 | Inf | 0.5346616 | 0.5558014 |

| contrast | estimate | SE | df | z.ratio | p.value |
|---|---|---|---|---|---|
| female - male | -0.0063983 | 0.0073087 | Inf | -0.8754332 | 0.3813383 |

##### GM Trend

| term | contrast | null.value | estimate | std.error | df | statistic | p.value |
|---|---|---|---|---|---|---|---|
| sex | male - female | 0 | -0.0039376 | 0.0025058 | 624.4119 | -1.571418 | 0.1165921 |

##### White Matter sex differences emmeans

| sex | emmean | SE | df | asymp.LCL | asymp.UCL |
|---|---|---|---|---|---|
| male | 0.4778242 | 0.0028938 | Inf | 0.4721525 | 0.4834959 |
| female | 0.4717174 | 0.0031637 | Inf | 0.4655168 | 0.4779181 |

| contrast | estimate | SE | df | z.ratio | p.value |
|---|---|---|---|---|---|
| female - male | -0.0061067 | 0.0042875 | Inf | -1.424309 | 0.154357 |

**WM Trend**

| term | contrast | null.value | estimate | std.error | df | statistic | p.value |
|---|---|---|---|---|---|---|---|
| sex | male - female | 0 | -0.001264 | 0.001658 | 663.0553 | -0.7623668 | 0.4461123 |

**Combined RSN sex differences emmeans**

| sex | emmean | SE | df | asymp.LCL | asymp.UCL |
|---|---|---|---|---|---|
| male | 0.5417905 | 0.0045696 | Inf | 0.5328343 | 0.5507468 |
| female | 0.5354386 | 0.0049960 | Inf | 0.5256465 | 0.5452306 |

| contrast | estimate | SE | df | z.ratio | p.value |
|---|---|---|---|---|---|
| female - male | -0.0063519 | 0.0067706 | Inf | -0.9381594 | 0.3481625 |

**Combined RSN trend**

| sex | scan_age.trend | p.value | conf.low | conf.high |
|---|---|---|---|---|
| female | 0.0109294 | 0 | 0.0086544 | 0.0132044 |
| male | 0.0098864 | 0 | 0.0075668 | 0.0122060 |

| contrast | estimate | SE | df | z.ratio | p.value |
|---|---|---|---|---|---|
| male - female | -0.001043 | 0.0016577 | Inf | -0.6291862 | 0.5292272 |

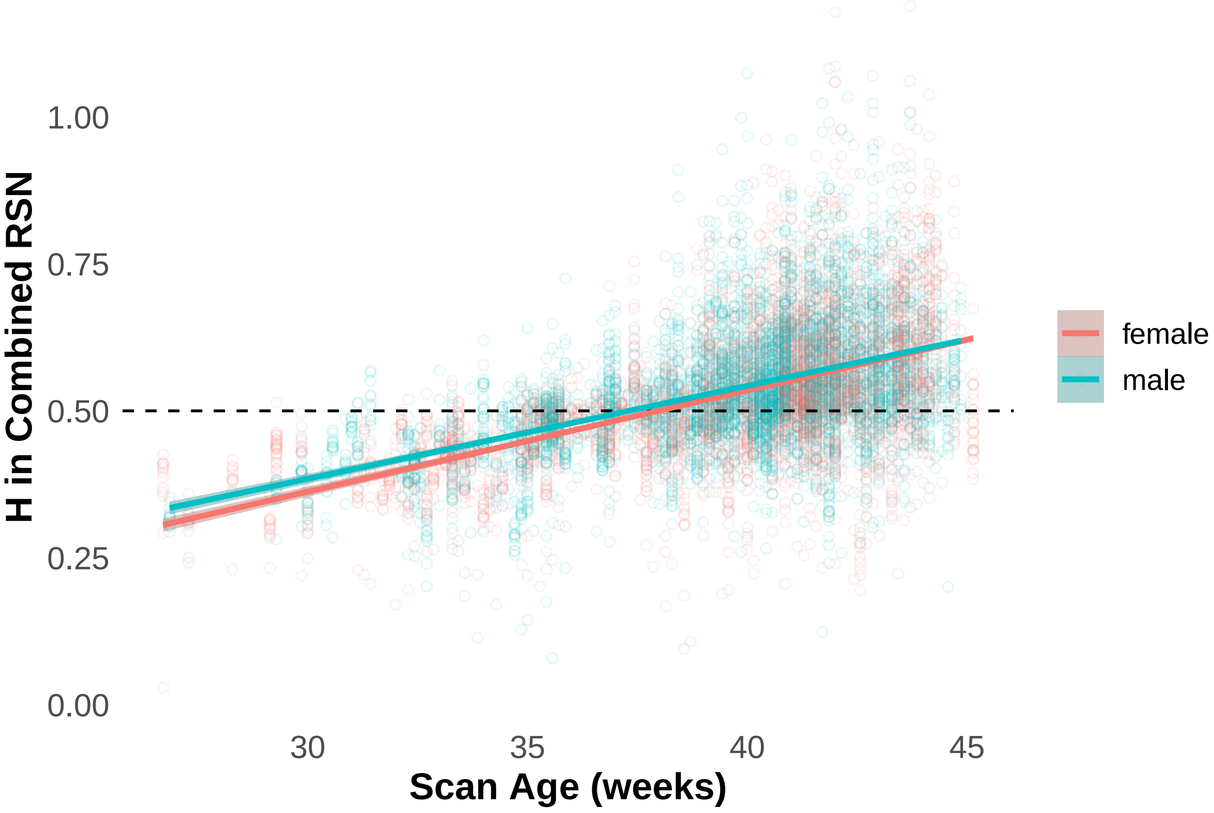

#### Individual RSN Sex Differences

| sex | RSN | emmean | SE | df | asymp.LCL | asymp.UCL |
|---|---|---|---|---|---|---|
| male | H_MotorMedial | 0.6297543 | 0.0047661 | Inf | 0.6204130 | 0.6390956 |
| female | H_MotorMedial | 0.6210321 | 0.0051849 | Inf | 0.6108700 | 0.6311943 |
| male | H_MotorLateral | 0.5998929 | 0.0047661 | Inf | 0.5905516 | 0.6092342 |
| female | H_MotorLateral | 0.5989598 | 0.0051849 | Inf | 0.5887976 | 0.6091219 |
| male | H_Visual | 0.5523317 | 0.0047661 | Inf | 0.5429904 | 0.5616731 |
| female | H_Visual | 0.5499075 | 0.0051849 | Inf | 0.5397453 | 0.5600696 |
| male | H_Auditory | 0.5671673 | 0.0047661 | Inf | 0.5578259 | 0.5765086 |
| female | H_Auditory | 0.5610736 | 0.0051849 | Inf | 0.5509115 | 0.5712358 |
| male | H_MotorAssociation | 0.5458618 | 0.0047661 | Inf | 0.5365205 | 0.5552031 |
| female | H_MotorAssociation | 0.5376202 | 0.0051849 | Inf | 0.5274580 | 0.5477823 |
| male | H_PosteriorParietal | 0.5664201 | 0.0047661 | Inf | 0.5570788 | 0.5757614 |
| female | H_PosteriorParietal | 0.5679882 | 0.0051849 | Inf | 0.5578261 | 0.5781504 |
| male | H_Somatosensory | 0.5502632 | 0.0047661 | Inf | 0.5409219 | 0.5596045 |
| female | H_Somatosensory | 0.5498628 | 0.0051849 | Inf | 0.5397006 | 0.5600249 |
| male | H_DorsalVisualStream | 0.5127851 | 0.0047661 | Inf | 0.5034438 | 0.5221265 |
| female | H_DorsalVisualStream | 0.5155872 | 0.0051849 | Inf | 0.5054251 | 0.5257494 |
| male | H_PosteriorCingulateCortex | 0.4983689 | 0.0047661 | Inf | 0.4890276 | 0.5077102 |
| female | H_PosteriorCingulateCortex | 0.5007377 | 0.0051849 | Inf | 0.4905755 | 0.5108998 |
| male | H_FrontalPole | 0.5170250 | 0.0047661 | Inf | 0.5076837 | 0.5263663 |
| female | H_FrontalPole | 0.5035927 | 0.0051849 | Inf | 0.4934306 | 0.5137549 |
| male | H_Midbrain | 0.4797200 | 0.0047661 | Inf | 0.4703787 | 0.4890613 |
| female | H_Midbrain | 0.4774874 | 0.0051849 | Inf | 0.4673252 | 0.4876495 |
| male | H_DorsolateralPrefrontal | 0.4988617 | 0.0047661 | Inf | 0.4895204 | 0.5082030 |
| female | H_DorsolateralPrefrontal | 0.4924662 | 0.0051849 | Inf | 0.4823041 | 0.5026284 |
| male | H_Cerebellum | 0.4344145 | 0.0047661 | Inf | 0.4250732 | 0.4437558 |
| female | H_Cerebellum | 0.4192718 | 0.0051849 | Inf | 0.4091096 | 0.4294339 |

| contrast | RSN | estimate | SE | df | z.ratio | p.value |
|---|---|---|---|---|---|---|
| female - male | H_MotorMedial | -0.0087222 | 0.0070426 | Inf | -1.2384931 | 1.0000000 |
| female - male | H_MotorLateral | -0.0009332 | 0.0070426 | Inf | -0.1325011 | 1.0000000 |
| female - male | H_Visual | -0.0024243 | 0.0070426 | Inf | -0.3442291 | 1.0000000 |
| female - male | H_Auditory | -0.0060936 | 0.0070426 | Inf | -0.8652516 | 1.0000000 |
| female - male | H_MotorAssociation | -0.0082416 | 0.0070426 | Inf | -1.1702531 | 1.0000000 |
| female - male | H_PosteriorParietal | 0.0015681 | 0.0070426 | Inf | 0.2226649 | 1.0000000 |
| female - male | H_Somatosensory | -0.0004004 | 0.0070426 | Inf | -0.0568580 | 1.0000000 |
| female - male | H_DorsalVisualStream | 0.0028021 | 0.0070426 | Inf | 0.3978750 | 1.0000000 |
| female - male | H_PosteriorCingulateCortex | 0.0023687 | 0.0070426 | Inf | 0.3363443 | 1.0000000 |
| female - male | H_FrontalPole | -0.0134323 | 0.0070426 | Inf | -1.9072901 | 0.6777964 |
| female - male | H_Midbrain | -0.0022326 | 0.0070426 | Inf | -0.3170200 | 1.0000000 |
| female - male | H_DorsolateralPrefrontal | -0.0063955 | 0.0070426 | Inf | -0.9081096 | 1.0000000 |
| female - male | H_Cerebellum | -0.0151428 | 0.0070426 | Inf | -2.1501661 | 0.4100470 |

#### Standard Deviation BOLD Analysis

Determining how H relates to basic signal variability of BOLD standard deviation. In adults, there is a positive correlation between BOLD SD and H. Correlation between BOLD SD and H in the whole brain and in the groups.

##### Correlations BOLD SD and H Whole Group

Gray Matter Whole Group

| estimate | statistic | p.value | parameter | conf.low | conf.high | method | alterna |
|---|---|---|---|---|---|---|---|
| -0.0470582 | -4.512777 | 6.5e-06 | 9176 | -0.0674523 | -0.0266249 | Pearson's product-moment correlation | two.sic |

Preterm Group GM

| estimate | statistic | p.value | parameter | conf.low | conf.high | method | alternativ |
|---|---|---|---|---|---|---|---|
| 0.0746183 | 4.28151 | 1.91e-05 | 3274 | 0.0404761 | 0.1085864 | Pearson's product-moment correlation | two.sided |

White Matter Whole Group

| estimate | statistic | p.value | parameter | conf.low | conf.high | method | alternative |
|---|---|---|---|---|---|---|---|
| 0.098176 | 9.450069 | 0 | 9176 | 0.0778734 | 0.1183971 | Pearson's product-moment correlation | two.sided |

Preterm Group WM

| estimate | statistic | p.value | parameter | conf.low | conf.high | method | alternative |
|---|---|---|---|---|---|---|---|
| 0.2579145 | 15.27434 | 0 | 3274 | 0.225662 | 0.2896022 | Pearson's product-moment correlation | two.sided |

Combined RSN Whole Group

| estimate | statistic | p.value | parameter | conf.low | conf.high | method | alterna |
|---|---|---|---|---|---|---|---|
| -0.2380667 | -23.47982 | 0 | 9176 | -0.2572726 | -0.2186728 | Pearson's product-moment correlation | two.sic |

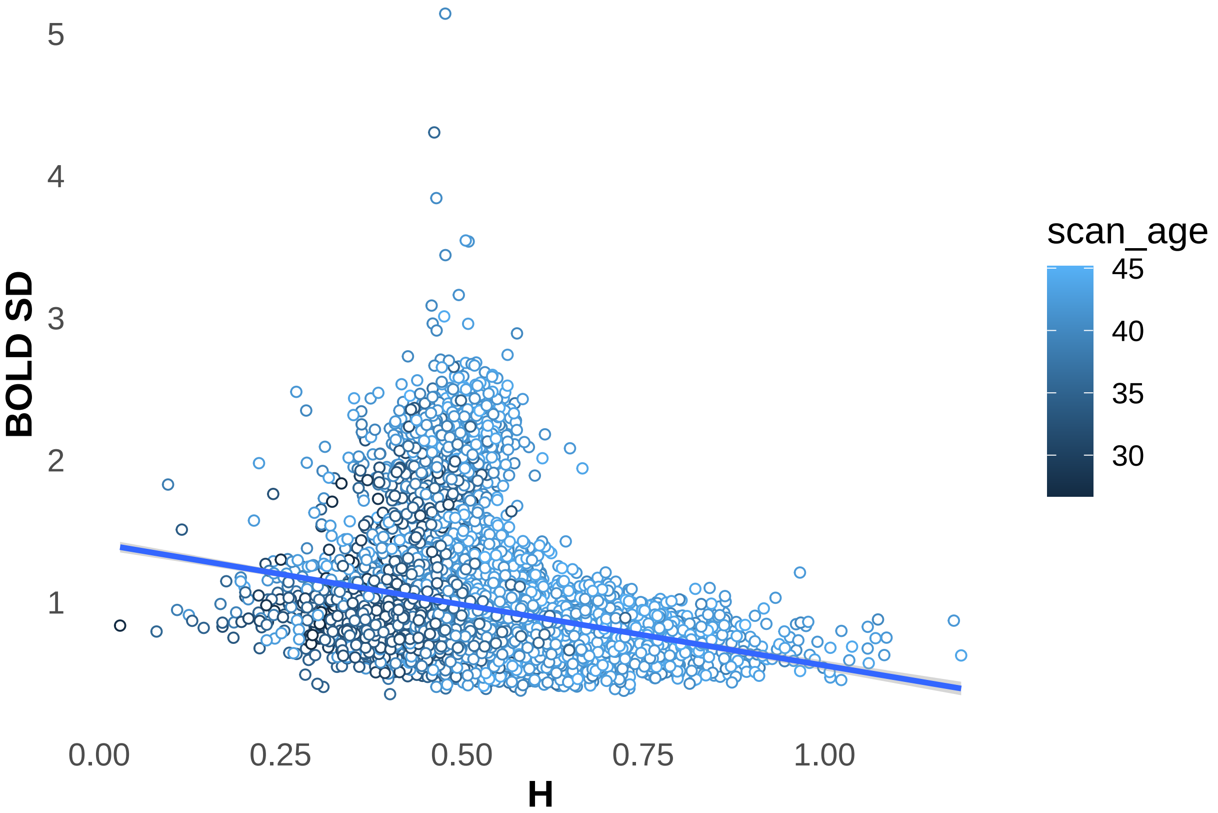


There is a weak negative correlation in the combined RSN.

Preterm Group combined RSN

| estimate | statistic | p.value | parameter | conf.low | conf.high | method | alterna |
|---|---|---|---|---|---|---|---|
| -0.1235593 | -7.124521 | 0 | 3274 | -0.1571401 | -0.0896932 | Pearson's product-moment correlation | two.sic |

Preterm RSN analysis BOLD SD and PMA

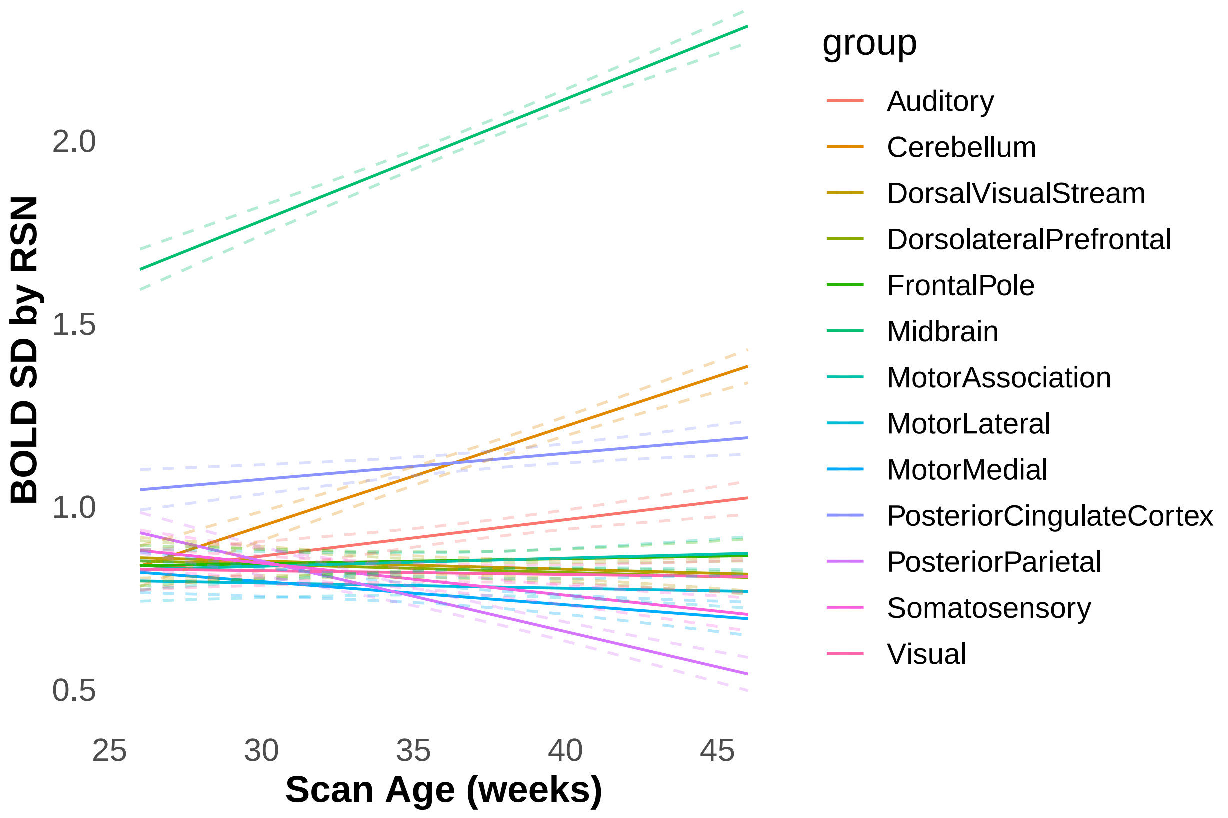

BOLD SD by RSN
2.0
1.5
1.0
0.5
25
30
35
40
45
Scan Age (weeks)
group
Auditory
Cerebellum
DorsalVisualStream
DorsolateralPrefrontal
FrontalPole
Midbrain
MotorAssociation
MotorLateral
MotorMedial
PosteriorCingulateCortex
PosteriorParietal
Somatosensory
Visual
