## Supplementary material for "Temporal Complexity of the BOLD-Signal In Preterm Versus Term Infants": All supplementary data: eLife_RAnalysis.Rmd

````
---
title: "dHCP H Study of Temporal Complexity (Nov, 2023)"
output:
  pdf_document: default
  html_document: default
date: "`r format(Sys.time(), '%d %B, %Y')`"
---

# Intro

We are assessing a measure of the BOLD temporal complexity, H, in preterm born infants compared to term born controls.

Manuscript Title: **Temporal complexity of BOLD-signal in preterm versus term infants**

Study Questions:
 1) How does H develop over time? \newline
 2) Do different RSNs develop differently with regards to H values? \newline
 3) Do term born infants have greater H compared to preterm infants at scan age? \newline
 4) Is H associated with indirect measures of myelination (FA and RD)? 

Linear mixed effects models will be created to take into account of random effects and interacting terms. Significant differences will be determined through using estimated marginal means (emmeans) and trends (emtrends). Multiple comparisons will be controlled for using Holm's method.

```{r, echo=F, message=F, warning=F}
#Loading all the required libraries
knitr::opts_chunk$set(echo = TRUE)
library(broom)
library(tidyverse)
library(ggplot2)
library(emmeans)
library(lme4)
library(ggeffects)
library(ggpubr)
library(directlabels) # for labelling lines on ggplot
library(forcats)
library(DescTools)
library(report)
library(rstatix)
library(kableExtra)
```

```{r, echo=F, message=F, warning=F}
#Housekeeping
theme_minimalism <- function(){

theme_minimal() + # ggplot′s minimal theme hides many unnecessary features of plot
theme( # make modifications to the theme
panel.grid.major.y=element_blank(), # hide major grid for y axis
panel.grid.minor.y=element_blank(), # hide minor grid for y axis
panel.grid.major.x=element_blank(), # hide major grid for x axis
panel.grid.minor.x=element_blank(), # hide minor grid for x axis
text=element_text(size=14), # font aesthetics
axis.text=element_text(size=12),
axis.title=element_text(size=14, face="bold"))
}

RSN_names <- list(
  'H_MotorMedial' = "Motor Medial",
  'H_MotorLateral' = "Motor Lateral",
  'H_Visual' = "Visual",
  'H_Auditory'= "Auditory",
  'H_MotorAssociation' = "Motor Association",
  'H_PosteriorParietal' = "Posterior Parietal",
  'H_Somatosensory' = "Somatosensory",
  'H_DorsalVisualStream' = "Dorsal Visual Stream",
  'H_PosteriorCingulateCortex' = "Posterior Cingulate Cortex",
  'H_FrontalPole'= "Frontal Pole",
  'H_Midbrain'= "Midbrain",
  'H_DorsolateralPrefrontal'= "Dorsolateral Prefrontal",
  'H_Cerebellum'= "Cerebellum"
)

RSN_labeller <- function(variable,value){
  return(RSN_names[value])
}

# Helper function for string wrapping. 
# Default 20 character target width.
swr = function(string, nwrap=10) {
  paste(strwrap(string, width=nwrap), collapse="\n")
}
swr = Vectorize(swr)
```

# Load data
```{r}
# Data Loading
##CSV file load
ClinicalInfo <- read.csv("csvfiles/ClinicalInfo.csv")
H_Tissue <- read.csv("csvfiles/new_Hvalues.csv")
H_RSN <- read.csv("csvfiles/new_HvalinRSN_wFIX.csv")
#Data merging 
fulldata <- merge(ClinicalInfo,H_Tissue)
fulldata <- merge(fulldata, H_RSN)
```

After loading our data, we have `r fulldata %>% nrow()` rows.

We will exclude subjects with large motion (meanFD)
```{r}
meanFD <- read.csv("csvfiles/meanFD.csv")
newdata <- merge(fulldata, meanFD)
newdata <- newdata %>% filter(meanFD < 0.5)
```

Which leaves us with `r newdata %>% nrow()` rows. That means we lost `r fulldata %>% nrow() - newdata %>% nrow()` rows.

```{r, echo=F, message=F, warning=F}
latexvariable <- c('Initial_Number_Rows', 'Number_Rows_After_meanFD', 'Number_of_rows_lost')
latexvalue <- c(fulldata %>% nrow(), newdata %>% nrow(), fulldata %>% nrow() - newdata %>% nrow())
```

```{r, echo=F, message=F, warning=F}
oldfulldata <- fulldata
fulldata <- newdata
```

# GAfactors

Sorting the participants into 3 groups according to their gestational age (i.e. birth age). \newline
VPT = very preterm \newline
MPT = moderately preterm \newline
THC = term healthy control \newline

```{r}
fulldata <- fulldata %>% mutate(GAfactor = case_when(
    birth_age <= 32  ~ "VPT",
    birth_age > 32 & birth_age <= 37 ~ "MPT",
    birth_age >37 ~ "THC"
    ))
fulldata$GAfactor <- as.factor(fulldata$GAfactor)
fulldata$GAfactor <- factor(fulldata$GAfactor, levels = c("VPT", "MPT", "THC"))
```

```{r, echo=F,message=F,warning=F}
oldfulldata <- oldfulldata %>% mutate(GAfactor = case_when(
    birth_age <= 32  ~ "VPT",
    birth_age > 32 & birth_age <= 37 ~ "MPT",
    birth_age >37 ~ "THC"
    ))
oldfulldata$GAfactor <- as.factor(oldfulldata$GAfactor)
oldfulldata$GAfactor <- factor(oldfulldata$GAfactor, levels = c("VPT", "MPT", "THC"))
```

## Setting ages for when we will estimate marginal means and trends

Using median birth ages in the VPT, MPT and THC groups \newline
VPT = 29 weeks GA \newline
MPT = 35 weeks GA \newline
THC = 41 weeks GA \newline

preterm comparison age = 35 weeks PMA \newline
term comparison age = 41 weeks PMA


```{r, echo=F,message=F,warning=F}
# MORE HOUSEKEEPING
#Rename RSN columns 
fulldata <- fulldata %>% rename(
  'H_MotorMedial' = RSN_Sept2022_0000,
  'H_MotorLateral' = RSN_Sept2022_0001,
  'H_Visual' = RSN_Sept2022_0002,
  'H_Auditory' = RSN_Sept2022_0003and0006,
  'H_MotorAssociation' = RSN_Sept2022_0004,
  'H_PosteriorParietal' = RSN_Sept2022_0005,
  'H_Somatosensory' = RSN_Sept2022_0007and0011,
  'H_DorsalVisualStream' = RSN_Sept2022_0008and0012,
  'H_PosteriorCingulateCortex' = RSN_Sept2022_0009, 
  'H_FrontalPole' = RSN_Sept2022_0013,
  'H_Midbrain' = RSN_Sept2022_0014,
  'H_DorsolateralPrefrontal' = RSN_Sept2022_0015, 
  'H_Cerebellum' = RSN_Sept2022_0029
  )

# PIVOT LONG
fulldata_long <- fulldata %>% pivot_longer(cols = starts_with("H_"), names_to = "RSN", values_to = "H")
fulldata_long$GAfactor <- factor(fulldata_long$GAfactor, levels = c("VPT", "MPT", "THC"))
fulldata_long$RSN <- factor(fulldata_long$RSN, levels = c("H_MotorMedial", "H_MotorLateral", "H_Visual", "H_Auditory", "H_MotorAssociation", "H_PosteriorParietal", "H_Somatosensory", "H_DorsalVisualStream", "H_PosteriorCingulateCortex","H_FrontalPole", "H_Midbrain", "H_DorsolateralPrefrontal", "H_Cerebellum"))
```

# Initial Linear Mixed Effects Model Checks

Our main model will be Hurst with fixed effects: Birth Age, Scan Age, and ROI

Subjects will be set as random effects to take into account multiple scans of the same participant.

The first question we have is should Birth Age and Scan Age be interactive terms? (i.e. Birth Age + Scan Age or Birth Age * Scan Age)

We will use an anova of the two models to decide:

```{r}
# Setting up our linear mixed effects model. Determining whether we will need to to take 
# into account of PMA and GA.
# First: Model with no interaction:
h.model1 <- lmer(data = fulldata_long, H ~ scan_age + birth_age + (1|Subject), REML = F)
```

```{r}
#Nect: Model with interaction
h.model.int2 <- lmer(data = fulldata_long, H ~ scan_age*birth_age + (1|Subject), REML = F)
```

```{r}
# test for significance of interaction
GAPMAanova <- anova(h.model.int2, h.model1)
GAPMAanova
# If p > 0.05, there is no difference between these models, so we wouldn't need to include 
# PMA and GA as interactive terms
```

As p < 0.05, (check: `r any(GAPMAanova$'Pr(>Chisq)' < 0.05)`), we will include Birth Age and Scan Age as interactive terms.


# Demographics

```{r}
distinctn_old <- oldfulldata %>% distinct(Subject) %>% summarise(Observations = n())
```

Our original number of distinct subjects was `r distinctn_old`

```{r}
distinctn <- fulldata %>% distinct(Subject) %>% summarise (Observations = n())
```

Our number of distinct subjects after filtering for meanFD is: `r distinctn`

```{r, echo=F, message=F, warning=F}
latexvalue <- as.numeric(latexvalue)
latexdf <- data.frame(latexvariable, latexvalue)
latexdf
```
```{r, echo=F, message=F, warning=F}
latexdf <- latexdf %>% add_row(latexvariable = 'distinctn', latexvalue = as.numeric(distinctn))
latexdf <- latexdf %>% add_row(latexvariable = 'distinctn_old', latexvalue = as.numeric(distinctn_old))
```

## Range of GA

```{r, echo=F, message=F, warning=F}
minga = min(fulldata$birth_age)
maxga = round(max(fulldata$birth_age),2)
```

The minimum and maximum gestational age of our entire cohort is `r minga` and `r maxga`, respectively.

```{r, echo=F, message=F, warning=F}
latexdf <- latexdf %>% add_row(latexvariable = 'minga', latexvalue = as.numeric(minga))
latexdf <- latexdf %>% add_row(latexvariable = 'maxga', latexvalue = as.numeric(maxga))
```

## Group Sample Sizes
```{r, echo=F, message=F, warning=F}
gafactordistinctn_old <- oldfulldata %>% group_by(GAfactor) %>% distinct(Subject) %>% summarise (Observations = n())
gafactordistinctn <- fulldata %>% group_by(GAfactor) %>% distinct(Subject) %>% summarise (Observations = n())
```

The initial amount of distinct subjects from each GAfactor group was:
```{r, echo=F, message=F, warning=F}
knitr::kable(gafactordistinctn_old)
```

After meanFD filtering, we are left with:
```{r, echo=F, message=F, warning=F}
knitr::kable(gafactordistinctn)
```

```{r, echo=F, message=F, warning=F}
latexdf <- latexdf %>% add_row(latexvariable = 'numorigthc', latexvalue = as.numeric(gafactordistinctn_old$Observations[3]))
latexdf <- latexdf %>% add_row(latexvariable = 'numnewthc', latexvalue = as.numeric(gafactordistinctn$Observations[3]))
```

The number of preterm infants with both preterm and term scans is:

```{r, echo=F, message=F, warning=F}
numpreandtermscan <- fulldata %>% filter(birth_age <= 37) %>% filter(duplicated(Subject) == TRUE) %>% group_by(GAfactor) %>% summarise (Observations = n())
knitr::kable(numpreandtermscan)
```

```{r, echo=F, message=F, warning=F}
numjustpreterm <- fulldata %>% filter(birth_age <= 37) %>% filter(scan_age <= 37) %>% group_by(GAfactor) %>% summarise (Observations = n())
latexdf <- latexdf %>% add_row(latexvariable = 'numjustpreterm', latexvalue = as.numeric(sum(numjustpreterm$Observations) - sum(numpreandtermscan$Observations)))
```

```{r, echo=F, message=F, warning=F}
numjusttea <- fulldata %>% filter(birth_age <= 37)  %>% filter(scan_age > 37) %>% group_by(GAfactor) %>% summarise (Observations = n())
latexdf <- latexdf %>% add_row(latexvariable = 'numjusttea', latexvalue = as.numeric(sum(numjusttea$Observations) - sum(numpreandtermscan$Observations)))
```

```{r, echo=F, message=F, warning=F}
latexdf <- latexdf %>% add_row(latexvariable = 'numbothpreandterm', latexvalue = as.numeric(sum(numpreandtermscan$Observations)))
latexdf <- latexdf %>% add_row(latexvariable = 'numpreterm', latexvalue = as.numeric(sum(gafactordistinctn$Observations[1] + gafactordistinctn$Observations[2])))
```


Thus, the resulting preterm group was composed of `r gafactordistinctn$Observations[1] + gafactordistinctn$Observations[2]` infants including `r sum(numjustpreterm$Observations) - sum(numpreandtermscan$Observations)` infants with preterm only scans, `r sum(numjusttea$Observations) - sum(numpreandtermscan$Observations)` infants with term-equivalent only scans, and `r sum(numpreandtermscan$Observations)` infants with both preterm and term-equivalent aged scans.


```{r, echo=F, message=F, warning=F}

library(VennDiagram)
venn.plot <- draw.pairwise.venn(area1 = sum(numjustpreterm$Observations), area2 = sum(numjusttea$Observations), cross.area = sum(numpreandtermscan$Observations), category = c("Preterm", "TEA"), fill = c("blue", "red"))
venn.plot
grid.draw(venn.plot);
grid.newpage();
```

## Basic Stats

Non-parametric stats because our data is not normally distributed.

### Birth ages
Preterm birth median age in VPT and MPT
```{r, echo=F,message=F,warning=F}
# Preterm groups median scan age at preterm scan without repeats 
knitr::kable(fulldata %>% filter(duplicated(Subject) == FALSE) %>% group_by(GAfactor) %>% summarise (Observations = n(), Median.BirthAge = round(median(birth_age),2), Range = round(range(birth_age, na.rm=TRUE),2)))
```

### Scan ages/ PMA (Median)
Preterm scan median age in VPT and MPT
```{r, echo=F,message=F,warning=F}
knitr::kable(fulldata %>% filter(scan_age <= 37) %>% group_by(GAfactor) %>% summarise (Observations = n(), Median.Scanage = median(scan_age), Range = range(scan_age, na.rm=TRUE)))
```

TEA scan median age in VPT and MPT
```{r, echo=F,message=F,warning=F}
# Preterm groups median scan age at TEA scan without repeats 
knitr::kable(fulldata %>% filter(scan_age > 37, !GAfactor == "THC") %>% group_by(GAfactor) %>% summarise (Observations = n(), Median.Scanage = median(scan_age), Range = range(scan_age, na.rm=TRUE)))
```

Preterm scan median age in combined preterm group
```{r, echo=F,message=F,warning=F}
# Combined VPT, MPT median scan age at preterm scan
knitr::kable(fulldata %>% filter(duplicated(Subject) == FALSE) %>% filter(scan_age <= 37) %>% summarise (Observations = n(), Median.Scanage = median(scan_age)))
```

TEA scan median age in combined preterm group
```{r, echo=F,message=F,warning=F}
# Combined VPT, MPT median scan age at TEA scan
knitr::kable(fulldata %>% filter(duplicated(Subject) == FALSE) %>% filter(scan_age > 37, !GAfactor == "THC") %>% summarise (Observations = n(), Median.Scanage = median(scan_age))) #preterm group at TEA scan
```

Term scan median age in all groups
```{r, echo=F,message=F,warning=F}
# All groups scan age at TEA
knitr::kable(fulldata %>% filter(duplicated(Subject) == FALSE) %>% filter(scan_age > 37) %>% group_by(GAfactor) %>% summarise (Observations = n(), Median.Scanage = median(scan_age), Range = range(scan_age, na.rm=TRUE)))
```

THC scan median age
```{r, echo=F,message=F,warning=F}
# THC group scan age 
knitr::kable(fulldata %>% filter(duplicated(Subject) == FALSE) %>% filter(scan_age > 37, GAfactor == "THC") %>% summarise (Observations = n(), Median.Scanage = median(scan_age))) #THC group
```


### Birth Age/Gestational Age

```{r, echo=F,message=F,warning=F}
knitr::kable(fulldata %>% filter(duplicated(Subject) == FALSE) %>% group_by(GAfactor) %>% summarise (Observations = n(), Median.Birthage = round(median(birth_age),2), Range = round(range(birth_age),2), n = n()))
```

### Birth Weight
```{r, echo=F,message=F,warning=F}
knitr::kable(fulldata %>% filter(duplicated(Subject) == FALSE) %>% group_by(GAfactor) %>% summarise (Observations = n(), Median.Birthweight = median(birth_weight)))
```

### Head Circumference
```{r, echo=F,message=F,warning=F}
knitr::kable(fulldata %>% filter(duplicated(Subject) == FALSE) %>% group_by(GAfactor) %>% summarise (Observations = n(), Median.Headcircumference = median(head_circumference_scan)))
```


### Sedation
```{r, echo=F,message=F,warning=F}
#knitr::kable(fulldata %>% filter(sedation == 'True')  %>% distinct(Subject) %>% summarise (Observations = n()))
knitr::kable(fulldata %>% filter(sedation == 'True') %>% group_by(GAfactor) %>% summarise (Observations = n()))
```

### Radiology scores
5=Incidental finding with possible / likely significance for both clinical and imaging analysis (e.g. Major lesions within white matter cortex, cerebellum and or basal ganglia; small head / brain < 1 st percentile

**Radiology Score of 1:**
```{r, echo=F,message=F,warning=F}
radscore1.tot <- fulldata %>% filter(radiology_score == 1) %>% distinct(Subject) %>% summarise (Observations = n())
```

Total = `r radscore1.tot`

Breakdown by GAfactor:

```{r, echo=F,message=F,warning=F}
knitr::kable(radscore1.ga <- fulldata %>% filter(radiology_score == 1) %>% group_by(GAfactor) %>% distinct(Subject) %>% summarise (Observations = n()))
```

**Radiology Score of 2:**

```{r, echo=F,message=F,warning=F}
radscore2.tot <- fulldata %>% filter(radiology_score == 2) %>% distinct(Subject) %>% summarise (Observations = n())
```

Total = `r radscore2.tot`

Breakdown by GAfactor:

```{r, echo=F,message=F,warning=F}
knitr::kable(fulldata %>% filter(radiology_score == 2) %>% group_by(GAfactor) %>% distinct(Subject) %>% summarise (Observations = n()))
```

**Radiology Score of 3:**

```{r, echo=F,message=F,warning=F}
radscore3.tot <- fulldata %>% filter(radiology_score == 3) %>% distinct(Subject) %>% summarise (Observations = n())
```

Total = `r radscore3.tot`

Breakdown by GAfactor:

```{r, echo=F,message=F,warning=F}
knitr::kable(fulldata %>% filter(radiology_score == 3) %>% group_by(GAfactor) %>% distinct(Subject) %>% summarise (Observations = n()))
```

**Radiology Score of 4:**

```{r, echo=F,message=F,warning=F}
radscore4.tot <- fulldata %>% filter(radiology_score == 4) %>% distinct(Subject) %>% summarise (Observations = n())
```

Total = `r radscore4.tot`

Breakdown by GAfactor:

```{r, echo=F,message=F,warning=F}
knitr::kable(fulldata %>% filter(radiology_score == 4) %>% group_by(GAfactor) %>% distinct(Subject) %>% summarise (Observations = n()))
```

**Radiology Score of 5:**

```{r, echo=F,message=F,warning=F}
radscore5.tot <- fulldata %>% filter(radiology_score == 5) %>% distinct(Subject) %>% summarise (Observations = n())
```

Total = `r radscore5.tot`

Breakdown by GAfactor:

```{r, echo=F,message=F,warning=F}
knitr::kable(fulldata %>% filter(radiology_score == 5) %>% group_by(GAfactor) %>% distinct(Subject) %>% summarise (Observations = n()))
```


### Sexes
**Female**
```{r, echo=F,message=F,warning=F}
totalfemale <- fulldata %>% filter(sex == 'female') %>% distinct(Subject)%>% summarise (Observations = n()) #female = 328
```

Total = `r totalfemale`

Breakdown by GAfactor:

```{r, echo=F,message=F,warning=F}
knitr::kable(fulldata %>%  group_by(GAfactor) %>% filter(sex == 'female') %>% distinct(Subject) %>% summarise (Observations = n()))
```


**Male**
```{r, echo=F,message=F,warning=F}
totalmale <- fulldata %>% filter(sex == 'male') %>% distinct(Subject)%>% summarise (Observations = n()) #male = 388
```

Total = `r totalmale`

Breakdown by GAfactor:

```{r, echo=F,message=F,warning=F}
knitr::kable(fulldata %>% group_by(GAfactor) %>% filter(sex == 'male') %>% distinct(Subject) %>% summarise (Observations = n()))
```

## Birth age and scan age colinearity?
Preterm (combined) and term group scan age histogram

Checking to see that both groups are approximately scanned at the same age to make sure we are not extrapolating data when predicted H values at set scan age. 

Emmeans at 41 weeks, controls for colinearity issue. 
```{r, echo=F,message=F,warning=F}
RemovedpretermGAfactors <- fulldata_long%>% filter(scan_age > 37)
```


```{r, echo=F, message=F, warning=F}
RemovedpretermGAfactors %>% distinct(Subject, scan_age, birth_age, GAfactor, RSN, H) %>% 
  pivot_wider(names_from = RSN, values_from=H) %>% 
  ggplot(aes(y=birth_age, x=scan_age)) + geom_point() + 
  geom_smooth(method="lm") + theme_minimalism() +
  xlab("Scan Age/PMA (weeks)") +
  ylab("Birth Age/GA (weeks)") + 
  ggtitle("GA vs PMA for Scan Ages > 37")
```
```{r, echo=F,message=F,warning=F}
ggsave("pngfigures/GAvsPMA_PMAgt37.png", width = 20, height = 10, units = "cm", dpi = 600)
```


Preterm groups and term group scan age histogram 
```{r,message=F}
RemovedpretermGAfactors %>% distinct(Subject, scan_age, GAfactor) %>% 
  filter(scan_age > 37) %>% 
  ggplot(aes(x=scan_age, color=GAfactor)) + 
  geom_histogram(fill="white",alpha=0.5, position="identity") + 
  theme_minimalism() + 
  scale_color_manual(values = c("indianred1", "goldenrod1", "deepskyblue"), 
                     name = "Group", labels = c("VPT", "MPT", "THC")) +
  xlab("Scan Age/PMA (weeks)") +
  ylab("Count") +
  ggtitle("PMA Histogram for Scan Ages > 37")
```
```{r, echo=F,message=F,warning=F}
ggsave("pngfigures/PMAHist_PMAgt37.png", width = 20, height = 10, units = "cm", dpi = 600)
```

Correlation test
```{r, echo=F,message=F,warning=F}
gapma.cortest <- RemovedpretermGAfactors %>% pivot_wider(names_from = RSN, values_from=H) %>% do(tidy(cor.test(.$scan_age,.$birth_age)))
knitr::kable(gapma.cortest %>% mutate(across(where(is.numeric), round, 3)) %>% select(estimate, p.value, conf.low, conf.high, method))
```

# Linear Mixed Effects Model: GM in Entire Cohort

Looking at GA as a continuous variable in the grey matter. 

Let's set-up our model for H in the grey matter, with scan age and birth age as interative terms

```{r}
GMwholecohortmodel <- lmer(data = fulldata, HinGM ~ scan_age*birth_age + 
                             (1|Subject), REML = F)
```

Making sure birth age and scan age should be interative terms
```{r, echo=F,message=F,warning=F}
#making sure they should be interative terms
GMwholecohortmodel_ <- lmer(data = fulldata, HinGM ~ scan_age + (1|Subject), REML = F)
knitr::kable(papeR::prettify(anova(GMwholecohortmodel,GMwholecohortmodel_)))
```

Summary of full model:
```{r, echo=F,message=F,warning=F}
knitr::kable(papeR::prettify(summary(GMwholecohortmodel)))
```
Report blurb:

```{r, echo=F, message=F, warning=F}
GMwholecohortreport <- report(GMwholecohortmodel)
```

`r GMwholecohortreport`

## Raw correlation

H in GM vs birth age:

```{r, echo=F,message=F,warning=F}
knitr::kable(fulldata %>% do(tidy(cor.test(.$HinGM, .$birth_age))))
```

H in GM vs scan age:

```{r, echo=F,message=F,warning=F}
knitr::kable(fulldata %>% do(tidy(cor.test(.$HinGM, .$scan_age))))
```

```{r,message=F}
(HGMvsGA <- ggplot(data = fulldata, aes(x = birth_age, y = HinGM)) +
  geom_point(shape=1, alpha = 1/3) + geom_smooth(method = "lm", se=T) + 
  theme_minimalism() +
  xlab("Birth age (weeks)") +
  ylab("H in the GM") +
  theme(legend.title=element_blank()))
```

```{r, echo=F,message=F,warning=F}
ggsave("pngfigures/HGMvsGA.png", width = 20, height = 10, units = "cm", dpi = 600)
```

Whole group GM plot

```{r, echo=F,message=F,warning=F}
(HGMvsSCplot <- ggplot(data = fulldata, aes(x = scan_age, y = HinGM)) +
  geom_point(shape=1, alpha = 1/3) + geom_smooth(method = "lm", se=T) + 
  theme_minimalism() +
  xlab("Scan Age (weeks)") +
  ylab("H in the GM") +
  theme(legend.title=element_blank()))
```

```{r, echo=F,message=F,warning=F}
ggsave("pngfigures/HGMvsPMA.png", width = 20, height = 10, units = "cm", dpi = 600)
```

## How does H develop with respect to birth age? 

Let's look at the estimated marginal trend, keeping scan age constant at 41weeks ("term age")

```{r, echo=F, message=F, warning=F}
GMwholecohortTREND <- emtrends(GMwholecohortmodel, ~ scan_age, at = list(scan_age = c(41)), var = "birth_age")
GMwholecohortTREND
```

So if we scan a baby at term age (41 weeks), the rate of change of H in GM with respect to birth age is `r round(tidy(GMwholecohortTREND)$birth_age.trend, 4)`


# Linear Mixed Effects Model: White Matter

Model, summary and report:

```{r, echo=F, message=F, warning=F}
WMwholecohortmodel <- (lmer(data = fulldata, HinWM ~ scan_age*birth_age + (1|Subject), REML = F))
summary(WMwholecohortmodel)
report(lmer(data = fulldata, HinWM ~ scan_age*birth_age + (1|Subject), REML = F))
```


# Linear Mixed Effects Model: RSN


Model summary:

```{r, echo=F, message=F, warning=F}
RSNwholecohortmodel <- (lmer(data = fulldata_long, H ~ scan_age*birth_age*RSN + (1|Subject), REML = F))
summary(RSNwholecohortmodel)
```
Report:

```{r, echo=F, message=F, warning=F}
report(RSNwholecohortmodel)
```


Report in just the Motor Medial as example:

```{r, echo=F, message=F, warning=F}
report(lmer(data = fulldata, H_MotorMedial ~ scan_age*birth_age + (1|Subject), REML = F))
```


# Just Preterm Group:

The preterm group will be assessed by comparison of the preterm scans to term age (TEA) scans and between the VPT and MPT groups.

First, let's remove the THC group from the data (not shown)
```{r, echo=F, message=F, warning=F}
RemovedTHC_fulldata <- fulldata %>% filter(!GAfactor == "THC") 
RemovedTHC_fulldata_long <- fulldata_long %>% filter(!GAfactor == "THC") 
```

Double checking that we are only analyzing preterm infants.
```{r, echo=F, message=F, warning=F}
numobs <- RemovedTHC_fulldata%>% filter(birth_age > 37) %>% summarise (Observations = n())
```

This number should be 0: `r numobs`

## Group Numbers 

Number of preterm and TEA scans. 

```{r, echo=F,message=F,warning=F}
RemovedTHC_fulldata <- RemovedTHC_fulldata %>% mutate(GAfactor2 = case_when(
    scan_age <= 37 ~ "PT",
    scan_age >37 ~ "T"
    ))
RemovedTHC_fulldata$GAfactor2 <- as.factor(RemovedTHC_fulldata$GAfactor2)
RemovedTHC_fulldata$GAfactor2 <- factor(RemovedTHC_fulldata$GAfactor2, levels = c("PT", "T"))

preandtea_table <- RemovedTHC_fulldata %>% group_by(Subject, GAfactor2) %>% summarise (Observations = n()) %>% group_by(Observations, GAfactor2) %>% count()

knitr::kable(preandtea_table)
```

2 preterm infants scanned twice at TEA (CC00098AN17 and CC00688XX21).

## Preterm vs TEA Comparison 
Will now compare the preterm group at preterm age (35 weeks) and at TEA (41 weeks) in the grey matter, white matter, combined RSN, and individual RSN. \newline
These ages were used as they the median values of the preterm group at preterm and TEA scan age (PMA).The estimated marginal means will be calculated. Linear mixed effects models are created with scan age. 

### Grey Matter H comparison Preterm vs TEA

Estimated Marginal Means:
```{r, echo=F,message=F,warning=F}
h.model.int.fullnoTHC_GM <- lmer(data = RemovedTHC_fulldata, HinGM ~ scan_age*birth_age + (1|Subject), REML = F)
(GMage34vs41 <- h.model.int.fullnoTHC_GM %>% emmeans(~scan_age, at = list(scan_age = c(34.785, 41))))

```

Comparison:

```{r,echo=F,message=F,warning=F}
GMage34vs41 %>% contrast("revpairwise") %>% summary(by = NULL, adjust = "holm")
```

Slopes / Trends:

```{r, echo=F,message=F,warning=F}
(GMpretermtrendTEA <- emtrends(h.model.int.fullnoTHC_GM, ~ birth_age, var = "scan_age"))
```

### White Matter H comparison Preterm vs TEA

Estimated marginal means:

```{r, echo=F,message=F,warning=F}
h.model.int.fullnoTHC_WM <- lmer(data = RemovedTHC_fulldata, HinWM ~ scan_age*birth_age + (1|Subject), REML = F)
(WMage34vs41 <- h.model.int.fullnoTHC_WM %>% emmeans(~scan_age, at = list(scan_age = c(34.785, 41))))
```

Contrast:

```{r, echo=F,message=F,warning=F}
WMage34vs41 %>% contrast("revpairwise") %>% summary(by = NULL, adjust = "holm")
```

Slopes:

```{r, echo=F,message=F,warning=F}
(WMpretermtrendTEA <- emtrends(h.model.int.fullnoTHC_WM, ~ birth_age, var = "scan_age"))
```


### Combined RSN H comparison Preterm vs TEA

Estimated marginal Means

```{r, echo=F,message=F,warning=F}
h.model.int.fullnoTHC_allRSN <- lmer(data = RemovedTHC_fulldata_long, H ~ scan_age*birth_age + (1|Subject), REML = F)
(All_RSNage34vs41 <- h.model.int.fullnoTHC_allRSN %>% emmeans(~scan_age, at = list(scan_age = c(34.785, 41))))
```

Contrast

```{r, echo=F,message=F,warning=F}
All_RSNage34vs41 %>% contrast("revpairwise") %>% summary(by = NULL, adjust = "holm")
```

Slopes:

```{r, echo=F,message=F,warning=F}
(AllRSNpretermtrendTEA <- emtrends(h.model.int.fullnoTHC_allRSN, ~ birth_age, var = "scan_age"))
```


## Individual RSN H comparison Preterm vs TEA

Estimated marginal means

```{r, echo=F,message=F,warning=F}
h.model.int.fullnoTHC_RSN <- lmer(data = RemovedTHC_fulldata_long, H ~ scan_age*RSN*birth_age + (1|Subject), REML = F)
knitr::kable(RSNage34vs41 <- h.model.int.fullnoTHC_RSN  %>% emmeans(~scan_age*RSN, at = list(scan_age = c(34.785, 41))))
```

Contrast:

```{r, echo=F,message=F,warning=F}
knitr::kable(RSNage34vs41 %>% contrast("revpairwise", by="RSN") %>% summary(by = NULL, adjust = "holm"))
```

### RSN Slopes Preterm to TEA
Determining which RSN has the greatest increase with age in the preterm group.

Trends in descending order:

```{r, echo=F,message=F,warning=F}
RSNslopes_pretermtoTEA <- emtrends(h.model.int.fullnoTHC_RSN, ~ RSN, var="scan_age")
knitr::kable(tidy(RSNslopes_pretermtoTEA) %>% arrange(desc(scan_age.trend)))
```

### Investigating when RSNs Surpass Ventricle

Let's treat H in ventricles as a 'noise ground floor'

When do various RSN's surpass the ventricle?
```{r, echo=F,message=F,warning=F}
Preterm_VentandRSN <- RemovedTHC_fulldata[,!names(RemovedTHC_fulldata) %in% c("HinGM", "HinWM")]
Preterm_VentandRSN <- Preterm_VentandRSN %>% pivot_longer(starts_with('H'), names_to = "RSN", values_to = "H")
Preterm_VentandRSN$RSN <- factor(Preterm_VentandRSN$RSN, levels = c("H_MotorMedial", "H_MotorLateral", "H_Visual", "H_Auditory", "H_MotorAssociation", "H_PosteriorParietal", "H_Somatosensory", "H_DorsalVisualStream", "H_PosteriorCingulateCortex","H_FrontalPole", "H_Midbrain", "H_DorsolateralPrefrontal", "H_Cerebellum", "HinVent"))
h.model_simplePretermVent <- lmer(data = Preterm_VentandRSN, H ~ scan_age*RSN*birth_age + (1|Subject), REML = F)
HpredictPretermVent <- ggpredict(h.model_simplePretermVent, terms = c("scan_age","RSN"), ci.lvl=0.95)
VentandRSNslopes <- emtrends(h.model_simplePretermVent, ~ RSN, var="scan_age")
```

Scan age (weeks) when RSN crosses Ventricles:
```{r, echo=F,message=F,warning=F}
HinVent <- HpredictPretermVent %>% dplyr::filter(group == "HinVent")
vent_func <- approxfun(HinVent$x, HinVent$predicted)
brain_regions <- unique(HpredictPretermVent$group)
brain_regions <- brain_regions[-14] ##remove HinVent from brain region list

intersections <- list()

for (i in 1:length(brain_regions)) {
  region <- brain_regions[i]
  H_region <- HpredictPretermVent %>% dplyr::filter(group == region)
  region_func <- approxfun(H_region$x, H_region$predicted)
  intersect <- optimize(function(x) abs(region_func(x) - vent_func(x)), interval = range(H_region$x))
  intersections[i] <- intersect$minimum
}

Ventricle_Crossing <- as.data.frame(brain_regions)
Ventricle_Crossing$intersect <- t(as.data.frame(intersections))
#Ventricle_Crossing[13, 2] <- NULL

knitr::kable(Ventricle_Crossing %>% arrange(intersect))
```


## VPT vs MPT Preterm Group Comparison 
Assessing the very preterm and moderately born preterm groups separately to determine if H values between groups are significantly different at preterm age and term age. 
The overall slopes will be compared in the regions. Only testing at preterm age since we will be doing the term age comparison later with the THC.

### Grey matter

Estimated marginal means:

```{r, echo=F,message=F,warning=F}
knitr::kable(GMage34wkPreGroup <- h.model.int.fullnoTHC_GM %>% emmeans(~scan_age*birth_age, at = list(scan_age = c(35), birth_age = c(29, 35))))
```

Contrast:

```{r, echo=F,message=F,warning=F}
knitr::kable(GMage34wkPreGroup %>% contrast("revpairwise") %>% summary(by = NULL, adjust = "holm"))
```


Slopes:
```{r, echo=F,message=F,warning=F}
GMpretermtrend <- emtrends(h.model.int.fullnoTHC_GM, ~ birth_age, at = list(birth_age = c(29,35)), var = "scan_age")
knitr::kable(GMpretermtrendtidy <- tidy(GMpretermtrend, conf.int = T) %>% dplyr::select(birth_age, scan_age.trend, p.value)) #conf.low, conf.high))
```
Contrast of slopes
```{r, echo=F,message=F,warning=F}
knitr::kable(GMpretermtrend %>% contrast("revpairwise") %>% summary(by = NULL, adjust = "holm"))
```

### White Matter

Estimated marginal means:

```{r, echo=F,message=F,warning=F}
h.model.int.fullnoTHC_WMPreGroup <- lmer(data = RemovedTHC_fulldata, HinWM ~ scan_age*birth_age + (1|Subject), REML = F)
knitr::kable(WMage34wkPreGroup <- h.model.int.fullnoTHC_WMPreGroup %>% emmeans(~scan_age*birth_age, at = list(scan_age = c(35), birth_age = c(29,35))))
```

Contrast:
```{r, echo=F,message=F,warning=F}
knitr::kable(WMage34wkPreGroup %>% contrast("revpairwise") %>% summary(by = NULL, adjust = "holm"))
```

Slopes / trends:

```{r, echo=F,message=F,warning=F}
# Compare slopes in GM in the VPT and MPT
WMpretermtrend <- emtrends(h.model.int.fullnoTHC_WMPreGroup, at = list(birth_age = c(29,35)), var = "scan_age")

knitr::kable(WMpretermtrendtidy <- tidy(WMpretermtrend, conf.int = T) %>% dplyr::select(birth_age, scan_age.trend, p.value)) #conf.low, conf.high))
```

Contrast of trends:

```{r, echo=F,message=F,warning=F}
knitr::kable(WMpretermtrend  %>% contrast("revpairwise") %>% summary(by = NULL, adjust = "holm"))
```

### Combined RSN

Estimated marginal means:

```{r, echo=F,message=F,warning=F}
h.model.int.fullnoTHC_allRSNPreGroup <- lmer(data = RemovedTHC_fulldata_long, H ~ scan_age*birth_age + (1|Subject), REML = F)
knitr::kable(allRSNage34wkPreGroup <- h.model.int.fullnoTHC_allRSNPreGroup %>% emmeans(~scan_age*birth_age, at = list(scan_age = c(35), birth_age = c(29,35))))
```

Contrast:

```{r, echo=F,message=F,warning=F}
knitr::kable(allRSNage34wkPreGroup %>% contrast("revpairwise") %>% summary(by = NULL, adjust = "holm"))
```

Slopes

```{r, echo=F,message=F,warning=F}
allRSNpretermtrend <- emtrends(h.model.int.fullnoTHC_allRSNPreGroup, at = list(birth_age = c(29,35)), var = "scan_age")
knitr::kable(allRSNpretermtrendtidy <- tidy(allRSNpretermtrend, conf.int = T) %>% dplyr::select(birth_age, scan_age.trend, p.value)) #conf.low, conf.high))
```

Contrast of slopes:

```{r, echo=F,message=F,warning=F}
knitr::kable(allRSNpretermtrend %>% contrast("revpairwise") %>% summary(by = NULL, adjust = "holm"))
```


### Individual RSNs

Emmeans:

```{r, echo=F,message=F,warning=F}
h.model.int.fullnoTHC_RSN <- lmer(data = RemovedTHC_fulldata_long, H ~ scan_age*birth_age*RSN + (1|Subject), REML = F)
RSNage34wkPreGroup <- h.model.int.fullnoTHC_RSN  %>% emmeans(~scan_age*birth_age*RSN, at = list(scan_age = c(35), birth_age = c(29,35)))
knitr::kable(RSNage34wkPreGroup)
```


Contrast

```{r, echo=F, message=F, warning=F}
RSNage34wkPreGroup.contrast <- RSNage34wkPreGroup %>% contrast("revpairwise", by = "RSN") %>% summary(by = NULL, adjust = "holm") %>% select(-c(SE, df, z.ratio))
RSNage34wkPreGroup.contrast %>% kbl() %>% kableExtra::column_spec(which(colnames(RSNage34wkPreGroup.contrast)=="p.value"), bold = ifelse(RSNage34wkPreGroup.contrast$p.value < 0.05,TRUE, FALSE))
```

Slopes:

```{r, echo=F,message=F,warning=F}
RSNslopes_pretermgroups <- tidy(emtrends(h.model.int.fullnoTHC_RSN, ~ birth_age*RSN, at = list(birth_age = c(29,35)), var="scan_age"))
```

```{r, echo=F,message=F,warning=F}
knitr::kable(RSNslopes_pretermgroups <- tidy(emtrends(h.model.int.fullnoTHC_RSN, ~ birth_age*RSN, at = list(birth_age = c(29,35)), var="scan_age")))
```

## Plots:

### GM, WM and Combined RSN Plot

Plot of H in the tissues in the preterm group. The horizontal line H=0.5
```{r, echo=F,message=F,warning=F}
RemovedTHCforplot <- RemovedTHC_fulldata_long %>% rename('HinCombinedRSN' = H)
RemovedTHCforplot$HinVent <- NULL
```

```{r, echo=F,message=F,warning=F}
PretermRegionData <- RemovedTHCforplot %>% pivot_longer(starts_with('Hin'), names_to = 'Region', values_to = 'H_region')
PretermRegionData$Region <- as.factor(PretermRegionData$Region)
PretermRegion.model <- lmer(H_region ~ scan_age*Region*birth_age + (1|Subject), REML=FALSE, data = PretermRegionData)
```

```{r, echo=F,message=F,warning=F}
Regionnames <- c("HinGM" = "GM", "HinWM" = "WM", "HinCombinedRSN" = "Combined RSN")
H_pretermregionpredict <- ggpredict(PretermRegion.model, terms = c("scan_age","Region"), ci.lvl=0.95)
region.colors <- c(HinGM = "#AF58BA", HinWM= "#EC3939", HinCombinedRSN = "#00CD6C")
pretermregionplot <- ggplot(H_pretermregionpredict, aes(x=x, y=predicted, color=group)) +
  theme_minimalism() +
  theme(legend.title=element_blank()) +
  geom_smooth(method="lm", se=F) +
  geom_hline(yintercept=0.5, linetype="dashed") +
  geom_line(aes(x=x, y=conf.high), linetype=2, alpha=0.3) +
  geom_line(aes(x=x, y=conf.low), linetype=2, alpha=0.3) +
  scale_color_manual(values=region.colors, limits=c("HinGM","HinWM","HinCombinedRSN"), labels = c("GM", "WM", "Combined RSN")) +
    labs(x='Scan Age (weeks)', y='Predicted H by Region') + theme(panel.grid.major = element_blank(), panel.grid.minor = element_blank())

pretermregionplot
```

```{r, echo=F, message=F, warning=F}
ggsave("pngfigures/VPTMPTtissueslopes.png", width = 20, height = 10, units = "cm", dpi = 600)
```

### Individual RSN Preterm Group Plot

```{r, echo=F,message=F,warning=F}
rsnnames2 <- c("Motor Medial", "Motor Lateral", "Posterior Parietal", "Visual","Auditory", "Somatosensory", "Motor Association", "Dorsal Visual Stream", "Frontal Pole", "Posterior Cingulate", "Dorsolateral Prefrontal", "Midbrain", "Cerebellum", "Ventricles")
rsnnames3 <- c("H_MotorMedial", "H_MotorLateral", "H_PosteriorParietal", "H_Visual","H_Auditory","H_Somatosensory", "H_MotorAssociation", "H_DorsalVisualStream","H_FrontalPole", "H_PosteriorCingulateCortex", "H_DorsolateralPrefrontal", "H_Midbrain", "H_Cerebellum", "HinVent")
rsn.colours <- c(H_MotorMedial = "#FB3434", H_MotorLateral = "#EEA19B", H_Visual = "#F79F5E", H_Auditory = "#6CD30B", H_MotorAssociation = "#B7EE82", H_PosteriorParietal = "#E3670B", H_Somatosensory = "#F4E315", H_DorsalVisualStream = "#14B4E8", H_FrontalPole = "#96D6EA",H_PosteriorCingulateCortex  = "#701BD6", H_DorsolateralPrefrontal = "#BB95E9", H_Midbrain= "#E814EB", H_Cerebellum = "#E79CE8", HinVent = "#000000")
HpredictPretermVent$group <- factor(HpredictPretermVent$group, levels = c("H_MotorMedial", "H_MotorLateral", "H_PosteriorParietal", "H_Visual","H_Auditory","H_Somatosensory", "H_MotorAssociation", "H_DorsalVisualStream","H_FrontalPole", "H_PosteriorCingulateCortex", "H_DorsolateralPrefrontal", "H_Midbrain", "H_Cerebellum", "HinVent"))
```

```{r, echo=F,message=F,warning=F}
ggplot(HpredictPretermVent, aes(x=x, y=predicted, color=group)) +
  geom_line()+ 
  theme_minimalism() +
  geom_line(aes(x=x, y=conf.high), linetype=2, alpha=0.3) +
  geom_line(aes(x=x, y=conf.low), linetype=2, alpha=0.3) +
  geom_hline(yintercept=0.5, linetype="dashed") +
  scale_color_manual(values=rsn.colours,limits = rsnnames3, labels = rsnnames2, name="RSN") +
    labs(x='Scan Age (weeks)', y='Predicted H by RSN') + theme_minimalism()
```

```{r, echo=F,message=F,warning=F}
ggsave("pngfigures/VPTMPTRSNlopes.png", width = 20, height = 10, units = "cm", dpi = 600)
```

### VPT and MPT RSN Barchart at 35 weeks
Comparing the preterm groups VPT and MPT at preterm age. Later will compare these groups at term to THC. 
```{r, echo=F,message=F,warning=F}
RSNage34wkPreGroup.df <- as.data.frame(RSNage34wkPreGroup)
```

```{r, echo=F,message=F,warning=F}
pos <- position_dodge(width=0.5)
emmeanplot1 <- RSNage34wkPreGroup.df  %>% mutate(RSN = fct_relevel(RSN,"H_MotorMedial", "H_MotorLateral", "H_PosteriorParietal", "H_Auditory", "H_Somatosensory", "H_Visual", "H_MotorAssociation", "H_DorsalVisualStream", "H_FrontalPole", "H_DorsolateralPrefrontal", "H_PosteriorCingulateCortex", "H_Midbrain", "H_Cerebellum")) %>%
ggplot(aes(x=RSN, y=emmean, fill= as.factor(birth_age))) + geom_col(position="dodge2") +
  geom_errorbar(aes(ymin=asymp.LCL, ymax=asymp.UCL), width =.3, position=position_dodge(0.9), alpha=0.5) +
  geom_hline(yintercept=0.5, linetype="dashed") +
  coord_cartesian(ylim = c(.2, .8)) + ylab("Estimated Mean H at 35 Weeks PMA") +
  scale_fill_manual(values = c("indianred1", "goldenrod1"), name = "Group", labels = c("VPT (29 weeks GA)", "MPT (35 weeks GA)")) + scale_color_manual(values = c("indianred1", "goldenrod1"), name = "Group", labels = c("VPT", "MPT")) + 
  xlab("RSN") +
  scale_x_discrete(limit = c("H_MotorMedial", "H_MotorLateral", "H_PosteriorParietal", "H_Auditory", "H_Somatosensory", "H_Visual", "H_MotorAssociation", "H_DorsalVisualStream", "H_FrontalPole", "H_DorsolateralPrefrontal", "H_PosteriorCingulateCortex", "H_Midbrain", "H_Cerebellum"),
                     labels = c("Motor Medial", "Motor Lateral", "Posterior Parietal", "Auditory", "Somatosensory", "Visual", "Motor Association", "Dorsal Visual Stream", "Frontal Pole", "Dorsolateral Prefrontal", "Posterior Cingulate", "Midbrain", "Cerebellum")) +
  theme(axis.text.x = element_text(angle = 45, hjust=1),panel.grid.major = element_blank(), panel.grid.minor = element_blank(),
panel.background = element_blank(), axis.line = element_line(colour = "black"))
emmeanplot1
```

```{r, echo=F,message=F,warning=F}
ggsave("pngfigures/VPTMPTRSNbar.png", width = 20, height = 10, units = "cm", dpi = 600)
```

### VPT and MPT Tissue Barchart at 35 weeks
Visual assessment of H values in the tissues compared in the groups. Need to add which are significant differences.
```{r, echo=F,message=F,warning=F}
GMage34wkPreGroup.df <- as.data.frame(GMage34wkPreGroup)
WMage34wkPreGroup.df <- as.data.frame(WMage34wkPreGroup)
allRSNage34wkPreGroup.df <- as.data.frame(allRSNage34wkPreGroup)
Tissue = c("Gray Matter")
GMage34wkPreGroup.df$Tissue <- Tissue
Tissue = c("White Matter")
WMage34wkPreGroup.df$Tissue <- Tissue
Tissue = c("Combined RSN")
allRSNage34wkPreGroup.df$Tissue <- Tissue
allRSNage34wkPreGroup.df <- allRSNage34wkPreGroup.df %>% rename(lower.CL = asymp.LCL, upper.CL = asymp.UCL)
# combine data frames
TissuesPreGroups.df <- rbind(GMage34wkPreGroup.df, WMage34wkPreGroup.df, allRSNage34wkPreGroup.df)
```

```{r, echo=F,message=F,warning=F}
pos <- position_dodge(width=0.5)
emmeanplot <- TissuesPreGroups.df %>%
ggplot(aes(x=Tissue, y=emmean, fill= as.factor(birth_age)  )) + geom_col(position="dodge2") +
  geom_errorbar(aes(ymin=lower.CL, ymax=upper.CL), width =.3, position=position_dodge(0.9), alpha=0.5) +
  geom_hline(yintercept=0.5, linetype="dashed") +
  coord_cartesian(ylim = c(.2, .7)) + ylab("Estimated Mean H at 35 Weeks PMA") +
  scale_fill_manual(values = c("indianred1", "goldenrod1"), name = "Group", labels = c("VPT (29 weeks GA)", "MPT (35 weeks GA)")) + scale_color_manual(values = c("indianred1", "goldenrod1"), name = "Group", labels = c("VPT", "MPT")) + 
  xlab("Tissue") +
  scale_x_discrete(limit = c("Gray Matter", "White Matter", "Combined RSN"),
                     labels = c("Gray Matter", "White Matter", "Combined RSN")) +
  theme(axis.text.x = element_text(angle = 45, hjust=1),panel.grid.major = element_blank(), panel.grid.minor = element_blank(),
panel.background = element_blank(), axis.line = element_line(colour = "black"))
emmeanplot
```

```{r, echo=F, message=F, warning=F}
ggsave("pngfigures/VPTMPTtissuebar.png", width = 20, height = 10, units = "cm", dpi = 600)
```

---

# All Group Comparisons:

Do term born infants have greater H compared to preterm infants at scan age?

VPT at TEA vs MPT at TEA vs THC

Assessing the very preterm and moderately born preterm groups at term age to the term born controls. 

Removing all the preterm scans.
```{r, echo=F, message=F, warning=F}
RemovedpretermGAfactors <- fulldata_long%>% filter(scan_age > 37)
medianteaage <- RemovedpretermGAfactors %>% filter(birth_age > 37) %>% summarise(Median.Age = median(birth_age))
```

Median age when looking at all scans > 37 weeks: `r medianteaage`

## GM H Comparison VPT vs MPT vs THC

```{r, echo=F,message=F,warning=F}
h.model.int.fullTEAgroup_GM <- lmer(data = RemovedpretermGAfactors, HinGM ~ scan_age*birth_age + (1|Subject), REML = F)
GMlm_TEAgroupvsTHC_41 <- h.model.int.fullTEAgroup_GM %>% emmeans(~scan_age*birth_age, at = list(scan_age = c(41), birth_age = c(29,35,41))) 
knitr::kable(GMlm_TEAgroupvsTHC_41)
```

Contrast:

```{r, echo=F, message=F, warning=F}
knitr::kable(GMlm_TEAgroupvsTHC_41 %>% contrast("revpairwise") %>% summary(by = NULL, adjust = "holm"))
```

## WM H Comparison VPT vs MPT vs THC
```{r, echo=F,message=F,warning=F}
h.model.int.fullTEAgroup_WM <- lmer(data = RemovedpretermGAfactors, HinWM ~ scan_age*birth_age + (1|Subject), REML = F)
WMlm_TEAgroupvsTHC_41 <- h.model.int.fullTEAgroup_WM %>% emmeans(~scan_age*birth_age, at = list(scan_age = c(41), birth_age = c(29,35,41) )) 
knitr::kable(WMlm_TEAgroupvsTHC_41)
```


```{r, echo=F, message=F, warning=F}
knitr::kable(WMlm_TEAgroupvsTHC_41 %>% contrast("revpairwise") %>% summary(by = NULL, adjust = "holm"))
```

## Combined RSN H Comparison VPT vs MPT vs THC
```{r, echo=F,message=F,warning=F}
h.model.int.fullTEAgroup_allRSN <- lmer(data = RemovedpretermGAfactors, H ~ scan_age*birth_age + (1|Subject), REML = F)
allRSNlm_TEAgroupvsTHC_41 <- h.model.int.fullTEAgroup_allRSN %>% emmeans(~scan_age*birth_age, at = list(scan_age = c(41), birth_age=c(29,35,41) )) 
knitr::kable(allRSNlm_TEAgroupvsTHC_41)
```

```{r, echo=F,message=F,warning=F}
knitr::kable(allRSNlm_TEAgroupvsTHC_41 %>% contrast("revpairwise") %>% summary(by = NULL, adjust = "holm"))
```

## Individual RSN H Comparison VPT vs MPT vs THC
```{r, echo=F,message=F,warning=F}
h.model.int.fullTEAgroup_RSN <- lmer(data = RemovedpretermGAfactors, H ~ scan_age*birth_age*RSN + (1|Subject), REML = F)
RSNageTEAGroupvsTHC_41 <- h.model.int.fullTEAgroup_RSN  %>% emmeans(~scan_age*birth_age*RSN, at = list(scan_age = c(41), birth_age = c(29,35,41) ))
knitr::kable(RSNageTEAGroupvsTHC_41)
```

Contrast

```{r, echo=F,message=F,warning=F}
RSNageTEAGroupvsTHC_41.contrast <- RSNageTEAGroupvsTHC_41 %>% contrast("revpairwise", by = "RSN") %>% summary(by = NULL, adjust = "holm") %>% select(-c(SE, df, z.ratio))
RSNageTEAGroupvsTHC_41.contrast %>% kbl() %>% kableExtra::column_spec(which(colnames(RSNageTEAGroupvsTHC_41.contrast)=="p.value"), bold = ifelse(RSNageTEAGroupvsTHC_41.contrast$p.value < 0.05,TRUE, FALSE))
```

Slopes:

```{r, echo=F, message=F, warning=F}
RSNslopes_Termgroups <- emtrends(h.model.int.fullTEAgroup_RSN, ~ birth_age*RSN, at = list(birth_age = c(29,35,41)), var="scan_age")
knitr::kable(RSNslopes_Termgroups)
```

## Preterm Groups vs THC Plots

### Group RSN Bar Graph
Visual assessment of H values in the RSN compared in the groups. 
```{r, echo=F,message=F,warning=F}
RSNageTEAGroupvsTHC_41.df <- as.data.frame(RSNageTEAGroupvsTHC_41)
```

```{r, echo=F,message=F,warning=F}
pos <- position_dodge(width=0.5)
emmeanplot <- RSNageTEAGroupvsTHC_41.df  %>% mutate(RSN = fct_relevel(RSN,"H_MotorMedial", "H_MotorLateral", "H_PosteriorParietal", "H_Auditory", "H_Somatosensory", "H_Visual", "H_MotorAssociation", "H_DorsalVisualStream", "H_FrontalPole", "H_DorsolateralPrefrontal", "H_PosteriorCingulateCortex", "H_Midbrain", "H_Cerebellum")) %>%
ggplot(aes(x=RSN, y=emmean, fill= as.factor(birth_age) )) + geom_col(position="dodge2") +
  geom_errorbar(aes(ymin=asymp.LCL, ymax=asymp.UCL), width =.3, position=position_dodge(0.9), alpha=0.5) +
  geom_hline(yintercept=0.5, linetype="dashed") +
  coord_cartesian(ylim = c(.2, .8)) + ylab("Estimated Mean H at 41 Weeks PMA") +
  scale_fill_manual(values = c("indianred1", "goldenrod1", "deepskyblue"), name = "Group", labels = c("VPT (29 weeks GA)", "MPT (35 weeks GA)", "THC (41 weeks GA)")) + scale_color_manual(values = c("indianred1", "goldenrod1", "deepskyblue"), name = "Group", labels = c("VPT", "MPT", "THC")) + 
  xlab("RSN") +
  scale_x_discrete(limit = c("H_MotorMedial", "H_MotorLateral", "H_PosteriorParietal", "H_Auditory", "H_Somatosensory", "H_Visual", "H_MotorAssociation", "H_DorsalVisualStream", "H_FrontalPole", "H_DorsolateralPrefrontal", "H_PosteriorCingulateCortex", "H_Midbrain", "H_Cerebellum"),
                     labels = c("Motor Medial", "Motor Lateral", "Posterior Parietal", "Auditory", "Somatosensory", "Visual", "Motor Association", "Dorsal Visual Stream", "Frontal Pole", "Dorsolateral Prefrontal", "Posterior Cingulate", "Midbrain", "Cerebellum")) +
  theme(axis.text.x = element_text(angle = 45, hjust=1),panel.grid.major = element_blank(), panel.grid.minor = element_blank(),
panel.background = element_blank(), axis.line = element_line(colour = "black"))
emmeanplot
```
```{r, echo=F, message=F, warning=F}
ggsave("pngfigures/ALLRSNbar.png", width = 20, height = 10, units = "cm", dpi = 600)
```


### Tissues Bar Graph
Visual assessment of H values in the tissues compared in the groups. 

```{r, echo=F,message=F,warning=F}
GMTEAGroupvsTHC_41.df <- as.data.frame(GMlm_TEAgroupvsTHC_41)
WMTEAGroupvsTHC_41.df <- as.data.frame(WMlm_TEAgroupvsTHC_41)
allRSNGroupvsTHC_41.df <- as.data.frame(allRSNlm_TEAgroupvsTHC_41)
Tissue = c("Gray Matter")
GMTEAGroupvsTHC_41.df$Tissue <- Tissue
Tissue = c("White Matter")
WMTEAGroupvsTHC_41.df$Tissue <- Tissue
Tissue = c("Combined RSN")
allRSNGroupvsTHC_41.df$Tissue <- Tissue
# combine data frames
TissuesGroups.df <- rbind(GMTEAGroupvsTHC_41.df, WMTEAGroupvsTHC_41.df, allRSNGroupvsTHC_41.df)
```

```{r, echo=F,message=F,warning=F}
pos <- position_dodge(width=0.5)
emmeanplot <- TissuesGroups.df %>%
ggplot(aes(x=Tissue, y=emmean, fill= as.factor(birth_age))) + geom_col(position="dodge2") +
  geom_errorbar(aes(ymin=asymp.LCL, ymax=asymp.UCL), width =.3, position=position_dodge(0.9), alpha=0.5) +
  geom_hline(yintercept=0.5, linetype="dashed") +
  coord_cartesian(ylim = c(.2, .7)) + ylab("Estimated Mean H at 41 Weeks PMA") +
  scale_fill_manual(values = c("indianred1", "goldenrod1", "deepskyblue"), name = "Group", labels = c("VPT (29 weeks GA)", "MPT (35 weeks GA)", "THC (41 weeks GA)")) + 
  scale_color_manual(values = c("indianred1", "goldenrod1", "deepskyblue"), name = "Group", labels = c("VPT", "MPT", "THC")) + 
  xlab("Tissue") +
  scale_x_discrete(limit = c("Gray Matter", "White Matter", "Combined RSN"),
                     labels = c("Gray Matter", "White Matter", "Combined RSN")) +
  theme(axis.text.x = element_text(angle = 45, hjust=1),panel.grid.major = element_blank(), panel.grid.minor = element_blank(),
panel.background = element_blank(), axis.line = element_line(colour = "black"))
emmeanplot
```

```{r, echo=F, message=F, warning=F}
ggsave("pngfigures/ALLtissuebar.png", width = 20, height = 10, units = "cm", dpi = 600)
```

---

# Is H associated with indirect measures of myelination (FA and RD)?

WM measures of FA and RD were calculated in the total white matter and in the RSNs.

Load data (hidden)
```{r, echo=F, message=F, warning=F}
myelindata1 <- read.csv("csvfiles/ODandFA_tractandWM_values.csv") %>% select(Subject, Session, ends_with("_WMFA"))
myelindata2 <- read.csv("csvfiles/RD_notract_values.csv")
myelindata <- merge(myelindata1, myelindata2)
# Merging WM measures to H data.
# Making new data frame with new H RSNs names

lookup <- c("MedialMotor_H" = "H_MotorMedial", "LateralMotor_H" = "H_MotorLateral", "Posterior_Parietal_H" = "H_PosteriorParietal", "Visual_H" = "H_Visual","Auditory_H" = "H_Auditory","Somatosensory_H" = "H_Somatosensory", "MotorAssociation_H" = "H_MotorAssociation", "Dorsal_Visual_Stream_H" = "H_DorsalVisualStream","Frontal_Pole_H" = "H_FrontalPole", "Posterior_Cingular_Cortex_H" = "H_PosteriorCingulateCortex", "Dorsolateral_Prefrontal_H" = "H_DorsolateralPrefrontal", "Hindbrain_H" = "H_Midbrain", "Cerebellum_H" = "H_Cerebellum")
fulldata_HandWM <- RemovedTHC_fulldata %>% rename(all_of(lookup))

MyelinandHdata <- merge(myelindata, fulldata_HandWM) %>% select(-X,-scan_number,-singleton,-sedation,-birth_weight,-head_circumference_scan,-radiology_score)

RSN_namesWMFA <- list(
  'MedialMotor' = "Motor Medial",
  'LateralMotor' = "Motor Lateral",
  'Visual' = "Visual",
  'Auditory'= "Auditory",
  'MotorAssociation' = "Motor Association",
  'Posterior_Parietal' = "Posterior Parietal",
  'Somatosensory' = "Somatosensory",
  'Dorsal_Visual_Stream' = "Dorsal Visual Stream",
  'Posterior_Cingular_Cortex' = "Posterior Cingulate Cortex",
  'Frontal_Pole'= "Frontal Pole",
  'Dorsolateral_Prefrontal'= "Dorsolateral Prefrontal"
)

RSN_labellerWMFA <- function(variable,value){
  return(RSN_namesWMFA[value])
}

MyelinandHdata_long_ <- MyelinandHdata %>% select(Subject,Session,ends_with("_WMFA")) %>% pivot_longer(cols = ends_with("_WMFA"), names_to = "RSN", values_to = "WMFA") %>% mutate(across('RSN', str_replace, '_WMFA', '')) %>% filter(RSN != "Hindbrain") %>% filter(RSN != "Cerebellum")

MyelinandHdata_long__ <- MyelinandHdata %>% select(Subject,Session,ends_with("_H"),GAfactor,scan_age, birth_age) %>% pivot_longer(cols = ends_with("_H"), names_to = "RSN", values_to = "H") %>% mutate(across('RSN', str_replace, '_H', '')) %>% filter(RSN != "Hindbrain") %>% filter(RSN != "Cerebellum")

MyelinandHdata_long___ <- MyelinandHdata %>% select(Subject,Session,ends_with("_RD")) %>% pivot_longer(cols = ends_with("_RD"), names_to = "RSN", values_to = "RD") %>% mutate(across('RSN', str_replace, '_RD', '')) %>% filter(RSN != "Hindbrain") %>% filter(RSN != "Cerebellum")

MyelinandHdata_long <- merge(MyelinandHdata_long_,MyelinandHdata_long__) %>% filter(WMFA > 0) %>% filter(!is.na(WMFA))

MyelinandHdata_longRD <- merge(MyelinandHdata_long, MyelinandHdata_long___)

rm(MyelinandHdata_long_,MyelinandHdata_long__, MyelinandHdata_long___)
```

## FA Analysis

### FA comparison Total WM Preterm vs TEA
```{r, echo=F, message=F, warning=F}
FA.model.fullnoTHC_WM <-lmer(data=MyelinandHdata_long, WMFA ~ scan_age*birth_age + (1|Subject), REML = F)
FA_WMage34vs41 <- FA.model.fullnoTHC_WM  %>% emmeans(~scan_age*birth_age, at = list(scan_age = c(41), birth_age = c(35,41)))
knitr::kable(FA_WMage34vs41)
```

Contrast:
```{r, echo=F, message=F, warning=F}
FA_WMage34vs41.contrast <- FA_WMage34vs41 %>% contrast("revpairwise") %>% summary(by = NULL, adjust = "holm") %>% select(-c(SE, df, t.ratio))
FA_WMage34vs41.contrast %>% kbl() %>% kableExtra::column_spec(which(colnames(FA_WMage34vs41.contrast)=="p.value"), bold = ifelse(FA_WMage34vs41.contrast$p.value < 0.05,TRUE, FALSE))
```

### FA vs scan age (PMA) plot 
```{r, echo=F, message=F, warning=F}
(FAvsSCplot <- ggplot(data = MyelinandHdata_long, aes(x = scan_age, y = WMFA)) +
  geom_point(shape=1, alpha = 1/10) + geom_smooth(method = "lm", se=T) + 
  theme_minimalism() +
  xlab("Scan Age/PMA (weeks)") +
  ylab("FA (Total WM)") +
  theme(legend.title=element_blank()))
```

### FA comparison individual RSN Preterm vs TEA
```{r, echo=F, message=F, warning=F}
FA.model.fullnoTHC_RSN <- MyelinandHdata_long %>% select(-H) %>% lmer(data=., WMFA ~ scan_age*RSN + (1|Subject), REML = F)
FA_RSNage34vs41 <- FA.model.fullnoTHC_RSN  %>% emmeans(~scan_age*RSN, at = list(scan_age = c(35, 41)))
knitr::kable(FA_RSNage34vs41)
```

Contrast:

```{r, echo=F, message=F, warning=F}
FA_RSNage34vs41.contrast <- FA_RSNage34vs41 %>% contrast("revpairwise") %>% summary(by = NULL, adjust = "holm") %>% select(-c(SE, df, t.ratio))
FA_RSNage34vs41.contrast %>% kbl() %>% kableExtra::column_spec(which(colnames(FA_RSNage34vs41.contrast)=="p.value"), bold = ifelse(FA_RSNage34vs41.contrast$p.value < 0.05,TRUE, FALSE))
```

### FA Slopes in RSN Preterm vs TEA
```{r, echo=F, message=F, warning=F}
FA_RSNslopes_pretermtoTEA <- emtrends(FA.model.fullnoTHC_RSN, ~ RSN, var="scan_age")
knitr::kable(tidy(FA_RSNslopes_pretermtoTEA) %>% arrange(desc(scan_age.trend)))
```

### FA plot in the RSN
```{r, echo=F, message=F, warning=F}
FApredictPreterm <- ggpredict(FA.model.fullnoTHC_RSN, terms = c("scan_age","RSN"), ci.lvl=0.95)
```

```{r, echo=F, message=F, warning=F}
rsnnames2 <- c("Motor Medial", "Motor Lateral", "Posterior Parietal", "Visual","Auditory", "Somatosensory", "Motor Association", "Dorsal Visual Stream", "Frontal Pole", "Posterior Cingulate", "Dorsolateral Prefrontal")
rsnnames3 <- c("MedialMotor", "LateralMotor", "Posterior_Parietal", "Visual","Auditory","Somatosensory", "MotorAssociation", "Dorsal_Visual_Stream","Frontal_Pole", "Posterior_Cingular_Cortex", "Dorsolateral_Prefrontal")
rsn.colours <- c(MedialMotor = "#FB3434", LateralMotor = "#EEA19B", Visual = "#F79F5E", Auditory = "#6CD30B", MotorAssociation = "#B7EE82", Posterior_Parietal = "#E3670B", Somatosensory = "#F4E315", Dorsal_Visual_Stream = "#14B4E8", Frontal_Pole = "#96D6EA",Posterior_Cingular_Cortex  = "#701BD6", Dorsolateral_Prefrontal = "#BB95E9")
FApredictPreterm$group <- factor(FApredictPreterm$group, levels = c("MedialMotor", "LateralMotor", "Posterior_Parietal", "Visual","Auditory","Somatosensory", "MotorAssociation", "Dorsal_Visual_Stream","Frontal_Pole", "Posterior_Cingular_Cortex", "Dorsolateral_Prefrontal"))
```

```{r, echo=F, message=F, warning=F}
ggplot(FApredictPreterm, aes(x=x, y=predicted, color=group)) +
  geom_line()+ 
  theme_minimalism() +
  geom_line(aes(x=x, y=conf.high), linetype=2, alpha=0.3) +
  geom_line(aes(x=x, y=conf.low), linetype=2, alpha=0.3) +
  
  scale_color_manual(values=rsn.colours,limits = rsnnames3, labels = rsnnames2, name="ROI") +
    labs(x='Scan Age (weeks)', y='Predicted FA by ROI') + theme_minimalism()
```

### FA and H Correlation test 

**Total WM**:

```{r, echo=F, message=F, warning=F}
knitr::kable(MyelinandHdata_long  %>% do(tidy(cor.test(.$H, .$WMFA))))
```

**White Matter Regions**:

```{r, echo=F, message=F, warning=F}
knitr::kable(MyelinandHdata_long %>% group_by(RSN)  %>% do(tidy(cor.test(.$H, .$WMFA)))  %>% arrange(desc(estimate)))
```

**FA H correlation preterm vs TEA**:

Preterm:

```{r, echo=F, message=F, warning=F}
knitr::kable(MyelinandHdata_long  %>% filter(scan_age < 37) %>% do(tidy(cor.test(.$H,.$WMFA))))
```

Term:

```{r, echo=F, message=F, warning=F}
knitr::kable(MyelinandHdata_long  %>% filter(scan_age > 37) %>% do(tidy(cor.test(.$H,.$WMFA))))
```


### FA and H Correlation Plot
```{r, echo=F, message=F, warning=F}
MyelinandHdata_long %>% ggplot(aes(x=H,y=WMFA,color=scan_age)) + geom_point(shape = 21, fill = "white") + 
  scale_colour_gradient(
  name = "Scan Age",
  low = "#F43030",
  high = "#2644EC",
  space = "Lab",
  na.value = "grey50",
  guide = "colourbar",
  aesthetics = "colour") +
  geom_smooth(method="lm") + 
  theme_minimalism() + 
  xlab("H") + ylab("FA")
```

```{r, echo=F, message=F, warning=F}
ggsave("pngfigures/FAvsH.png", width = 22, height = 10, units = "cm", dpi = 600)
```

### FA and H Correlation RSN Plot
```{r, echo=F, message=F, warning=F}
MyelinandHdata_long %>% group_by(RSN)  %>% ggplot(aes(x=H,y=WMFA,color=scan_age)) + geom_point(shape = 21, fill = "white") +
   scale_colour_gradient(
  name = "Scan Age",
  low = "#F43030",
  high = "#2644EC",
  space = "Lab",
  na.value = "grey50",
  guide = "colourbar",
  aesthetics = "colour") +
  geom_smooth(method="lm") + facet_wrap(~RSN, labeller=RSN_labellerWMFA) + theme_minimalism() + xlab("H") + ylab("FA") 
```

```{r, echo=F, message=F, warning=F}
ggsave("pngfigures/FAvsH_RSN.png", width = 22, height = 10, units = "cm", dpi = 600)
```

## RD Analysis

### RD comparison Total WM Preterm vs TEA:

```{r, echo=F, message=F, warning=F}
RD.model.fullnoTHC_WM <-lmer(data=MyelinandHdata_longRD, RD ~ scan_age*birth_age + (1|Subject), REML = F)
knitr::kable(RD_WMage35vs41 <- RD.model.fullnoTHC_WM  %>% emmeans(~scan_age*birth_age, at = list(scan_age = c(41), birth_age = c(35,41))))
```

Contrast:

```{r, echo=F, message=F, warning=F}
RD_WMage35vs41.contrast <- RD_WMage35vs41 %>% contrast("revpairwise") %>% summary(by = NULL, adjust = "holm") %>% select(-c(SE, df, t.ratio))
RD_WMage35vs41.contrast %>% kbl() %>% kableExtra::column_spec(which(colnames(RD_WMage35vs41.contrast)=="p.value"), bold = ifelse(RD_WMage35vs41.contrast$p.value < 0.05,TRUE, FALSE))
```

### RD vs scan age (PMA) plot 
```{r, echo=F, message=F, warning=F}
(RDvsSCplot <- ggplot(data = MyelinandHdata_longRD, aes(x = scan_age, y = RD)) +
  geom_point(shape=1, alpha = 1/10) + geom_smooth(method = "lm", se=T) + 
  theme_minimalism() +
  xlab("Scan Age/PMA (weeks)") +
  ylab("RD (Total WM)") +
   #coord_cartesian(ylim = c(.0005, .0025)) +
  theme(legend.title=element_blank()))
```

### RD comparison individual RSN Preterm vs TEA
```{r, echo=F, message=F, warning=F}
RD.model.fullnoTHC_RSN <- MyelinandHdata_longRD %>% select(-H) %>% lmer(data=., RD ~ scan_age*RSN + (1|Subject), REML = F)
RD_RSNage34vs41 <- RD.model.fullnoTHC_RSN  %>% emmeans(~scan_age*RSN, at = list(scan_age = c(35, 41)))
knitr::kable(RD_RSNage34vs41)

```


```{r, echo=F, message=F, warning=F}
RD_RSNage34vs41.contrast <- RD_RSNage34vs41 %>% contrast("revpairwise", by = "RSN") %>% summary(by = NULL, adjust = "holm") %>% select(-c(SE, df, t.ratio))
RD_RSNage34vs41.contrast %>% kbl() %>% kableExtra::column_spec(which(colnames(RD_RSNage34vs41.contrast)=="p.value"), bold = ifelse(RD_RSNage34vs41.contrast$p.value < 0.05,TRUE, FALSE))

```

### RD Slopes in RSN Preterm vs TEA
```{r, echo=F, message=F, warning=F}
RD_RSNslopes_pretermtoTEA <- emtrends(RD.model.fullnoTHC_RSN, ~ RSN, var="scan_age")
knitr::kable(tidy(RD_RSNslopes_pretermtoTEA) %>% arrange(desc(scan_age.trend)))
```

### RD plot in the RSN
```{r, echo=F, message=F, warning=F}
RDpredictPreterm <- ggpredict(RD.model.fullnoTHC_RSN, terms = c("scan_age","RSN"), ci.lvl=0.95)
```

```{r, echo=F, message=F, warning=F}
rsnnames2 <- c("Motor Medial", "Motor Lateral", "Posterior Parietal", "Visual","Auditory", "Somatosensory", "Motor Association", "Dorsal Visual Stream", "Frontal Pole", "Posterior Cingulate", "Dorsolateral Prefrontal")
rsnnames3 <- c("MedialMotor", "LateralMotor", "Posterior_Parietal", "Visual","Auditory","Somatosensory", "MotorAssociation", "Dorsal_Visual_Stream","Frontal_Pole", "Posterior_Cingular_Cortex", "Dorsolateral_Prefrontal")
rsn.colours <- c(MedialMotor = "#FB3434", LateralMotor = "#EEA19B", Visual = "#F79F5E", Auditory = "#6CD30B", MotorAssociation = "#B7EE82", Posterior_Parietal = "#E3670B", Somatosensory = "#F4E315", Dorsal_Visual_Stream = "#14B4E8", Frontal_Pole = "#96D6EA",Posterior_Cingular_Cortex  = "#701BD6", Dorsolateral_Prefrontal = "#BB95E9")
FApredictPreterm$group <- factor(FApredictPreterm$group, levels = c("MedialMotor", "LateralMotor", "Posterior_Parietal", "Visual","Auditory","Somatosensory", "MotorAssociation", "Dorsal_Visual_Stream","Frontal_Pole", "Posterior_Cingular_Cortex", "Dorsolateral_Prefrontal"))
```

```{r, echo=F, message=F, warning=F}
ggplot(RDpredictPreterm, aes(x=x, y=predicted, color=group)) +
  geom_line()+ 
  theme_minimalism() +
  geom_line(aes(x=x, y=conf.high), linetype=2, alpha=0.3) +
  geom_line(aes(x=x, y=conf.low), linetype=2, alpha=0.3) +
  
  scale_color_manual(values=rsn.colours,limits = rsnnames3, labels = rsnnames2, name="ROI") +
    labs(x='Scan Age (weeks)', y='Predicted RD by ROI') + theme_minimalism()
```

## RD and H Correlation tests

#### Total WM
```{r, echo=F, message=F, warning=F}
knitr::kable(MyelinandHdata_longRD  %>% do(tidy(cor.test(.$H, .$RD))))
```

#### White Matter Regions
```{r, echo=F, message=F, warning=F}
RDH_RSN_precorr <- MyelinandHdata_longRD %>% group_by(RSN)  %>% do(tidy(cor.test(.$H, .$RD)))
RDH_RSN_precorr %>% kbl() %>% kableExtra::column_spec(which(colnames(RDH_RSN_precorr)=="p.value"), bold = ifelse(RDH_RSN_precorr$p.value < 0.05,TRUE, FALSE))
```

```{r, echo=F, message=F, warning=F}
MyelinandHdata_longRD.cortest <- MyelinandHdata_longRD %>% group_by(RSN)  %>% do(tidy(cor.test(.$H, .$RD)))  %>% arrange(desc(estimate)) 

MyelinandHdata_longRD.cortest %>% kbl() %>% kableExtra::column_spec(which(colnames(MyelinandHdata_longRD.cortest)=="p.value"), bold = ifelse(MyelinandHdata_longRD.cortest$p.value < 0.05,TRUE, FALSE))
```

### RD and H Correlation Plot
```{r, echo=F, message=F, warning=F}
MyelinandHdata_longRD %>% ggplot(aes(x=H,y=RD,color=scan_age)) + geom_point(shape = 21, fill = "white") + 
  scale_colour_gradient(
  name = "Scan Age",
  low = "#F43030",
  high = "#2644EC",
  space = "Lab",
  na.value = "grey50",
  guide = "colourbar",
  aesthetics = "colour") +
  geom_smooth(method="lm") + theme_minimalism() + xlab("H") + ylab("RD") +coord_cartesian(ylim = c(.0007, .0014)) 
```

```{r, echo=F, message=F, warning=F}
ggsave("pngfigures/RDvsH.png", width = 22, height = 10, units = "cm", dpi = 600)
```

### RD and H Correlation RSN Plot
```{r, echo=F, message=F, warning=F}
MyelinandHdata_longRD %>% group_by(RSN)  %>% ggplot(aes(x=H,y=RD,color=scan_age)) + geom_point(shape = 21, fill = "white") +
  scale_colour_gradient(
  name = "Scan Age",
  low = "#F43030",
  high = "#2644EC",
  space = "Lab",
  na.value = "grey50",
  guide = "colourbar",
  aesthetics = "colour") +
  geom_smooth(method="lm") + facet_wrap(~RSN, labeller=RSN_labellerWMFA) + theme_minimalism() + xlab("H") + ylab("RD") 
```

```{r, echo=F, message=F, warning=F}
ggsave("pngfigures/RDvsH_RSN.png", width = 22, height = 10, units = "cm", dpi = 600)
```

---

# Supplementary Analysis

## Sex Differences Analysis
Comparing both sexes in total group
Better to look at overall means, sexes may differ at a specific time point.

### Grey Matter sex differences emmeans
```{r, echo=F,message=F,warning=F}
h.model.int.fullGM_sex <- lmer(data = fulldata_long, HinGM ~ sex + (1|Subject), REML = F)
knitr::kable(GMMvsF <- h.model.int.fullGM_sex%>% emmeans(~sex, at = list(sex = c('male', 'female'))))


knitr::kable(GMMvsF %>% contrast("revpairwise") %>% summary(by = NULL, adjust = "holm"))
```

### GM Trend
```{r, echo=F,message=F,warning=F}
GMlm_sex <- lmer(data = fulldata, HinGM ~ scan_age*sex*birth_age + (1|Subject), REML = F)
GMsextrend <- emtrends(GMlm_sex, ~ sex, var = "scan_age")
GMsextrendtidy <- tidy(GMsextrend, conf.int = T) %>% dplyr::select(sex, scan_age.trend, p.value, conf.low, conf.high)

knitr::kable(GMsextrendtidy <- GMsextrend %>% contrast("revpairwise") %>% tidy(.))
```

### White Matter sex differences emmeans
```{r, echo=F,message=F,warning=F}
h.model.int.fullWM_sex <- lmer(data = fulldata_long, HinWM ~ sex + (1|Subject), REML = F)
WMMvsF <- h.model.int.fullWM_sex%>% emmeans(~sex, at = list(sex = c('male', 'female')))
knitr::kable(WMMvsF)

knitr::kable(WMMvsF %>% contrast("revpairwise") %>% summary(by = NULL, adjust = "holm"))
```

### WM Trend
```{r, echo=F,message=F,warning=F}
WMlm_sex <- lmer(data = fulldata, HinWM ~ scan_age*sex*birth_age + (1|Subject), REML = F)
WMsextrend <- emtrends(WMlm_sex, ~ sex, var = "scan_age")
WMsextrendtidy <- tidy(WMsextrend, conf.int = T) %>% dplyr::select(sex, scan_age.trend, p.value, conf.low, conf.high)

WMsextrendtidy <- WMsextrend %>% contrast("revpairwise") %>% tidy(.)
knitr::kable(WMsextrendtidy)
```

### Combined RSN sex differences emmeans
```{r, echo=F,message=F,warning=F}
h.model.int.fullallRSN_sex <- lmer(data = fulldata_long, H ~ sex + (1|Subject), REML = F)
All_RSNMvsF <- h.model.int.fullallRSN_sex%>% emmeans(~sex, at = list(sex = c('male', 'female')))
knitr::kable(All_RSNMvsF)

knitr::kable(All_RSNMvsF %>% contrast("revpairwise") %>% summary(by = NULL, adjust = "holm"))
```

### Combined RSN trend
```{r, echo=F,message=F,warning=F}
AllRSNlm_sex <- lmer(data = fulldata_long, H ~ scan_age*sex*birth_age + (1|Subject), REML = F)
AllRSNsextrend <- emtrends(AllRSNlm_sex, ~ sex, var = "scan_age")
knitr::kable(tidy(AllRSNsextrend, conf.int = T) %>% dplyr::select(sex, scan_age.trend, p.value, conf.low, conf.high))

knitr::kable(AllRSNsextrend %>% contrast("revpairwise") %>% summary(by = NULL, adjust = "holm"))
```

```{r, echo=F, message=F, warning=F}
(Sexdiffplot <- ggplot(data = fulldata_long, aes(x = scan_age, y = H, color = sex)) +
  geom_point(shape=1, alpha = 1/10) + geom_smooth(method = "lm", se=T) + 
   geom_hline(yintercept=0.5, linetype="dashed") +
  stat_smooth(method = "lm", aes(fill=sex), alpha=0.2) +
  theme_minimalism() +
  xlab("Scan Age (weeks)") +
  ylab("H in Combined RSN") +
  theme(legend.title=element_blank()))
```

## Individual RSN Sex Differences
```{r, echo=F,message=F,warning=F}
h.model.int.fullRSN_sex <- lmer(data = fulldata_long, H ~ sex*RSN*birth_age + (1|Subject), REML = F)
RSNMvsF <- h.model.int.fullRSN_sex%>% emmeans(~sex*RSN, at = list(sex = c('male', 'female')))
knitr::kable(RSNMvsF)

knitr::kable(RSNMvsF %>% contrast("revpairwise", by = "RSN") %>% summary(by = NULL, adjust = "holm"))
```

## Standard Deviation BOLD Analysis
Determining how H relates to basic signal variability of BOLD standard deviation. 
In adults, there is a positive correlation between BOLD SD and H. 
Correlation between BOLD SD and H in the whole brain and in the groups.

```{r, echo=F,message=F,warning=F}
BOLD_SD <- read.csv("csvfiles/new_BOLD_SDvalinRSN_wFIX.csv")
```

```{r, echo=F,message=F,warning=F}
#Rename RSN columns 
BOLD_SD <- BOLD_SD %>% rename(
  'SD_MotorMedial' = BOLD_SD_0000,
  'SD_MotorLateral' = BOLD_SD_0001,
  'SD_Visual' = BOLD_SD_0002,
  'SD_Auditory' = BOLD_SD_0003and0006,
  'SD_MotorAssociation' = BOLD_SD_0004,
  'SD_PosteriorParietal' = BOLD_SD_0005,
  'SD_Somatosensory' = BOLD_SD_0007and0011,
  'SD_DorsalVisualStream' = BOLD_SD_0008and0012,
  'SD_PosteriorCingulateCortex' = BOLD_SD_0009, 
  'SD_FrontalPole' = BOLD_SD_0013,
  'SD_Midbrain' = BOLD_SD_0014,
  'SD_DorsolateralPrefrontal' = BOLD_SD_0015, 
  'SD_Cerebellum' = BOLD_SD_0029
  )
```

```{r, echo=F,message=F,warning=F}
BOLD_SD_longT <- BOLD_SD %>% pivot_longer(cols = starts_with("SD_"), names_to = "RSN", values_to = "BOLDSD") %>% mutate(across('RSN', str_replace, 'SD_', ''))
BOLD_SD_longHT <- fulldata_long %>% mutate(across('RSN', str_replace, 'H_', ''))
BOLD_SD_Hdatalong <- merge(BOLD_SD_longT, BOLD_SD_longHT) %>% select(-X,-scan_number,-singleton,-sedation,-birth_weight,-head_circumference_scan,-radiology_score)
rm(BOLD_SD_longT, BOLD_SD_longHT)
```

```{r, echo=F,message=F,warning=F}
RSN_namesBOLDsd <- list(
  'MotorMedial' = "Motor Medial",
  'MotorLateral' = "Motor Lateral",
  'Visual' = "Visual",
  'Auditory'= "Auditory",
  'MotorAssociation' = "Motor Association",
  'PosteriorParietal' = "Posterior Parietal",
  'Somatosensory' = "Somatosensory",
  'DorsalVisualStream' = "Dorsal Visual Stream",
  'PosteriorCingulateCortex' = "Posterior Cingulate Cortex",
  'FrontalPole'= "Frontal Pole",
  'Midbrain'= "Midbrain",
  'DorsolateralPrefrontal'= "Dorsolateral Prefrontal",
  'Cerebellum'= "Cerebellum"
)

RSN_labeller3 <- function(variable,value){
  return(RSN_names[value])
}
```


```{r, echo=F,message=F,warning=F}
BOLDSD_data <- merge(BOLD_SD, fulldata)
```

### Correlations BOLD SD and H Whole Group

Gray Matter Whole Group
```{r, echo=F,message=F,warning=F}
knitr::kable(BOLD_SD_Hdatalong %>% do(tidy(cor.test(.$HinGM, .$BOLD_SD_GM))))
```

Preterm Group GM
```{r, echo=F,message=F,warning=F}
knitr::kable(BOLD_SD_Hdatalong %>% filter(birth_age<= 37) %>% do(tidy(cor.test(.$HinGM, .$BOLD_SD_GM))))
```


White Matter Whole Group
```{r, echo=F,message=F,warning=F}
knitr::kable(BOLD_SD_Hdatalong %>% do(tidy(cor.test(.$HinWM, .$BOLD_SD_WM))))
```

Preterm Group WM
```{r, echo=F,message=F,warning=F}
knitr::kable(BOLD_SD_Hdatalong %>% filter(birth_age<= 37) %>% do(tidy(cor.test(.$HinWM, .$BOLD_SD_WM))))
```


Combined RSN Whole Group
```{r, echo=F,message=F,warning=F}
knitr::kable(BOLD_SD_Hdatalong %>% do(tidy(cor.test(.$H, .$BOLDSD))))
```

```{r, echo=F,message=F,warning=F}
BOLD_SD_Hdatalong  %>% ggplot(aes(x=H,y=BOLDSD,color=scan_age)) + geom_point(shape = 21, fill = "white") + geom_smooth(method="lm") + theme_minimalism() + xlab("H") + ylab("BOLD SD")
```
There is a weak negative correlation in the combined RSN. 

Preterm Group combined RSN
```{r, echo=F,message=F,warning=F}
knitr::kable(BOLD_SD_Hdatalong %>% filter(birth_age<= 37) %>% do(tidy(cor.test(.$H, .$BOLDSD))))
```

Preterm RSN analysis BOLD SD and PMA
```{r, echo=F,message=F,warning=F}
BOLD_SD_Hdatalong_preterm <- BOLD_SD_Hdatalong %>% filter(birth_age <= 37)
```

```{r, echo=F,message=F,warning=F}
BOLDSDh.model_preLongplot <- lmer(data = BOLD_SD_Hdatalong_preterm, BOLDSD ~ scan_age*RSN*birth_age + (1|Subject), REML = F)
BOLDSDpredictPretermLong <- ggpredict(BOLDSDh.model_preLongplot, terms = c("scan_age","RSN"), ci.lvl=0.95)
```

```{r, echo=F,message=F,warning=F}
ggplot(BOLDSDpredictPretermLong, aes(x=x, y=predicted, color=group)) +
  geom_line()+ 
  theme_minimalism() +
  #geom_ribbon(aes(ymax=conf.high, ymin=conf.low, color=group), alpha=0.3) +
  geom_line(aes(x=x, y=conf.high), linetype=2, alpha=0.3) +
  geom_line(aes(x=x, y=conf.low), linetype=2, alpha=0.3) +
    labs(x='Scan Age (weeks)', y='BOLD SD by RSN') + theme_minimalism()
```


```{r, echo=F, message=F, warning=F}
write.table(latexdf, "../latexvariables.csv", append = FALSE, sep = ",",  row.names = F)
```
````
