## Supplementary material for "Temporal Complexity of the BOLD-Signal In Preterm Versus Term Infants": All supplementary data: SupplementaryMaterial.docx

Supplementary Materials

### Tables

**Supplementary Table 1.** Preterm group H values. Estimated marginal mean H values with asymptotic confidence intervals and p-values from the VPT (n = 70) and MPT group (n = 76) comparison at 35 weeks PMA in the tissues and RSNs. NS = not significant.

| **Region** | **VPT (29 Weeks GA)** | **MPT (35 Weeks GA)** | **p-value** |
| --- | --- | --- | --- |
| Grey Matter | 0.448 [0.439, 0.456] | 0.465 [0.456, 0.473] | **0.0029** |
| White Matter | 0.422 [0.414, 0.429] | 0.448 [0.441, 0.455] | **< 0.0001** |
| Combined RSN | 0.455 [0.437, 0.454] | 0.463 [0.453, 0.472] | **0.0032** |
| Motor Medial | 0.501 [0.490, 0.513] | 0.504 [0.490, 0.518] | NS |
| Motor Lateral | 0.468 [0.456, 0.479] | 0.486 [0.473, 0.500] | NS |
| Motor Association | 0.457 [0.457, 0.484] | 0.474 [0.461, 0.488] | NS |
| Somatosensory | 0.455 [0.443, 0.466] | 0.468 [0.454, 0.481] | NS |
| Auditory | 0.454 [0.442, 0.465] | 0.470 [0.457, 0.484] | NS |
| Visual | 0.444 [0.432, 0.456] | 0.453 [0.439, 0.466] | NS |
| Posterior Parietal | 0.477 [0.465, 0.488] | 0.470 [0.456, 0.484] | NS |
| Posterior Cingulate Cortex | 0.441 [0.429, 0.452] | 0.457 [0.443, 0.470] | NS |
| Dorsal Visual Stream | 0.437 [0.426, 0.449] | 0.452 [0.439, 0.466] | NS |
| Frontal Pole | 0.446 [0.435, 0.458] | 0.474 [0.461, 0.488] | **0.0061** |
| Dorsolateral Prefrontal | 0.428 [0.417, 0.440] | 0.455 [0.442, 0.469] | **0.010** |
| Midbrain | 0.449 [0.438, 0.461] | 0.468 [0.454, 0.481] | NS |
| Cerebellum | 0.333 [0.321, 0.344] | 0.388 [0.374, 0.402] | **< 0.0001** |

**Supplementary Table 2.** Term age H values by group. Estimated marginal mean H values with asymptotic confidence intervals and p-values in the VPT, MPT and THC groups compared at 41 weeks PMA in the tissues and RSNs. NS = not significant.

| **Region** | **VPT (29 Weeks GA)** | **MPT (35 Weeks GA)** | **THC (41 Weeks GA)** | **p-value** |
| --- | --- | --- | --- | --- |
| Grey Matter | 0.547 [0.541, 0.554] | 0.553 [0.550, 0.556] | 0.559 [0.556, 0.562] | **0.0081** |
| White Matter | 0.443 [0.440, 0.447] | 0.474 [0.472, 0.476] | 0.505 [0.503, 0.506] | **<0.0001** |
| Combined RSN | 0.519 [0.502, 0.535] | 0.545 [0.537, 0.553] | 0.571 [0.563, 0.578] | **<0.0001** |
| Motor Medial | 0.644 [0.625, 0.664] | 0.658 [0.649, 0.668] | 0.673 [0.663, 0.682] | NA |
| Motor Lateral | 0.570 [0.551, 0.590] | 0.614 [0.605, 0.624] | 0.658 [0.649, 0.667] | **< 0.0001** |
| Motor Association | 0.522 [0.502, 0.542] | 0.552 [0.542, 0.561] | 0.582 [0.573, 0.591] | **< 0.0001** |
| Somatosensory | 0.538 [0.518, 0.557] | 0.564 [0.555, 0.574] | 0.590 [0.581, 0.600] | **0.0004** |
| Auditory | 0.537 [0.517, 0.556] | 0.575 [0.566, 0.585] | 0.614 [0.604, 0.623] | **< 0.0001** |
| Visual | 0.545 [0.525, 0.565] | 0.571 [0.561, 0.580] | 0.596 [0.587, 0.605] | **0.0005** |
| Posterior Parietal | 0.582 [0.562, 0.601] | 0.592 [0.582, 0.601] | 0.602 [0.593, 0.611] | NA |
| Posterior Cingulate Cortex | 0.496 [0.476, 0.515] | 0.510 [0.500, 0.519] | 0.524 [0.515, 0.533] | NA |
| Dorsal Visual Stream | 0.500 [0.480, 0.519] | 0.524 [0.514, 0.533] | 0.548 [0.539, 0.557] | **0.0012** |
| Frontal Pole | 0.484 [0.464, 0.504] | 0.513 [0.504, 0.523] | 0.542 [0.533, 0.551] | **< 0.0001** |
| Dorsolateral Prefrontal | 0.469 [0.450, 0.489] | 0.499 [0.489, 0.508] | 0.528 [0.519, 0.537] | **< 0.0001** |
| Midbrain | 0.486 [0.466, 0.505] | 0.487 [0.478, 0.497] | 0.489 [0.479, 0.498] | NA |
| Cerebellum | 0.356 [0.336, 0.376] | 0.417 [0.408, 0.427] | 0.479 [0.469, 0.488] | **< 0.0001** |

**Supplementary Table 3.** R and p-values of FA and RD correlations with H.

| **Region** | **FA** | | **RD** | |
| --- | --- | --- | --- | --- |
|  | **R value** | **p-value** | **R value** | **p-value** |
| Total WM | 0.418 | **< 0.0001** | -0.348 | **< 0.0001** |
| Motor Lateral | 0.606 | **< 0.0001** | 0.475 | **< 0.0001** |
| Somatosensory | 0.589 | **< 0.0001** | -0.499 | **< 0.0001** |
| Motor Medial | 0.552 | **< 0.0001** | -0.565 | **< 0.0001** |
| Posterior Parietal | 0.523 | **< 0.0001** | -0.322 | **< 0.0001** |
| Auditory | 0.507 | **< 0.0001** | -0.211 | **0.001** |
| Motor Association | 0.470 | **< 0.0001** | -0.338 | **< 0.0001** |
| Dorsal Visual Stream | 0.461 | **< 0.0001** | -0.151 | **0.020** |
| Visual | 0.411 | **< 0.0001** | -0.259 | **< 0.0001** |
| Dorsolateral Prefrontal | 0.425 | **< 0.0001** | -0.131 | **0.044** |
| Frontal Pole | 0.370 | **< 0.0001** | -0.008 | NS |
| Posterior Cingulate Cortex | 0.352 | **< 0.0001** | -0.300 | **< 0.0001** |

### Figures

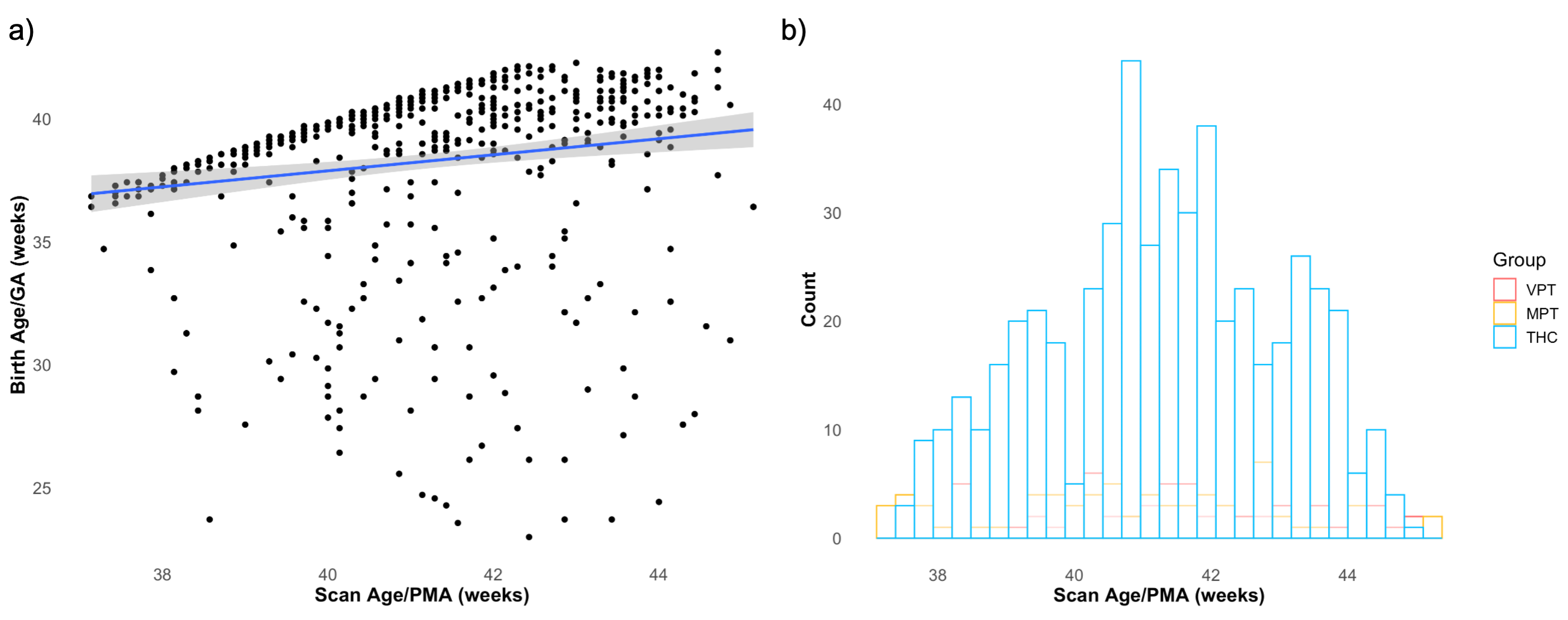
**Supplementary Figure 1**. Birth age and scan age relationships. a) Scatter plot between the two variables, r = 0.15. Birth age and scan age are not collinear. b) Histogram of the infant group at term age. An even distribution suggests we are not extrapolating our data for analysis at 41 weeks PMA.

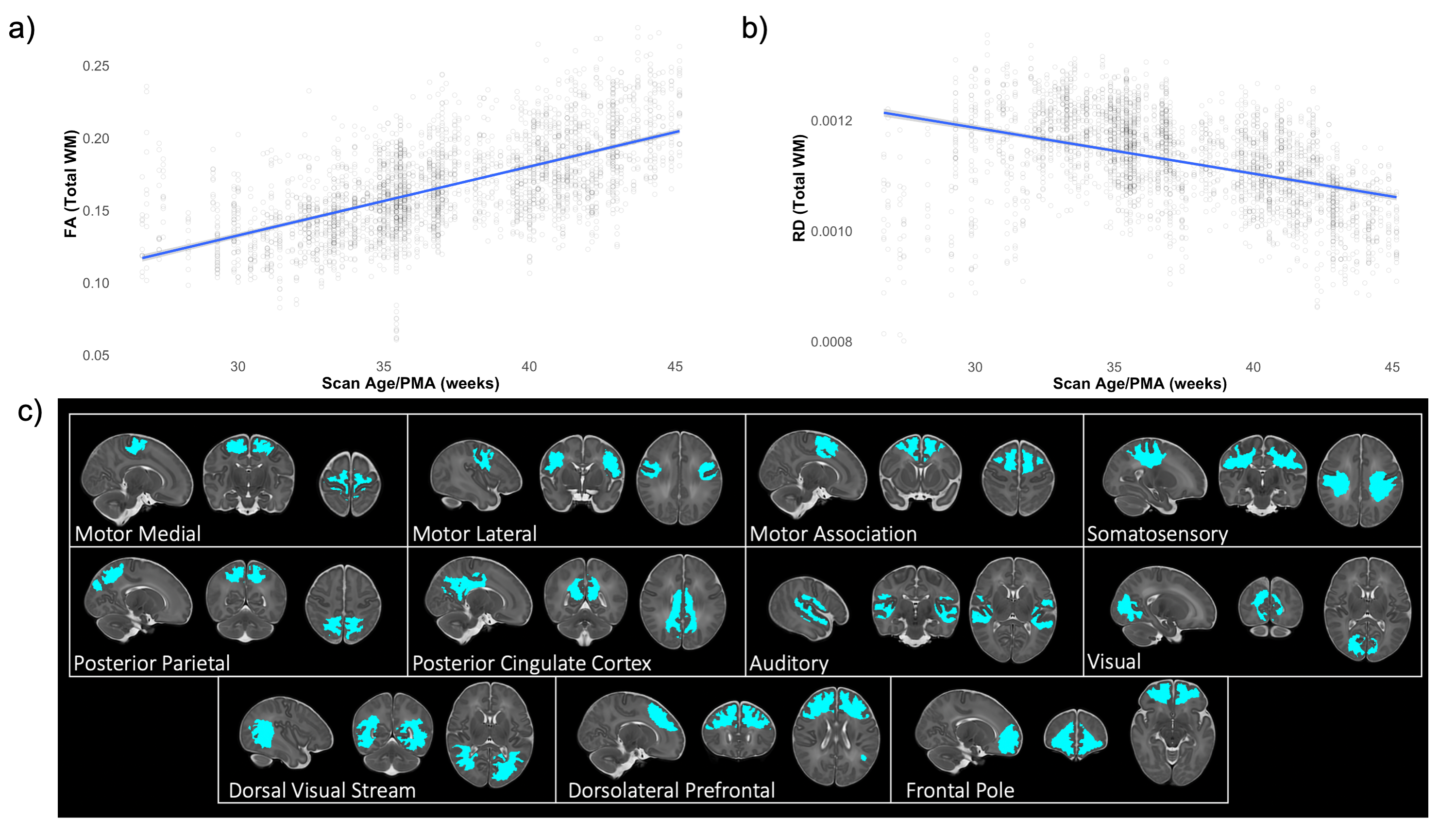

**Supplementary Figure 2.** DTI scalar plots and WM regions in the preterm group. a) FA plot with scan age (PMA) in the total white matter. b) RD plot with scan age. c) White matter regions analyzed using RSNs are regions of interests. FA = fractional anisotropy, RD = radial diffusivity, PMA = postmenstrual age, RSNs = resting state networks.

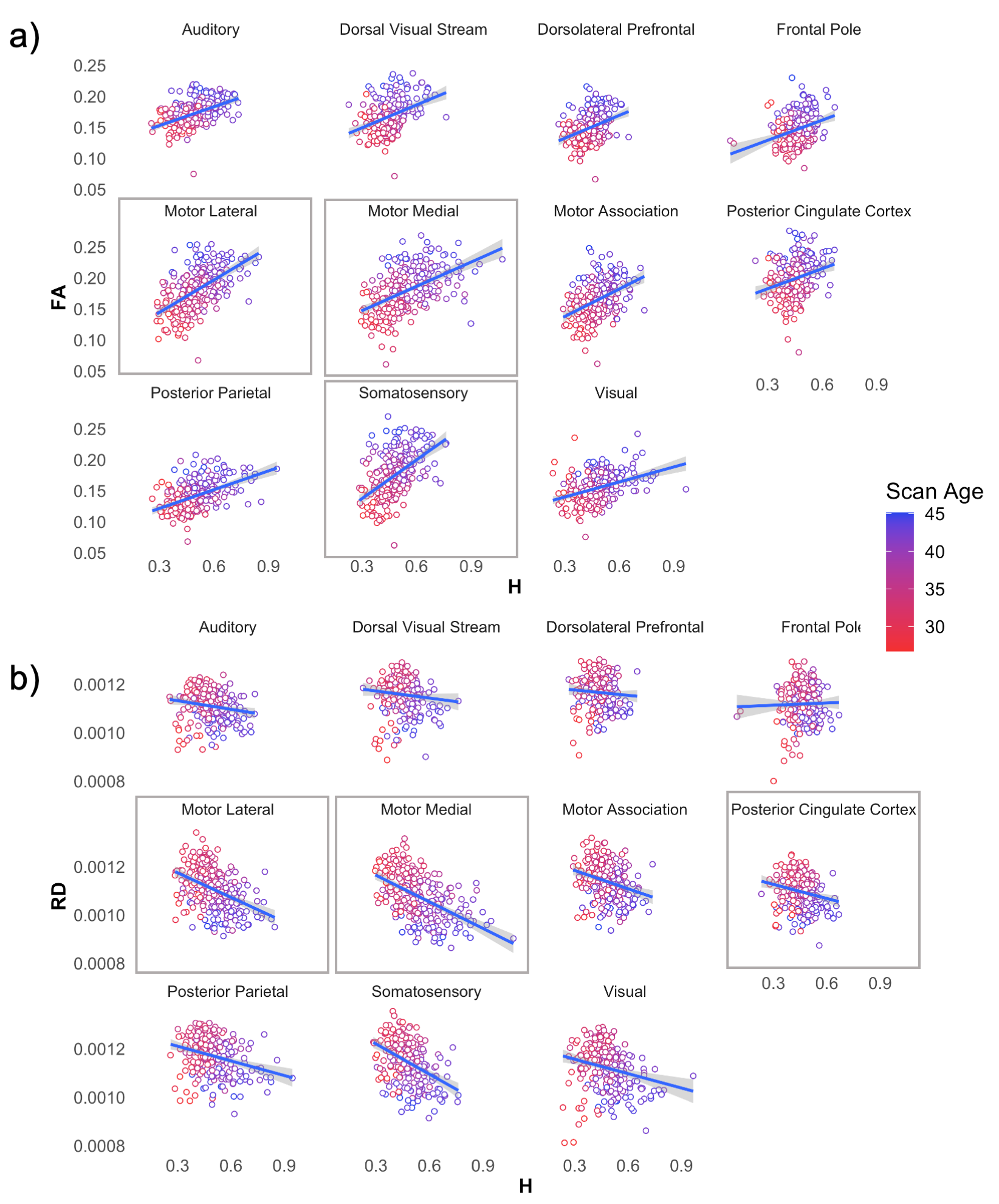

**Supplementary Figure 3.** Individual WM region correlations. a) FA and H correlations in the individual white matter regions. The motor lateral, somatosensory, and motor medial regions had the strongest correlations. b) RD and H correlations in the individual white matter regions. The motor medial, motor lateral and posterior cingulate regions had the strongest correlations. Pearson correlation and p-values can be found in Supplementary Table 3.
