## Supplementary figures and images for "Temporal Complexity of the BOLD-Signal In Preterm Versus Term Infants"

### ALLRSNbar.png

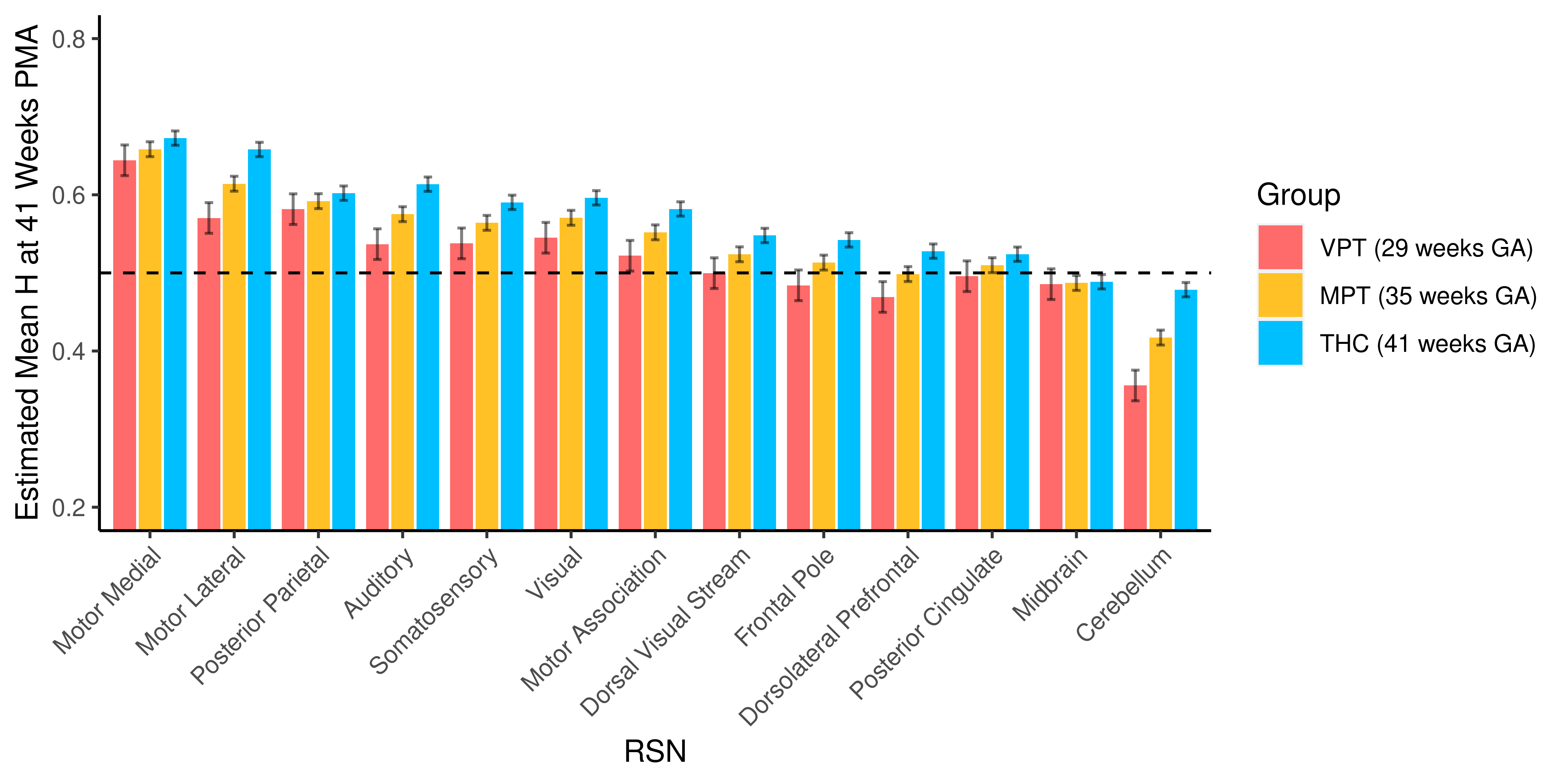

### ALLtissuebar.png

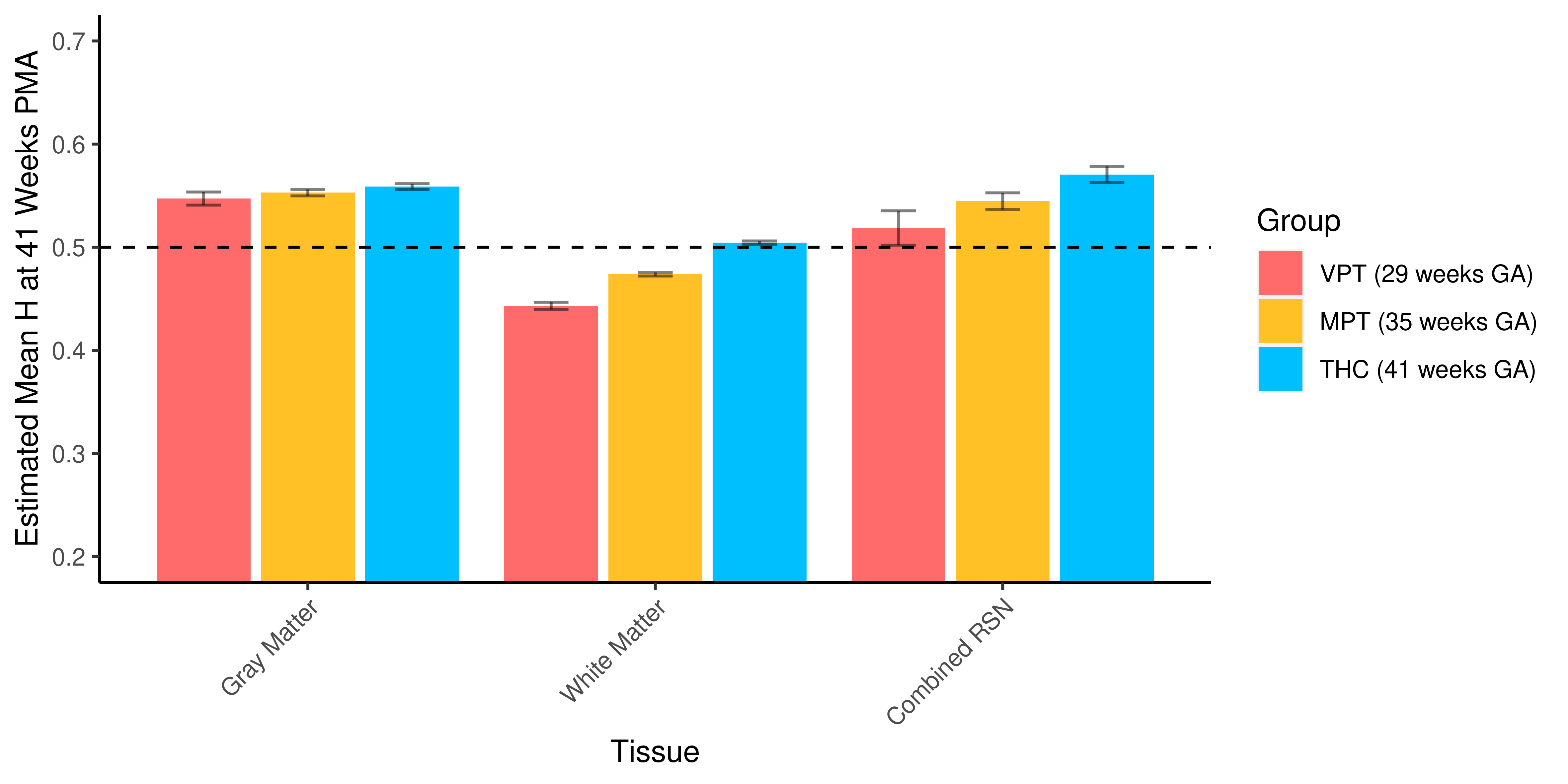

### FAvsH.png

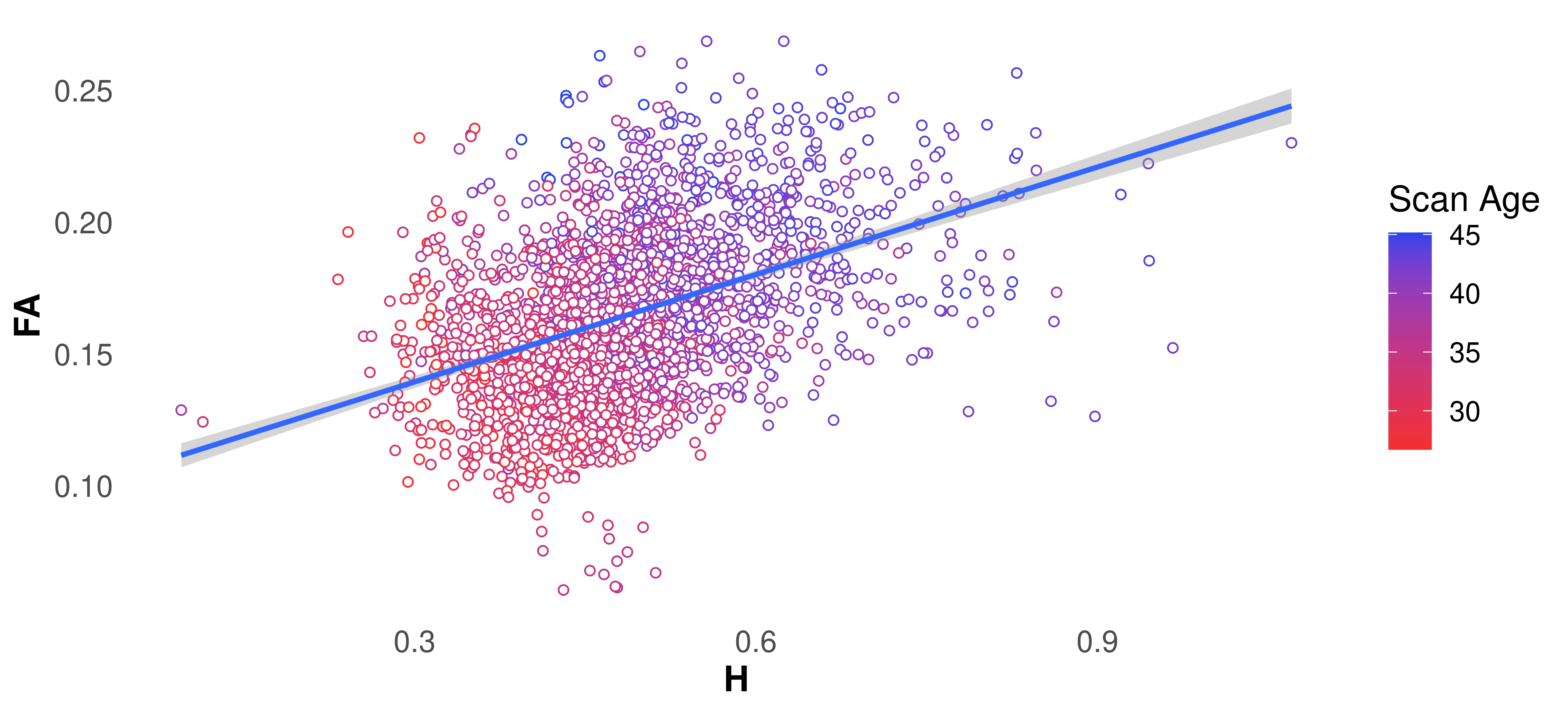

### FAvsH_RSN.png

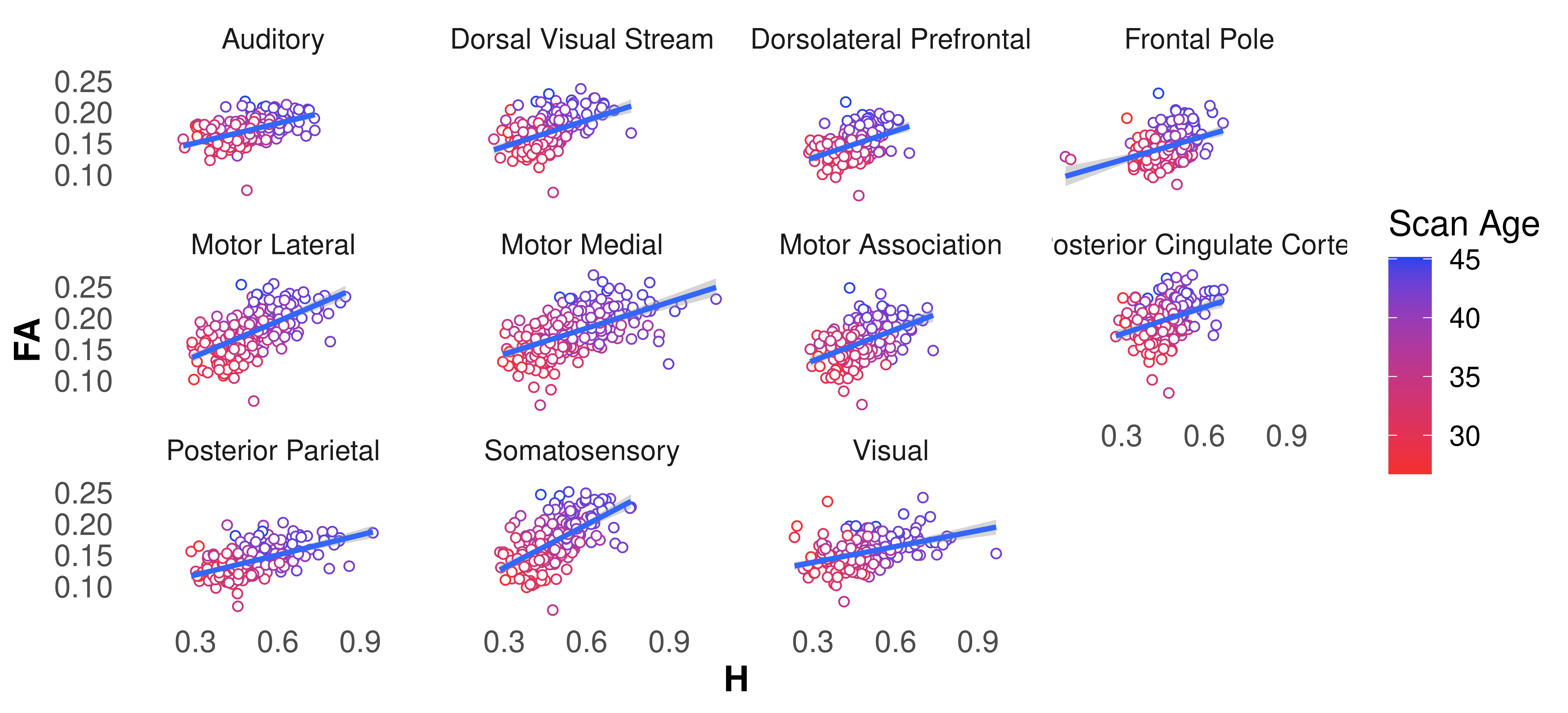

### GAvsPMA_PMAgt37.png

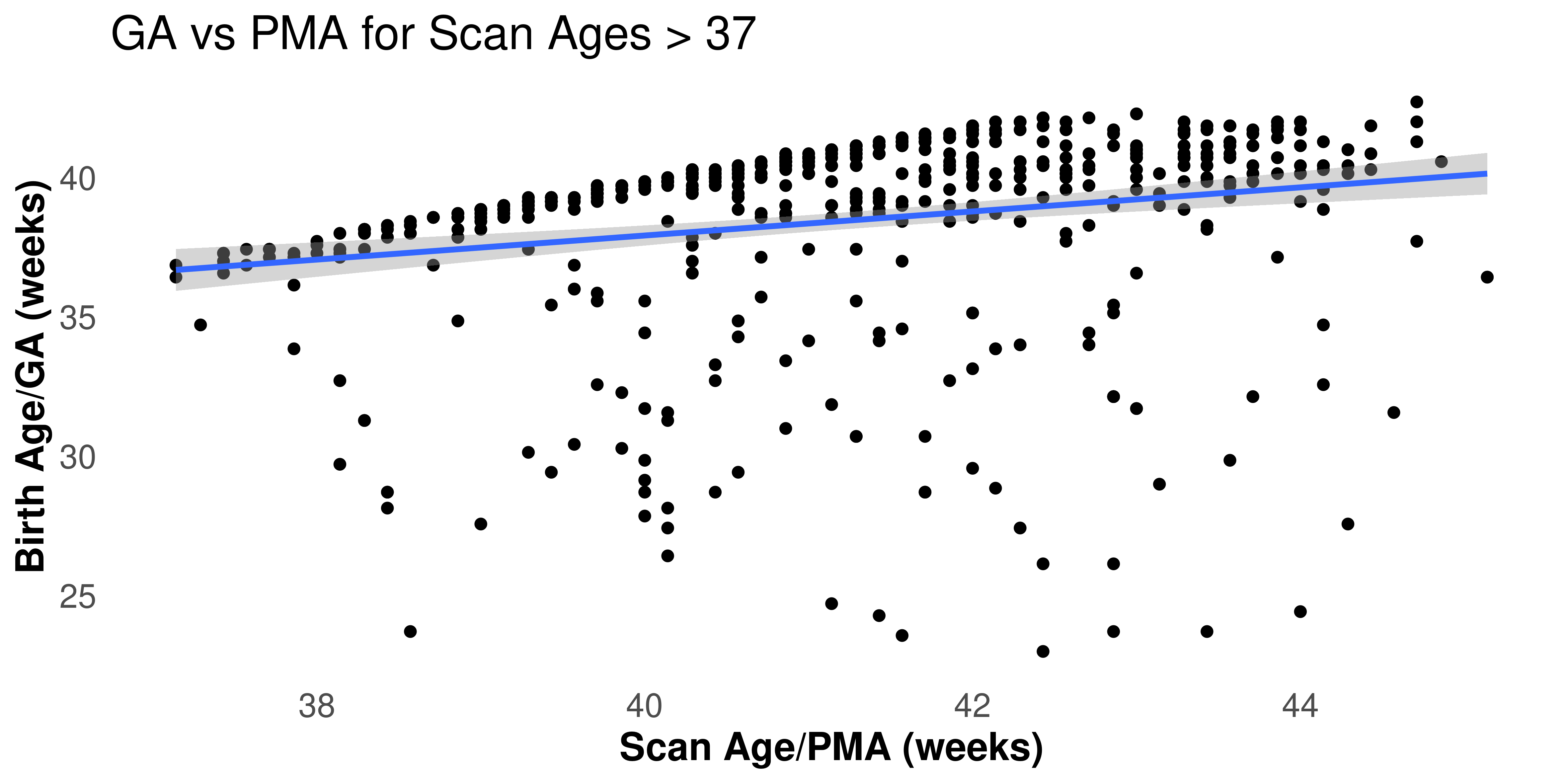

### HGMvsGA.png

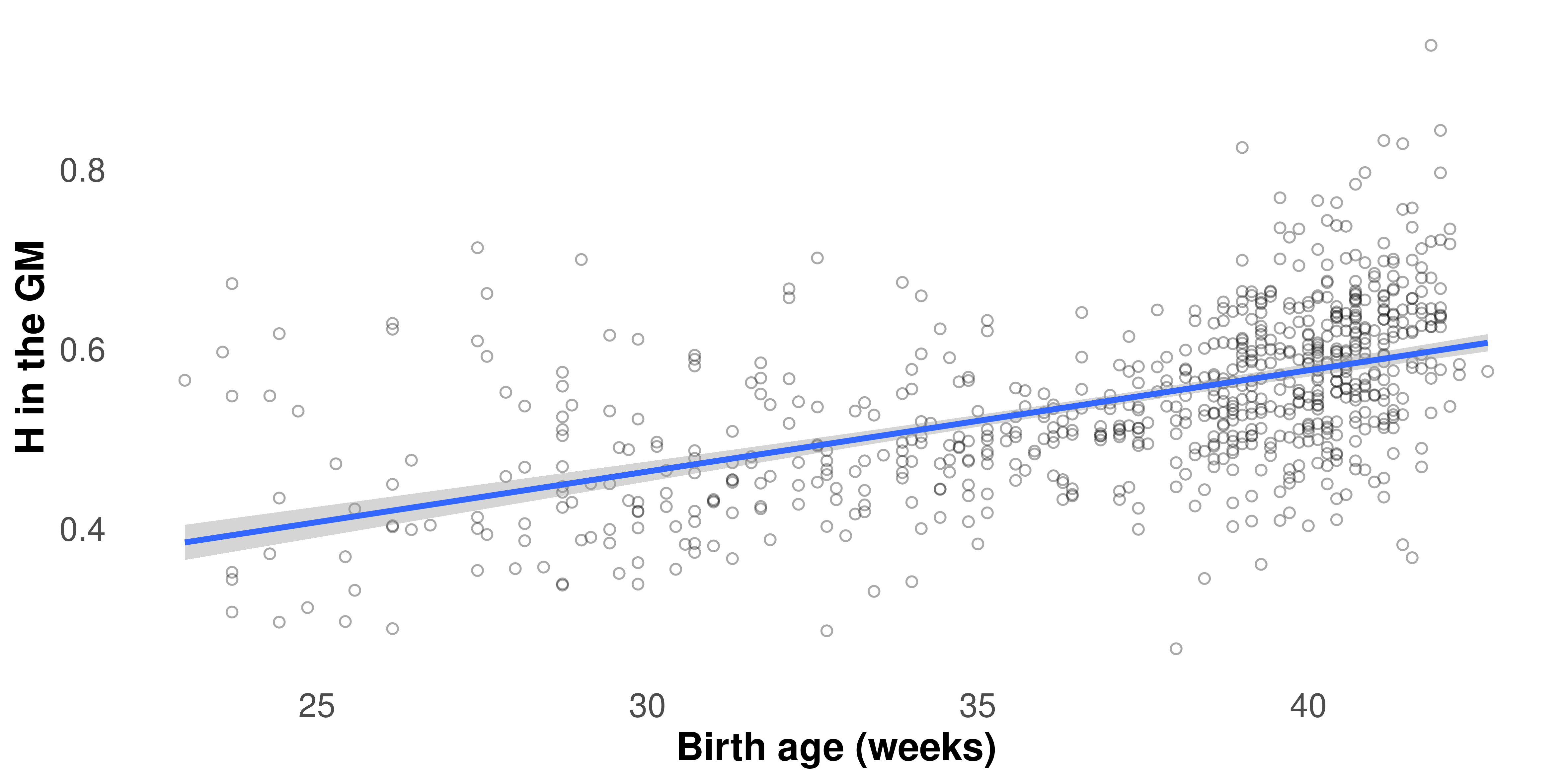

### HGMvsPMA.png

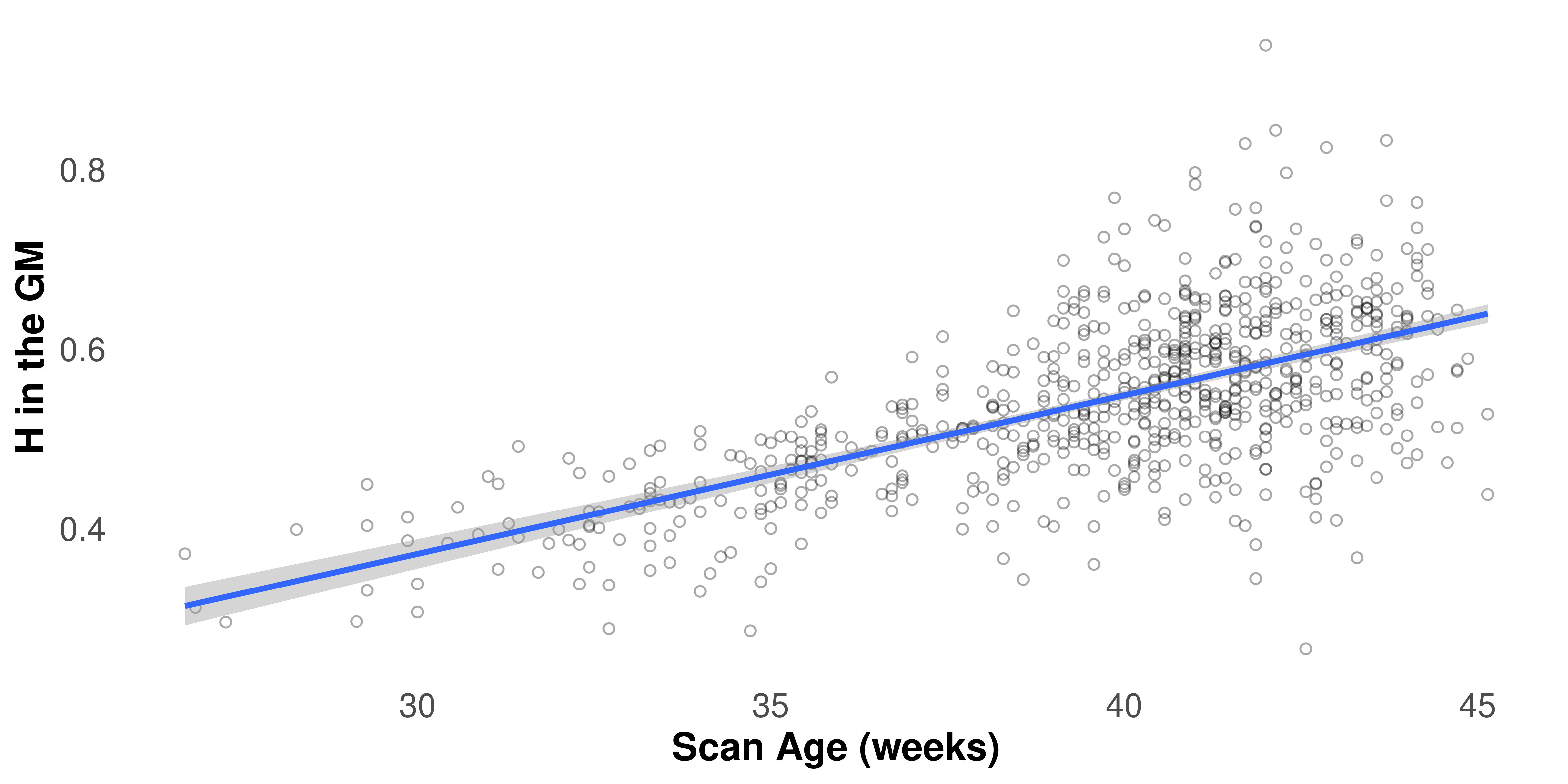

### PMAHist_PMAgt37.png

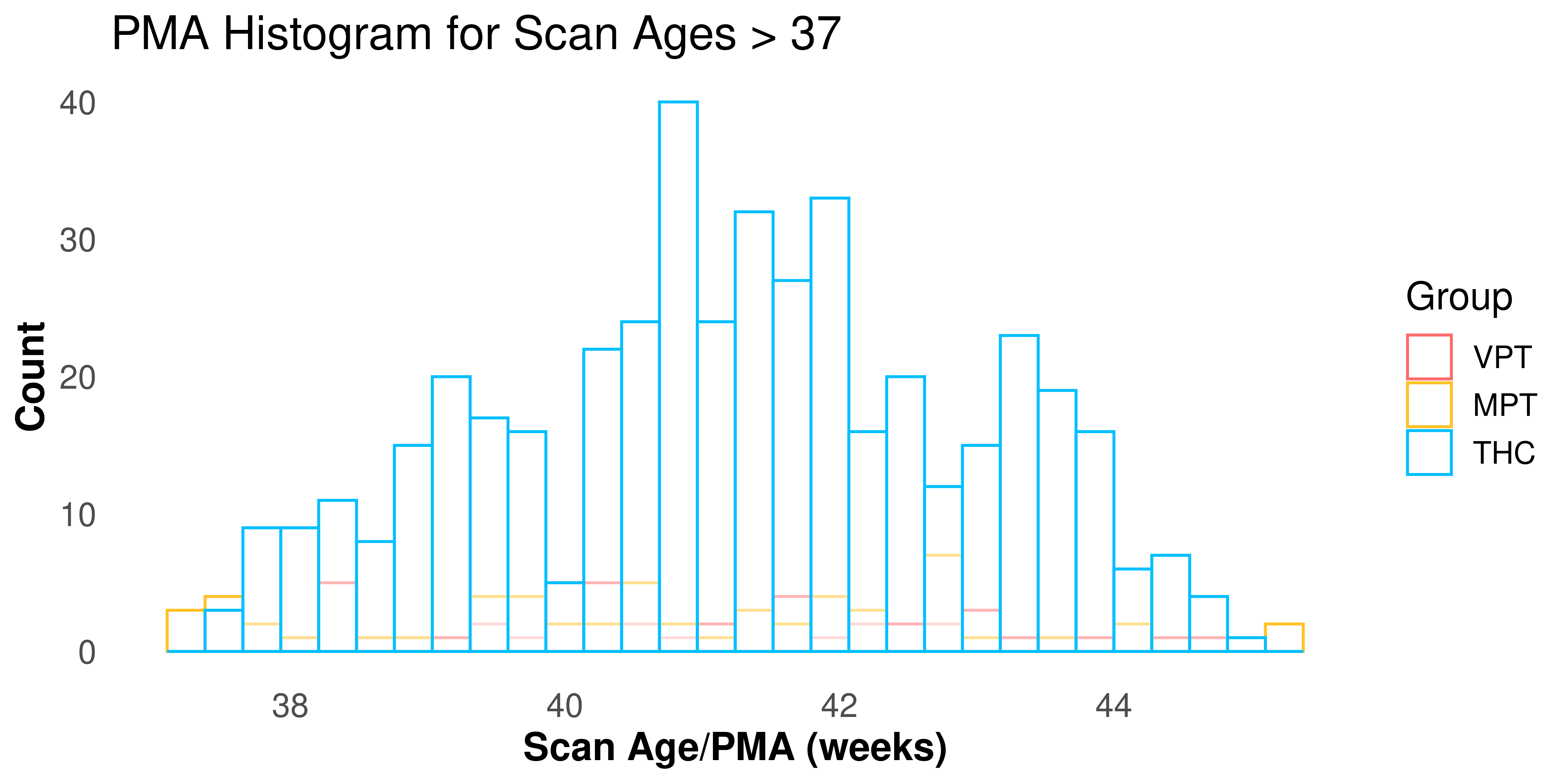

### RDvsH.png

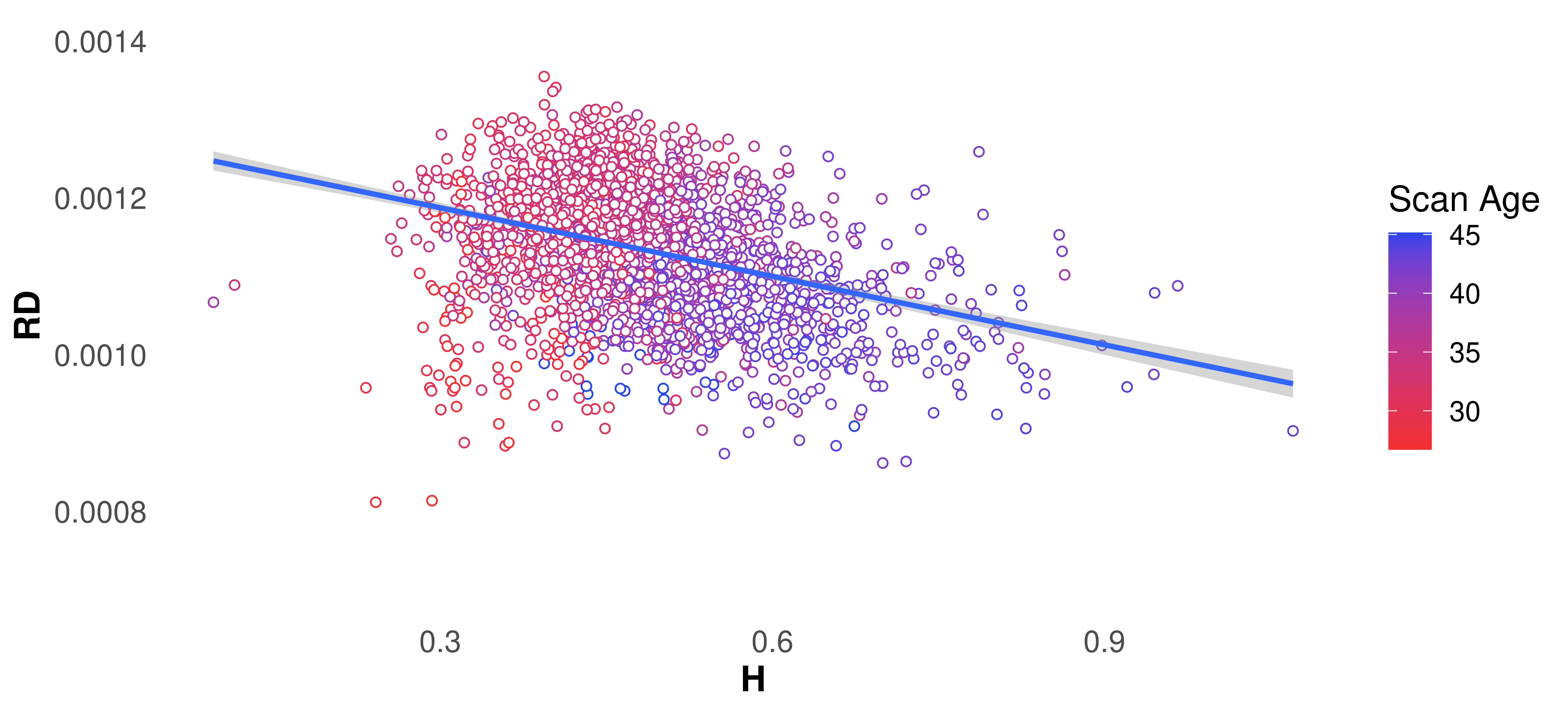

### RDvsH_RSN.png

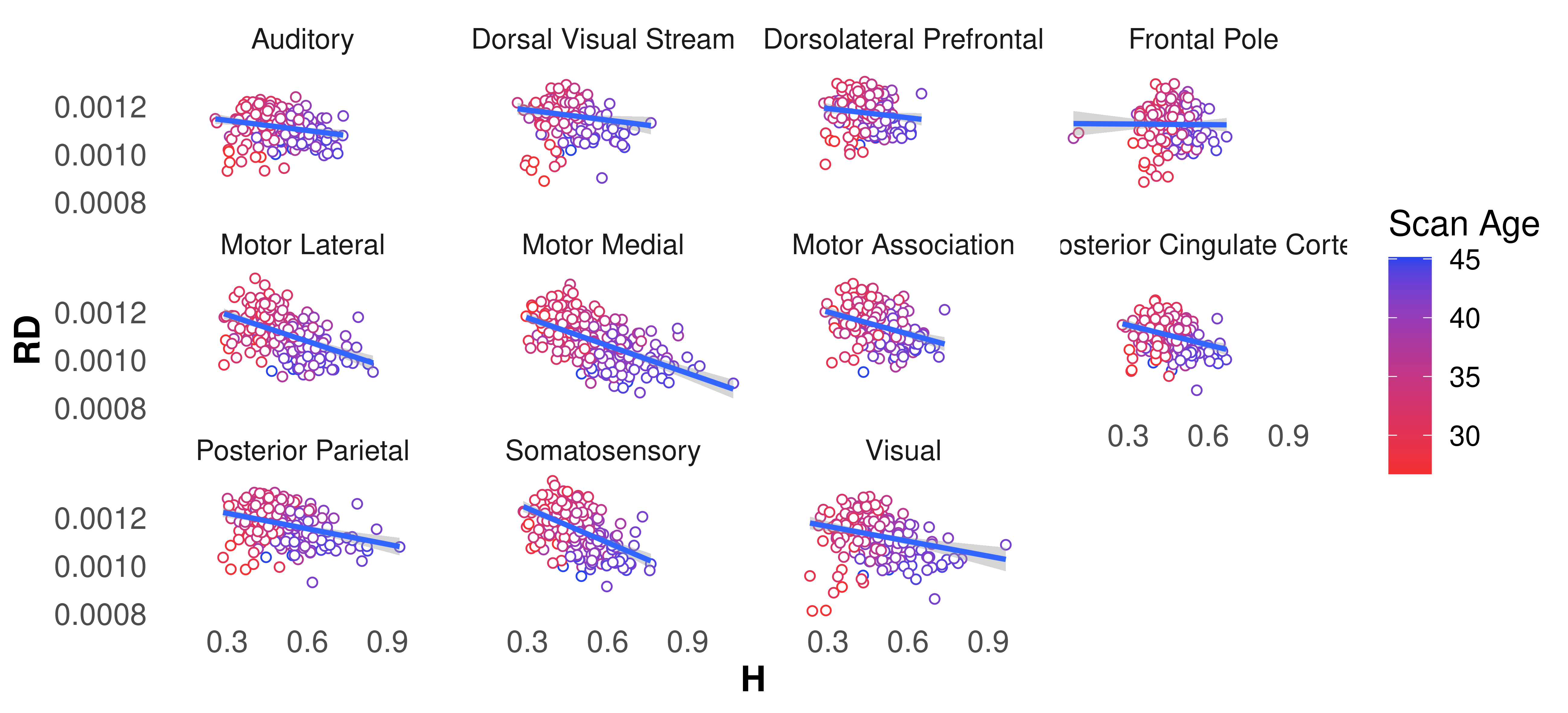

### VPTMPTRSNbar.png

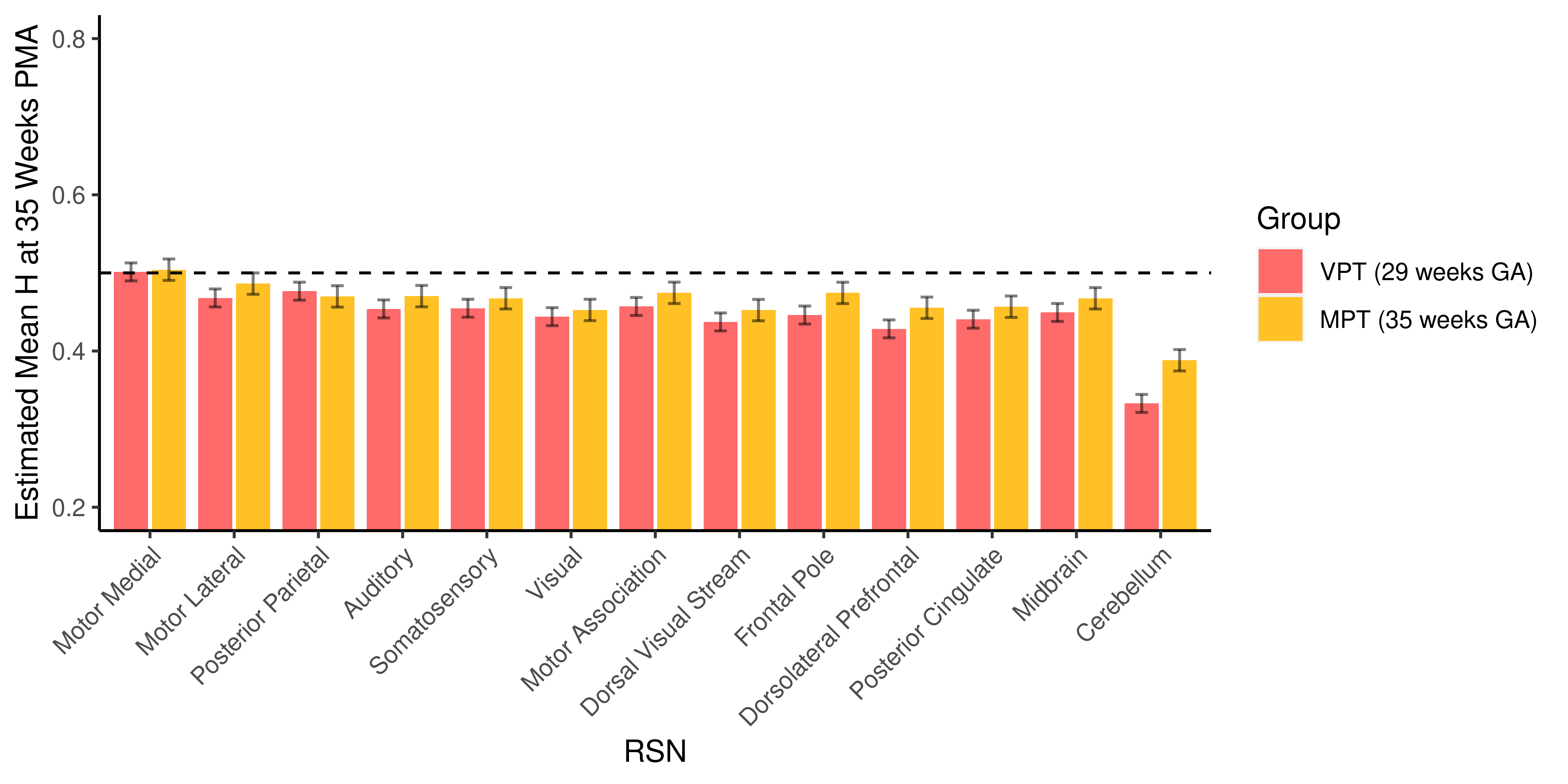

### VPTMPTRSNlopes.png

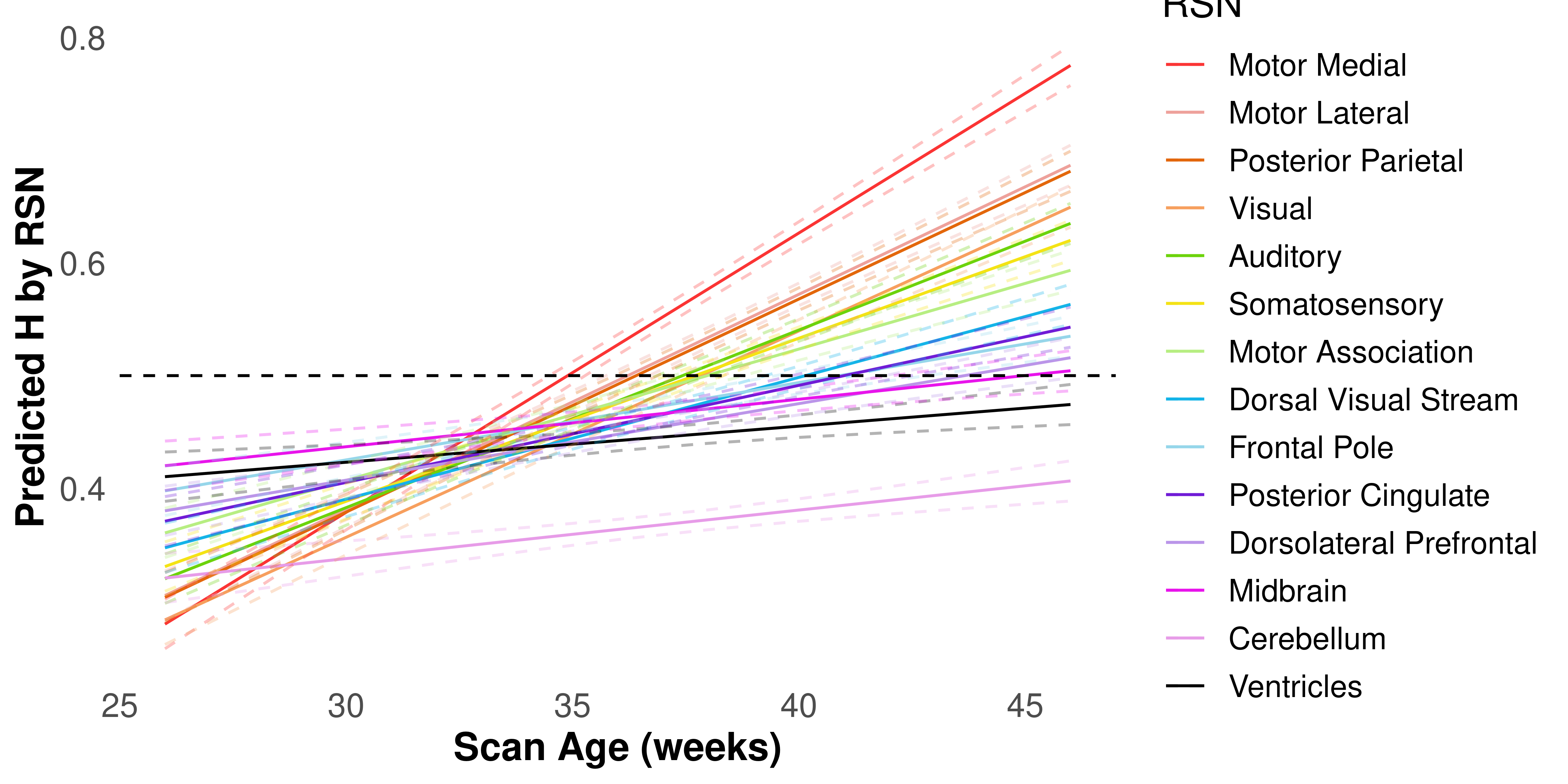

### VPTMPTtissuebar.png

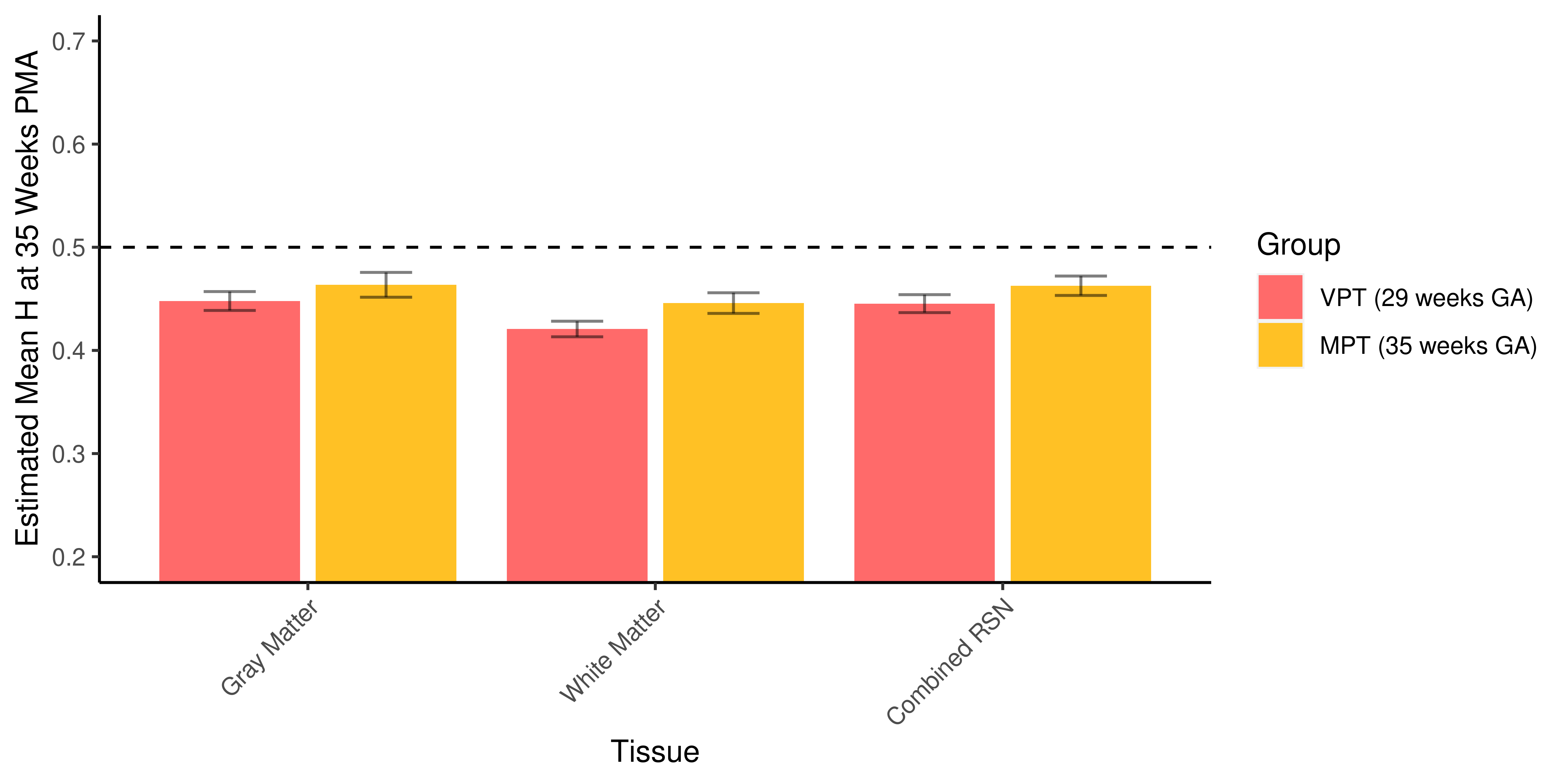

### VPTMPTtissueslopes.png

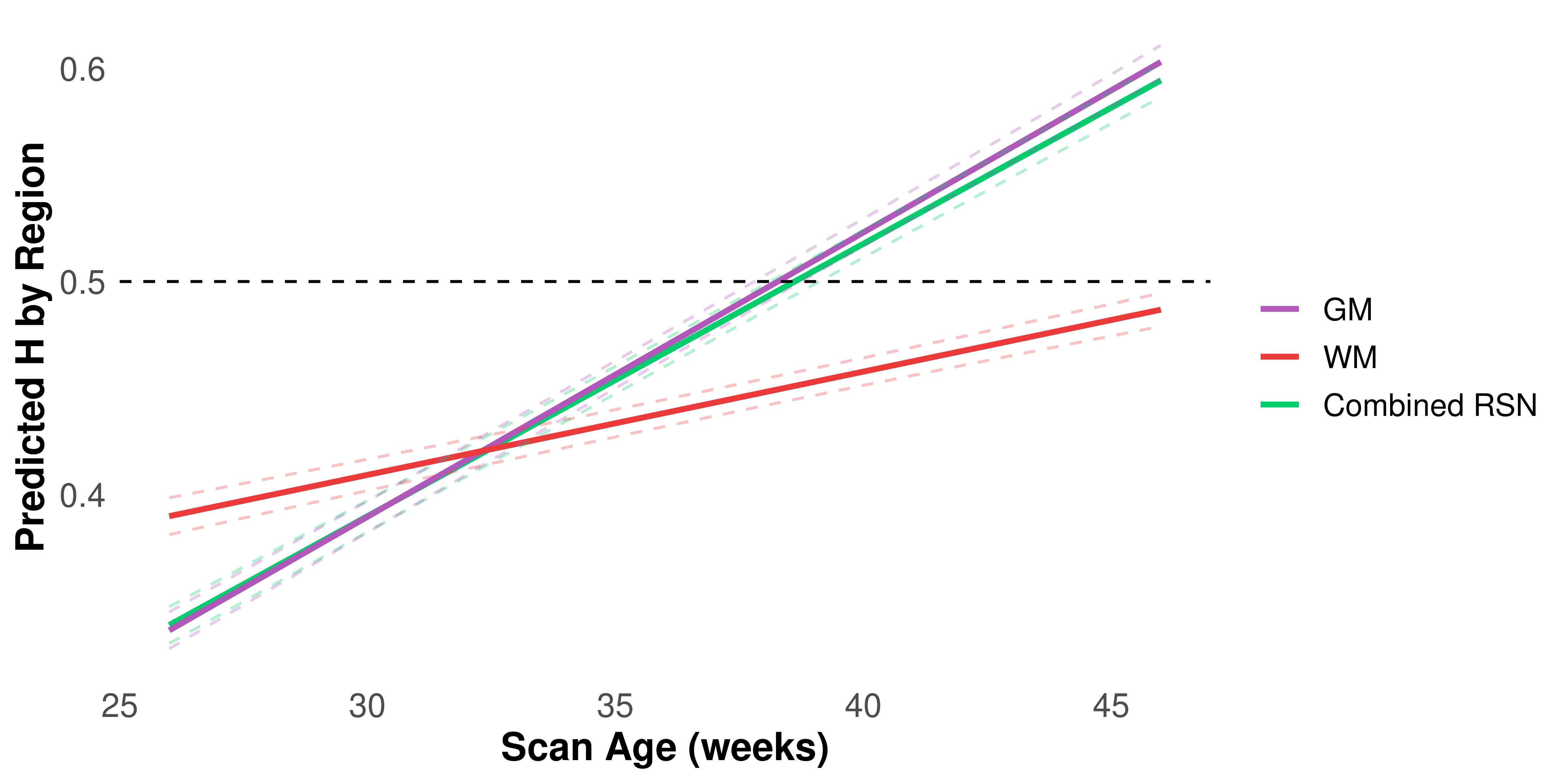
